# The origins and molecular evolution of SARS-CoV-2 lineage B.1.1.7 in the UK

**DOI:** 10.1101/2022.03.08.481609

**Authors:** Verity Hill, Louis Du Plessis, Thomas P. Peacock, Dinesh Aggarwal, Rachel Colquhoun, Alesandro M. Carabelli, Nicholas Ellaby, Eileen Gallagher, Natalie Groves, Ben Jackson, JT McCrone, Áine O’Toole, Anna Price, Theo Sanderson, Emily Scher, Joel Southgate, Erik Volz, The COVID-19 genomics UK (COG-UK) consortium, Wendy S. Barclay, Jeffrey C. Barrett, Meera Chand, Thomas Connor, Ian Goodfellow, Ravindra K. Gupta, Ewan M. Harrison, Nicholas Loman, Richard Myers, David L Robertson, Oliver G Pybus, Andrew Rambaut

**Affiliations:** Institute of Evolutionary Biology, University of Edinburgh, Edinburgh, UK; Department of Zoology, University of Oxford, Oxford, UK; Department of Biosystems Science and Engineering, ETH Zürich, Switzerland; Department of Infectious Disease, Imperial College London, London, W2 1PG, UK; Wellcome Sanger Institute, Wellcome Genome Campus, Hinxton, UK; UK Health Security Agency, London, UK; University of Cambridge, Department of Medicine, Cambridge, UK; Cambridge University Hospital NHS Foundation Trust, Cambridge, UK; School of Biosciences, The Sir Martin Evans Building, Cardiff University, Cardiff, UK; The Francis Crick Institute, 1 Midland Rd, London NW1 1AT, UK; MRC Centre for Global Infectious Disease Analysis and Department of Infectious Disease Epidemiology, Imperial College London; Guy’s and St. Thomas’ Hospital NHS Trust, London, UK; Pathogen Genomics Unit, Public Health Wales NHS Trust, Cardiff, UK; Department of Pathology, University of Cambridge, Cambridge, UK; Africa Health Research Institute, Durban, South Africa; Department of Public Health and Primary Care, University of Cambridge, Cambridge, UK; Institute of Microbiology and Infection, University of Birmingham, Birmingham, UK; MRC-University of Glasgow Centre for Virus Research, Glasgow, G61 1QH, Scotland, UK; Department of Pathobiology and Population Science, The Royal Veterinary College, London, UK

## Abstract

The first SARS-CoV-2 variant of concern (VOC) to be designated was lineage B.1.1.7, later labelled by the World Health Organisation (WHO) as Alpha. Originating in early Autumn but discovered in December 2020, it spread rapidly and caused large waves of infections worldwide. The Alpha variant is notable for being defined by a long ancestral phylogenetic branch with an increased evolutionary rate, along which only two sequences have been sampled. Alpha genomes comprise a well-supported monophyletic clade within which the evolutionary rate is more typical of SARS-CoV-2. The Alpha epidemic continued to grow despite the continued restrictions on social mixing across the UK, and the imposition of new restrictions, in particular the English national lockdown in November 2020. While these interventions succeeded in reducing the absolute number of cases, the impact of these non-pharmaceutical interventions was predominantly to drive the decline of the SARS-CoV-2 lineages which preceded Alpha. We investigate the only two sampled sequences that fall on the branch ancestral to Alpha. We find that one is likely to be a true intermediate sequence, providing information about the order of mutational events that led to Alpha. We explore alternate hypotheses that can explain how Alpha acquired a large number of mutations yet remained largely unobserved in a region of high genomic surveillance: an under-sampled geographical location, a non-human animal population, or a chronically-infected individual. We conclude that the last hypothesis provides the best explanation of the observed behaviour and dynamics of the variant, although we find that the individual need not be immunocompromised, as persistently-infected immunocompetent hosts also display a higher within-host rate of evolution. Finally, we compare the ancestral branches and mutation profiles of other VOCs to each other, and identify that Delta appears to be an outlier both in terms of the genomic locations of its defining mutations, and its lack of rapid evolutionary rate on the ancestral branch. As new variants, such as Omicron, continue to evolve (potentially through similar mechanisms) it remains important to investigate the origins of other variants to identify ways to potentially disrupt their evolution and emergence.

## Introduction

In early December 2020, one of the four UK public health agencies (Public Health England, PHE, now known as UK Health Security Agency, UKHSA) began tracking and investigating a rapid increase in COVID-19 incidence in South East England, centred on Kent and East London. Numbers of new cases had grown more rapidly than expected over the previous four weeks, despite an elevated level of non-pharmaceutical interventions (NPIs) in the region, and increased incidence had begun to be observed in other locations in the UK, indicating further spread (Public Health England 2020). A corresponding genomic cluster was detected separately within the COG-UK SARS-CoV-2 genomic surveillance dataset, and the genome sequences carried a substantially larger than usual number of genetic changes (Rambaut et al. 2020a). At a routine PHE meeting on the 8th December 2020, the link between the genomic cluster and the Kent epidemiological situation was made and investigations were rapidly initiated to characterise the mutations and to estimate the growth rate of the cluster. Evidence accumulated that this cluster was growing rapidly and had expanded throughout November, during a national lockdown in England. The cluster was designated B.1.1.7 under the Pango lineage naming system (Rambaut et al. 2020b), and was later labelled as variant of concern (VOC) Alpha under the World Health Organisation (WHO) variant nomenclature (Konings et al. 2021).

Since its discovery, substantial analytical effort has been put into teasing apart the contributions of human behavioural factors and true virological effects on the rapid growth of the lineage (Kraemer et al. 2021). It is now clear that Alpha was associated with a higher transmission rate than the background D614G lineages that dominated in the UK at the time (Davies, Abbott, et al. 2021; Leung et al. 2021; Volz et al. 2021) as well as a higher case fatality rate (Davies, Jarvis, et al. 2021).

Alpha contains 14 lineage-specific amino acid replacements and three deletions compared to contemporaneous lineages (Rambaut et al. 2020a), which was unprecedented in the global virus genomic dataset for the COVID-19 pandemic at the time of its emergence (Table S1). This mutational constellation included several mutations that have arisen independently in other VOCs. For example, N501Y in the Spike protein is also found in Beta (B.1.351), Gamma (P.1) and Omicron (B.1.1.529 descendants) and is a key contact residue in the receptor binding domain (RBD); experimental data has determined that it increases binding affinity to human and murine ACE2 (Starr et al. 2020; Tian et al. 2021) and it has been associated with increased infectivity and virulence in a mouse model (Gu et al. 2020). N501Y alone has also been associated with higher infectivity and transmissibility of SARS-CoV-2 (Liu et al. 2021). There are also two deletions of interest in Alpha’s Spike gene: six base pairs at position 21765 (amino acid positions 69-70) and three base pairs at position 21991 (amino acid position 144). Both have previously arisen in chronically-infected individuals (Avanzato et al. 2020; Choi et al. 2020; Kemp et al. 2021; McCarthy et al. 2021). The former was also associated with the rapid outbreak in mink in Denmark (Oude Munnink et al. 2021), and has been shown *in vitro* to increase infectivity (Meng et al. 2021); and the latter has been shown to prevent monoclonal antibody and, to a lesser extent, convalescent antisera binding (Andreano et al. 2020; Collier et al. 2021), as well as exhibiting decreased neutralisation efficiency (Weigang et al. 2021). Further, Alpha contains a 9 base pair deletion in NSP6, also found in the VOCs Beta, Gamma and Omicron, which is on the outside of the autophagy vesicle, theoretically limiting autophagosome expansion (Benvenuto et al. 2020). There is also a mutation in the accessory protein ORF8, which truncates the protein from 121 to only 27 amino acids in length, likely resulting in loss of function and allowing further downstream mutations to accrue. Subsequent work has found that the ORF8 deletion has only a modest deleterious effect on virus replication in human primary airway cells compared to viruses without the deletion (Gamage et al. 2020). However, these mutations and deletions have arisen multiple times during the pandemic, and are not always associated with rapid growth or VOCs. This suggests that there are epistatic effects between many of the mutations present in Alpha that together lead to its increased fitness, as well as some hitchhiking mutations which are selectively neutral.

While this constellation of mutations appears to have arisen in one evolutionary leap, two sequences have been identified in the COG-UK genomic surveillance dataset that contain some, but not all, of the Alpha-defining mutations, hence they may represent intermediate steps in the evolution of the Alpha lineage. These sequences could provide clues to the evolutionary processes underlying the evolution and emergence of VOCs and information on the timings of mutational events.

The existence of only two potential intermediate samples must be explained. Due to the high level of SARS-CoV-2 genomic surveillance in the UK, it is unlikely that Alpha would transmit and evolve in a conventional way (i.e. transmitting between individuals in the general UK population) without numerous intermediate genomes being sampled. Instead, the lineage may have been evolving in an unsampled population before being detected in the general population in the UK, with the potential intermediates indicating early introductions from this population into the general population. We propose three possible alternatives for the nature of this unsampled population: first, Alpha may have evolved conventionally in a location with little or no virus genomic surveillance before being introduced into the UK in Kent; second, it may have evolved in a non-human animal population before a zoonotic event re-introduced it into the human population in the UK; or finally, it may have evolved in a single or small number of chronically-infected individuals, which were not sampled, before a single transmission event into the general population.

The sudden appearance, in late 2021, of the VOC Omicron, designated a descendant of B.1.1.529, has renewed interest in the processes underlying the emergence of variants exhibiting major leaps in evolution: it is defined by 45 non-synonymous mutations, and exhibits increased transmissibility, increased ability to bind to ACE2 compared to Delta, and marked change in its antigenic profile enabling antibody escape from much of the current population immunity (Meng et al. 2022; Viana et al. 2022; World Health Organisation 2021). Whilst Omicron is the most extreme example to have emerged to date, here we consider the origins of B.1.1.7/Alpha and the evidence for this lineage being the result of a similar process. In this study, we explore the processes on the branch leading up to the B.1.1.7 lineage, by using a Bayesian phylogenetic analysis to provide temporal and evolutionary rate estimates; as well as examining the two sequences which appear to be evolutionary intermediates. We then conduct coalescent and birth-death analyses to explore any differences in growth rates between Alpha and background lineages. Finally, in order to identify any common patterns among VOCs, we perform similar evolutionary rate analyses on all VOCs and VOIs, and compare their mutation profiles.

## Results

### Characterising the ancestral branch of B.1.1.7

While the first sequence of B.1.1.7 was sampled on 20th September 2020 (GISAID Accession ID: EPI_ISL_601443), the lineage diverged from other concurrently circulating background lineages in the UK in late April 2020 (time of most recent common ancestor (TMRCA) 2020-04-24, 95% highest posterior density (HPD) 2020-03-26 to 2020-05-24). However, it appears that rapid transmission of this lineage within the UK only began later in the year, with the TMRCA of the Alpha clade estimated at 2020-09-19 (95% HPD 2020-09-18 to 2020-09-20, Fig. 1A).

**Figure 1.**
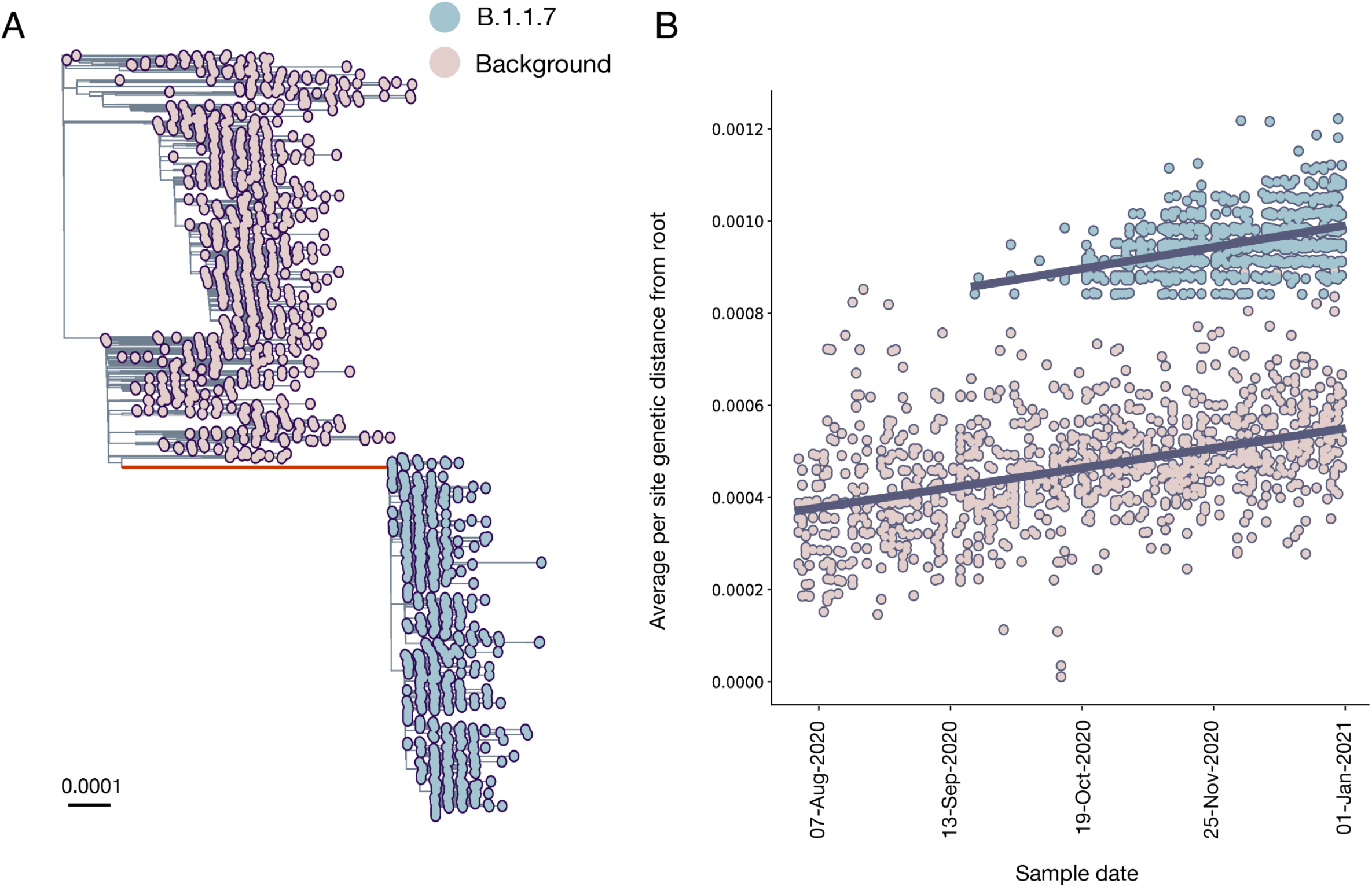
A) Maximum likelihood phylogeny showing the well-supported monophyletic clade that constitutes B.1.1.7. The ancestral branch with the higher rate of evolution is highlighted in red, and branch lengths represent substitutions/site B) Regression of root-to-tip genetic distances against sampling dates, for sequences belonging to lineage B.1.1.7 (blue) and those in its immediate outgroup in the global phylogenetic tree (pink). The regression lines are fitted to the two datasets independently. The regression gradient is an estimate of the rate of sequence evolution. These rates are 4.6×10^−4^ and 4.3×10^−4^ nucleotide changes/site/year for the B.1.1.7 and outgroup data sets, respectively

The ancestral branch leading to the B.1.1.7 lineage is exceptionally long, both in terms of time (mean=147 days, 95% HPD: 112 days to 173 days) and genetic changes: there are 23 nucleotide changes with the majority being amino acid altering (14 non-synonymous mutations and three deletions). We found that the evolutionary rate of the ancestral branch was an average of 2.3 times higher than the background rate (95% HPD = 1.4 to 3.6). There is however little evidence for an increased rate of evolution within the B.1.1.7 clade: a regression of root-to-tip of genetic distances against genome sampling date (Figure 1B) shows that the rates within the B.1.1.7 clade and the background sequences are very similar (4.6×10^−4^ and 4.3×10^−4^ nucleotide changes/site/year respectively).

Two sequences lie along the branch leading to the B.1.1.7 clade, and both contain some, but not all, of the Alpha-defining mutations. The earlier of the two (COG-UK identifier CAMC-946506, gisaid ID EPI_ISL_556680) was sampled on 15th July 2020, and the more recent genome (MILK-B154B6, gisaid ID EPI_ISL_2735517) was sampled on 23rd October 2020. If these two sequences are truly intermediate - i.e. they represent midpoints in the accrual of the 23 lineage-defining mutations for Alpha - then they may provide insight into the order of mutational accumulation during VOC evolution.

The more recent sequence, MILK-B154B6, is ambiguous at several sites, including at position 501 in Spike, with 80% of reads encoding N (asparagine, found in background lineages) and 20% Y (tyrosine, found in Alpha). These ambiguous sites imply either a coinfection of two different virus populations or laboratory contamination. If a coinfection, the sample could have been an individual who was infected by an early Alpha sequence and a background lineage (possible in late October 2020 in the South East of England, as both lineages were circulating there at the time). MILK-B154B6 contains the synonymous mutation C5986T, which is also found in 971 of the 976 of the early Alpha sequences, but in none of the 1100 background sequences used in this study. It also carries two more mutations (C15279T and C913T) that are respectively found in 974 and 970 (out of 976) sequences in the Alpha dataset, but only once in the background dataset. As this sequence contains mutations that are shared by most Alpha sequences and not found frequently in earlier clades, it suggests that its intermediate status is due to a coinfection of an Alpha sequence (which contains these mutations), and a background clade that was co-circulating, and so the consensus sequence contains only some of the Alpha-defining mutations; or cross-contamination in a laboratory handling samples from both Alpha and background lineages. MILK-B154B6 can therefore not be treated as an evolutionary intermediate.

The older sequence, CAMC-946506, contains no ambiguous sites and carries four of the Alpha-defining mutations: N501Y in Spike, the 9 base pair deletion in NSP6, as well as R52I and Q27 to stop codon mutation (Q27*) in ORF8. In the UK, prior to 1st September 2020, 37 sequences were sampled with Q27*, 5 with R52I (all in July and August, one of which also had Q27*), and none with the NSP6 deletion or N501Y (n=34,291). This makes it unlikely either that a virus containing all of these mutations existed in early 2020, prior to when Alpha diverged from the background lineages, or that this sequence has convergently acquired these mutations and has been placed erroneously in the tree. Instead, the evidence suggests that CAMC-94506 could be a true intermediate sequence, however it must be noted that it may also simply contain mutations shared by the common ancestor of the hypothesised cryptic population and CAMC-94506 (Fig. 2A).

**Figure 2.**
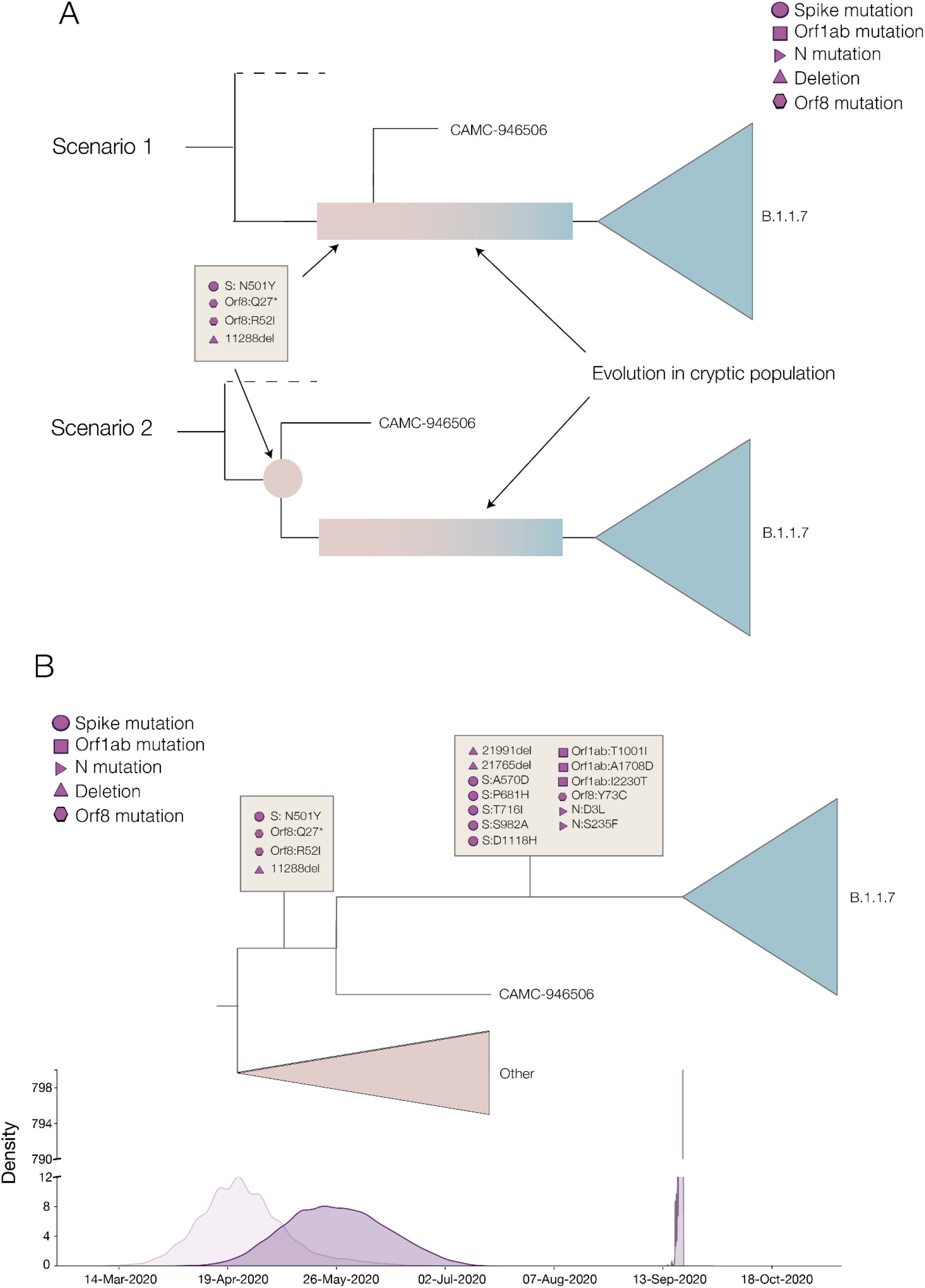
A) Two different scenarios of how the shared mutations between CAMC-946506 and the B.1.1.7 clade could have arisen. Scenario 1 shows CAMC-946506 as resulting from a transmission chain spilling over from an isolated cryptic population, such as a chronically infected individual, and the mutations arising early in the infection. Scenario 2 shows the mutations as being shared by the common ancestor of CAMC-946506 and a cryptic population. B) Schematic of the time tree showing possible timings for B.1.1.7 lineage-defining mutations. Densities of the most recent common ancestors for (respectively) the background lineages and all B.1.1.7, the intermediate sequence and B.1.1.7, and all B.1.1.7 are shown along the bottom.

This intermediate sequence provides evidence that the Spike mutation N501Y, the two ORF8 mutations Q27* and R52I, and the 9 base pair deletion in NSP6 all evolved early in the evolutionary history of Alpha (see Fig. 2B), between 2020-04-24 (95% HPD 2020-03-26 to 2020-05-24) and 2020-05-26 (95% HPD 2020-04-27 to 2020-06-30), i.e. between the TMRCA of B.1.1.7 and all background sequences, and the TMRCA of CAMC-934506 and B.1.1.7.

### Early growth rate of B.1.1.7 in the UK and interaction with November lockdown in England

Using a non-parametric coalescent model, we found that the growth of B.1.1.7 in England in the second half of 2020 was rapid compared to the background lineages present at the time (Fig. 3A). At the start of 2021, B.1.1.7 continued to grow, while other lineages began to decrease. These trends are broadly similar when comparing B.1.1.7 to just the B.1.177 lineage (Fig. S1B), which spread rapidly across the UK and became the dominant lineage over the summer of 2020 (Hodcroft et al. 2021).

**Figure 3.**
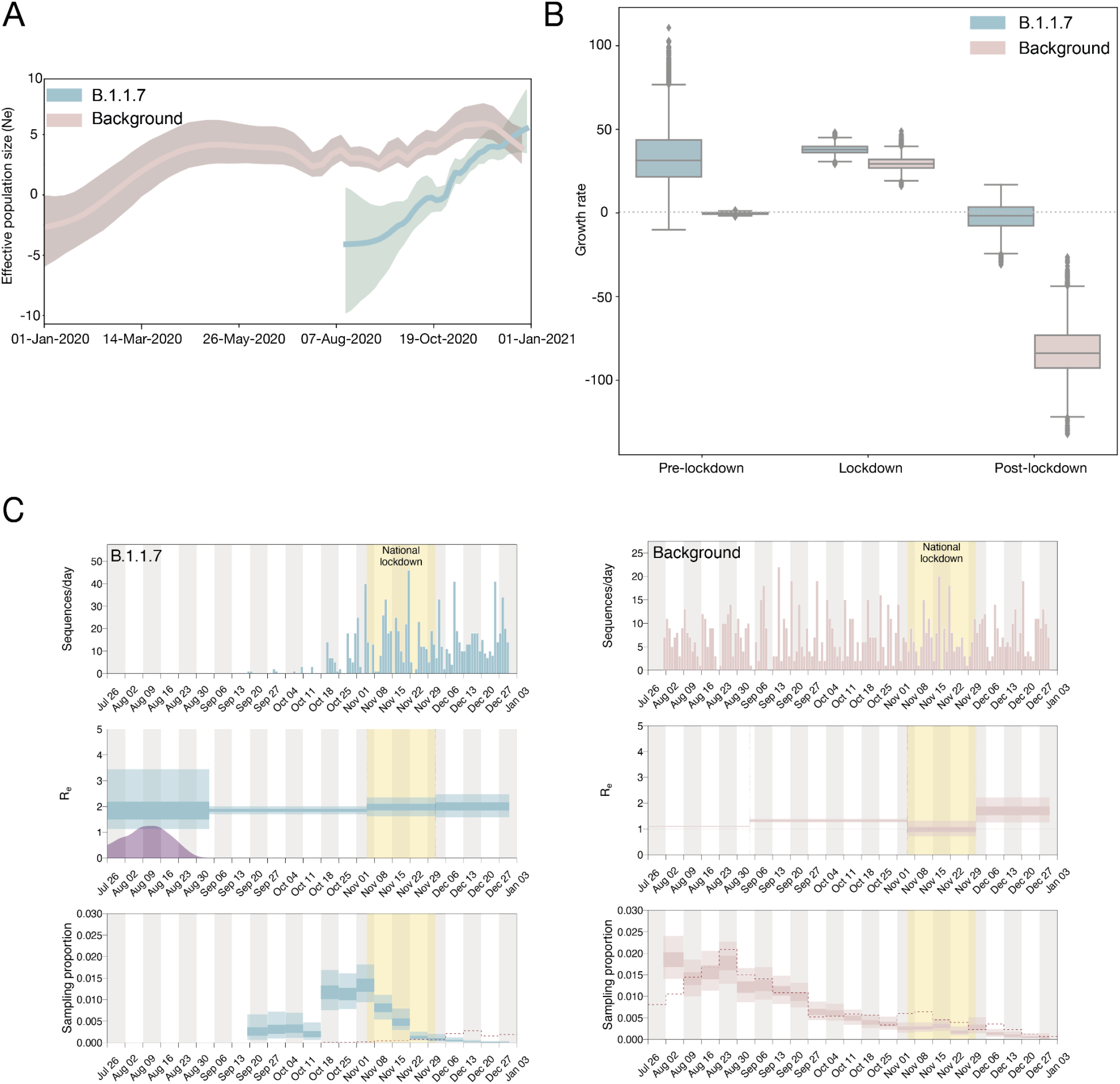
A) Effective population sizes for background lineages (pink) and B.1.1.7 (blue), generated from independent BEAST analyses. 95% HPDs are shown as shaded areas. B) Growth rate estimates with fixed transition times at pre-lockdown, lockdown, and post-lockdown, split by background lineage and B.1.1.7. C) Independent birth-death skyline analyses showing the number of sequences per day, the effective reproduction number (R_e_) and sampling proportion (which is allowed to vary on a weekly basis), for B.1.1.7 and the background. 95% HPDs are shown as light shaded areas and 50% HPDs as dark shaded areas. The English national lockdown in November is highlighted in all plots.

To further investigate the growth of B.1.1.7, we tested the difference between three standard population growth models: logistic, exponential, and epoch-based. For the last, wherein different epochs are permitted to have different growth rates, we estimated growth rates in the pre-lockdown period (2020-09-01 to 2020-11-04), during lockdown (2020-11-05 to 2020-12-04) and post-lockdown (2020-12-05 to 2020-12-31). Using a marginal likelihood estimation (MLE) approach, we found that the three-epoch model provided the best fit to the genomic data (Table S2). For the second time period, B.1.1.7 has a positive growth rate, and the post-lockdown period estimation includes zero; whereas the background lineages have a very strong negative growth rate in the most recent time period (Fig. 3B). This suggests that while the national lockdown in England in November significantly reduced the growth rates of both lineages, it was not sufficiently strict to push the growth rate of B.1.1.7 below zero. This reduced, but non-negative, growth rate for B.1.1.7 during the November lockdown has also been shown on a spatial level in (Kraemer et al. 2021).

In order to explore this further, we also estimated the effective reproduction number (R_e_), using a birth-death approach, which allowed the sampling proportion to vary to account for changes in genomic surveillance intensity across time (Fig. 3C). This showed that while both the background lineages and B.1.1.7 had an R_e_ above 1 (i.e. the epidemic was growing) in September and October, the English national lockdown in November was sufficient to push the R_e_ of the background lineages to around 1 (i.e. the epidemic was stable). However, the R_e_ of B.1.1.7 remained above 1, matching epidemiological information which showed growth of S-gene target failure positive cases despite the November lockdown (Kraemer et al. 2021).

### Other variants of concern

Under current WHO designations, there are four VOCs other than Alpha: Beta, discovered in South Africa at the end of 2020; Gamma, discovered in Brazil at the start of 2021; Delta, discovered in India at the start of 2021; and Omicron, discovered in South Africa and Botswana at the end of 2021. There are also two variants of interest (VOI): Lambda, discovered in Peru in mid-2021 and Mu, discovered in Colombia at the start of 2021. Each of these variants has had differing impacts across different regions, but the current Omicron wave has displaced almost all other lineages (outbreak.info).

Similarly to B.1.1.7, Omicron has a long ancestral branch (Fig. 4A), and the root-to-tip plot shows that Omicron sequences are distinct from the background diversity (Fig. 4B). However, as Omicron did not evolve out of the dominant circulating variant (i.e. Delta, B.1.617.2 and its descendants), it is more difficult to identify the clear pattern that can be observed in B.1.1.7, which evolved out of the dominant lineage at the time (i.e. B.1.1). Further, Omicron contains two distinct sibling clades, BA.1 and BA.2 which may represent two independent introductions into the general population. There is also a third lineage, which appears to be a recombinant of ancestral BA.1 and BA.2 sequences due to its mixture of some mutations found in both of them (Viana et al. 2022). The circumstances under which Omicron arose are clearly more complex than those that led to the evolution of Alpha. This could indicate a chronically infected individual or individuals with more contact with the general population (Maponga et al. 2022), or perhaps a non-human animal population.

**Figure 4.**
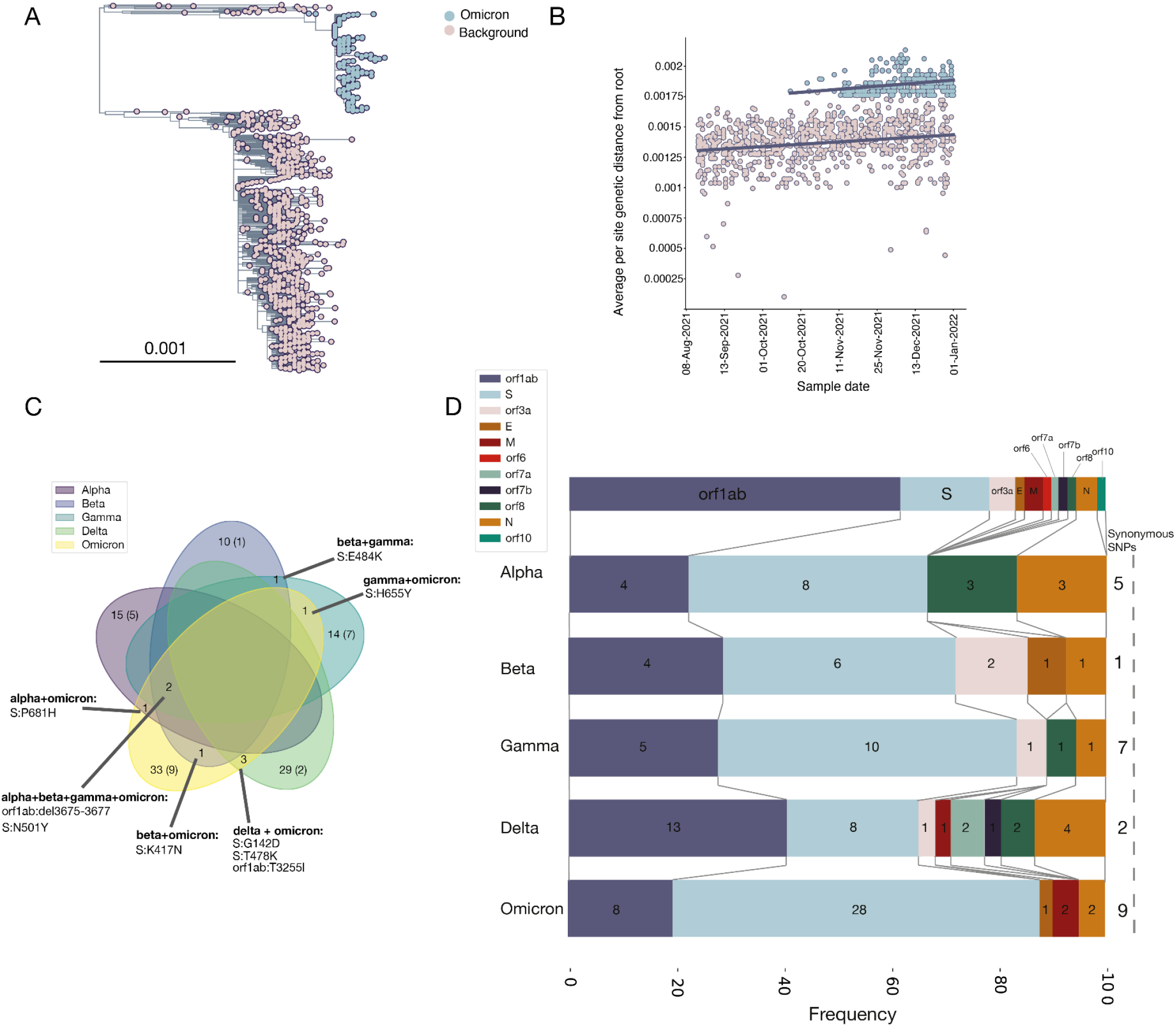
A) Phylogeny showing Omicron in blue and background sequences in pink. The large background group is the Delta variant, the dominant variant globally in the second half of 2021. B) Separate regressions of distance from the root against sample time for background sequences and Omicron sequences. Note that the parallel lines indicate similar rates of evolution within each clade (5.03 × 10^−4^ and 3.88 × 10^−4^ for Omicron and background lineages respectively). C) Venn diagram showing numbers of mutations shared between different variants of concern, with synonymous mutations in brackets. Zeroes, denoting no shared mutations acquired on the ancestral branch, are omitted. D) Frequency of non-synonymous mutations acquired on the ancestral branch in different parts of the genome between variants of concern. A schematic of the genome is shown along the top, numbers on each slice represent the absolute numbers of non-synonymous mutations or deletions in that gene, and numbers of synonymous mutations are shown along the right hand side.

There is no clear increase in evolutionary rate on the ancestral branches leading to Delta, Lambda or Mu (Fig. S3), suggesting that these variants may have arisen under more traditional evolutionary processes involving intense between-host transmission. On the other hand, Beta and Gamma show *some* evidence of an increased evolutionary rate on the ancestral branch (although this is stronger in Gamma than Beta which remains somewhat inconclusive, Fig. S3), and so may have a similar process of emergence involving a potential chronic infection.

Looking at mutations or patterns shared by variants of concern could provide evidence of common emergence routes or evolutionary pressures. We therefore collated mutations for each VOC (Alpha, Beta, Gamma, Delta and Omicron) compared to the background that they emerged from (see Methods) to identify any similarities. In general, there were very few shared mutations, and in particular, none shared by all variants. No synonymous mutations were shared between any variants (Fig. 4C).

The most commonly shared mutations were N501Y in Spike, and the nine base pair deletion in NSP6, which were both found in Alpha, Beta, Gamma and Omicron (Fig. 4C): all four of which show evidence of an increased evolutionary rate prior to their emergence. Notably, the intermediate genome for B.1.1.7 (CAMC-94506) also exhibits both N501Y and the NSP6 deletion. N501Y in particular has been monitored throughout the pandemic, due to its ability to increase binding to the ACE2 of human and murine cells. However, it appears that by itself, it is not necessarily enough to create a VOC, as there was a cluster in Wales in late 2020 defined by N501Y, but without the NSP6 deletion, which was rapidly outcompeted by Alpha (Fig. S2B).

Between Alpha and Omicron, which are the two variants with the *most* arguments for arising from chronic infections, there is only one unique shared mutation that was acquired during the evolution of the variant: P681H in the furin cleavage site of the Spike protein (Fig. 4C), which enhances Spike cleavage (Peacock et al. 2022). It must be noted also that Delta carries P681R, but shares no other mutations compared to the background lineages with Alpha and Omicron. The polybasic furin cleavage site is found in other coronaviruses, although not in any other Sarbecovirus, and it is required for SARS-CoV-2 virus entry into human lung cells (Hoffmann et al. 2020). Mutations in this area may be an adaptation to the human host, providing evidence for evolution in a human cryptic population. Of note, Peacock et al (Peacock et al. 2021) found that mutants with deletions in the furin cleavage site were rare; we speculate that this could be part of the fitness valley that Alpha (and Delta and Omicron) had to cross, as the furin cleavage site appears to be relatively conserved and so possibly many mutations are deleterious. It is also worth noting that while it is not a defining mutation of all sub-lineages of Omicron, the 69-70 deletion in the Spike protein is present in BA.1, which at the time of writing dominates most Omicron epidemics.

Finally, between Beta and Omicron, the variants with the most evidence for immune evasion, the single common mutation is K417N in the Spike protein (with a similar mutation, K417T found in Gamma, Fig. 4C). This mutation has been found to confer reduced susceptibility to neutralisation by specific monoclonal antibody therapies (Starr et al. 2021). This mutation also arose in AY.1, the so-called “Delta plus” variant descended from B.1.617.2 (Kannan et al. 2021), but this variant did not appear to acquire any noticeable advantage compared to the background Delta wave (outbreak.info).

In terms of the frequency of regions of mutations (Fig. 4D), all bar Delta have the highest frequency of non-synonymous mutations in the Spike protein, and Delta has the highest frequency in orf1ab (~38% of its mutations). Omicron has the highest frequency of Spike mutations (56%), Gamma the highest frequency of synonymous mutations (28%) and Alpha and Beta have the highest frequency of deletions (~13%). Overall there doesn’t appear to be a discernable pattern in the types or locations of deletions.

In comparing all of the VOCs, Alpha, Gamma and Omicron share the most evidence for a faster rate of evolution along their ancestral branch and Beta shows remains somewhat inconclusive, but with some evidence of the lineage having distinctly more root-to-tip divergence than expected. The VOIs and Delta show no evidence of a faster rate of evolution along their ancestral branches. Further, Delta carries mutations spread more evenly across its genome than the other VOCs, and consequently has fewer Spike mutations. This provides evidence for an alternate route of emergence for the Delta variant as compared to the other VOCs.

## Discussion

B.1.1.7/Alpha was first sampled in Kent, in South East England on 20th September 2020, and spread quickly across the rest of the UK (Kraemer et al. 2021; Volz et al. 2021). It was able to grow rapidly in the context of the NPIs applied in November 2020, which did not include school closure and were not as strict in restricting mixing; although these measures were sufficient to reduce the background lineage growth rate significantly. While these trends have been described previously (Kraemer et al. 2021; Volz et al. 2021), we have here reproduced them using only phylodynamic techniques and a small but representative genomic dataset. This finding will be useful for investigating VOCs which arise in areas with less genomic sequencing, or tracking those without e.g. SGTF drop-out.

Any proposed origin of B.1.1.7 must explain three observations: first, the long branch leading to the B.1.1.7 clade with at most one intermediate sequence, despite high genomic surveillance; second, an increased evolutionary rate along this branch; and third, a single geographical and evolutionary origin of B.1.1.7 (Kraemer et al. 2021).

In a country with an extensive virus genomic surveillance programme like the UK, which includes random and relatively dense sampling (an average of 7.9% of weekly reported cases in Kent and Medway between 24th April 2020 and 19th September 2020 were sequenced), it is unlikely that a precursor lineage was circulating in Kent over the summer of 2020 and was not detected. It is worth noting also that B.1.1.7 was captured by this surveillance programme within at most two days of its origin - the MRCA of the clade is the 19th September 2020 (see above), and the first sample was taken on 20th of the same month.

One possible explanation for the lack of detection of the precursor lineage to B.1.1.7 in the UK surveillance data is simply that it was not in the UK prior to its expansion in South East England, but in a region of the world with little or no genomic surveillance. However, this hypothesis requires that the lineage was introduced twice into the UK (firstly for the intermediate sequence and secondly for the B.1.1.7 lineage) without being exported and establishing transmission anywhere else. Genomic surveillance has since been scaled up in many regions, and the fact that no descendents of such a cryptic population have been sampled to date indicates either that this population went extinct (unlikely given the fitness advantages conferred by the lineage-defining mutations) or that no such population existed. Furthermore, transmission between humans, even if rapid, would not explain the higher rate of evolution observed along the branch. These explanations would also apply to a population in the UK which is disproportionately under-sampled, for example vulnerable communities, such as individuals experiencing homelessness, who are unlikely or unable to seek healthcare.

An alternative explanation is a zoonotic event, as SARS-CoV-2 has been shown to spread in non-human animals, for example in mink (Oreshkova et al. 2020; Oude Munnink et al. 2021), white-tailed deer (Chandler et al. 2021), and Syrian hamsters (Yen et al. 2022). In this hypothesis, there would have been a reverse zoonosis from humans, an increased rate of molecular evolution among animals, perhaps due to natural selection for the new host species, followed by at most two zoonotic events in the course of several months (the intermediate and the final clade). For the former, in an animal population that had sufficient contact with humans for a reverse zoonotic event and then two later zoonoses, it is unlikely that there would be only two spillovers in five months: in mink farms in the Netherlands, it was estimated that there were 43 spillovers between April and November 2020 (Lu et al. 2021); and in a pet shop in Hong Kong, there were at least two spillovers in the space of a few weeks (Yen et al. 2022). Further, transmission between animals has not been observed to lead to a higher rate of evolution: even in a large outbreak among densely farmed mink, the rate of evolution was estimated to be similar to what is expected between humans, at approximately 7.9×10^−4^ nucleotide changes/site/year (Lu et al. 2021). Further, it is unlikely that a variant so effective at spreading in the human population would have evolved in a non-human population; as the mutations required to be successful in a human population may well be different due to differences in cell receptors, as well as behaviour. Common mutations appear in animal populations, such as N501T and Y453F in mink across different continents (Eckstrand et al. 2021; Lu et al. 2021), but are rarely found in human infections, and not in any of the VOCs. While it would be possible for a two-step evolutionary process wherein there is first some adaptation in an animal population, allowing the crossing of a fitness valley, followed by spillover and human adaptation through conventional transmission, it would once again be difficult to explain the intermediate sample we observe through this transmission process.

We propose that the most likely explanation for the emergence of B.1.1.7 is that an individual was chronically infected with SARS-CoV-2 over the course of months providing an evolutionary environment conducive to the virus making adaptive jumps. The evolutionary environment within a single host is different to that at the between-host scale, with a large effective population size and the opportunity for recombination (Jackson et al. 2021). This large effective population size can be established and maintained partially due to the different compartments that a respiratory virus can establish infection in, for example, upper and lower respiratory tract, as well as deeper into the lung (e.g. (Lakdawala et al. 2015; Richard et al. 2020). Conversely, the effective population size at the between-host level is small due to tight bottlenecks occurring on transmission (Lythgoe et al. 2021). Furthermore, although a persistent infection will provide the time and selective environment for a period of adaptive evolution, the exact cause of the persistence may affect the traits of the virus that are selected for.

There are a number of studies on chronically or persistently infected individuals which contain longitudinal sampling of the viruses present (Avanzato et al. 2020; Choi et al. 2020; Clark et al. 2021; Karim et al. 2021; Kemp et al. 2021; Ramírez et al. 2021; Stanevich et al. 2021; Voloch et al. 2020; Weigang et al. 2021; Williamson et al. 2021). Across these studies, there were an average of 4.0 (95% confidence interval: 0.63 to 9.76) evolutionary events (i.e. gaining or losing a mutation compared to the individual’s first sample) per week (Table S3), compared to the approximately 0.5 mutations expected in between-host transmission. Further, in these studies, deletions (notably, the 69/70 deletion found in Alpha and the BA.1 sub-lineage of Omicron) have been found to both increase and decrease in frequency along the time period of infection (Kemp et al. 2021; Stanevich et al. 2021). As it is unlikely that a deletion could be reverted, this is evidence for multiple coexisting viral populations (Lythgoe et al. 2021).

Much of the discussion of variant emergence from within-host evolution has focussed on the idea of individuals with compromised immune systems, either pathologically (e.g., a lymphoma) or medically (e.g., post-transplant suppression or chemotherapy) induced (Karim et al. 2021; Maponga et al. 2022). Persistent infections have also been recorded in apparently immunocompetent individuals (Ramírez et al. 2021; Voloch et al. 2020) and although the average length of these infections is shorter than in immunocompromised cases – from the studies above, an average of 32 days and 174 days respectively (Figure S2B). In one case an immunocompetent individual was infected for approximately 64 days (Voloch et al. 2020). Furthermore, there may be an observation bias towards data from immunocompromised hosts and long term infections in immunocompetent hosts are less likely to be recorded.

The degree of immunocompromisation is variable and will depend on whether antibody or cellular immunity (or both) are affected by the condition of the individual. Grenfell and colleagues used a simple population genetic model to posit that the rate of viral adaptation is a non-linear function of immune (selection) pressure, because of the opposing effects of raised immune pressure on virus population size and average selection coefficients. Hence the highest viral adaptation rate is predicted to arise from intermediate immune pressures (Grenfell et al. 2004). Although there is currently no evidence of a difference in numbers of evolutionary events between immunocompromised and immunocompetent hosts (Fig. S2A), we propose that this may be because these cases lie on either side of the peak of viral adaptation rate in the Grenfell et al. model. Thus within-host virus evolution in both healthy and immunocompromised hosts could lead to an increased evolutionary rate; and we have not yet observed an individual at the part of the immune spectrum which would lead to the explosive adaptation seen in Alpha and Omicron.

The absence of more than one sampled transmission chain arising from intermediate combinations of the Alpha mutations, even on the background of mild NPIs in the summer of 2020 in the UK; as well as this constellation not evolving elsewhere across the phylogeny, suggests that the fitness peak is difficult to reach, despite its large selective advantage compared to background lineages. This implies a complex fitness landscape, and a large fitness valley prior to the peak that Alpha represents for between-host fitness. The different selective pressures inside a host could enable the crossing of such a valley due to a relaxation of selection on transmission-based adaptations in favour of factors such as evading neutralising antibodies, as found in longitudinal samples in (Weigang et al. 2021), or focussing on cell entry. The intermediate sequence is likely part of a transmission chain that was ultimately very short, as the virus was still not particularly well adapted to transmitting between hosts at this time point.

However, Alpha and other VOCs clearly became well adapted to spreading between hosts as well as within a host. While a transmission advantage would not be explicitly selected for within hosts, traits which are useful within a host could also lead to better transmission. For example, increased ACE2 binding which increases the efficiency of cell entry (Ozono et al. 2021) would allow a virion to enter and use a host cell faster than its competitor within a host, leading to faster growth; and would also make it easier for a virus population to establish an infection in a new host. The evasion of a host immune response is another clear advantage within a host, allowing fewer virions to be destroyed by the immune system, and on a population level, especially in the immune context of widespread previous infection and vaccination. This is seen most clearly in Omicron, and provides a possible explanation as to why there are three sub-lineages of Omicron which all arose at once: the sub-lineages all acquired immune evasion properties within a host, and then spread from diverse viral populations within that host. Finally, and non-exclusively, an efficient and host-adapted infection could lead to a large amount of viral shedding, increasing transmissibility.

From the mutational profiles and the evolutionary rates of the VOCs, it appears that Delta is an outlier: it does not contain N501Y or the deletion in NSP6, nor does it appear to have a higher rate of evolution leading up to its emergence, and it has fewer non-synonymous mutations in its Spike protein as well as more mutations in other parts of the genome compared to the other VOCs. We therefore hypothesise that it may have followed an alternative route of emergence, perhaps simply intense between-host transmission in an undersampled location. Given that the estimate of the TMRCA is several months prior to the first sample (McCrone et al. 2021), it is plausible that the lineage-defining mutations could have been acquired sequentially, prior to a larger explosion of cases once the full constellation was present. The common patterns found in the emergence of Alpha, Beta, Gamma and Omicron provide some evidence that Beta, Gamma and Omicron may also have arisen through chronically infected individuals. It is notable that Southern Africa, where Beta and Omicron were first sampled, has a high prevalence of people living with HIV. An individual with a poorly-controlled HIV infection would provide another avenue for a large viral population to be maintained in an individual over a long period of time and therefore new between-host fitness peaks to be explored. Indeed, an individual with an uncontrolled HIV infection accumulated more than 20 mutations in the course of 9 months (Maponga et al. 2022). In this case the HIV virus was controlled and the SARS-CoV-2 cleared through antiretroviral treatment. However, in other circumstances progression of an HIV infection could then allow the partially controlled SARS-CoV-2 infection to proliferate significantly allowing for shedding and therefore transmission. Accessible antiretroviral therapy is therefore a key element of mitigating the risk of further SARS-CoV-2 variant emergence in countries with significant numbers of people living with HIV, as called for by Maponga *et al (Maponga et al. 2022).* More generally, equitable and universal access to SARS-CoV-2 vaccination and antiviral drugs will be a critical strategy given the apparent diversity of circumstances by which VOCs have emerged thus far.

Chronic COVID-19 cases are relatively rare, but as another wave of transmission sweeps across the world, there will be many more as has been seen recently with the Omicron variant (Viana et al. 2022). If all persistent infections present a risk of a new, highly transmissible or immune evasive variant, then simply shielding the vulnerable and selective vaccination will not be sufficient to prevent the emergence of another wave of morbidity and mortality. Without urgent and truly widespread vaccination efforts and dispersal of antiviral medication, we expect to see the delayed impacts of uncontrolled transmission resulting from vaccine and antiviral inequity into the future.

## Methods

### Genomic dataset

The COG-UK alignment and metadata of all SARS-CoV-2 genomes from 2021-04-21 was restricted to between 2020-08-01 and 2020-12-31 and surveillance (i.e. pillar 2) sequences from England. This dataset was then subsampled in a time-homogenous way to generate approximately 1000 sequences per sequence set which comprised 50 sequences per week for non-B.1.1.7 sequences and 100 per week for B.1.1.7 sequences. The B.1.1.7 dataset was checked for molecular clock outliers (sequences that have disproportionately too much or too little root-to-tip divergence for its sampling time (Hill & Baele 2019)) using TempEST, and one sequence was identified and removed (England/CAMC-BBDA45/2020). The resulting dataset was 1100 background sequences and 976 B.1.1.7 sequences.

For mutation scanning, the sequences were aligned to the reference sequence Wuhan-Hu-1 using minimap2 (Li 2018) and gofasta (https://github.com/cov-ert/gofasta). Variants were determined using type_variants.py (https://github.com/cov-ert/type_variants).

### Evolutionary rate calculation

First, a maximum likelihood tree was generated using IQTree v2.1.2 (Minh et al. 2020) and an HKY substitution model (Hasegawa et al. 1985). This was used to generate the plots showing a linear regression of root-to-tip genetic distance against sampling date and provide estimates of the rate of evolution in background sequences and within the B.1.1.7 clade (slope of the linear regression).

To estimate the rate of evolution in the branch leading up to B.1.1.7, a local clock model in BEAST v1.10.4 (Suchard et al. 2018) was used. Briefly, a strict clock model was applied to each of the three groups so that their clock rates could be estimated independently, each with a Gamma prior. Preliminary analyses suggested that the within-B.1.1.7 evolutionary rate was similar to the background rate, and so the same clock rate model was placed on both the background and the within-B.1.1.7 clade. Nested taxon sets containing the one plausible intermediate sequence (CAMC-946506) were used to estimate the dates of the most recent common ancestor of B.1.1.7 and the intermediate sequence. A nonparametric coalescent Skygrid model (Gill et al. 2012) with 64 change-points spanning 15 months was placed on the background sequences including the intermediate sequence, and an exponential growth coalescent model was placed on B.1.1.7. For both sequence alignments an HKY substitution model was used. Two chains with 100 million states were run, and following assessment via Tracer (Rambaut et al. 2018), 45 million states from each were removed as burn-in.

### Growth rate calculations

To describe general patterns in the growth of B.1.1.7 compared to the background rate, as well as B.1.177 (N=1069) and a Welsh cluster containing N501Y (N=478), we ran a series of non-parametric Skygrid analyses (Gill et al. 2012). These were run independently in BEAST, each for two chains of 100 million states. For B.1.1.7, B.1.177 and the Welsh cluster, 77 change-points were used, spaced equidistantly between the most recent tip and 0.75 of a year before it (approximately 9 months). For the background dataset, 64 change-points were used with 1.25 of a year as the cutoff. All analyses assumed a strict molecular clock model and the HKY substitution model (Hasegawa et al. 1985).

To compare parametric growth models, a logistic, exponential and 3-epoch growth model were run only on the B.1.1.7 dataset. The 3-epoch model used fixed transition times and exponential growth rates for within each epoch. Each of them used an HKY substitution model and a strict clock model, and two independent chains of 100 million states were run. A marginal likelihood estimation (MLE) analysis, a commonly used form of Bayesian model selection integrated into the BEAST software package, was used to distinguish between these three models (Suchard et al. 2001).

To infer growth rates using the 3-epoch model, we combined the B.1.1.7 and background lineage datasets to simultaneously estimate growth rates for both lineages during each of the epochs (N=2076). Transition times were fixed at the start and end of the lockdown in England (2020-11-05 and 2020-12-02), and an earlier transition time was also placed on 2020-09-01 (the TMRCA of B.1.1.7) at the start of the study period to focus on the growth after B.1.1.7 began to diversify. For variable transition times, the oldest transition time remains fixed, and the other two are allowed to vary under normally distributed priors with means at the start and end of the lockdown, respectively, and standard deviations of two weeks. Both analyses were run for two chains independently for 100 million states.

To estimate differences in the effective reproduction number (R_e_) between B.1.1.7 and background lineages, a Bayesian birth-death skyline (Stadler et al. 2013) model was run independently on the B.1.1.7 and background datasets. An HKY substitution model was used along with a strict clock model, and a Gamma prior with *α*=0.001 and *β*=1000 was placed on the clock rate. A lognormal prior with mean 0.8 and standard deviation 0.5 was placed on R_e_ and a Beta prior with *α*=2 and *β*=1000 on the sampling proportion. R_e_ was parameterised into 4 epochs with transition times fixed at the start and end of the lockdown in England (2020-11-05 and 2020-12-02), and an earlier transition time placed at 2020-09-04. The sampling proportion was fixed to 0 before the first week containing a sample and then estimated for each week thereafter, resulting in 16 epochs for B.1.1.7 (from 2020-09-19) and 23 for the background dataset (from 2020-08-01). The becoming-uninfectious rate was assumed to be constant and fixed at 36.5, which is equivalent to a mean period of 10 days from infection to loss of infectiousness (through recovery, isolation or death). Analyses were started from initial trees estimated in IQTree v2.1.2 (Minh et al. 2020) and scaled to calendar time using TreeTime (Sagulenko et al. 2018). For both datasets four chains of 100 million iterations were run independently, sampling states and trees every 10'000 iterations. Chains were combined after removing 10% of states as burn-in and convergence assessed using Tracer (Rambaut et al. 2018). Convergence was assessed using the R-package coda (Plummer et al. 2006) and 10% of states were removed to account for burn-in.

### Sequencing proportion

Case data was collated from the UK government dashboard (https://coronavirus.data.gov.uk/). Cases and sequences with sample dates between 2020-04-24 and 2020-09-19 (the first day of the week of the most common recent ancestors of the stem of B.1.1.7 and the clade of B.1.1.7 respectively) were aggregated by week. Cases and sequences with the locations of “Kent” or “Medway”, a county surrounded by Kent, were included.

### Rates of evolution in chronically-infected individuals

For each of the eight studies identified containing longitudinal samples of chronically-infected individuals, we counted the number of mutations present per individual in the paper. A mutation to the derived state and back to the reference allele each counted as a separate evolutionary event, relative to the first individual sample available, which was within the first week of infection start for all papers other than Karim *et al* (day 12) and Avanzato *et al* (day 49). For papers (e.g. Kemp *et al*) where proportions of variants were given, a 25% cutoff was used for presence/absence of a mutation. Ambiguous or missing data was treated as the reference allele.

The rate of mutation events was calculated by dividing the number of events by the number of days the individual was followed up divided by 7 to change the denominator to weeks. Note that for Karim *et al,* only non-synonymous substitutions were provided in the paper, meaning that the estimate of 2.04 events per week is an underestimate.

Start of infection was taken to be the start of symptoms or the date of the first positive PCR test depending on what was available for each study. End of infection was the date of the final negative PCR test in the study (Avanzato et al. 2020; Karim et al. 2021; Stanevich et al. 2021; Voloch et al. 2020; Weigang et al. 2021; Williamson et al. 2021), death (Choi et al. 2020) or when the individual was lost to follow up (Ramírez et al. 2021).

### Other variant analyses

For performing the linear regression of root-to-tip genetic distance against sampling date to provide estimates of the rate of evolution in background sequences and within each VOC lineage, background datasets were obtained from the master COG-UK alignment. For Delta, Lambda and Mu, sequences sampled globally between 2021-01-01 and 2021-06-01 that were not any of the above variants were downsampled to 50 per week where possible. For Gamma and Beta, the same background dataset as Alpha was used (see above). Then sequences from the correct time period for each variant were taken, and all were downsampled to 50 sequences per week, where possible. Finally, all sequences were run through metadata and sequence quality control. The final dataset sizes were 390 for the background set, 141 for Beta, 31 for Gamma, 373 for Delta, 246 for Lambda, and 208 for Mu.

For Omicron, B.1 sequences sampled between 2021-09-01 and 2022-01-01 and all Omicron sequences until the same cut-off were taken. These were then also downsampled to 50 per week, resulting in 900 background sequences and 390 Omicron sequences. Omicron sequences with any reference calls, any Delta mutations and more than three missing SNPs were excluded, as well as two molecular clock outliers (see above) in the background dataset. The final dataset comprised 898 background and 328 Omicron sequences.

To undertake the mutation analysis for each variant, the first ten sequences for each variant other than Delta were taken from after the oldest reported sample in the original paper or report describing the variant. These were from 2020-09-20 for Alpha (Rambaut et al. 2020a), 2020-10-15 for Beta (Tegally et al. 2021), 2020-12-17 for Gamma (Faria et al. 2021) and 2021-11-15 for Omicron (Viana et al. 2022). For Delta, due to an unclear starting point and large amounts of missing data in the sequences, all Delta sequences in March 2021 (following the estimate of the start of Delta expansion in March in (McCrone et al. 2021)) were extracted from the COG-UK alignment, and run through Scorpio (https://github.com/cov-lineages/scorpio), filtering to allow no reference alleles (from Wuhan-Hu-1) alleles and a maximum of 2 missing alleles in lineage-defining positions. This resulted in 98 sequences, ten of which were from India, which were taken as the representative group. For all variants, mutations which were common to all representative ten sequences were taken, and supplemented with any lineage-defining mutations which were missing (due to a small amount of missing data), based again on the original defining publication for each variant.

These were then compared to a representative background set. For each variant, this entailed up to ten sequences from the month of the first reported sample of the parent lineage: B.1.1 for Alpha and Omicron, B.1 for Beta and Omicron, and B.1.1.28 for Gamma.

## Supporting information

COG authorship

GISAID Acknowledgements table

## Data Availability

UK genome sequences used were generated by the COVID-19 Genomics UK consortium (COG-UK, https://www.cogconsortium.uk/). Non-UK data was from GISAID (gisaid.org), and an acknowledgements table for the sequences used can be found in the supplementary materials.

## Acknowledgements

COG-UK is supported by funding from the Medical Research Council (MRC) part of UK Research & Innovation (UKRI), the National Institute of Health Research (NIHR) [grant code: MC_PC_19027], and Genome Research Limited, operating as the Wellcome Sanger Institute. V.H. was supported by the Biotechnology and Biological Sciences Research Council (BBSRC) [grant number BB/M010996/1]. T.P.P and W.S.B are supported by the G2P-UK National Virology Consortium funded by the MRC [grant number MR/W005611/1]. J.T.M, R.C. and A.R. acknowledge support from the Wellcome Trust [Collaborators Award 206298/Z/17/Z - ARTIC network]. A.R. is also supported by the European Research Council [grant agreement number 725422 - ReservoirDOCS] and Bill & Melinda Gates Foundation [OPP1175094 – HIV-PANGEA II]. A.OT is supported by the Wellcome Trust Hosts, Pathogens & Global Health Programme [grant number: grant.203783/Z16/Z] and Fast Grants [award number: 2236]. O.G.P. and L.dP. acknowledge support from the Oxford Martin School. D.A. is a Wellcome Clinical PhD Fellow and gratefully supported by the Wellcome Trust (Grant number: 222903/Z/21/Z). I.G. is a Wellcome Senior Fellow and supported by the Wellcome Trust (Grant number: 207498/Z/17/Z). E.V. is supported by the Wellcome Trust (Grant number: 220885/Z/20/Z).

