## Supplementary Information for "The origins and molecular evolution of SARS-CoV-2 lineage B.1.1.7 in the UK"

**Table S1** | Non-synonymous mutations and deletions inferred to occur on the branch leading to lineage B.1.1.7.

| gene | nucleotide | amino acid | Present in CAMC-946506? | Present in MILK-B154B6? | Amplicon |
|---|---|---|---|---|---|
| ORF1ab | C3267T | T1001I | No | No | 11 |
| | C5388A | A1708D | No | No | 18 |
| | T6954C | I2230T | No | Yes | 23 |
| | 11288-11296 deletion | SGF 3675-3677 deletion | Yes | No | 37 |
| Spike | 21765-21770 deletion | HV 69-70 deletion | No | Yes | 72 |
| | 21991-21993 deletion | Y144 deletion | No | Yes | 72/73 |
| | A23063T | N501Y | Yes | Mixture | 76 |
| | C23271A | A570D | No | No | 77 |
| | C23604A | P681H | No | Yes | 78 |
| | C23709T | T716I | No | Yes | 78 |
| | T24506G | S982A | No | Mixture | 81 |
| | G24914C | D1118H | No | No | 82 |
| Orf8 | C27972T | Q27stop | Yes | No | 92 |
| | G28048T | R52I | Yes | No | 92 |
| | A28111G | Y73C | No | No | 92/93 |
| N | 28280 GAT->CTA | D3L | No | No | 93 |

| | | | | |
|---|---|---|---|---|
| C28977T | S235F | No | Yes | 95 |

**Table S2** | Marginal Likelihood Estimation of different growth rate models

| Model | Path sampling | Stepping stone sampling | Order |
|---|---|---|---|
| Three epoch | -55785.30 | -55802.68 | Best |
| Logistic | -55842.26 | -55858.27 | Second |
| Exponential | -55844.18 | -55861.28 | Third |

**Table S3** | Information about individuals used in chronic individual analysis

| Paper | Individual number | Length of infection (days) | Time between first and last sequence (days) | Type of immunocompromis-ation | Number of mutation events |
|---|---|---|---|---|---|
| Karim *et al* 2021 | 1 | 228 | 190 | HIV positive | 57 |
| Weigang *et al* 2021 | 1 | 149 | 140 | Organ transplant recipient | 59 |
| Ramirez *et al* 2021 | 1 | 48 | 45 | N/A | 4 |
| Ramirez *et al* 2021 | 2 | 36 | 30 | N/A | 1 |
| Choi *et al* 2020 (further described in Clarke *et al* 2021) | 1 | 157 | 152 | B cell depletion | 52 |
| Kemp *et al* 2021 | 1 | 102 | 101 | Combined immunodeficiency | 150 |

| | | | | | |
|---|---|---|---|---|---|
| Williamson *et al* 2021 | 1 | 311 | 290 | B cell depletion | 124 |
| Stanevich *et al* 2021 | 1 | 318 | 308 | B cell depletion | 79 |
| Avanzato *et al* 2020 | 1 | 105 | 56 | B cell depletion | 10 |
| Voloch *et al* 2021 | 5 | 28 | 26 | N/A | 9 |
| Voloch *et al* 2021 | 6 | 40 | 28 | N/A | 8 |
| Voloch *et al* 2021 | 8 | 42 | 11 | N/A | 14 |
| Voloch *et al* 2021 | 9 | 41 | 27 | N/A | 15 |
| Voloch *et al* 2021 | 11 | 32 | 23 | N/A | 16 |
| Voloch *et al* 2021 | 12 | 45 | 39 | N/A | 14 |
| Voloch *et al* 2021 | 13 | 29 | 20 | N/A | 17 |
| Voloch *et al* 2021 | 14 | 52 | 10 | N/A | 4 |
| Voloch *et al* 2021 | 15 | 35 | 20 | N/A | 9 |
| Voloch *et al* 2021 | 16 | 28 | 20 | N/A | 15 |
| Voloch *et al* 2021 | 19 | 30 | 12 | N/A | 3 |
| Voloch *et al* 2021 | 21 | 27 | 21 | N/A | 11 |
| Voloch *et al* 2021 | 22 | 24 | 21 | N/A | 7 |
| Voloch *et al* 2021 | 23 | 64 | 22 | N/A | 0 |
| Voloch *et al* | 25 | 30 | 21 | N/A | 12 |

| | | | | | |
|---|---|---|---|---|---|
| 2021 | | | | | |
| Voloch *et al* 2021 | 26 | 23 | 20 | HIV positive | 10 |
| Voloch *et al* 2021 | 27 | 23 | 20 | N/A | 13 |
| Voloch *et al* 2021 | 29 | 22 | 21 | N/A | 6 |
| Voloch *et al* 2021 | 31 | 21 | 17 | N/A | 45 |
| Voloch *et al* 2021 | 32 | 21 | 19 | N/A | 3 |
| Voloch *et al* 2021 | 33 | 35 | 14 | N/A | 2 |
| Voloch *et al* 2021 | 34 | 21 | 19 | N/A | 2 |
| Voloch *et al* 2021 | 35 | 31 | 11 | N/A | 9 |
| Voloch *et al* 2021 | 36 | 28 | 11 | N/A | 12 |
| Voloch *et al* 2021 | 37 | 28 | 11 | N/A | 12 |
| Voloch *et al* 2021 | 38 | 27 | 14 | N/A | 11 |
| Voloch *et al* 2021 | 39 | 28 | 17 | N/A | 2 |
| Voloch *et al* 2021 | 40 | 48 | 20 | N/A | 7 |
| Voloch *et al* 2021 | 41 | 20 | 12 | N/A | 3 |
| Voloch *et al* 2021 | 43 | 26 | 5 | N/A | 7 |
| Voloch *et al* 2021 | 44 | 26 | 6 | N/A | 5 |
| Voloch *et al* 2021 | 45 | 25 | 14 | N/A | 6 |

| | | | | | |
|---|---|---|---|---|---|
| Voloch *et al* 2021 | 46 | 18 | 16 | N/A | 16 |

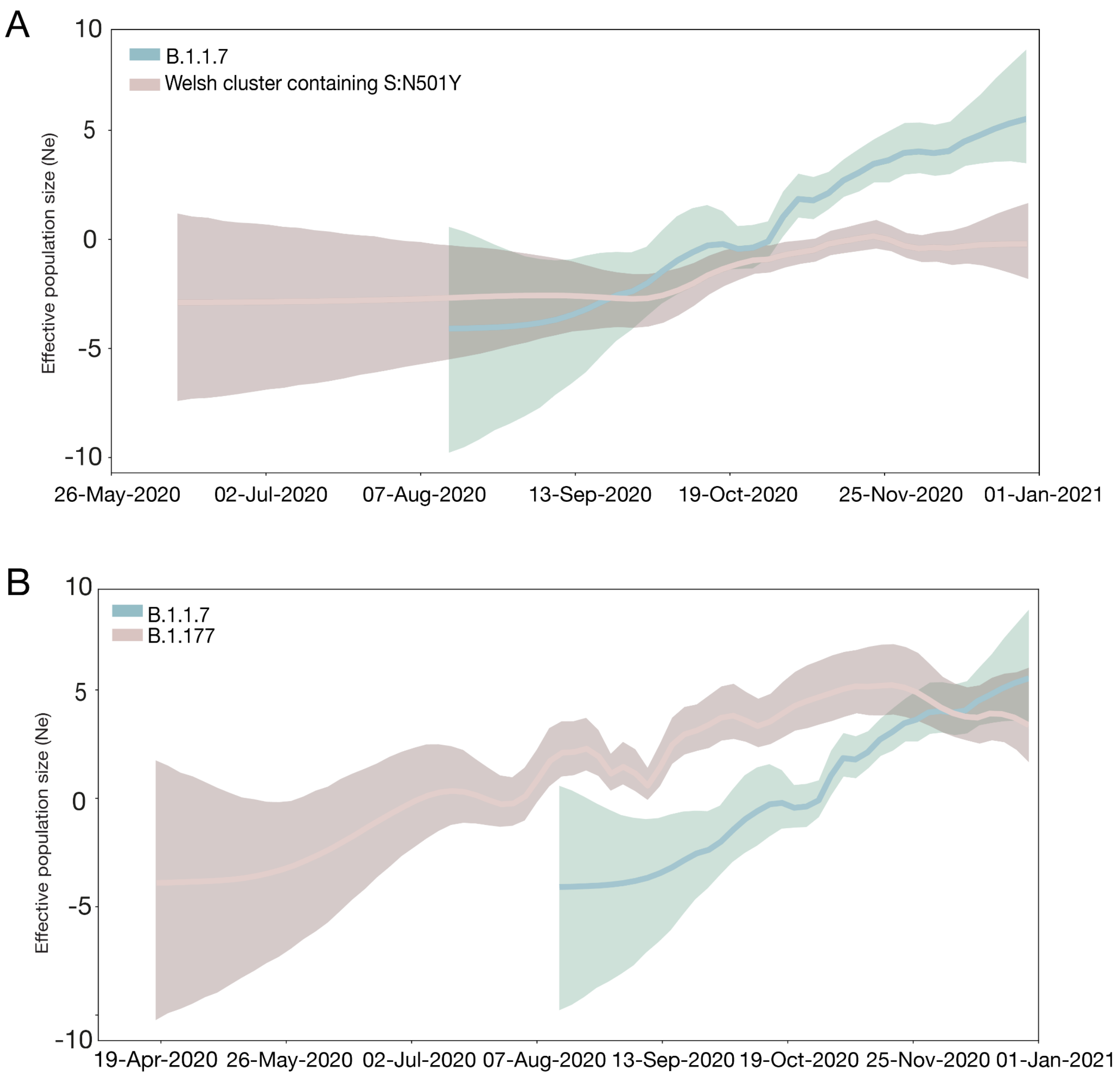


***Figure S1*** | *Effective population sizes of B.1.1.7 (blue) against: A) a Welsh cluster defined by N501Y B) B.1.177. Both are generated from independent BEAST analyses. 95% HPDs shown as shaded areas.*

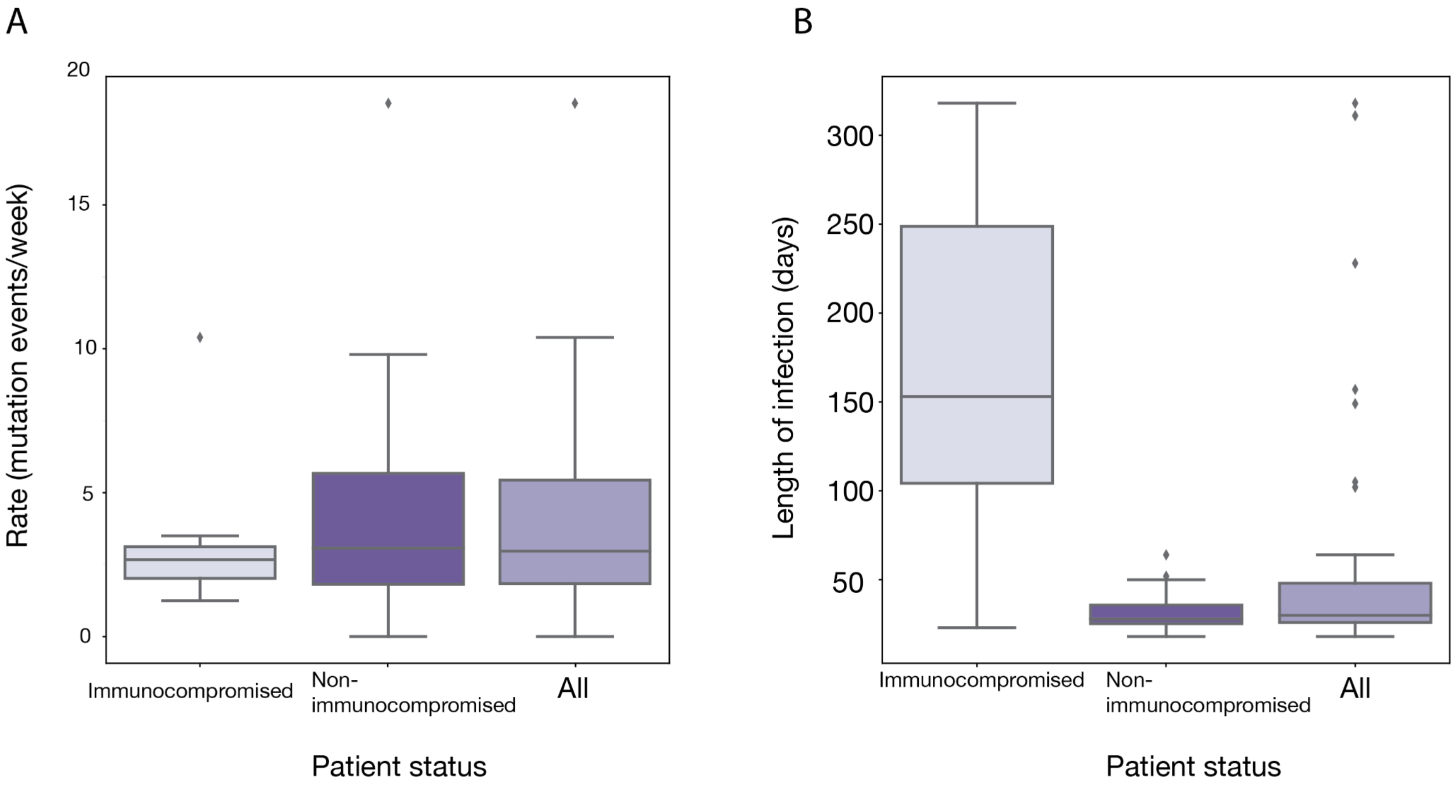


***Figure S2*** | *Results from longitudinally-sampled patients from across seven papers. A) Estimated mutation rate in terms of number of mutation events per week. B) Length of infection in days.*

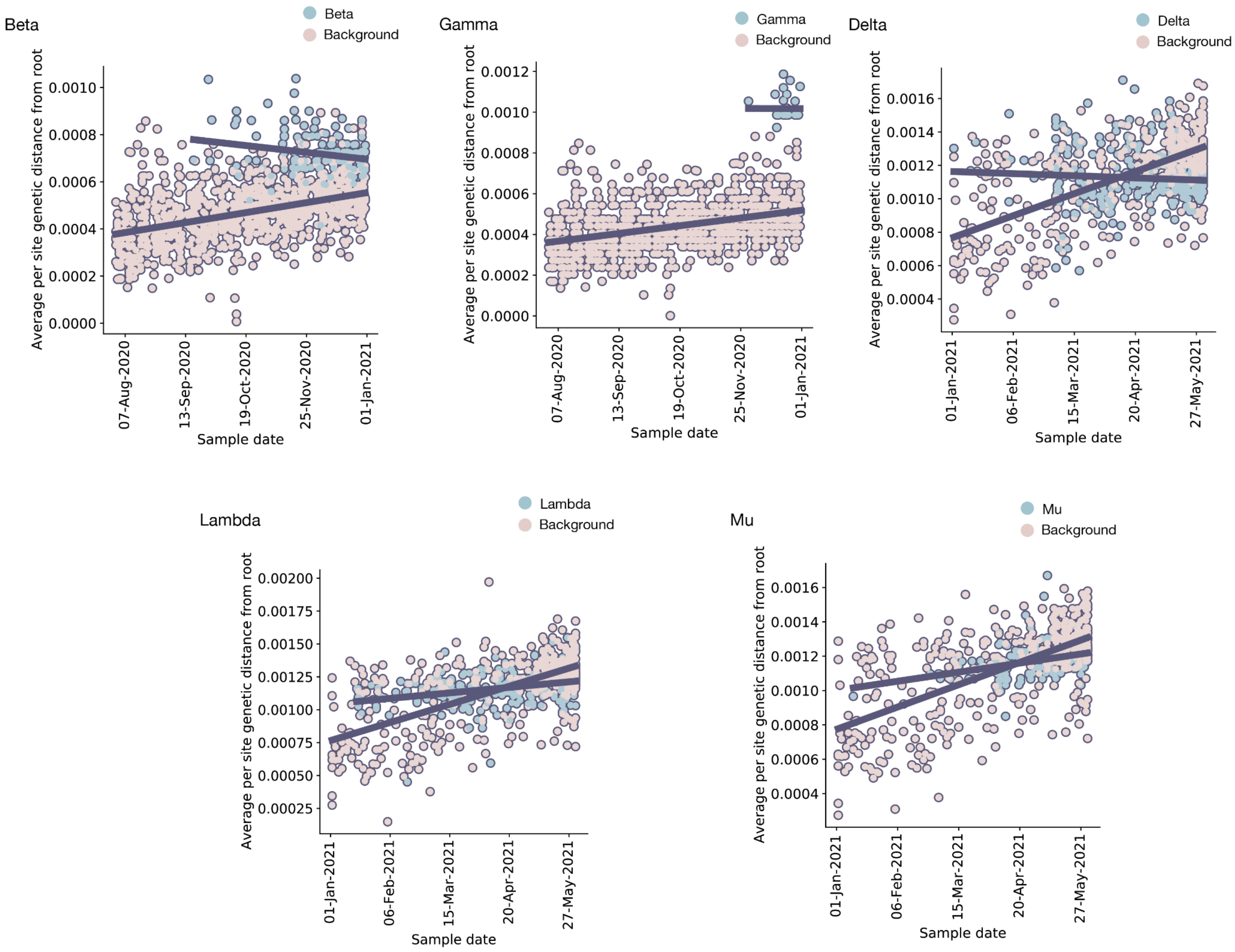


***Figure S3*** | *Root to tip divergence plots for the five other variants of concern (Beta, Gamma and Delta) or variants of interest (Lambda and Mu) as designated by the WHO. None show any noticeable difference in evolutionary rate (shown by the lines) between the background sequences and the relevant variant.*
