## Supplementary material for "The origins and molecular evolution of SARS-CoV-2 lineage B.1.1.7 in the UK": COG authorship

**June 2021 V.1**


**Funding acquisition, Leadership and supervision, Metadata curation, Project administration, Samples and logistics, Sequencing and analysis, Software and analysis tools, and Visualisation:**
Samuel C Robson [13, 84]

**Funding acquisition, Leadership and supervision, Metadata curation, Project administration, Samples and logistics, Sequencing and analysis, and Software and analysis tools:**
Thomas R Connor [11, 74] and Nicholas J Loman [43]

**Leadership and supervision, Metadata curation, Project administration, Samples and logistics, Sequencing and analysis, Software and analysis tools, and Visualisation:**
Tanya Golubchik [5]

**Funding acquisition, Leadership and supervision, Metadata curation, Samples and logistics, Sequencing and analysis, and Visualisation:**
Rocio T Martinez Nunez [46]

**Funding acquisition, Leadership and supervision, Project administration, Samples and logistics, Sequencing and analysis, and Software and analysis tools:**
David Bonsall [5]

**Funding acquisition, Leadership and supervision, Project administration, Sequencing and analysis, Software and analysis tools, and Visualisation:**
Andrew Rambaut [104]

**Funding acquisition, Metadata curation, Project administration, Samples and logistics, Sequencing and analysis, and Software and analysis tools:**
Luke B Snell [12]

**Leadership and supervision, Metadata curation, Project administration, Samples and logistics, Software and analysis tools, and Visualisation:**
Rich Livett [116]

**Funding acquisition, Leadership and supervision, Metadata curation, Project administration, and Samples and logistics:**
Catherine Ludden [20, 70]

**Funding acquisition, Leadership and supervision, Metadata curation, Samples and logistics, and Sequencing and analysis:**
Sally Corden [74] and Eleni Nastouli [96, 95, 30]

**Funding acquisition, Leadership and supervision, Metadata curation, Sequencing and analysis, and Software and analysis tools:**
Gaia Nebbia [12]

**Funding acquisition, Leadership and supervision, Project administration, Samples and logistics, and Sequencing and analysis:**
Ian Johnston [116]

**Leadership and supervision, Metadata curation, Project administration, Samples and logistics, and Sequencing and analysis:**
Katrina Lythgoe [5], M. Estee Torok [19, 20] and Ian G Goodfellow [24]
**Leadership and supervision, Metadata curation, Project administration, Samples and logistics, and Visualisation:**
Jacqui A Prieto [97, 82] and Kordo Saeed [97, 83]

**Leadership and supervision, Metadata curation, Project administration, Sequencing and analysis, and Software and analysis tools:**
David K Jackson [116]

**Leadership and supervision, Metadata curation, Samples and logistics, Sequencing and analysis, and Visualisation:**
Catherine Houlihan [96, 94]

**Leadership and supervision, Metadata curation, Sequencing and analysis, Software and analysis tools, and Visualisation:**
Dan Frampton [94, 95]

**Metadata curation, Project administration, Samples and logistics, Sequencing and analysis, and Software and analysis tools:**
William L Hamilton [19] and Adam A Witney [41]

**Funding acquisition, Samples and logistics, Sequencing and analysis, and Visualisation:**
Giselda Bucca [101]

**Funding acquisition, Leadership and supervision, Metadata curation, and Project administration:**
Cassie F Pope [40, 41]

**Funding acquisition, Leadership and supervision, Metadata curation, and Samples and logistics:**
Catherine Moore [74]

**Funding acquisition, Leadership and supervision, Metadata curation, and Sequencing and analysis:**
Emma C Thomson [53]

**Funding acquisition, Leadership and supervision, Project administration, and Samples and logistics:**
Ewan M Harrison [116, 102]

**Funding acquisition, Leadership and supervision, Sequencing and analysis, and Visualisation:**
Colin P Smith [101]

**Leadership and supervision, Metadata curation, Project administration, and Sequencing and analysis:**
Fiona Rogan [77]

**Leadership and supervision, Metadata curation, Project administration, and Samples and logistics:**
Shaun M Beckwith [6], Abigail Murray [6], Dawn Singleton [6], Kirstine Eastick [37], Liz A Sheridan [98], Paul Randell [99], Leigh M Jackson [105], Cristina V Ariani [116] and Sónia Gonçalves [116]

**Leadership and supervision, Metadata curation, Samples and logistics, and Sequencing and analysis:**
Derek J Fairley [3, 77], Matthew W Loose [18] and Joanne Watkins [74]

**Leadership and supervision, Metadata curation, Samples and logistics, and Visualisation:**
Samuel Moses [25, 106]

**Leadership and supervision, Metadata curation, Sequencing and analysis, and Software and analysis tools:**
Sam Nicholls [43], Matthew Bull [74] and Roberto Amato [116]

**Leadership and supervision, Project administration, Samples and logistics, and Sequencing and analysis:**
Darren L Smith [36, 65, 66]

**Leadership and supervision, Sequencing and analysis, Software and analysis tools, and Visualisation:**
David M Aanensen [14, 116] and Jeffrey C Barrett [116]

**Metadata curation, Project administration, Samples and logistics, and Sequencing and analysis:**
Dinesh Aggarwal [20, 116, 70], James G Shepherd [53], Martin D Curran [71] and Surendra Parmar [71]

**Metadata curation, Project administration, Sequencing and analysis, and Software and analysis tools:**
Matthew D Parker [109]

**Metadata curation, Samples and logistics, Sequencing and analysis, and Software and analysis tools:**
Catryn Williams [74]

**Metadata curation, Samples and logistics, Sequencing and analysis, and Visualisation:**
Sharon Glaysher [68]

**Metadata curation, Sequencing and analysis, Software and analysis tools, and Visualisation:**
Anthony P Underwood [14, 116], Matthew Bashton [36, 65], Nicole Pacchiarini [74], Katie F Loveson [84] and Matthew Byott [95, 96]

**Project administration, Sequencing and analysis, Software and analysis tools, and Visualisation:**
Alessandro M Carabelli [20]

**Funding acquisition, Leadership and supervision, and Metadata curation:**
Kate E Templeton [56, 104]

**Funding acquisition, Leadership and supervision, and Project administration:**
Thushan I de Silva [109], Dennis Wang [109], Cordelia F Langford [116] and John Sillitoe [116]

**Funding acquisition, Leadership and supervision, and Samples and logistics:**
Rory N Gunson [55]

**Funding acquisition, Leadership and supervision, and Sequencing and analysis:**
Simon Cottrell [74], Justin O'Grady [75, 103] and Dominic Kwiatkowski [116, 108]

**Leadership and supervision, Metadata curation, and Project administration:**
Patrick J Lillie [37]

**Leadership and supervision, Metadata curation, and Samples and logistics:**
Nicholas Cortes [33], Nathan Moore [33], Claire Thomas [33], Phillipa J Burns [37], Tabitha W Mahungu [80] and Steven Liggett [86]

**Leadership and supervision, Metadata curation, and Sequencing and analysis:**
Angela H Beckett [13, 81] and Matthew TG Holden [73]

**Leadership and supervision, Project administration, and Samples and logistics:**
Lisa J Levett [34], Husam Osman [70, 35] and Mohammed O Hassan-Ibrahim [99]

**Leadership and supervision, Project administration, and Sequencing and analysis:**
David A Simpson [77]

**Leadership and supervision, Samples and logistics, and Sequencing and analysis:**
Meera Chand [72], Ravi K Gupta [102], Alistair C Darby [107] and Steve Paterson [107]

**Leadership and supervision, Sequencing and analysis, and Software and analysis tools:**
Oliver G Pybus [23], Erik M Volz [39], Daniela de Angelis [52], David L Robertson [53], Andrew J Page [75] and Inigo Martincorena [116]

**Leadership and supervision, Sequencing and analysis, and Visualisation:**
Louise Aigrain [116] and Andrew R Bassett [116]

**Metadata curation, Project administration, and Samples and logistics:**
Nick Wong [50], Yusri Taha [89], Michelle J Erkiert [99] and Michael H Spencer Chapman [116, 102]

**Metadata curation, Project administration, and Sequencing and analysis:**
Rebecca Dewar [56] and Martin P McHugh [56, 111]

**Metadata curation, Project administration, and Software and analysis tools:**
Siddharth Mookerjee [38, 57]

**Metadata curation, Project administration, and Visualisation:**
Stephen Aplin [97], Matthew Harvey [97], Thea Sass [97], Helen Umpleby [97] and Helen Wheeler [97]

**Metadata curation, Samples and logistics, and Sequencing and analysis:**
James P McKenna [3], Ben Warne [9], Joshua F Taylor [22], Yasmin Chaudhry [24], Rhys Izuagbe [24], Aminu S Jahun [24], Gregory R Young [36, 65], Claire McMurray [43], Clare M McCann [65, 66], Andrew Nelson [65, 66] and Scott Elliott [68]

**Metadata curation, Samples and logistics, and Visualisation:**
Hannah Lowe [25]

**Metadata curation, Sequencing and analysis, and Software and analysis tools:**
Anna Price [11], Matthew R Crown [65], Sara Rey [74], Sunando Roy [96] and Ben Temperton [105]

**Metadata curation, Sequencing and analysis, and Visualisation:**

Sharif Shaaban [73] and Andrew R Hesketh [101]

**Project administration, Samples and logistics, and Sequencing and analysis:**
Kenneth G Laing [41], Irene M Monahan [41] and Judith Heaney [95, 96, 34]

**Project administration, Samples and logistics, and Visualisation:**
Emanuela Pelosi [97], Siona Silviera [97] and Eleri Wilson-Davies [97]

**Samples and logistics, Software and analysis tools, and Visualisation:**
Helen Fryer [5]

**Sequencing and analysis, Software and analysis tools, and Visualization:**
Helen Adams [4], Louis du Plessis [23], Rob Johnson [39], William T Harvey [53, 42], Joseph Hughes [53], Richard J Orton [53], Lewis G Spurgin [59], Yann Bourgeois [81], Chris Ruis [102], Áine O'Toole [104], Marina Gourtovaia [116] and Theo Sanderson [116]

**Funding acquisition, and Leadership and supervision:**
Christophe Fraser [5], Jonathan Edgeworth [12], Judith Breuer [96, 29], Stephen L Michell [105] and John A Todd [115]

**Funding acquisition, and Project administration:**
Michaela John [10] and David Buck [115]

**Leadership and supervision, and Metadata curation:**
Kavitha Gajee [37] and Gemma L Kay [75]

**Leadership and supervision, and Project administration:**
Sharon J Peacock [20, 70] and David Heyburn [74]

**Leadership and supervision, and Samples and logistics:**
Katie Kitchman [37], Alan McNally [43, 93], David T Pritchard [50], Samir Dervisevic [58], Peter Muir [70], Esther Robinson [70, 35], Barry B Vipond [70], Newara A Ramadan [78], Christopher Jeanes [90], Danni Weldon [116], Jana Catalan [118] and Neil Jones [118]

**Leadership and supervision, and Sequencing and analysis:**
Ana da Silva Filipe [53], Chris Williams [74], Marc Fuchs [77], Julia Miskelly [77], Aaron R Jeffries [105], Karen Oliver [116] and Naomi R Park [116]

**Metadata curation, and Samples and logistics:**
Amy Ash [1], Cherian Koshy [1], Magdalena Barrow [7], Sarah L Buchan [7], Anna Mantzouratou [7], Gemma Clark [15], Christopher W Holmes [16], Sharon Campbell [17], Thomas Davis [21], Ngee Keong Tan [22], Julianne R Brown [29], Kathryn A Harris [29, 2], Stephen P Kidd [33], Paul R Grant [34], Li Xu-McCrae [35], Alison Cox [38, 63], Pinglawathee Madona [38, 63], Marcus Pond [38, 63], Paul A Randell [38, 63], Karen T Withell [48], Cheryl Williams [51], Clive Graham [60], Rebecca Denton-Smith [62], Emma Swindells [62], Robyn Turnbull [62], Tim J Sloan [67], Andrew Bosworth [70, 35], Stephanie Hutchings [70], Hannah M Pymont [70], Anna Casey [76], Liz Ratcliffe [76], Christopher R Jones [79, 105], Bridget A Knight [79, 105], Tanzina Haque [80], Jennifer Hart [80], Dianne Irish-Tavares [80], Eric Witele [80], Craig Mower [86], Louisa K Watson [86], Jennifer Collins [89], Gary Eltringham [89], Dorian Crudgington [98], Ben Macklin [98], Miren Iturriza-Gomara [107], Anita O Lucaci [107] and Patrick C McClure [113]

**Metadata curation, and Sequencing and analysis:**

Matthew Carlile [18], Nadine Holmes [18], Christopher Moore [18], Nathaniel Storey [29], Stefan Rooke [73], Gonzalo Yebra [73], Noel Craine [74], Malorie Perry [74], Nabil-Fareed Alikhan [75], Stephen Bridgett [77], Kate F Cook [84], Christopher Fearn [84], Salman Goudarzi [84], Ronan A Lyons [88], Thomas Williams [104], Sam T Haldenby [107], Jillian Durham [116] and Steven Leonard [116]

**Metadata curation, and Software and analysis tools:**
Robert M Davies [116]

**Project administration, and Samples and logistics:**
Rahul Batra [12], Beth Blane [20], Moira J Spyer [30, 95, 96], Perminder Smith [32, 112], Mehmet Yavus [85, 109], Rachel J Williams [96], Adhyana IK Mahanama [97], Buddhini Samaraweera [97], Sophia T Girgis [102], Samantha E Hansford [109], Angie Green [115], Charlotte Beaver [116], Katherine L Bellis [116, 102], Matthew J Dorman [116], Sally Kay [116], Liam Prestwood [116] and Shavanthi Rajatileka [116]

**Project administration, and Sequencing and analysis:**
Joshua Quick [43]

**Project administration, and Software and analysis tools:**
Radoslaw Poplawski [43]

**Samples and logistics, and Sequencing and analysis:**
Nicola Reynolds [8], Andrew Mack [11], Arthur Morriss [11], Thomas Whalley [11], Bindi Patel [12], Iliana Georgana [24], Myra Hosmillo [24], Malte L Pinckert [24], Joanne Stockton [43], John H Henderson [65], Amy Hollis [65], William Stanley [65], Wen C Yew [65], Richard Myers [72], Alicia Thornton [72], Alexander Adams [74], Tara Annett [74], Hibo Asad [74], Alec Birchley [74], Jason Coombes [74], Johnathan M Evans [74], Laia Fina [74], Bree Gatica-Wilcox [74], Lauren Gilbert [74], Lee Graham [74], Jessica Hey [74], Ember Hilvers [74], Sophie Jones [74], Hannah Jones [74], Sara Kumziene-Summerhayes [74], Caoimhe McKerr [74], Jessica Powell [74], Georgia Pugh [74], Sarah Taylor [74], Alexander J Trotter [75], Charlotte A Williams [96], Leanne M Kermack [102], Benjamin H Foulkes [109], Marta Gallis [109], Hailey R Hornsby [109], Stavroula F Louka [109], Manoj Pohare [109], Paige Wolverson [109], Peijun Zhang [109], George MacIntyre-Cockett [115], Amy Trebes [115], Robin J Moll [116], Lynne Ferguson [117], Emily J Goldstein [117], Alasdair Maclean [117] and Rachael Tomb [117]

**Samples and logistics, and Software and analysis tools:**
Igor Starinskij [53]

**Sequencing and analysis, and Software and analysis tools:**
Laura Thomson [5], Joel Southgate [11, 74], Moritz UG Kraemer [23], Jayna Raghwani [23], Alex E Zarebski [23], Olivia Boyd [39], Lily Geidelberg [39], Chris J Illingworth [52], Chris Jackson [52], David Pascall [52], Sreenu Vattipally [53], Timothy M Freeman [109], Sharon N Hsu [109], Benjamin B Lindsey [109], Keith James [116], Kevin Lewis [116], Gerry Tonkin-Hill [116] and Jaime M Tovar-Corona [116]

**Sequencing and analysis, and Visualisation:**
MacGregor Cox [20]

**Software and analysis tools, and Visualisation:**
Khalil Abudahab [14, 116], Mirko Menegazzo [14], Ben EW Taylor MEng [14, 116], Corin A Yeats [14], Afrida Mukaddas [53], Derek W Wright [53], Leonardo de Oliveira Martins [75], Rachel Colquhoun [104], Verity Hill [104], Ben Jackson [104], JT McCrone [104], Nathan Medd [104], Emily Scher [104] and Jon-Paul Keatley [116]

**Leadership and supervision:**

Tanya Curran [3], Sian Morgan [10], Patrick Maxwell [20], Ken Smith [20], Sahar Eldirdiri [21], Anita Kenyon [21], Alison H Holmes [38, 57], James R Price [38, 57], Tim Wyatt [69], Alison E Mather [75], Timofey Skvortsov [77] and John A Hartley [96]

**Metadata curation:**

Martyn Guest [11], Christine Kitchen [11], Ian Merrick [11], Robert Munn [11], Beatrice Bertolusso [33], Jessica Lynch [33], Gabrielle Vernet [33], Stuart Kirk [34], Elizabeth Wastnedge [56], Rachael Stanley [58], Giles Idle [64], Declan T Bradley [69, 77], Jennifer Poyner [79] and Matilde Mori [110]

**Project administration:**

Owen Jones [11], Victoria Wright [18], Ellena Brooks [20], Carol M Churcher [20], Mireille Fragakis [20], Katerina Galai [20, 70], Andrew Jermy [20], Sarah Judges [20], Georgina M McManus [20], Kim S Smith [20], Elaine Westwick [20], Stephen W Attwood [23], Frances Bolt [38, 57], Alisha Davies [74], Elen De Lacy [74], Fatima Downing [74], Sue Edwards [74], Lizzie Meadows [75], Sarah Jeremiah [97], Nikki Smith [109] and Luke Foulser [116]

**Samples and logistics:**

Themoula Charalampous [12, 46], Amita Patel [12], Louise Berry [15], Tim Boswell [15], Vicki M Fleming [15], Hannah C Howson-Wells [15], Amelia Joseph [15], Manjinder Khakh [15], Michelle M Lister [15], Paul W Bird [16], Karlie Fallon [16], Thomas Helmer [16], Claire L McMurray [16], Mina Odedra [16], Jessica Shaw [16], Julian W Tang [16], Nicholas J Willford [16], Victoria Blakey [17], Veena Raviprakash [17], Nicola Sheriff [17], Lesley-Anne Williams [17], Theresa Feltwell [20], Luke Bedford [26], James S Cargill [27], Warwick Hughes [27], Jonathan Moore [28], Susanne Stonehouse [28], Laura Atkinson [29], Jack CD Lee [29], Dr Divya Shah [29], Adela Alcolea-Medina [32, 112], Natasha Ohemeng-Kumi [32, 112], John Ramble [32, 112], Jasveen Sehmi [32, 112], Rebecca Williams [33], Wendy Chatterton [34], Monika Pusok [34], William Everson [37], Anibolina Castigador [44], Emily Macnaughton [44], Kate El Bouzidi [45], Temi Lampejo [45], Malur Sudhanva [45], Cassie Breen [47], Graciela Sluga [48], Shazaad SY Ahmad [49, 70], Ryan P George [49], Nicholas W Machin [49, 70], Debbie Binns [50], Victoria James [50], Rachel Blacow [55], Lindsay Coupland [58], Louise Smith [59], Edward Barton [60], Debra Padgett [60], Garren Scott [60], Aidan Cross [61], Mariyam Mirfenderesky [61], Jane Greenaway [62], Kevin Cole [64], Phillip Clarke [67], Nichola Duckworth [67], Sarah Walsh [67], Kelly Bicknell [68], Robert Impey [68], Sarah Wyllie [68], Richard Hopes [70], Chloe Bishop [72], Vicki Chalker [72], Ian Harrison [72], Laura Gifford [74], Zoltan Molnar [77], Cressida Auckland [79], Cariad Evans [85, 109], Kate Johnson [85, 109], David G Partridge [85, 109], Mohammad Raza [85, 109], Paul Baker [86], Stephen Bonner [86], Sarah Essex [86], Leanne J Murray [86], Andrew I Lawton [87], Shirelle Burton-Fanning [89], Brendan AI Payne [89], Sheila Waugh [89], Andrea N Gomes [91], Maimuna Kimuli [91], Darren R Murray [91], Paula Ashfield [92], Donald Dobie [92], Fiona Ashford [93], Angus Best [93], Liam Crawford [93], Nicola Cumley [93], Megan Mayhew [93], Oliver Megram [93], Jeremy Mirza [93], Emma Moles-Garcia [93], Benita Percival [93], Megan Driscoll [96], Leah Ensell [96], Helen L Lowe [96], Laurentiu Maftei [96], Matteo Mondani [96], Nicola J Chaloner [99], Benjamin J Cogger [99], Lisa J Easton [99], Hannah Huckson [99], Jonathan Lewis [99], Sarah Lowdon [99], Cassandra S Malone [99], Florence Munemo [99], Manasa Mutingwende [99], Roberto Nicodemi [99], Olga Podplomyk [99], Thomas Somassa [99], Andrew Beggs [100], Alex Richter [100], Claire Cormie [102], Joana Dias [102], Sally Forrest [102], Ellen E Higginson [102], Mailis Maes [102], Jamie Young [102], Rose K Davidson [103], Kathryn A Jackson [107], Lance Turtle [107], Alexander J Keeley [109], Jonathan Ball [113], Timothy Byaruhanga [113], Joseph G Chappell [113], Jayasree Dey [113], Jack D Hill [113], Emily J Park [113], Arezou Fanaie [114], Rachel A Hilson [114], Geraldine Yaze [114] and Stephanie Lo [116]

**Sequencing and analysis:**

Safiah Afifi [10], Robert Beer [10], Joshua Maksimovic [10], Kathryn McCluggage [10], Karla Spellman [10], Catherine Bresner [11], William Fuller [11], Angela Marchbank [11], Trudy Workman [11], Ekaterina Shelest [13, 81], Johnny Debebe [18], Fei Sang [18], Marina Escalera Zamudio [23], Sarah Francois [23], Bernardo Gutierrez [23], Tetyana I Vasylyeva [23], Flavia Flaviani [31], Manon Ragonnet-Cronin [39], Katherine L Smollett [42], Alice Broos [53], Daniel Mair [53], Jenna Nichols [53], Kyriaki Nomikou [53], Lily Tong [53], Ioulia Tsatsani [53], Sarah

O'Brien [54], Steven Rushton [54], Roy Sanderson [54], Jon Perkins [55], Seb Cotton [56], Abbie Gallagher [56], Elias Allara [70, 102], Clare Pearson [70, 102], David Bibby [72], Gavin Dabrera [72], Nicholas Ellaby [72], Eileen Gallagher [72], Jonathan Hubb [72], Angie Lackenby [72], David Lee [72], Nikos Manesis [72], Tamyo Mbisa [72], Steven Platt [72], Katherine A Twohig [72], Mari Morgan [74], Alp Aydin [75], David J Baker [75], Ebenezer Foster-Nyarko [75], Sophie J Prosolek [75], Steven Rudder [75], Chris Baxter [77], Sílvia F Carvalho [77], Deborah Lavin [77], Arun Mariappan [77], Clara Radulescu [77], Aditi Singh [77], Miao Tang [77], Helen Morcrette [79], Nadua Bayzid [96], Marius Cotic [96], Carlos E Balcazar [104], Michael D Gallagher [104], Daniel Maloney [104], Thomas D Stanton [104], Kathleen A Williamson [104], Robin Manley [105], Michelle L Michelsen [105], Christine M Sambles [105], David J Studholme [105], Joanna Warwick-Dugdale [105], Richard Eccles [107], Matthew Gemmell [107], Richard Gregory [107], Margaret Hughes [107], Charlotte Nelson [107], Lucille Rainbow [107], Edith E Vamos [107], Hermione J Webster [107], Mark Whitehead [107], Claudia Wierzbicki [107], Adrienn Angyal [109], Luke R Green [109], Max Whiteley [109], Emma Betteridge [116], Iraad F Bronner [116], Ben W Farr [116], Scott Goodwin [116], Stefanie V Lensing [116], Shane A McCarthy [116, 102], Michael A Quail [116], Diana Rajan [116], Nicholas M Redshaw [116], Carol Scott [116], Lesley Shirley [116] and Scott AJ Thurston [116]

**Software and analysis tools:**
Will Rowe [43], Amy Gaskin [74], Thanh Le-Viet [75], James Bonfield [116], Jennifier Liddle [116] and Andrew Whitwham [116]

**1** Barking, Havering and Redbridge University Hospitals NHS Trust, **2** Barts Health NHS Trust, **3** Belfast Health & Social Care Trust, **4** Betsi Cadwaladr University Health Board, **5** Big Data Institute, Nuffield Department of Medicine, University of Oxford, **6** Blackpool Teaching Hospitals NHS Foundation Trust, **7** Bournemouth University, **8** Cambridge Stem Cell Institute, University of Cambridge, **9** Cambridge University Hospitals NHS Foundation Trust, **10** Cardiff and Vale University Health Board, **11** Cardiff University, **12** Centre for Clinical Infection and Diagnostics Research, Department of Infectious Diseases, Guy's and St Thomas' NHS Foundation Trust, **13** Centre for Enzyme Innovation, University of Portsmouth, **14** Centre for Genomic Pathogen Surveillance, University of Oxford, **15** Clinical Microbiology Department, Queens Medical Centre, Nottingham University Hospitals NHS Trust, **16** Clinical Microbiology, University Hospitals of Leicester NHS Trust, **17** County Durham and Darlington NHS Foundation Trust, **18** Deep Seq, School of Life Sciences, Queens Medical Centre, University of Nottingham, **19** Department of Infectious Diseases and Microbiology, Cambridge University Hospitals NHS Foundation Trust, **20** Department of Medicine, University of Cambridge, **21** Department of Microbiology, Kettering General Hospital, **22** Department of Microbiology, South West London Pathology, **23** Department of Zoology, University of Oxford, **24** Division of Virology, Department of Pathology, University of Cambridge, **25** East Kent Hospitals University NHS Foundation Trust, **26** East Suffolk and North Essex NHS Foundation Trust, **27** East Sussex Healthcare NHS Trust**, 28** Gateshead Health NHS Foundation Trust, **29** Great Ormond Street Hospital for Children NHS Foundation Trust, **30** Great Ormond Street Institute of Child Health (GOS ICH), University College London (UCL), **31** Guy's and St. Thomas' Biomedical Research Centre, **32** Guy's and St. Thomas' NHS Foundation Trust, **33** Hampshire Hospitals NHS Foundation Trust, **34** Health Services Laboratories, **35** Heartlands Hospital, Birmingham, **36** Hub for Biotechnology in the Built Environment, Northumbria University, **37** Hull University Teaching Hospitals NHS Trust, **38** Imperial College Healthcare NHS Trust, **39** Imperial College London, **40** Infection Care Group, St George's University Hospitals NHS Foundation Trust, **41** Institute for Infection and Immunity, St George's University of London, **42** Institute of Biodiversity, Animal Health & Comparative Medicine, **43** Institute of Microbiology and Infection, University of Birmingham, **44** Isle of Wight NHS Trust, **45** King's College Hospital NHS Foundation Trust, **46** King's College London, **47** Liverpool Clinical Laboratories, **48** Maidstone and Tunbridge Wells NHS Trust, **49** Manchester University NHS Foundation Trust, **50** Microbiology Department, Buckinghamshire Healthcare NHS Trust, **51** Microbiology, Royal Oldham Hospital, **52** MRC Biostatistics Unit, University of Cambridge, **53** MRC-University of Glasgow Centre for Virus Research, **54** Newcastle University, **55** NHS Greater Glasgow and Clyde, **56** NHS Lothian, **57** NIHR Health Protection Research Unit in HCAI and AMR, Imperial College London, **58** Norfolk and Norwich University Hospitals NHS Foundation Trust, **59** Norfolk County Council, **60** North Cumbria Integrated Care NHS Foundation Trust, **61** North Middlesex University Hospital NHS Trust, **62** North Tees and Hartlepool NHS Foundation Trust, **63** North West London Pathology, **64** Northumbria Healthcare NHS Foundation Trust, **65** Northumbria University, **66** NU-OMICS, Northumbria University, **67** Path Links, Northern Lincolnshire and Goole NHS Foundation Trust, **68** Portsmouth Hospitals University NHS Trust, **69** Public Health Agency, Northern Ireland, **70** Public Health England, **71** Public Health England, Cambridge, **72** Public Health England,

Colindale, **73** Public Health Scotland, **74** Public Health Wales, **75** Quadram Institute Bioscience, **76** Queen Elizabeth Hospital, Birmingham, **77** Queen's University Belfast, **78** Royal Brompton and Harefield Hospitals, **79** Royal Devon and Exeter NHS Foundation Trust, **80** Royal Free London NHS Foundation Trust, **81** School of Biological Sciences, University of Portsmouth, **82** School of Health Sciences, University of Southampton, **83** School of Medicine, University of Southampton, **84** School of Pharmacy & Biomedical Sciences, University of Portsmouth, **85** Sheffield Teaching Hospitals NHS Foundation Trust, **86** South Tees Hospitals NHS Foundation Trust, **87** Southwest Pathology Services, **88** Swansea University, **89** The Newcastle upon Tyne Hospitals NHS Foundation Trust, **90** The Queen Elizabeth Hospital King's Lynn NHS Foundation Trust, **91** The Royal Marsden NHS Foundation Trust, **92** The Royal Wolverhampton NHS Trust, **93** Turnkey Laboratory, University of Birmingham, **94** University College London Division of Infection and Immunity**, 95** University College London Hospital Advanced Pathogen Diagnostics Unit**, 96** University College London Hospitals NHS Foundation Trust, **97** University Hospital Southampton NHS Foundation Trust, **98** University Hospitals Dorset NHS Foundation Trust, **99** University Hospitals Sussex NHS Foundation Trust, **100** University of Birmingham, **101** University of Brighton, **102** University of Cambridge, **103** University of East Anglia, **104** University of Edinburgh, **105** University of Exeter, **106** University of Kent, **107** University of Liverpool, **108** University of Oxford, **109** University of Sheffield, **110** University of Southampton, **111** University of St Andrews, **112** Viapath, Guy's and St Thomas' NHS Foundation Trust, and King's College Hospital NHS Foundation Trust, **113** Virology, School of Life Sciences, Queens Medical Centre, University of Nottingham, **114** Watford General Hospital, **115** Wellcome Centre for Human Genetics, Nuffield Department of Medicine, University of Oxford, **116** Wellcome Sanger Institute, **117** West of Scotland Specialist Virology Centre, NHS Greater Glasgow and Clyde, **118** Whittington Health NHS Trust
