## Supplementary material for "The origins and molecular evolution of SARS-CoV-2 lineage B.1.1.7 in the UK": GISAID Acknowledgements table

We gratefully acknowledge the following Authors from the Originating laboratories responsible for obtaining the specimens, as well as the Submitting laboratories where the genome data were generated and shared via GISAID, on which this research is based.

All Submitters of data may be contacted directly via www.gisaid.org

Authors are sorted alphabetically.

Acknowledgement EPI_SET Identifier: EPI_SET_20220223fy

| Accession ID | Originating Laboratory | Submitting Laboratory | Authors |
|---|---|---|---|
| EPI_ISL_8258238 | "HI Dept. of Health, State Laboratories Division" | Centers for Disease Control and Prevention Division of Viral Diseases, Pathogen Discovery | Alex Burgin; Ben Rambo-Martin; Clinton Paden; Dakota Howard; Dave Wentworth; Dhwani Batra; Jasmine Padilla; Joseph Madden; Justin Lee; Kristen Knipe; Kristine Lacek; Mark Burroughs; Matthew Schmerer; Meghan Bentz; Mili Sheth; Peter Cook; Sam Shepard; Sarah Nobles; Vivien Dugan; Yvette Unoarumhi |
| EPI_ISL_1226072 | "NYSDOH Wadsworth Center, Virology Lab" | Genomics and Discovery, Respiratory Viruses Branch, Division of Viral Diseases, Centers for Disease Control and Prevention | Anna Montmayeur; Anna Uehara; Ben L. Rambo-Martin; Clinton R. Paden; Dhwani Batra; Haibin Wang; Jasmine Padilla; Jing Zhang; Justin Lee; Katie Dillon; Krista Queen; Kristen Knipe; Kristine Lacek; Lori Rowe; Mark Burroughs; Matthew Schmerer; Mili Sheth; Peter W. Cook; Rachel Marine; Sam Shepard; Sarah Nobles; Shoshona Le; Suxiang Tong; Yan Li; Ying Tao |
| EPI_ISL_1372414 | 1.AO Universitaria 'S. Giovanni di Dio e Ruggi D'Aragona, Scuola Medica Salernitana' Hospital / 2.UOC di Virologia e Microbiologia, Università della Campania 'L. Vanvitelli' / 3.AO Universitaria 'Federico II' Napoli Hospital / 4.AORN 'San Giuseppe Moscati' Avellino Hospital / 5.AO 'San Pio - presidio G. Rummo' Benevento Hospital / 6.AO 'Sant'Anna e San Sebastiano' Caserta Hospital / 7.PO 'Maria Santissima Addolorata' Eboli Hospital / 8.Biogem Istituto di Ricerche Genetiche / 9. U.O.C. di Genetica Medica e di Laboratorio A.O.R.N., Azienda Ospedaliera di Rilievo Nazionale Antonio Cardarelli, Napoli / 10. Centro di riferimento Oncologico della Basilicata (IRCCS-CROB), Rionero in Vulture (PZ) / 11. Presidio Ospedaliero di Agropoli, Agropoli (SA). | 1. Genome Research Center for Health (CRGS) / 2. Laboratory of Molecular Medicine and Genomics(LMMGe) / 3. Center for Research in Pure and Applied Mathematics (CRMPA) | Alessandro Weisz (Corresponding Author); Alessia Cossu; Andreina Baj; Aniello Gentile; Annamaria Salvati; Antonello Saccomanno; Arnolfo Petruzziello; Assunta Sellitto; Caivano Antonella; Carlo Ferravante; D'Auria Fiorella; De Luca Luciana.; Domenico Memoli; Domenico Palumbo; Edmondo Adorisio; Elena Alexandrova; Emilia Vaccaro; Fausto Sessa; Francesca Marciano; Francesca Rizzo (Corresponding Author); Francesco Curcio; Gianluigi Franci; Giorgio Dirani; Giorgio Giurato (Corresponding Author); Giovanni Nassa; Giovanni Pecoraro; Giuseppe Fenza; Giuseppe Portella; Giusy Giordano; Gregorio Goffredi; Ilaria Terenzi; Jessica Lamberti; Maddalena Schioppa; Marcello Ametrano; Maria Grazia Foti; Maria Landi; Marianna Scrima; Mariano Fiorenza; Mariarosaria Ingino; Marina Tarsitano; Massimiliano Chetta; Massimiliano Galdiero; Maurizio Fumi; Michela Iacobellis; Michele Caraglia; Michele Cennamo; Morena D'Avenia; Oriana Strianese; Pasquale Pagliano; Rita Greco; Roberta Tarallo; Rosanna Piluscio; Silvia Zanoli; Simona Semprini; Sonia Amabile; Stefania Marzinotto; Teresa Rocco; Valeria Mirici Cappa; Vincenzo Rocco; Viola Melone; Vittoria Letizia; Vittorio Sambri; Ylenia D'Agostino |
| EPI_ISL_8296777, EPI_ISL_8297168, EPI_ISL_8297432 | 35259 CS SAINT EXUPERY 1 | CNR Virus des Infections Respiratoires - France SUD | Antonin Bal; Bruno Lina; Bruno Simon; Gregory Destras; Gwendolyne Burfin; Hadrien Regue; Laurence Josset; Martine Valette; Quentin Semanas |
| EPI_ISL_7954432, EPI_ISL_7954436 | AGES-Institute for medical Microbiology and Hygiene Vienna | AGES-Institute for medical Microbiology and Hygiene Vienna | Alexander Indra; Florian Heger; Julia Klikovits; Kathrin Lippert; Magdalena Purker; Marion Blaschitz; Patrick Hyden; Peter Hufnagl |
| EPI_ISL_1663507 | AIIMS, Patna | Institute of Life Sciences - INSACOG | Ajay Parida; Amol M. Kanampalliwar; Arup Ghosh; Atimukta Jha; INSACOG Consortium; Punit Prasad; Rajeeb Swain; Rupesh Dash; Safal Walia; Shifu Aggarwal; Sunil K. Raghav |
| EPI_ISL_8247338, EPI_ISL_8286643 | ALAB | WSSE w Warszawie | Dorota Wagrocka - Roczniak |
| EPI_ISL_7803621, EPI_ISL_8164892, EPI_ISL_8164991 | ALAB Siedlce | WSSE w Warszawie | Dorota Wagrocka - Roczniak |
| EPI_ISL_1820928 | ANALIZAR LABORATORIO CLINICO AUTOMATIZADO - SYNLAB | Instituto Nacional de Salud- Dirección de Investigación en Salud Pública | Carlos Franco-Muñoz; Carmen Osorio; Christian Romero; Diana Malo; Diego A. Álvarez-Díaz; Diego Andrés Prada; Gerardo Santamaría; Hector Alejandro Ruiz-Moreno; Jhonnatan Reales-González; Jorge Rivera; Juan Camilo Martinez; Julian Naizaque; Katherine Laiton-Donato; Lisseth Pardo; Magdalena Wiesner; Marcela Mercado-Reyes; Maria T. Herrera-Sepúlveda; Marta Lopez Blanco; Martha Lucia Ospina Martinez; Paola Rojas; Patricia del Portillo; Sergio Gomez; Sheryll Corchuelo; Ángela Alarcon Cruz |
| EPI_ISL_2231597, EPI_ISL_2231639, EPI_ISL_2441209, EPI_ISL_7235183, EPI_ISL_7468068 | AP SSO | CSIR-Centre for Cellular and Molecular Biology-INSACOG | Amareshwar Vodapalli; Ara Sreenivas; Archana Bharadwaj Siva; B Himasri; Blessy B John; Divya Tej Sowpati; Jandhyala Sai Krishna; Karthik Bharadwaj Tallapaka; Lamuk Zaveri; Malini Nemalikanti; Onkar Kulkarni; Payel Mukherjee; Priya Nurkurthy; Rakesh K Mishra; Sharath Chandra Thota; Shreekant Verma; Sofia Banu; Sreelekshmi MS; Sumedha Avadhanula; Surabhi Srivastava; Tulasi Nagabandi; Valli Nagalakshmi Undamatla; Vidhyadhari Methuku; Viswagithe S L |
| EPI_ISL_2451784 | AR Dept. of Health-PHL, Molecular Diagnostics | Centers for Disease Control and Prevention Division of Viral Diseases, Pathogen Discovery | Alison Laufer Halpin; Ben L. Rambo-Martin; Clinton R. Paden; Dakota Howard; Darlene Wagner; Dave Wentworth; Dhwani Batra; Jasmine Padilla; Justin Lee; Katie Dillon; Krista Queen; Kristen Knipe; Kristine Lacek; Mark Burroughs; Matthew Schmerer; Mili Sheth; Peter Cook; Sam Shepard; Sarah Nobles; Shoshona Le; Suxiang Tong; Vivien Dugan; Yvette Unoarumhi |
| EPI_ISL_6963529 | AREA DE SALUD GOICOECHEA 2 - CLINICA DR. JIMENEZ NUÑEZ | Inciensa, Instituto Costarricense de Investigación y Enseñanza en Nutrición y Salud | Adriana Godínez; Claudio Soto-Garita; Estela Cordero; Francisco Duarte; Hebleen Porras; José Luis Vargas; Mariela Gutiérrez & Sofia Herrera& Juan Carlos Cartes Parra; Melany Calderón |
| EPI_ISL_1298997 | ASL Napoli 1 Centro | AMES Centro Polidiagnostico Strumentale S.r.l. | "Giovanni Savarese; Antonella Di Carlo; Antonio Fico"; Eloisa Evangelista; Luigi D'Amore; Luisa Circelli; Maurizio D'Amora; Monica Ianniello; Nadia Petrillo; Raffaella Ruggiero; Roberto Sirica |
| EPI_ISL_4659138 | AUSTRAL-omics, UACh | AUSTRAL-omics, UACh | Andrea Silva; Carolina Encina; Cristian Molina; Daniela Plaza; Luis Guzmán; Suany Quesada |
| EPI_ISL_2376311, EPI_ISL_2376334, EPI_ISL_2377043 | AZDelta | AZ Delta Medical Laboratories in Roeselare, Belgium | Dieter De Smet; Geert Martens; Merijn Vanhee; on behalf of AZ Delta COVID-19 Genomics core (member of Genomic surveillance of SARS-CoV-2 in Belgium network) |
| EPI_ISL_1561156, EPI_ISL_1562503, EPI_ISL_1650631, EPI_ISL_1736859, EPI_ISL_2038982, EPI_ISL_2041726, EPI_ISL_2147681, EPI_ISL_2181374, EPI_ISL_2202297, EPI_ISL_2204595, EPI_ISL_2204609, EPI_ISL_2204778, EPI_ISL_2244480, EPI_ISL_2280451, EPI_ISL_2281160, EPI_ISL_2283826, EPI_ISL_2321232, EPI_ISL_2321237, EPI_ISL_2368269, EPI_ISL_2368887, EPI_ISL_2368979, EPI_ISL_2370661, EPI_ISL_2489151, EPI_ISL_2489335, EPI_ISL_2489338, EPI_ISL_2489524, EPI_ISL_2489669, EPI_ISL_2489687, EPI_ISL_2489725, EPI_ISL_2489737, EPI_ISL_2528472, EPI_ISL_2528673, EPI_ISL_2528961, EPI_ISL_4535536, EPI_ISL_5111690, EPI_ISL_5372243, EPI_ISL_5512262, EPI_ISL_5514079, EPI_ISL_5519023, EPI_ISL_5577552, EPI_ISL_6062890, EPI_ISL_6063535, EPI_ISL_6067686, EPI_ISL_6163045, EPI_ISL_6385014, EPI_ISL_6514749, EPI_ISL_6573162, EPI_ISL_6631111, EPI_ISL_6649927, EPI_ISL_7227549, EPI_ISL_7230319, EPI_ISL_7232492, EPI_ISL_7241315, EPI_ISL_7241707, EPI_ISL_7243351, EPI_ISL_7243552, EPI_ISL_7408682, EPI_ISL_7411142, EPI_ISL_7411707, EPI_ISL_7435464, EPI_ISL_7702671, EPI_ISL_7779518, EPI_ISL_7780039, EPI_ISL_7781093, EPI_ISL_7781112, EPI_ISL_7782483, EPI_ISL_7928841, EPI_ISL_7929075, EPI_ISL_7929186, EPI_ISL_7930460, EPI_ISL_7931108, EPI_ISL_7933229, EPI_ISL_8010722, EPI_ISL_8011583, EPI_ISL_8011987, EPI_ISL_8012141, EPI_ISL_8013458, EPI_ISL_8034050, EPI_ISL_8081657, EPI_ISL_8142559, EPI_ISL_8149820, EPI_ISL_8150287, EPI_ISL_8179184, EPI_ISL_8258994, EPI_ISL_8259153, EPI_ISL_8259757, EPI_ISL_8260157, EPI_ISL_8289666, EPI_ISL_8311186, EPI_ISL_8312484, EPI_ISL_8312489 | | | |
| see above | Aegis Sciences Corporation | Centers for Disease Control and Prevention Division of Viral Diseases, Pathogen Discovery | Adrian Paskey; Alec Vest; Benjamin Rambo-Martin; Christopher Gulvick; Clinton Paden; Clinton R. Paden; Cyndi Clark; Dakota Howard; Darlene Wagner; Dhwani Batra; Dillon Nall; Duncan MacCannell; Erisa Sula; Ethan Sanders; Holly Houdeshell; Jason Caravas; Kara Moser; Kristine Lacek; Matthew Hardison; Matthew Schmerer; Ola Kvalvaag; Patrick Campbell; Peter Cook; Peter W. Cook; Rob Case; Scott Sammons; Shatavia Morrison; Shaun Westlund; Tymeckia Kendall; Victoria Caban Figueroa; Vikramsinha Ghorpade; Yvette Unoarumhi |
| EPI_ISL_6775032 | Akademicki Ośrodek Diagnostyki Patomorfologicznej i Genetyczno-Molekularnej Sp. z.o.o. | 1. Academic Center for Pathomorphological and Genetic-Molecular Diagnostics ltd, Bialystok, Poland 2. National Institute of Public Health - National Institute of Hygiene, Warsaw, Poland | Anetta Sulewska; Jacek Nikliński; Janusz Dzięcioł; Joanna Kiśluk; Katarzyna Zacharczuk; Konrad Raczkowski; Magdalena Nowakowska; Małgorzata Sadkowska-Todys; Piotr Karabowicz; Piotr Majewski; Przemysław Biecek. Joanna Reszeć; Radosław Charkiewicz; Tomasz Wołkowicz |
| EPI_ISL_2491489, EPI_ISL_4274093, EPI_ISL_5633696 | Akershus University Hospital, Department for Microbiology and Infectious Disease Control | Norwegian Institute of Public Health, Department of Virology | Atiya R Ali; Debech Nadia; Engebretsen Serina Beate; Garcia Llorente Ignacio; Hilde Elshaug; Hilde Vollan; Jon Bråte; Kamilla Heddeland Instefjord; Karoline Bragstad; Kathrine Stene-Johansen; Line Victoria Moen; Marie Paulsen Madsen; Olav Hungnes; Pedersen Benedikte Nevjen; Rasmus Riis Kopperud |
| EPI_ISL_2101761, EPI_ISL_2158264, EPI_ISL_2158265, EPI_ISL_2158266, EPI_ISL_2158267, EPI_ISL_2421687, EPI_ISL_2421688, EPI_ISL_2421694, EPI_ISL_2421706, EPI_ISL_2421709, EPI_ISL_2422048, EPI_ISL_2422053, EPI_ISL_2422062, EPI_ISL_2422082, EPI_ISL_2454576, EPI_ISL_2455417, EPI_ISL_2455418, EPI_ISL_2458226, EPI_ISL_2458227, EPI_ISL_2467800, EPI_ISL_2467801, EPI_ISL_2467804, EPI_ISL_2473688 | | | |
| see above | Alaska State Virology Laboratory | Alaska State Virology Laboratory | Elva House; Jack Chen; Jacob Zidek; Lisa Smith; Ph.D.; Stephanie DeRonde |
| EPI_ISL_2426234 | All India Institute of Medical Sciences, Delhi | CSIR Institute of Genomics and Integrative Biology | Abhinav Jain; Afra Shamnath; Anjali Bajaj; Arvinden VR; Bani Jolly; COVID CBNAAT CORE GROUP; Kiran Bala; Mercy Rophina; Mohamed Imran; Mohit Kumar Divakar; Nayer Jamshed; Praveen Aggarwal; Rahul C. Bhoyar; Rama Chaudhry; Randeep Guleria; Ritu Gupta; Sridhar Sivasubbu; Subrata Sinha; Urvashi B Singh; Vigneshwar Senthivel; Vinod Scaria |
| EPI_ISL_5990102, EPI_ISL_8281560, EPI_ISL_8281577, EPI_ISL_8281612 | Allergy, Immunology and Cell Biology Unit (AICBU) | Allergy, Immunology and Cell Biology Unit (AICBU) | Ayesha Wijesinghe; Chandima Jeewandara; Deshni Jayathilaka; Dinuka Ariyarathne; Dinuka Ariyaratne; Diyanath Ranasinghe; Dumni Guasinghe; Dumni Gunasinghe; Farha Bary; Gathsaurie Neelika Malavige; Heshan Kuruppu; Tibutius Thanesh |
| EPI_ISL_6914020, EPI_ISL_6914026, EPI_ISL_6914035 | Ampath Laboratories | National Institute for Communicable Diseases of the National Health Laboratory Service | Amoako DG; Bhiman JN; Everatt J; Ismail A; Mahlangu B; Mnguni A; Mohale T; Ntuli N; Scheepers C; Wolter N |
| EPI_ISL_6035992, EPI_ISL_6036134 | Analytica | Institute of Medical Virology | Alexandra Trkola; Annette Audigé; Catharine Aquino; Cyril Shah; Daniel Ehrsam; Gabriela Ziltener; Guido Bloemberg; Hubert Rehrauer; Isabel Stürmer; Joel Wirz; Jon Huder; Jürg Böni; Kevin Steiner; Maria Grünberg; Maryam Zaheri; Michael Huber; Riccarda Capaul; Stefan Schmutz; Verena Kufner; Weihong Qi |
| EPI_ISL_1494722 | Area of Virology, Serology and Virology Division (SAViD), New South Wales Health Pathology Randwick | Virology Research Laboratory; Area of Virology, Serology and Virology Division (SAViD), New South Wales Health Pathology Randwick | Au, J.; Bull, R.; Deveson, I.; Foster, C.; Rawlinson, W.; Ruiz Silva, M.; Van Hal, S. |
| EPI_ISL_5461564, EPI_ISL_7898329, EPI_ISL_8163180 | Arizona State University | Arizona State University | Ajeet Bains; Efrem S. Lim; Joshua LaBaer; LaRinda A. Holland; Matthew F. Smith; Nathaniel Johnson; Nicholas J. Mellor; Rabia Maqsood; Regan A. Sullins; Steven C. Holland; Vel Murugan |
| EPI_ISL_6481752 | Atlanta Veterans Affairs Medical Center | Centers for Disease Control and Prevention Division of Viral Diseases, Pathogen Discovery | Alex Burgin; Alexis Whitmire; Aron Hall; Ben Rambo-Martin; Clinton Paden; Cynthia Lucero-Obusan; Dakota Howard; Dave Wentworth; David Beenhouwer; Dhwani Batra; Diya Surie; Elissa Meites; Evan Goldin; Gilberto Rivera-Dominguez; Jasmine Padilla; Jennifer Verani; Justin Lee; Krista Queen; Kristen Knipe; Kristina Bajema; Kristine Lacek; Maraia Tremarelli; Maria Rodriguez-Barradas; Mark Burroughs; Mark Holodniy; Matthew Schmerer; Meghan Bentz; Meredith McMorrow; Mila Prill; Mili Sheth; Miwako Kobayashi; Peter Cook; Rebecca Dahl; Rosalba Morones; Sam Shepard; Sarah Nobles; Sheldon Brown; Stephanie Schrag; Steve Evener; Suxiang Tong; Vincent Marconi; Vivien Dugan; Yvette Unoarumhi |
| EPI_ISL_6091487, EPI_ISL_7798420 | Austrian Agency for Health and Food Safety (AGES) | Bergthaler laboratory, CeMM Research Center for Molecular Medicine of the Austrian Academy of Sciences | Andreas Bergthaler; Anna Schedl; Bekir Erguner; Benedikt Agerer; Christoph Bock; Fabian Amman; Jan Laine; Lukas Endler; Martin Senekowitsch; Matthew Thornton; Michael Schuster; Michelle Chan; Petr Triska; Thomas Penz |
| EPI_ISL_2438009 | Ayudas diagnosticas SURA | Universidad Nacional de Colombia - Laboratorio Genómico One Health | Andres F. Cardona-Rios; Carlos Franco-Muñoz; Carolina Muñoz-Arango; Celeny Ortiz; Daniel O. Maldonado-Perez; Diego A. Álvarez-Díaz; Hector Alejandro Ruiz-Moreno; Idabely Betancur Ortiz; Jorge E. Osorio; Juan P. Hernandez-Ortiz; Karl A Ciuoderis; Katherine Laiton-Donato; Laura Silvana Perez; Lina M. Hurtado; Marcela Mercado-Reyes; Maria Angélica Maya; Maria Stella López; Rita Almanza Payares; Sandra Ines Cano; |

| | | | |
|---|---|---|---|
| | | | Simón Villegas Velásquez |
| EPI_ISL_2522101 | Azienda Ospedaliera San Camillo Forlanini | INMI Lazzaro Spallanzani IRCCS | A D'Agostino; A Di Caro; B Bartolini; CEM Gruber; E Giombini; F Basile; F Messina; F Santini; G Bonfiglio; G Parisi; M Rueca; MR Capobianchi; O Butera |
| EPI_ISL_2331418, EPI_ISL_2331437 | Azienda Ospedaliero - Universitaria di Modena Policlinico - Virologia e Microbiologia Molecolare | Istituto Zooprofilattico Sperimentale della Lombardia e dell'Emilia Romagna (IZSLER), Risk Analysis and Genomic Epidemiology Unit | Erika Scaltriti; Giulia Fregni Serpini; Ilaria Menozzi; Marina Morganti; Monica Pecorari; Stefano Pongolini; William Gennari |
| EPI_ISL_1222789, EPI_ISL_2372852 | Azienda Sanitaria dell'Alto Adige Laboratorio Aziendale di Microbiologia e Virologia | Istituto di Genomica Applicata | Davide Scaglione; Eleonora Paparelli; Elisa Masi; Elisabetta Giacobazzi; Elisabetta Pagani; Gabriele Magris; Irena Jurman; Irene Bianconi; Michele Morgante; Stefanie Wieser; Vera Vendramin |
| EPI_ISL_1969249 | BBLK Palembang | National Institute of Health Research and Development | Arie Ardiansyah Nugraha; Hana Apsari Pawestri; Hartanti Dian Ikawati; Kartika Dewi Puspa; Krisna Pangesti; Nelly Puspandari; Subangkit; Triyani Soekarso; Vivi Setiawaty |
| EPI_ISL_7678300 | BIO LBS St Romain de Colbosc | CERBA HealthCare | Bénédicte Roquebert; Laura Verdurme; Sabine Trombert; Stéphanie Haim-Boukobza |
| EPI_ISL_7678522 | BIOALLIANCE-SITE CHARTRES | CERBA HealthCare | Bénédicte Roquebert; Laura Verdurme; Sabine Trombert; Stéphanie Haim-Boukobza |
| EPI_ISL_7898563 | BIOLAB UNILABS SITE BOUCICAUT | CERBA HealthCare | Bénédicte Roquebert; Laura Verdurme; Sabine Trombert; Stéphanie Haim-Boukobza |
| EPI_ISL_2373691 | BIOMNIS LYON | CNR Virus des Infections Respiratoires - France SUD | Antonin Bal; Bruno Lina; Bruno Simon; Gregory Destras; Gwendolyne Burfin; Hadrien Regue; Laurence Josset; Martine Valette; Quentin Semanas |
| EPI_ISL_7976432 | BIOPATH LABM | Department of Virology, Henri Mondor University Hospital, Assistance Publique Hôpitaux de Paris, Université Paris-Est Créteil, INSERM U955 | Alexandre Soulier; Christophe Rodriguez; Elisabeth Trawinski; Guillaume Gricourt; Jean-Michel Pawlotsky; Melissa N'Debi; Slim Fourati; Vanessa Demontant |
| EPI_ISL_2321586 | BIOREFERENCIA | Universidad Nacional de Colombia - Laboratorio Genómico One Health | Andres F. Cardona-Rios; Carlos Franco-Muñoz; Carolina Muñoz-Arango; Celeny Ortiz; Daniel O. Maldonado-Perez; Diego A. Álvarez-Díaz; Hector Alejandro Ruiz-Moreno; Idabely Betancur Ortiz; Jorge E. Osorio; Juan P. Hernandez-Ortiz; Karl A Ciuoderis; Katherine Laiton-Donato; Laura Silvana Perez; Lina M. Hurtado; Marcela Mercado-Reyes; Maria Angélica Maya; Maria Stella López; Rita Almanza Payares; Sandra Ines Cano; Simón Villegas Velásquez |
| EPI_ISL_2342878 | BRLSABVM Government Medical College, Rajnandgaon, Chhattisgarh | Institute of Life Sciences - INSACOG | Ajay Parida; Amol M. Kanampalliwar; Arup Ghosh; Atimukta Jha; INSACOG Consortium; Omprakash Shiriwas; Punit Prasad; Rajeeb Swain; Rupesh Dash; Safal Walia; Sana Fatma; Shifu Aggarwal; Sunil K. Raghav |
| EPI_ISL_5142661, EPI_ISL_5142662 | BTC, Khalifa University | BTC, Khalifa University | Habiba Al Safar et al |
| EPI_ISL_2373506, EPI_ISL_2373545 | Banaras Hindu University | CSIR-Centre for Cellular and Molecular Biology-INSACOG | ; Abhay Kumar Yadav; Ajay Kumar Yadav; Amareshwar Vodapalli; Ara Sreenivas; Archana Bharadwaj Siva; Ashish; B Himasri; Blessy B John; Chetan Sahni; Deepa Devadas; Divya Tej Sowpati; Gunjan Rai; Gyaneshwer Chaubey; Jaya Chakraborty; Karthik Bharadwaj Tallapaka; Lamuk Zaveri; Maneesha Upadhyay; Manpreet Kaur; Nitish Kumar Singh; Payel Mukherjee; Priyoneel Basu; Rakesh K Mishra; Royana Singh; Saumya Singh; Saurabh Singh; Shani Vishwakarma; Sharath Chandra Thota; Shivam Tiwari; Shivani Mishra; Shreekant Verma; Sofia Banu; Surendra Pratap Mishra; Tulasi Nagabandi; Umesh Choudhary; Valli Nagalakshmi Undamatla; Viswagithe S L |
| EPI_ISL_2315704 | Bayerisches Landesamt für Gesundheit und Lebensmittelsicherheit (LGL) | Robert Koch Institute | |
| EPI_ISL_8253538 | Berkeley Medical Center | WVU and Marshall University Combined Genomics Core Facilities | James Denvir; Peter Perrotta; Peter Stoilov; Ryan Percifield; Wesley Kimble |
| EPI_ISL_1406151 | Biolab Diagnostic Laboratories | Biolab Diagnostic Laboratories | Ahmad Tibi; Amid Abdelnour; Badia Saddedin; Eiad Atwa; Issa Abu-Dayyeh; Lama Hussein; Shayma Ali |
| EPI_ISL_1697435 | Bioptická laboratoř, s.r.o. | Bioptická laboratoř, s.r.o. | Martina Putzová; Michaela Říhová; Nikola Bílá; Petr Šteiner; Silva Vondráková; Tomáš Vaněček |
| EPI_ISL_2108737, EPI_ISL_2313387, EPI_ISL_2388599, EPI_ISL_8221436 | Bioscientia Labor Wermsdorf | Robert Koch Institute | |
| EPI_ISL_2313579 | Bioscientia MVZ Labor Karlsruhe GmbH | Robert Koch Institute | |
| EPI_ISL_1970089, EPI_ISL_1970223, EPI_ISL_1970224 | Biotechnology Division, NCDC Delhi | NCDC Delhi, Biotechnology Division | Hema Gogia; Hemlata Lall; Kalaiarasan Ponnusamy; Mahesh S Dhar; Manoj K Singh; Meena Datta; Partha Rakshit; Preeti Madan; Priyanka Singh; Radhakrishnan V. S; Robin Marwal; Sandhya Kabra; Sujeet K Singh; Uma Sharma |
| EPI_ISL_745158 | Bishop Lavis CDC wc BLP | National Health Laboratory Service (NHLS), Tygerberg | Bronwyn Kleinhans; Eduan Wilkindon; Gert van Zyl; Houriiyah Tegally; Kayla Delaney; Susan Engelbrecht; Tulio de Oliveira; Wolfgang Preiser |
| EPI_ISL_2504070, EPI_ISL_6640916, EPI_ISL_6640917, EPI_ISL_6640919, EPI_ISL_6670244, EPI_ISL_6752026, EPI_ISL_6774083, EPI_ISL_6774089, EPI_ISL_6774093 | | | |
| see above | Botswana Harvard HIV Reference Laboratory | Botswana Harvard HIV Reference Laboratory | Boitumelo Zuze; Botshelo Radibe; Dorcas Maruapula; Joseph Makhema; Keoratile Ntshambiwa; Kgomotso Moruisi; Legodile Kooepile; Madisa Mine; Modisa Motswaledi; Mosepele Mosepele; Mphaphi B. Mbulawa; Ontlametse T. Bareng; Pamela Smith-Lawrence; Roger Shapiro; Sefetogi Ramaologa; Shahin Lockman; Sikhulile Moyo; Simani Gaseitsiwe; Thongbotho Mphoyakgosi; Wonderful T. Choga |
| EPI_ISL_4839236 | British Columbia Centre For Disease Control | BCCDC Public Health Laboratory | Ana Pacagnella; Corrinne Ng; Dan Fornika; John Tyson; Kim Macdonald; Kimia Kamelian; Linda Hoang; Loretta Janz; Mel Krajden; Prystajecky Natalie; Robert Azana; Shannon Russell |
| EPI_ISL_1710085, EPI_ISL_2312405, EPI_ISL_2312423, EPI_ISL_4285069, EPI_ISL_4286289, EPI_ISL_4516698, EPI_ISL_4517827, EPI_ISL_4638683, EPI_ISL_4729194, EPI_ISL_4970955, EPI_ISL_4971531, EPI_ISL_5386578, EPI_ISL_5746284, EPI_ISL_5794752, EPI_ISL_5795095, EPI_ISL_6035126, EPI_ISL_6448163, EPI_ISL_6754719, EPI_ISL_6754961, EPI_ISL_6828852, EPI_ISL_7340999, EPI_ISL_7387217, EPI_ISL_7544594, EPI_ISL_7658932, EPI_ISL_7824203, EPI_ISL_8214815, EPI_ISL_8214883 | | | |
| see above | Broad Institute Clinical Research Sequencing Platform | Infectious Disease Program, Broad Institute of Harvard and MIT | Adams, G.; B.L.; B.W.; Bauer, M.; Birren; Blumenstiel, B.; Brown, C.; Carter, A.; Chaluvadi, S.; D.J.; DeFelice, M.; DeRuff, K.; Dodge, S.; Gabriel, S.; Gallagher, G.; Gladden-Young, A.; Granger, B.; J.E.; K.J.; Lagerborg, K.; Larkin, K.; Lee, M.; Lemieux; Lennon, N.; Loreth, C.; Madoff, L.; McGovern, S.; Meldrim, J.; Normandin, E.; P.C.; Park; Pearlman, L.; Reilly, S.; Rudy, M.; Sabeti; Siddle; Smole, S.; Tomkins-Tinch, C.; Vicente, G.; and MacInnis |
| EPI_ISL_7263830 | CAP Roger de Flor | Banc de Sang i Teixits | Carlos Hobeich; Francisco Vidal; Irene Corrales; Lorena Ramírez; Maria Glòria Soria; Natália Comes; Nina Borràs; Noemí Gonzalez; Silvia Sauleda |
| EPI_ISL_2362583 | CDI | Instituto Nacional de Salud- Dirección de Investigación en Salud Pública | Carlos Franco-Muñoz; Carmen Osorio; Diana Malo; Diego A. Álvarez-Díaz; Diego Andrés Prada; Gerardo Santamaría; Hector Alejandro Ruiz-Moreno; Jhonnatan Reales-González; Jorge Rivera; Juan Camilo Martinez; Julian Naizaque; Katherine Laiton-Donato; Lisseth Pardo; Magdalena Wiesner; Marcela Mercado-Reyes; Maria T. Herrera-Sepúlveda; Marta Lopez Blanco; Martha Lucia Ospina Martinez; Paola Rojas; Sergio Gomez; Sheryll Corchuelo; Ángela Alarcon Cruz |
| EPI_ISL_1060898, EPI_ISL_1064745 | CDL Laboratorio Santos e Vidal LTDA. | Instituto de Medicina Tropical de Sao Paulo | Brazil-UK Centre for Arbovirus Discovery Diagnosis Genomics and Epidemiology (CADDE) Genomic Network - Instituto de Medicina Tropical |
| EPI_ISL_5445440, EPI_ISL_5445475, EPI_ISL_5900176, EPI_ISL_5900247, EPI_ISL_6265191, EPI_ISL_6265443, EPI_ISL_6862324, EPI_ISL_7999040 | | | |
| see above | CDPH VBL | California Department of Public Health | CDPH COVIDNet; Emily Smith on behalf of CDPH-COVIDNet; Emily Smith on behalf of CDPH-COVIDNet and UCLA Technology Center for Genomics & Bioinformatics |
| EPI_ISL_2268036, EPI_ISL_4933474 | CENTOGENE Frankfurt Laboratory: Niederlassung Industriepark Höchst | Robert Koch Institute | |
| EPI_ISL_7890669 | CENTRE HOSPITALIER DE BOURG EN BRESSE | CNR Virus des Infections Respiratoires - France SUD | Antonin Bal; Bruno Lina; Bruno Simon; Gregory Destras; Gwendolyne Burfin; Hadrien Regue; Laurence Josset; Martine Valette; Quentin Semanas |
| EPI_ISL_7491050 | CERBALLIANCE IDF OUEST SITE JOUY EN JOSA | CERBA HealthCare | Bénédicte Roquebert; Laura Verdurme; Sabine Trombert; Stéphanie Haim-Boukobza |
| EPI_ISL_6370560 | CERBALLIANCE IDF OUEST SITE LES CLAYES | CERBA HealthCare | Bénédicte Roquebert; Laura Verdurme; Sabine Trombert; Stéphanie Haim-Boukobza |
| EPI_ISL_7044771 | CERBALLIANCE IDF SUD SITE DE LISSES | CERBA HealthCare | Bénédicte Roquebert; Laura Verdurme; Sabine Trombert; Stéphanie Haim-Boukobza |
| EPI_ISL_7206964 | CH GIVORS | CNR Virus des Infections Respiratoires - France SUD | Antonin Bal; Bruno Lina; Bruno Simon; Gregory Destras; Gwendolyne Burfin; Hadrien Regue; Laurence Josset; Martine Valette; Quentin Semanas |
| EPI_ISL_8190556 | CHC Dakor | Gujarat Biotechnology Research Centre | Akhilesh Modi; Apurvasinh Puvar; Bhadreshsinh Gohil; Chaitanya Joshi; Disha Vora; Janvi Raval; Jaykumar Rangani; Madhvi Joshi; Navendu Joshi; Nimesh Patel; Nitin Savaliya; Nitin Shukla; Priyank Chavda; Ramesh Pandit; Roshani Mishra; Sonal Sharma; Tasnim Trivedi; Zarna Patel |
| EPI_ISL_2090827 | CHOR | UMR PIMIT | Dr Camille Lebarbenchon; Dr David A Wilkinson; Dr Patrick Mavingui; Magali Turpin |
| EPI_ISL_860907 | CHU Tours - Virologie | National Reference Center for Viruses of Respiratory Infections, Institut Pasteur, Paris | Angela Brisebarre; Camille Capel; Etienne Simon-Lorière; Gaudy Graffin Catherine; Marion Barbet; Maud Vanpeene; Méline Bizard; Sylvie Behillil; Sylvie van der Werf; Vincent Enouf |
| EPI_ISL_4632159 | CHUV | Laboratory of genomics and metagenomics | Claire Bertelli; Damien Jacot; Gilbert Greub; Sébastien Aeby; Trestan Pillonel |
| EPI_ISL_8189703 | CLINA-LANCET LABORATORIES | National Reference Laboratory, Nigeria Centre for Disease Control | Anthony Ahumibe; Catherine Okoi; Chimaobi Chukwu; Dr Ifedayo Adetifa; Dr Ndodo Nnaemeka; Dr Omoare Adesuyi; Nwando Mba; Olusola Anuoluwapo Akanbi; Oyeronke Ayansola |
| EPI_ISL_1313954, EPI_ISL_6846282, EPI_ISL_7890962 | CNR Virus des Infections Respiratoires - France SUD | CNR Virus des Infections Respiratoires - France SUD | Antonin Bal; Bruno Lina; Bruno Simon; Gregory Destras; Gwendolyne Burfin; Hadrien Regue; Laurence Josset; Martine Valette; Quentin Semanas |
| EPI_ISL_1838446, EPI_ISL_1838463, EPI_ISL_1838638, EPI_ISL_1838646, EPI_ISL_1838712, EPI_ISL_2156950, EPI_ISL_2373429, EPI_ISL_2441487, EPI_ISL_2441639 | | | |
| see above | CSIR-Centre for Cellular and Molecular Biology | CSIR-Centre for Cellular and Molecular Biology-INSACOG | Amareshwar Vodapalli; Ara Sreenivas; Archana Bharadwaj Siva; B Himasri; Blessy B John; Divya Tej Sowpati; Karthik Bharadwaj Tallapaka; Lamuk Zaveri; Onkar Kulkarni; Payel Mukherjee; Priya Nurkurthy; Rakesh K Mishra; Sharath Chandra Thota; Shreekant Verma; Sofia Banu; Sumedha Avadhanula; Tulasi Nagabandi; Valli Nagalakshmi Undamatla; Vidhyadhari Methuku; Viswagithe S L |
| EPI_ISL_1916465, EPI_ISL_1916468, EPI_ISL_1916472, EPI_ISL_1916475, EPI_ISL_1916486 | CSIR-National Environmental Engineering Research Institute | CSIR-Centre for Cellular and Molecular Biology - INSACOG | Amareshwar Vodapalli; Ara Sreenivas; B Himasri; Divya Tej Sowpati; Karthik Bharadwaj Tallapaka; Krishna Khairnar; Lamuk Zaveri; Onkar Kulkarni; Rakesh K Mishra; Sharath Chandra Thota; Shreekant Verma; Sofia Banu; Viswagithe S L |
| EPI_ISL_2441372, EPI_ISL_2441385 | CSIR-National Environmental Engineering Research Institute | CSIR-Centre for Cellular and Molecular Biology-INSACOG | Amareshwar Vodapalli; Ara Sreenivas; Archana Bharadwaj Siva; B Himasri; Divya Tej Sowpati; Karthik Bharadwaj Tallapaka; Krishna Khairnar; Lamuk Zaveri; Onkar Kulkarni; Payel Mukherjee; Priya Nurkurthy; Rakesh K Mishra; Shreekant Verma; Sofia Banu; Sumedha Avadhanula; Tulasi Nagabandi; Valli Nagalakshmi Undamatla; Vidhyadhari Methuku |
| EPI_ISL_2234344 | California Department of Public Health | California Department of Public Health | CDPH IDLB COVIDNet et al |
| EPI_ISL_5115025, EPI_ISL_5741799 | Cantacuzino National Military-Medical Institute, Viral Respiratory Infections Laboratory | Cantacuzino National Military-Medical Institute, Viral Respiratory Infections Laboratory | Carmen Cherciu; Luiza Ustea; Mihaela Lazar; Mihaela Oprea; Nicoleta Paraschiv; Sorin Dinu |
| EPI_ISL_2290652 | Cell-T | 1. National Institute of Public Health - National Institute of | Dominik Strapagiel; Izabela Dróżdż; Jakub Lach; Katarzyna Zacharczuk; Klaudyna Królikowska; Maciej Borowiec; Magdalena Nowakowska; Magdalena Traczyk-Borszyńska; Marcin Słomka; Marta Sobalska-Kwapis; |

| | | | |
|---|---|---|---|
| | | Hygiene, Warsaw, Poland 2. Biobank Lab, University of Lodz 3. Laboratory of Respiratory Viruses, Teaching and Clinical Center of the Medical University of Lodz | Małgorzata Sadkowska-Todys; Tomasz Płoszaj; Tomasz Wołkowicz |
| EPI_ISL_2450042 | Center for Laboratory Medicine | Center for Laboratory Medicine | Yannick Gerth |
| EPI_ISL_5064723 | Central Diagnostic Laboratory | National Public Health Laboratory | National Public Health Laboratory Team |
| EPI_ISL_2464017 | Central Public Health Lab, National Public Health Organization | Central Public Health Lab, National Public Health Organization | Gregory Spanakos et al; Kyriaki Tryfinopoulou |
| EPI_ISL_1585389 | Centro de Investigación Biomédica del Noreste (CIBIN) | Instituto Nacional de Enfermedades Respiratorias (INER): Centro de Investigación en Enfermedades Infecciosas (CIENI) | Alejandro Sanchez-Flores; Alfredo Herrera-Estrella; Alicia Ocaña-Mondragón; Angel Gustavo Salas-Lais; Bernardo Martínez-Miguel; Blanca Taboada; Brenda Irasema Maldonado-Meza; Carla Ivón Herrera-Najera; Carlos F. Arias; Celia Boukadida; Clara Esperanza Santacruz-Tinoco; Concepción Grajales-Muñiz; Consorcio Mexicano de Vigilancia Genómica (CoViGen-Mex). Authors (in alphabetical order): Julio Elias Alvarado-Yaah; Célida Duque Molina; Fernando Fontove-Herrera; Francisco Pulido; Gloria Elena Espinosa-Ayala; Gloria María Molina-Salinas; Gloria Vazquez; Hector Esteban Paz-Juárez; Hector Montoya-Fuentes; Helen Haydee Fernanda Ramírez-Plascencia; José Antonio Enciso-Moreno; José Esteban Muñoz-Medina; José de Jesús Nuñez-Contreras; Juan Bautista Chale-Dzul; Luis Alberto Ochoa-Carrera; Margarita Matías-Florentino; María Guadalupe Santiago-Mauricio; María Guadalupe de Jesús Mireles-Rivera; Nelly Sélem-Mojica; Pavel Isa; Ricardo Grande; Santiago Ávila-Ríos; Victor Eduardo Garcia-Arias; Víctor Hugo Borja-Aburto |
| EPI_ISL_6700542 | Centrum Medycyny Klinicznej dr. n.med. Karol Majewski | Wojewodzka Stacja Sanitarno-Epidemiologiczna w Lodzi, Oddzial Laboratoryjny Mikrobiologii i Parazytologii | Agnieszka Karnowska; Aleksandra Krawczyk; Danuta Bartczak; Elian Terpo; Katarzyna Kosowska; Lucyna Kochanowska; Marta Gawronska; Martyna Matera; Szymon Konkolewski |
| EPI_ISL_7337469, EPI_ISL_7337471, EPI_ISL_7337486, EPI_ISL_7337501, EPI_ISL_7337503, EPI_ISL_7337511 | Charlotte Maxeke Johannesburg Academic Hospital | National Institute for Communicable Diseases of the National Health Laboratory Service | Amoako DG; Bhiman JN; Everatt J; Ismail A; Mahlangu B; Mnguni A; Mohale T; Ntuli N; Scheepers C |
| EPI_ISL_940104, EPI_ISL_940120, EPI_ISL_940124, EPI_ISL_1239299 | Charlotte Maxeke Johannesburg Academic Hospital, National Health Laboratory Services, Gauteng, South Africa | National Institute for Communicable Diseases of the National Health Laboratory Service | Allam M; Amoako DG; Bhiman JN; Ismail A; Mahlangu B; Mohale T; Ntuli N; Scheepers C |
| EPI_ISL_1239334, EPI_ISL_1239341 | Chris Hani Baragwanath Hospital, National Health Laboratory Services, Gauteng, South Africa | National Institute for Communicable Diseases of the National Health Laboratory Service | Amoako DG; Bhiman JN; Ismail A; Mahlangu B; Mohale T; Ntuli N; Scheepers C |
| EPI_ISL_7337525, EPI_ISL_7337541, EPI_ISL_7337553, EPI_ISL_7337560 | Chris Hani Baragwanath Laboratory | National Institute for Communicable Diseases of the National Health Laboratory Service | Amoako DG; Bhiman JN; Everatt J; Ismail A; Mahlangu B; Mnguni A; Mohale T; Ntuli N; Scheepers C |
| EPI_ISL_2523856 | City Hospital No 40 | WHO National Influenza Centre Russian Federation | Andrey Komissarov; Artem Fadeev; Daria Danilenko; Dmitry Lioznov; Elena Nabieva; Georgii Bazykin; Kirill Varchenko; Ksenia Safina; Kseniya Komissarova; Maria Pisareva; Maria Timofeeva; Mikhail Bakaev; Nikita Yolshin; Olga Shneider; Oula Mansour; Sergey Scherbak; Tamila Musaeva; Veronika Eder |
| EPI_ISL_1821066, EPI_ISL_1821067 | Clinica los Nogales | Instituto Nacional de Salud- Dirección de Investigación en Salud Pública | Carlos Franco-Muñoz; Carmen Osorio; Diana Malo; Diego A. Álvarez-Díaz; Diego Andrés Prada; Gerardo Santamaría; Hector Alejandro Ruiz-Moreno; Jhonnatan Reales-González; Jorge Rivera; Juan Camilo Martinez; Julian Naizaque; Katherine Laiton-Donato; Lisseth Pardo; Magdalena Wiesner; Marcela Mercado-Reyes; Maria T. Herrera-Sepúlveda; Marta Lopez Blanco; Martha Lucia Ospina Martinez; Paola Rojas; Sergio Gomez; Sheryll Corchuelo; Ángela Alarcon Cruz |
| EPI_ISL_7266083 | Clinical Microbiology Laboratory, Tel Aviv Sourasky Medical Center | Clinical Microbiology Laboratory, Tel Aviv Sourasky Medical Center | Alon Ziv; Amos Adler; Katya Levytskyi; Lior Handler; Ora Halutz |
| EPI_ISL_1373506, EPI_ISL_2380818, EPI_ISL_7782150 | Clinical Molecular Microbiology Laboratory, UNC Hospitals | Jeremy Wang | Alexander Rubinsteyn; Bravada Hill; Colleen Rice; Corbin Jones; Jason Smedberg; Jeremy Wang; Katie Martin; Melissa Miller; Robert Hagan; Sayal Guirales-Medrano; Shawn Hawken; Tara Skelly |
| EPI_ISL_7977173 | Clinical Virology | Clinical Bacteriology, University Hospital Basel | Adrian Egli; Alfredo Mari; Fanny Wegner; Hans Hirsch; Helena MB Seth-Smith; Julia Bielicki; Karoline Leuzinger; Manuel Battegay; Tim Roloff |
| EPI_ISL_745153 | CoVid EC Nelson Mandela Bay Metro | National Health Laboratory Service (NHLS), Tygerberg | Bronwyn Kleinhans; Eduan Wilkindon; Gert van Zyl; Houriiyah Tegally; Kayla Delaney; Susan Engelbrecht; Tulio de Oliveira; Wolfgang Preiser |
| EPI_ISL_1300064, EPI_ISL_2309379, EPI_ISL_2309975, EPI_ISL_2310477, EPI_ISL_4967132, EPI_ISL_5415394, EPI_ISL_5708873, EPI_ISL_6676339, EPI_ISL_6815245, EPI_ISL_6911309, EPI_ISL_6913099, EPI_ISL_7150010, EPI_ISL_7331814, EPI_ISL_7333523, EPI_ISL_7458772, EPI_ISL_7459080, EPI_ISL_7501690, EPI_ISL_7805070, EPI_ISL_7812204, EPI_ISL_7830313, EPI_ISL_7888000, EPI_ISL_8064172, EPI_ISL_8065109, EPI_ISL_8158786, EPI_ISL_8160529, EPI_ISL_8162212, EPI_ISL_8162447 | | | |
| see above | Colorado Department of Public Health and Environment | Colorado Department of Public Health and Environment | Alexandria Rossheim; Arianna Smith; Diana Ir; Emily A. Travanty; Laura Bankers; Mandy Waters; Michael A. Martin; Michael Martin; Molly C. Hetherington-Rauth; Sarah Elizabeth Totten; Shannon Ely; Shannon R. Matzinger |
| EPI_ISL_1398198, EPI_ISL_1789713 | Columbia University Irving Medical Center | Wadsworth Center, New York State Department of Health | Alexis Russel; Alexis Russell; Catharine Prussing; Daryl M. Lamson; Erasmus Schneider; Erica Lasek-Nesselquist; John Kelly; Jonathan Plitnick; Kirsten St. George; Matthew Shudt; Melissa A Leisner; Navjot Singh |
| EPI_ISL_1660442 | Communicable Disease Laboratory, Public Health Directorate | Communicable Disease Laboratory, Public Health Directorate | AlAbbas, Z.; AlHujairi, Z.; Almoamen, G.; Alwasti, H.; Marhoon, A. |
| EPI_ISL_700441, EPI_ISL_700492, EPI_ISL_2621065 | Conville CDC wc CVC | NHLS/UCT | Arash Iranzadeh; Bruna Galvao; Carolyn Williamson; Deelan Doolabh; Diana Hardie; Gert Marais; Houriiyah Tegally; Innocent Mudau; Kruger Marais; Lynn Tyers; Marvin Hsiao; Stephen Korsman |
| EPI_ISL_960122, EPI_ISL_960134 | Conville CDC wc CVC | National Health Laboratory Service/UCT | Arash Iranzadeh; Bruna Galvao; Carolyn Williamson; Deelan Doolabh; Diana Hardie; Innocent Mudau; Kruger Marais; Lynn Tyers; Marvin Hsiao; Stephen Korsman |
| EPI_ISL_1821399 | County of San Luis Obispo Public Health Laboratory | Chan-Zuckerberg Biohub | CZB Cliahub Consortium |
| EPI_ISL_4910276 | County of San Luis Obispo Public Health Laboratory | County of San Luis Obispo Public Health Laboratory | Frances Sidhu; Katrina Erwin |
| EPI_ISL_7451089 | Cruz Vermelha Portuguesa | Instituto Nacional de Saude (INSA) | Borges et al |
| EPI_ISL_5523909 | Curative Labs | Curative Labs | Elias L. Salfati; Eugenia Khorosheva; George Way; J.Cesar Ignacio-Espinoza; Janet Chen; Mikhail Hanewich-Hollatz; Nabjot Sandhu; Sophia Quasem; Vladimir Slepnev; Zhiyi Xie |
| EPI_ISL_700470 | D'Almeida Clinic wc DAL | NHLS/UCT | Arash Iranzadeh; Bruna Galvao; Carolyn Williamson; Deelan Doolabh; Diana Hardie; Houriiyah Tegally; Innocent Mudau; Kruger Marais; Lynn Tyers; Marvin Hsiao; Stephen Korsman |
| EPI_ISL_833167, EPI_ISL_833171, EPI_ISL_833172, EPI_ISL_833173, EPI_ISL_833174 | DB Diagnosticos do Brasil | Instituto Adolfo Lutz, Interdiciplinary Procedures Center, Strategic Laboratory | Claudia Regina Gonçalves; Claudio Tavares Sacchi; Erica Valessa Ramos Gomes; Karoline Rodrigues Campos |
| EPI_ISL_1060914, EPI_ISL_1060918 | DB Diagnosticos do Brasil | Instituto de Medicina Tropical de Sao Paulo | Brazil-UK Centre for Arbovirus Discovery Diagnosis Genomics and Epidemiology (CADDE) Genomic Network - Instituto de Medicina Tropical |
| EPI_ISL_804823, EPI_ISL_804824 | DB Diagnosticos do Brasil | Laboratório de Parasitologia Médica - Instituto de Medicina Tropical - Universidade de São Paulo | Andrew Rambaut; CADDE Genomic Network.; CDL; Camila A. Maia da Silva; Cecília da Cunha Camilo; DB; Darlan Candido; Erika Regina Manuli; Ester C. Sabino; Flavia Cristina Sales; HEMOAM; Ingra Morales Claro; Lucas A. Moyses Franco; Maria do Perpétuo Socorro Sampaio Carvalho; Myuki Alfaia Esashika Crispim; Nelson Abrahim Fraiji; Nelson Gaburo; Nick Loman; Nuno Faria; Oliver G. Pybus; Pamela dos Santos Andrade; Renato A. Santana; Thais de Moura Coletti |
| EPI_ISL_2158345 | DIRECCION DE SANIDAD EJERCITO NACIONAL | Instituto Nacional de Salud- Dirección de Investigación en Salud Pública | Carlos Franco-Muñoz; Carmen Osorio; Diana Malo; Diego A. Álvarez-Díaz; Diego Andrés Prada; Gerardo Santamaría; Hector Alejandro Ruiz-Moreno; Jhonnatan Reales-González; Jorge Rivera; Juan Camilo Martinez; Julian Naizaque; Katherine Laiton-Donato; Lisseth Pardo; Magdalena Wiesner; Marcela Mercado-Reyes; Maria T. Herrera-Sepúlveda; Marta Lopez Blanco; Martha Lucia Ospina Martinez; Paola Rojas; Sergio Gomez; Sheryll Corchuelo; Ángela Alarcon Cruz |
| EPI_ISL_966386 | DOHMH Corona | New York City Public Health Laboratory | Jade Wang; et al. |
| EPI_ISL_6962496 | DR P BHASIN PATH LABS | ILBS | Arjun Bhugra; Chhagan Bihari Sharma; Ekta Gupta; Pramod Gautam; Rahul Garg; Reshu Agarwal; Shiv Kumar Sarin; Varun Suroliya; santanu das; urvinder kaur |
| EPI_ISL_745171 | Delft CHC wc DFP | National Health Laboratory Service (NHLS), Tygerberg | Bronwyn Kleinhans; Eduan Wilkindon; Gert van Zyl; Houriiyah Tegally; Kayla Delaney; Susan Engelbrecht; Tulio de Oliveira; Wolfgang Preiser |
| EPI_ISL_2100636, EPI_ISL_2106198 | Delhi North District | CSIR-Institute of Genomics and Integrative Biology | Animesh Ray; Asangala Kamai; Ashwin Varadarajan; Ayush Goel; Bharathram Uppili*; Devasish Desai; Manish Kumar; Mohammed Ahmed; Mohammed Faruq; Naveet Wig; Pooja Sharma*; Rajesh Pandey; Sarafaraz Alam; Saruchi Wadhwa; Satish Swain; Sheeba Saifi; Sushma Rajpoot; Umang Arora |
| EPI_ISL_4634765, EPI_ISL_7467005 | Department Medical Microbiology, Baerum Hospital, Vestre Viken Health Trust | Norwegian Institute of Public Health, Department of Virology | Atiya R Ali; Debech Nadia; Engebretsen Serina Beate; Garcia Llorente Ignacio; Hilde Elshaug; Hilde Vollan; Jon Bråte; Kamilla Heddeland Instefjord; Karoline Bragstad; Kathrine Stene-Johansen; Line Victoria Moen; Marie Paulsen Madsen; Olav Hungnes; Pedersen Benedikte Nevjen; Rasmus Riis Kopperud |
| EPI_ISL_8267929 | Department for Virology, Molecular Biology and Genome Research, R. G. Lugar Center for Public Health Research, National Center for Disease Control and Public Health (NCDC) of Georgia. | Department for Virology, Molecular Biology and Genome Research, R. G. Lugar Center for Public Health Research, National Center for Disease Control and Public Health (NCDC) of Georgia. | Adam Kotorashvili; Amiran Gamkrelidze.; Ana Papkiauri; Ann Machablishvili; Anna Kasradze; Davit Tsaguria; Ekaterine Khmaladze; Ekaterine Zangaladze; Ekaterine Zhghenti; Giorgi Gogoladze; Giorgi Tomashvili; Gvantsa Brachveli; Gvantsa Chanturia; Irma Burjanadze; Ketevan Sidamonidze; Khatuna Zakhashvili; Lela Sabadze; Lela Urushadze; Magda Dgebuadze; Maia Alkhazashvili; Mari Gavashelidze; Mariam Zakalashvili; Marine Murtskhvaladze; Meri Pantsulaia; Nato Kotaria; Nino Berishvili; Paata Imnadze; Roena Sukhiashvili; Salome Javashvili; Tamar Jashiashvili; Tata Imnadze; Tea Tevdoradze |
| EPI_ISL_2413521, EPI_ISL_2414083, EPI_ISL_2414530, EPI_ISL_4355884, EPI_ISL_4355927, EPI_ISL_4626180, EPI_ISL_4886786, EPI_ISL_4977880, EPI_ISL_5057802, EPI_ISL_5098516, EPI_ISL_5882663, EPI_ISL_5916862, EPI_ISL_6099754, EPI_ISL_6241663, EPI_ISL_6244823, EPI_ISL_6248300, EPI_ISL_6305788, EPI_ISL_6321391, EPI_ISL_6505724, EPI_ISL_6506714, EPI_ISL_6722844, EPI_ISL_6723810, EPI_ISL_6730278, EPI_ISL_6829729, EPI_ISL_6848310, EPI_ISL_6848904, EPI_ISL_6965656, EPI_ISL_7278972, EPI_ISL_7281271, EPI_ISL_7283401, EPI_ISL_7286547, EPI_ISL_7647702, EPI_ISL_7904633, EPI_ISL_7905897, EPI_ISL_7909233, EPI_ISL_7911063, EPI_ISL_7916486, EPI_ISL_8002846, EPI_ISL_8053434, EPI_ISL_8171038, EPI_ISL_8201911, EPI_ISL_8249443, EPI_ISL_8249749, EPI_ISL_8250459 | | | |
| see above | Department of Bacteria, Parasites and Fungi, Statens Serum Institut, Copenhagen, Denmark | Statens Serum Institut Bioinformatics and Microbial Genomics | Danish Covid-19 Genome Consortium |
| EPI_ISL_1826809 | Department of Clinical Microbiology | GIGA Medical Genomics | Bouchra Boujemla; Cécile Meex; Keith Durkin; Maria Artesi; Marie-Pierre Hayette; Nathalie Renotte; Pierrette Melin; Raphaël Boreux; Sébastien Bontems; Vincent Bours |
| EPI_ISL_8053996 | Department of Clinical Microbiology and Center for Genomic Medicine, Rigshospitalet, Copenhagen, Denmark | Statens Serum Institut Bioinformatics and Microbial Genomics | Danish Covid-19 Genome Consortium |
| EPI_ISL_7192723 | Department of Health Technology and Informatics, The Hong Kong Polytechnic University | Department of Health Technology and Informatics, The Hong Kong Polytechnic University | Alan Ka-Lun Wu; Alex Yat-Man Ho; Barry Kin-Chung Wong; Chloe Toi-Mei Chan; David Ho-Keung Shum; Denise Sze-Hang Wong; Gilman Kit-Hang Siu; Hiu-Yin Lao; Hoi-Ching Jim; Ivan Tak-Fai Wong; Jake Siu-Lun Leung; Kam-Tong Yip; Kenneth Siu-Sing Leung; Kingsley King-Gee Tam; Kitty Sau-Chun Fung; Kristine Luk; Lam-Kwong Lee; Miranda Chong-Yee Yau; Sandy Ka-Yee Chau; Shea Ping Yip; Tak-Lun Que; Timothy Ting-Leung Ng; Wing Cheong Yam; Wing-Hei Lo; Wing-Kin To; Yvette Wai-Man Lai |
| EPI_ISL_1200004 | Department of Infectious Diseases, Istituto Superiore di Sanità, Rome, Italy; Università degli Studi di Perugia, Perugia, Italy | Istituto Superiore di Sanità (ISS) | Alessandra Lo Presti; Angela Di Martino; Antonella Mencacci; Barbara Camilloni; Manuela Marra; Marco Crescenzi; Maria Carollo; Paola Stefanelli; Stefano Fiore |

| | | | |
|---|---|---|---|
| EPI_ISL_2358356, EPI_ISL_5767724 | Department of Infectious Diseases, Kobe Institute of Health | Department of Infectious Diseases, Kobe Institute of Health | Kentaro Itokawa; Makoto Kuroda; Masanori Hashino; Noriko Nakanishi; Rina Tanaka; Ryohei Nomoto; Tomotada Iwamoto; Tsuyoshi Sekizuka |
| EPI_ISL_1678824 | Department of Medical Microbiology & Infection prevention, Amsterdam University Medical Centers location AMC | Department of Medical Microbiology & Infection prevention, Amsterdam University Medical Centers location AMC | Fokla Zorgdrager; Janke Schinkel; Marcel Jonges; Matthijs Welkers; Menno de Jong; Robin van Houdt; Sebastien Matamoros; Sjoerd Rebers |
| EPI_ISL_6960417 | Department of Medical Microbiology - section Molde, Molde Hospital | Norwegian Institute of Public Health, Department of Virology | Atiya R Ali; Debech Nadia; Engebretsen Serina Beate; Garcia Llorente Ignacio; Hilde Elshaug; Hilde Vollan; Jon Bråte; Kamilla Heddeland Instefjord; Karoline Bragstad; Kathrine Stene-Johansen; Line Victoria Moen; Marie Paulsen Madsen; Olav Hungnes; Pedersen Benedikte Nevjen; Rasmus Riis Kopperud |
| EPI_ISL_1254622 | Department of Medical Microbiology, St. Olavs hospital | Norwegian Institute of Public Health, Department of Virology | Atiya R Ali; Debech Nadia; Engebretsen Serina Beate; Garcia Llorente Ignacio; Hilde Elshaug; Hilde Vollan; Jon Bråte; Kamilla Heddeland Instefjord; Karoline Bragstad; Kathrine Stene-Johansen; Marie Paulsen Madsen; Olav Hungnes; Pedersen Benedikte Nevjen; Rasmus Riis Kopperud |
| EPI_ISL_5429642 | Department of Microbiology, National Institute of Public Health of Kosovo | Department of Microbiology, National Institute of Public Health of Kosovo | Aferdita Kuqi-Hyseni; Blendi Jerliu; Donjeta Hajdari; Nazmi Mehmeti; Pranvera Abazi; Robert Ramadani; Xhevat Jakupi; Zana Deva |
| EPI_ISL_6841981 | Department of Microbiology, The University of Hong Kong | Department of Microbiology, The University of Hong Kong | Kelvin K.W. To; Kwok-Yung Yuen |
| EPI_ISL_1677920 | Department of Public Health Microbiology Ljubljana, National Laboratory for Health, Environment and Food | Department for Public Health Microbiology Ljubljana, National Laboratory for Health, Environment and Food | José Gonçalves; Katarina Prosenc; Martin Bosilj; Natasa Berginc; Tom Koritnik |
| EPI_ISL_1842071, EPI_ISL_2031216 | Department of Virology and Immunology, University of Helsinki and Helsinki University Hospital, Huslab Finland | Department of Virology, Faculty of Medicine, University of Helsinki, Helsinki, Finland | Essi Korhonen; Hanna Jarva; Hanna Liimatainen; Hannimari Kallio-Kokko; Harri Kangas; Hussein Alburkat; Jenni Virtanen; Maija Lappalainen; Maija Suvanto; Olli Vapalahti; Pekka Ellonen; Phuoc Truong; Ravi Kant; Sari Hannula; Satu Kurkela; Teemu Smura |
| EPI_ISL_927941, EPI_ISL_928361, EPI_ISL_928878, EPI_ISL_1024059, EPI_ISL_1868200, EPI_ISL_1868965, EPI_ISL_1870608, EPI_ISL_1890559, EPI_ISL_1896543, EPI_ISL_2025772, EPI_ISL_2026288, EPI_ISL_2296712, EPI_ISL_2296939, EPI_ISL_2297353, EPI_ISL_2298986, EPI_ISL_2299141, EPI_ISL_2300125 | | | |
| see above | Department of Virus and Microbiological Special Diagnostics, Statens Serum Institut, Copenhagen, Denmark | Aalborg University | Danish Covid-19 Genome Consortium |
| EPI_ISL_4739755 | Dept. of Medical Microbiology, Stavanger University Hospital, Helse Stavanger HF | Norwegian Institute of Public Health, Department of Virology | Atiya R Ali; Debech Nadia; Engebretsen Serina Beate; Garcia Llorente Ignacio; Hilde Elshaug; Hilde Vollan; Jon Bråte; Kamilla Heddeland Instefjord; Karoline Bragstad; Kathrine Stene-Johansen; Line Victoria Moen; Marie Paulsen Madsen; Olav Hungnes; Pedersen Benedikte Nevjen; Rasmus Riis Kopperud |
| EPI_ISL_2357998, EPI_ISL_4919852, EPI_ISL_8297713 | Dept. of Microbiology and Infection Control, Akershus University Hospital HF | Dept. of Microbiology and Infection Control, Akershus University Hospital HF | Alexander Hesselberg Løvestad; Hanne Berggreen; Hege Vangstein Aamot |
| EPI_ISL_8056909 | Dhaulagiri Hopital PCR COVID Lab | National Public Health Laboratory | National Public Health Labortaory Team/Dhaulagiri Hopital PCR COVID Lab |
| EPI_ISL_2455178 | Diagnostyka | 1. Tricity SARS-CoV-2 sequencing consortium: University of Gdansk, Medical University of Gdansk, Vaxican Ltd., Invicta Ltd. 2. National Institute of Public Health - National Institute of Hygiene, Warsaw, Poland | Celina Cybulska; Karolina Gackowska; Katarzyna Groth; Katarzyna Zacharczuk; Krystyna Bienkowska Szewczyk; Lukasz Rabalski; Maciej Grzybek; Maciej Kosinski; Magdalena Nowakowska; Marcin Lubocki; Małgorzata Sadkowska-Todys; Tomasz Wołkowicz |
| EPI_ISL_8285748 | Diagnostyka Sp. z o. o. Wrocław | Wojewodzka Stacja Sanitarno-Epidemiologiczna w Gorzowie Wielkopolskim | Elzbieta Justynska; Klaudia Kobendza-Wlodarczak; Patrycja Faberska and Marek Margol; Renata Siegel |
| EPI_ISL_4633123 | Diagnostyka. Laboratoria Medyczne. | 1. ViroGenetics - BSL3 Laboratory of Virology, Małopolska Centre of Biotechnology, Jagiellonian University; 2. Diagtron Laboratoria Łukasz Rąbalski | Gromowski, T.; Kowalski, M.; Labaj; Maciej Kosinski; Mazur-Panasiuk, N.; Natalia Derewonko; P.P.; Pyrc, K.; Rabalski L.; Rogalska-Kupiec M.; Swadzba J.; Sylwia Januszczak; Szulc, P.; Wydmanski, W. |
| EPI_ISL_5331701 | Diagnostyka. Laboratoria Medyczne. | 1. ViroGenetics - BSL3 Laboratory of Virology, Małopolska Centre of Biotechnology, Jagiellonian University; 2. genXone SA, Research & Development Laboratory | Aleksandra Gidlewicz; Anna Brylak; Gromowski, T.; Grzegorz Nowicki; Jakub Grabowski; Kowalski, M.; Labaj; Maciej Sykulski; Mazur-Panasiuk, N.; Michał Kaszuba; Natalia Drweska-Matelska; P.P.; Pyrc, K.; Ruslan Herasymenko; Sylwia Januszczak; Szulc, P.; Wydmanski, W.; Łukasz Krych |
| EPI_ISL_2403771, EPI_ISL_2433667, EPI_ISL_5804495 | Dianalabs SA | Genesupport | Geraldine Jost; Katia Jaton; Nadia Liassine; Tanguy ARAUD |
| EPI_ISL_5004040 | District Hospital | CDFD | Ashwin Dalal; Asmita Gupta; Divya Vashisht; Murali Bashyam; Nagamani Kammili; Vinay Donipadi |
| EPI_ISL_1489532, EPI_ISL_2284527, EPI_ISL_2332570, EPI_ISL_5007658 | Division of Emerging Infectious Diseases, Bureau of Infectious Diseases Diagnosis Control, Korea Disease Control and Prevention Agency | Division of Emerging Infectious Diseases, Bureau of Infectious Diseases Diagnosis Control, Korea Disease Control and Prevention Agency | Ae Kyung Park; Chae Young Lee; Eun-Jin Kim; Heui Man Kim; Il-Hwan Kim; Jeong-Ah Kim; Jeong-Min Kim; Jin Sun No |
| EPI_ISL_1502131, EPI_ISL_1502141 | Division of Medical Virology, National Health Laboratory Service (NHLS), Tygerberg Hospital / Stellenbosch University | Division of Medical Virology, Stellenbosch University and NHLS Tygerberg Hospital | Bronwyn Kleinhans; Gert van Zyl; Susan Engelbrecht; Wolfgang Preiser |
| EPI_ISL_2379462, EPI_ISL_2379496, EPI_ISL_2379497, EPI_ISL_2379498, EPI_ISL_2379546, EPI_ISL_2379646, EPI_ISL_2379647 | | | |
| see above | Dr S Raju, Director of Public Heath and Preventive Medicine | inStem NCBS - INSACOG | Uma Ramakrishnan Dasaradhi Palakodeti Aswin SaiNarain |
| EPI_ISL_6032030 | Dr S raju, Director of public heath and preventive medicine | inStem NCBS – INSACOG | Aswin SaiNarain; Dasaradhi Palakodeti; Uma Ramakrishnan |
| EPI_ISL_745112 | Dr Van Niekerk Hospital | National Health Laboratory Service (NHLS), Tygerberg | Bronwyn Kleinhans; Eduan Wilkindon; Gert van Zyl; Houriiyah Tegally; Kayla Delaney; Susan Engelbrecht; Tulio de Oliveira; Wolfgang Preiser |
| EPI_ISL_4483387 | Dutch COVID-19 response team | Erasmus Medical Center | Anne van der Linden; Annemiek van der Eijk; Bas Oude Munnink; Corine GeurtsvanKessel; David Nieuwenhuijse; Emmanuelle Munger; Irina Chestakova; Marion Koopmans; Marjan Boter; Reina Sikkema; Richard Molenkamp; on behalf of the Dutch national COVID-19 response team. |
| EPI_ISL_2030459, EPI_ISL_8191040 | Dutch COVID-19 response team | Medical Microbiology, Maastricht University Medical Centre | Brian van der Veer*; Carmen Reumkens; Christian Hoebe; Erik Beuken; Jozef Dingemans*; Lieke van Alphen; Paul Savelkoul |
| EPI_ISL_1232268, EPI_ISL_1962046, EPI_ISL_1962290, EPI_ISL_1962951, EPI_ISL_1962952, EPI_ISL_2094359, EPI_ISL_2094360, EPI_ISL_2094366, EPI_ISL_2094367, EPI_ISL_2094371, EPI_ISL_2220135, EPI_ISL_2220453, EPI_ISL_2220458, EPI_ISL_2220461, EPI_ISL_2302439, EPI_ISL_2302933, EPI_ISL_2303943, EPI_ISL_2303944, EPI_ISL_2303946, EPI_ISL_2303947, EPI_ISL_2405403, EPI_ISL_2405469, EPI_ISL_2405492, EPI_ISL_2405505, EPI_ISL_2405516, EPI_ISL_2405527, EPI_ISL_2405557, EPI_ISL_2405582, EPI_ISL_2406147, EPI_ISL_2406319, EPI_ISL_2406427, EPI_ISL_2406455, EPI_ISL_2474731, EPI_ISL_2475352, EPI_ISL_2476272, EPI_ISL_2476273, EPI_ISL_2476274, EPI_ISL_2476275, EPI_ISL_2476277, EPI_ISL_2476278, EPI_ISL_2476279, EPI_ISL_2476280, EPI_ISL_2476281, EPI_ISL_2476282, EPI_ISL_2476283, EPI_ISL_2476284, EPI_ISL_2476285, EPI_ISL_2476286, EPI_ISL_2476287, EPI_ISL_2476288, EPI_ISL_4401056, EPI_ISL_4840718, EPI_ISL_4845426, EPI_ISL_5206516, EPI_ISL_5206779, EPI_ISL_5430614, EPI_ISL_5819539, EPI_ISL_6841610, EPI_ISL_7470340, EPI_ISL_7785155, EPI_ISL_7786217 | | | |
| see above | Dutch COVID-19 response team | National Institute for Public Health and the Environment (RIVM) | Adam Meijer; AnneMarie van den Brandt; Annelies Kroneman; Bas van der Veer; Chantal Reusken; Dennis Schmitz; Dirk Eggink; Euníce Then; Florian Zwagemaker; Harry Vennema; Ivo van Walle; James Groot; Jeroen Cremer; Jil Kocken; Karim Hajji; Kim Freriks; Linda van Someren; Linda van de Nes; Lisa Wijsman; Lynn Aarts; Melissa van Tuil; Ryanne Jaarsma; Sanne Bos; Sharon van den Brink; Sjoerd Kuiling; Stijn van Rossum; on behalf of the national COVID-19 response team |
| EPI_ISL_1582978 | E.S.E. HOSPITAL SAN JOSE DE MAICAO | Instituto Nacional de Salud- Dirección de Investigación en Salud Pública | Carlos Franco-Muñoz; Carmen Osorio; Diana Malo; Diego A. Álvarez-Díaz; Diego Andrés Prada; Gerardo Santamaría; Hector Alejandro Ruiz-Moreno; Jhonnatan Reales-González; Juan Camilo Martinez; Julian Naizaque; Katherine Laiton-Donato; Lisseth Pardo; Magdalena Wiesner; Marcela Mercado-Reyes; Maria T. Herrera-Sepúlveda; Marta Lopez Blanco; Martha Lucia Ospina Martinez; Paola Rojas; Sergio Gomez; Sheryll Corchuelo; Ángela Alarcon Cruz |
| EPI_ISL_5564769 | ELISABETH PHARMACON | University Hospital Brno, CMBG | Bezdicek Matej; Dolejska Monika; Kristyna Dufkova; Lengerova Martina; Svaton Jan |
| EPI_ISL_6845886 | EUROFINS CBM 69 MEDIPOLE | CNR Virus des Infections Respiratoires - France SUD | Antonin Bal; Bruno Lina; Bruno Simon; Gregory Destras; Gwendolyne Burfin; Hadrien Regue; Laurence Josset; Martine Valette; Quentin Semanas |
| EPI_ISL_1811676, EPI_ISL_1816573 | EXCITE Lab | Andersen lab at Scripps Research | Abigail Schnapper; Alexandre Bolze; Alice Summerfield; Angela Scioscia; Celena Andrade; Charlotte Rivera-Garcia; Cheryl Anderson; Chip Schooley; David Becker; Efren Sandoval; Elizabeth Cirulli; Francisco Tanudjaja; Geraint Levan; Greg Humphrey; Helena Tubb; James Lu + SEARCH; Jason Nguyen; Jimmy Ramirez; Kelly Schiabor Barrett; Magnus Isaksson; Marc Laurent; Natasha Martin; Nicole L Washington; Ryan Cho; Sawyer Farmer; Sherry Wang; Simon White; Smruthi Karthikeyan; Tommy Valles + SEARCH; Tyler Cassens; William Lee |
| EPI_ISL_7123680, EPI_ISL_7738576 | Edmonton Provincial Lab | Alberta Precision Labs (APL) | Buss; Croxen M; Deo A; Dieu P; E; Ferrato C; Gill K; Khan F; Koleva P; Li V; Lloyd C; Lynch T; Ma R; Murphy S; Pabbaraju K; Shokoples S; Thayer J; Tipples G; Whitehouse M; Wong A; Yu C; Zelyas N |
| EPI_ISL_4561211 | Edmonton Provincial Lab | Public Health Agency of Canada (PHAC) National Microbiology Laboratory | Buss; Croxen M; Deo A; Dieu P; E; Ferrato C; Gill K; Khan F; Koleva P; Li V; Lloyd C; Lynch T; Ma R; Murphy S; Pabbaraju K; Shokoples S; Thayer J; Tipples G; Whitehouse M; Wong A; Yu C; Zelyas N |
| EPI_ISL_745119, EPI_ISL_745178 | Eerste River Hospital wc ERH | National Health Laboratory Service (NHLS), Tygerberg | Bronwyn Kleinhans; Eduan Wilkindon; Gert van Zyl; Houriiyah Tegally; Kayla Delaney; Susan Engelbrecht; Tulio de Oliveira; Wolfgang Preiser |
| EPI_ISL_1444970 | Erie County Public Health (ECPHL) | University at Buffalo Genomics and Bioinformatics Core | Alyssa Pohlman; Amanda Boccolucci; Brandon Marzullo; Donald Yergeau; Jennifer Surtees; Jonathan Bard; Natalie Lamb; Norma Nowak |
| EPI_ISL_1355181, EPI_ISL_1845854, EPI_ISL_2110547, EPI_ISL_4937282, EPI_ISL_8220440, EPI_ISL_8240359 | Eurofins LifeCodexx GmbH | Robert Koch Institute | |
| EPI_ISL_2316409, EPI_ISL_8235164, EPI_ISL_8235173, EPI_ISL_8235401 | Eurofins MVZ Labor Gelsenkirchen | Robert Koch Institute | |
| EPI_ISL_7547732 | Everight diagnostics (Abuja) | National Reference Laboratory, Nigeria Centre for Disease Control | Catherine Okoi; Chimaobi Chukwu; Dr Ifedayo Adetifa; Dr Ndodo Nnaemeka; Dr Omoare Adesuyi; Nwando Mba; Olajumoke Babatunde; Olusola Anuoluwapo Akanbi; Oyeronke Ayansola |
| EPI_ISL_2341694 | FM Medical College, Balasore | Institute of Life Sciences - INSACOG | Ajay Parida; Amol M. Kanampalliwar; Arup Ghosh; Atimukta Jha; INSACOG Consortium; Omprakash Shiriwas; Punit Prasad; Rajeeb Swain; Rupesh Dash; Safal Walia; Sana Fatma; Shifu Aggarwal; Sunil K. Raghav |
| EPI_ISL_2158349 | FUNDACION HOSPITAL SAN PEDRO | Instituto Nacional de Salud- Dirección de Investigación en Salud Pública | Carlos Franco-Muñoz; Carmen Osorio; Diana Malo; Diego A. Álvarez-Díaz; Diego Andrés Prada; Gerardo Santamaría; Hector Alejandro Ruiz-Moreno; Jhonnatan Reales-González; Jorge Rivera; Juan Camilo Martinez; Julian Naizaque; Katherine Laiton-Donato; Lisseth Pardo; Magdalena Wiesner; Marcela Mercado-Reyes; Maria T. Herrera-Sepúlveda; Marta Lopez Blanco; Martha Lucia Ospina Martinez; Paola Rojas; Sergio Gomez; Sheryll Corchuelo; Ángela Alarcon Cruz |
| EPI_ISL_6193073 | Fakultní nemocnice u sv. Anny v Brně | Fakultní nemocnice u sv. Anny v Brně | Filip Růžička; Kareřina Zettlová; Milada Dvořáčková. Jan Vrbský |

| | | | |
|---|---|---|---|
| EPI_ISL_2140672, EPI_ISL_2375947 | False Bay Hospital wc FBH | NHLS/UCT | Arash Iranzadeh; Bruna Galvao; Carolyn Williamson; Deelan Doolabh; Diana Hardie; Innocent Mudau; Kruger Marais; Lynn Tyers; Marvin Hsiao; Stephen Korsman |
| EPI_ISL_2229947 | Florida Bureau of Public Health Laboratories | Florida Bureau of Public Health Laboratories | Jason Blanton; Sarah Schmedes |
| EPI_ISL_2441850, EPI_ISL_2441884, EPI_ISL_2480021, EPI_ISL_2480190, EPI_ISL_4183726, EPI_ISL_4184046, EPI_ISL_4243111, EPI_ISL_4243899, EPI_ISL_4243932, EPI_ISL_4245318, EPI_ISL_4358771, EPI_ISL_4455862, EPI_ISL_4457159, EPI_ISL_4535263, EPI_ISL_4566086, EPI_ISL_4567880, EPI_ISL_4828069, EPI_ISL_4901726, EPI_ISL_4901880, EPI_ISL_5091759, EPI_ISL_5092814, EPI_ISL_5234081, EPI_ISL_5234678, EPI_ISL_5308178, EPI_ISL_5319169, EPI_ISL_5320447, EPI_ISL_5378600, EPI_ISL_5587486, EPI_ISL_5615261, EPI_ISL_5615518, EPI_ISL_5891261, EPI_ISL_6000855, EPI_ISL_6001241, EPI_ISL_6071610, EPI_ISL_6071740, EPI_ISL_6072009, EPI_ISL_6340995, EPI_ISL_6341585, EPI_ISL_6511699, EPI_ISL_6623665, EPI_ISL_6628521, EPI_ISL_6654549, EPI_ISL_6892297, EPI_ISL_7153926, EPI_ISL_7313060, EPI_ISL_7447749, EPI_ISL_7476157, EPI_ISL_7567233, EPI_ISL_7567268, EPI_ISL_7567830, EPI_ISL_7671186, EPI_ISL_7671259, EPI_ISL_7847319, EPI_ISL_8019467, EPI_ISL_8019535, EPI_ISL_8022555, EPI_ISL_8080961, EPI_ISL_8091653, EPI_ISL_8157113, EPI_ISL_8183123, EPI_ISL_8183332, EPI_ISL_8183357, EPI_ISL_8281385, EPI_ISL_8282314, EPI_ISL_8282768, EPI_ISL_8284534, EPI_ISL_8284977, EPI_ISL_8285006, EPI_ISL_8285042, EPI_ISL_8285237, EPI_ISL_8285287, EPI_ISL_8292776, EPI_ISL_8293314, EPI_ISL_8293672 | | | |
| see above | Fulgent Genetics | Centers for Disease Control and Prevention Division of Viral Diseases, Pathogen Discovery | Adrian Paskey; Becky Tsai; Benafsh Sapra; Benjamin Rambo-Martin; Christopher Gulvick; Clinton Paden; Clinton R. Paden; Dakota Howard; Darlene Wagner; Dhwani Batra; Doreen Ng; Duncan MacCannell; Erisa Sula; Harry Gao; James Xie; Jason Caravas; John Gao; Joseph Fierro; Kara Moser; Kristine Lacek; Matthew Schmerer; Mickey Li; Peter Cook; Peter W. Cook; Scott Sammons; Shatavia Morrison; Tymeckia Kendall; Victoria Caban Figueroa; Yan Meng; Yvette Unoarumhi |
| EPI_ISL_1708318 | Fundação Hospitalar de Hematologia e Hemoterapia do Amazonas | Laboratório de Estudos de Vírus Emergentes | Adriana S. S. Duarte; Alessandro S. Farias; Arilson Bernardo S. P. Gomes; Audrey B. Zangirolami; Bruno D. Benites; Camila A.M. Silva; Camila L. Simeoni; Carolina Costa-Lima; Cecilia C. Camilo; Chieh-Hsi Wu; Christopher Dye; Clarice W. Arns; Daniel A. Toledo-Teixeira; Darlan S. Candido; Erika R. Manuli; Esmenia C. Rocha; Ester C. Sabino; Fabia; Fernando R. Spilki; Flavia C. Sales; Giulia M. Ferreira; Grazielle C. Maktura; Henrique Marques-Souza; Ingra M. Claro; Jaqueline G. de Jesus; Karina Bispo-dos-Santos; Lais D. Coimbra; Leandro M. Souza; Lucas A.M. Franco; Lucas I. Buscaratti; Luciana S. Mofatto; Magnun N.N. Santos; Marcelo A.S. Mori; Marcelo Addas-Carvalho; Maria L. Moretti; Mariana C. Pinho; Mariana S. Ramundo; Mariene R. Amorim; Michael S. Diamond; Myuki A.E. Crispim; Natalia S. Brunetti; Nelson Gaburo; Nuno R. Faria; Oliver G. Pybus; Pamela S. Andrade; Pierina L. Parise; Priscilla P. Barbosa; Rafael E. Marques; Renata Sesti-Costa; Rodrigo N. Angerami; Thais M. Coletti; Vitor A. Costa; William M. Souza |
| EPI_ISL_7776984 | Furst Medical Laboratory | Norwegian Institute of Public Health, Department of Virology | Atiya R Ali; Debech Nadia; Engebretsen Serina Beate; Garcia Llorente Ignacio; Hilde Elshaug; Hilde Vollan; Jon Bråte; Kamilla Heddeland Instefjord; Karoline Bragstad; Kathrine Stene-Johansen; Line Victoria Moen; Marie Paulsen Madsen; Olav Hungnes; Pedersen Benedikte Nevjen; Rasmus Riis Kopperud |
| EPI_ISL_5637812 | GH A.CHENEVIER-H.MONDOR | Department of Virology, Henri Mondor University Hospital, Assistance Publique Hôpitaux de Paris, Université Paris-Est Créteil, INSERM U955 | Alexandre Soulier; Christophe Rodriguez; Elisabeth Trawinski; Guillaume Gricourt; Jean-Michel Pawlotsky; Melissa N'Debi; Slim Fourati; Vanessa Demontant |
| EPI_ISL_6938613 | GMERS Medical College, Valsad | Gujarat Biotechnology Research Centre | Apurvasinh Puvar; Bhadreshsinh Gohil; Chaitanya Joshi; Dinesh Kumar; Janvi Raval; Madhvi Joshi; Nimesh Patel; Nitin Savaliya; Nitin Shukla; Priyank Chavda; Ramesh Pandit; Sonal Sharma; Vicky Gandhi; Zarna Patel |
| EPI_ISL_2017757 | GMERS, Government Medical College (GMC), Gotri | Gujarat Biotechnology Research Centre | Bithika Duttaroy; Chaitanya Joshi; Dinesh Kumar; Janvi Raval; Madhvi Joshi; Nitesh Shah; Nitin Savaliya; Ramesh Pandit; Sonal Sharma; Twinkle Soni; Umang Mishra; Zarna Patel; Zuber Saiyed |
| EPI_ISL_5816145, EPI_ISL_6903858 | Gandhi Hospital | Centre for DNA Fingerprinting and Diagnostics | Arunkumar Karunanidhi; Ashwin Dalal; Asmita Gupta; Divya Vashisht; Murali Bashyam; Nagamani Kammili; Vinay Donipadi |
| EPI_ISL_6858569 | GeneMe Labs, ul. Potokowa 10A, 80-283 Gdańsk | 1. Tricity SARS-CoV-2 sequencing consortium: University of Gdansk, Medical University of Gdansk, Vaxican Ltd., Invicta Ltd. 2. National Institute of Public Health - National Institute of Hygiene, Warsaw, Poland | Celina Cybulska; Karolina Gackowska; Katarzyna Groth; Katarzyna Zacharczuk; Krystyna Bienkowska Szewczyk; Lukasz Rabalski; Maciej Grzybek; Maciej Kosinski; Magdalena Nowakowska; Marcin Lubocki; Małgorzata Sadkowska-Todys; Tomasz Wołkowicz |
| EPI_ISL_1300520, EPI_ISL_1321476, EPI_ISL_1321477, EPI_ISL_1321487, EPI_ISL_1470428, EPI_ISL_1470434, EPI_ISL_1470453, EPI_ISL_1470458, EPI_ISL_1470459, EPI_ISL_1470462, EPI_ISL_1470468, EPI_ISL_1470502, EPI_ISL_1470509, EPI_ISL_1470514, EPI_ISL_1470539, EPI_ISL_1470559, EPI_ISL_1534549, EPI_ISL_1534561, EPI_ISL_1534582, EPI_ISL_1534594, EPI_ISL_1541022, EPI_ISL_1541049, EPI_ISL_1633370, EPI_ISL_1633474, EPI_ISL_1633477, EPI_ISL_1633517, EPI_ISL_1633547, EPI_ISL_1712331, EPI_ISL_1712356, EPI_ISL_2009240, EPI_ISL_2009246, EPI_ISL_2009265, EPI_ISL_2009288, EPI_ISL_2009560, EPI_ISL_2009626, EPI_ISL_2009635, EPI_ISL_2009645, EPI_ISL_2391020, EPI_ISL_2391024, EPI_ISL_2391025, EPI_ISL_2391056, EPI_ISL_2391112, EPI_ISL_2391222, EPI_ISL_2391232, EPI_ISL_2391294, EPI_ISL_2391295, EPI_ISL_2391309, EPI_ISL_2391337, EPI_ISL_2391364, EPI_ISL_2391365, EPI_ISL_2391369, EPI_ISL_2391370, EPI_ISL_2391371, EPI_ISL_2391373, EPI_ISL_2391378, EPI_ISL_2391379, EPI_ISL_2391386, EPI_ISL_2391387, EPI_ISL_2391399, EPI_ISL_2391423, EPI_ISL_2543506, EPI_ISL_2543514, EPI_ISL_2543516, EPI_ISL_2543518, EPI_ISL_2543521, EPI_ISL_2543530, EPI_ISL_2543531, EPI_ISL_2543532, EPI_ISL_2543533, EPI_ISL_2543535, EPI_ISL_2543538, EPI_ISL_2543541, EPI_ISL_2543556, EPI_ISL_2543578, EPI_ISL_2543579, EPI_ISL_2543587, EPI_ISL_2543589, EPI_ISL_2543592, EPI_ISL_2543595, EPI_ISL_2543597, EPI_ISL_2543599, EPI_ISL_2543601, EPI_ISL_2543605, EPI_ISL_2543609, EPI_ISL_2543629, EPI_ISL_2543637, EPI_ISL_2543650, EPI_ISL_2543654, EPI_ISL_2543659, EPI_ISL_2543660, EPI_ISL_2543663 | | | |
| see above | Genetica Molecular and Subdepartamento de Virologia ISP Chile | Instituto de Salud Publica de Chile | Andres Castillo; Barbara Parra; Constanza Campano; Gisselle Barra; Jaime Lagos; Javier Tognarelli; Jorge Fernandez; Karen Orostica; Loredana Arata; Patricia Bustos; Rodrigo Fasce; Soledad Ulloa |
| EPI_ISL_6892620 | Germano de Sousa | Instituto Nacional de Saude (INSA) | Borges et al |
| EPI_ISL_5622746, EPI_ISL_5623514, EPI_ISL_6339838, EPI_ISL_8255660 | Gravity Diagnostics, LLC | Gravity Diagnostics, LLC | Gravity Diagnostics |
| EPI_ISL_1040673, EPI_ISL_1040697, EPI_ISL_1040704, EPI_ISL_1040722, EPI_ISL_1040745, EPI_ISL_1534315, EPI_ISL_1534323, EPI_ISL_1706560, EPI_ISL_1706561, EPI_ISL_2140683, EPI_ISL_2375965 | | | |
| see above | Groote Schuur Hospital wc GSH | NHLS/UCT | Arash Iranzadeh; Bruna Galvao; Carolyn Williamson; Deelan Doolabh; Diana Hardie; Emmanuel SJ; Innocent Mudau; Kruger Marais; Lynn Tyers; Marvin Hsiao; Stephen Korsman; Tegally H; de Oliveira T |
| EPI_ISL_960109, EPI_ISL_960136 | Groote Schuur Hospital wc GSH | National Health Laboratory Service/UCT | Arash Iranzadeh; Bruna Galvao; Carolyn Williamson; Deelan Doolabh; Diana Hardie; Innocent Mudau; Kruger Marais; Lynn Tyers; Marvin Hsiao; Stephen Korsman |
| EPI_ISL_2339796, EPI_ISL_2339799, EPI_ISL_2339801, EPI_ISL_2339802, EPI_ISL_2339805, EPI_ISL_2339810, EPI_ISL_2339812, EPI_ISL_2339813, EPI_ISL_2339814, EPI_ISL_2339815, EPI_ISL_2339845, EPI_ISL_2339847, EPI_ISL_2339850, EPI_ISL_2339851, EPI_ISL_2339853 | | | |
| see above | Grupo de Investigación en Enfermedades Tropicales del Ejército (GINETEJ), Laboratorio de Referencia e Investigación, Dirección de Sanidad Ejército, Bogotá, Colombia | Centro de Investigaciones en Microbiología y Biotecnología-UR (CIMBIUR), Facultad de Ciencias Naturales, Universidad del Rosario, Bogotá, Colombia | Camilo A. Correa-Cárdenas; Carolina Oliveros; Claudia Méndez; Elizabeth K. Márquez; Frank de los Santos Ortiz; Juan David Ramírez; Julie Pérez; Lorena Albarracín; Luz H. Patiño; Maria Clara Duque; Marina Muñoz; María Teresa Alvarado; Nathalia Ballesteros; Sergio Castañeda; Sergio Gutíerrez-Riveros; Yanira Romero; Zulma Cucunubá |
| EPI_ISL_1040675 | Guguletu CHC wc GDH | NHLS/UCT | Arash Iranzadeh; Bruna Galvao; Carolyn Williamson; Deelan Doolabh; Diana Hardie; Innocent Mudau; Kruger Marais; Lynn Tyers; Marvin Hsiao; Stephen Korsman |
| EPI_ISL_7846620 | H Braga | Instituto Nacional de Saude (INSA) and Instituto Gulbenkian de Ciencia (IGC) | Borges et al |
| EPI_ISL_6900139, EPI_ISL_6900142, EPI_ISL_6900143, EPI_ISL_7740782, EPI_ISL_7740834, EPI_ISL_7740840 | HELEN JOSEPH LABORATORY | National Institute for Communicable Diseases of the National Health Laboratory Service | Amoako DG; Bhiman JN; Everatt J; Ismail A; Mahlangu B; Mnguni A; Mohale T; Ntuli N; Scheepers C; Wolter N |
| EPI_ISL_1652551 | HELIX LLC | WHO National Influenza Centre Russian Federation | Andrey Komissarov; Anna Ivanova; Artem Fadeev; Daria Danilenko; Dmitry Lioznov; Elena Nabieva; Georgii Bazykin; Ksenia Safina; Kseniya Komissarova; Maria Pisareva; Maria Timofeeva; Tamila Musaeva; Veronika Eder |
| EPI_ISL_1821074 | HIC | Instituto Nacional de Salud- Dirección de Investigación en Salud Pública | Carlos Franco-Muñoz; Carmen Osorio; Diana Malo; Diego A. Álvarez-Díaz; Diego Andrés Prada; Gerardo Santamaría; Hector Alejandro Ruiz-Moreno; Jhonnatan Reales-González; Jorge Rivera; Juan Camilo Martinez; Julian Naizaque; Katherine Laiton-Donato; Lisseth Pardo; Magdalena Wiesner; Marcela Mercado-Reyes; Maria T. Herrera-Sepúlveda; Marta Lopez Blanco; Martha Lucia Ospina Martinez; Paola Rojas; Sergio Gomez; Sheryll Corchuelo; Ángela Alarcon Cruz |
| EPI_ISL_1821071 | HIGUERA ESCALANTE | Instituto Nacional de Salud- Dirección de Investigación en Salud Pública | Carlos Franco-Muñoz; Carmen Osorio; Diana Malo; Diego A. Álvarez-Díaz; Diego Andrés Prada; Gerardo Santamaría; Hector Alejandro Ruiz-Moreno; Jhonnatan Reales-González; Jorge Rivera; Juan Camilo Martinez; Julian Naizaque; Katherine Laiton-Donato; Lisseth Pardo; Magdalena Wiesner; Marcela Mercado-Reyes; Maria T. Herrera-Sepúlveda; Marta Lopez Blanco; Martha Lucia Ospina Martinez; Paola Rojas; Sergio Gomez; Sheryll Corchuelo; Ángela Alarcon Cruz |
| EPI_ISL_2187882 | HLAGYN - Laboratorio de Imunologia de Transplantes de Goias | HLAGYN - Laboratorio de Imunologia de Transplantes de Goias | Alessandro Leonardo Alvares Magalhaes; Daniel Ferreira de Sousa; Erika Lopes Rocha Batista; Fernando Antonio Vinhal dos Santos; Frederico Rodrigues Vinhal; Lucas Carlos Gomes Pereira; Sabrina Sara Moreira Duarte |
| EPI_ISL_1904994 | HOPITAL SAINT ANDRE | CNR Virus des Infections Respiratoires - France SUD | Antonin Bal; Bruno Lina; Bruno Simon; Gregory Destras; Gwendolyne Burfin; Hadrien Regue; Laurence Josset; Martine Valette; Quentin Semanas |
| EPI_ISL_1940001, EPI_ISL_1940002, EPI_ISL_1940008, EPI_ISL_1940013, EPI_ISL_1940014, EPI_ISL_1940015 | HOSAHALLI | inStem NCBS - INSACOG | Uma Ramakrishnan Dasaradhi Palakodeti Aswin SaiNarain |
| EPI_ISL_1967071 | HOSP MUN CIDADE TIRADENTES CARMEN PRUDENTE | Instituto Butantan / ESALQ-USP (Piracicaba) | Antonio Jorge Martins; Bianca Cechetto Carlos. Mendelics: Bibiana Santos; Claudia Renata dos Santos Barros; Cíntia Bittar; David Schlesinger. Hemocentro Ribeirão Preto: Simone Kashima; Debora Botequio Moretti; Elaine Cristina Marqueze; Elaine Vieira dos Santos; Elisangela Chicaroni Mattos; Erika Freitas; Evandra Strazza Rodrigues; Felipe Allan da Silva da Costa; Flavia Aburjaile; Fábio Sossai Possebon; Guilherme Campos; Guilherme Targino Valente; Heidge Fukumasu. USP-Botucatu: Rejane Maria Tommasini Grotto; Helena Lage Ferreira; Instituto Butantan: Dimas Tadeu Covas; Jardelina de Souza Todao Bernardino; Jayme A. Souza-Neto; Jessika Cristina Chagas Lesbon; Jorge A. Petroli Marchesi; José Salvatore Leister Patané; João Paulo Kitajima; João Pessoa Araújo Jr.; Leila Sabrina Ullmann; Loyze Paola Oliveira de Lima; Luiz Aurelio de Campos Crispin. Centro de Genômica Funcional da ESALQ: Luiz Lehmann Coutinho; Luiz Carlos Junior de Alcantara; Luiz Sacchetto; Maisa C. Pereira Parra; Maria Carolina Elias; Marta Giovanetti; Marília Moraes; Maurício Lacerda Nogueira. Prefeitura de Sao Paulo: Melissa Palmieri.; Patricia Akemi Assato; Paula Rahal; Paulo Inacio da Costa; Rafael dos Santos Bezerra; Raquel de Lello Rocha Campos Cassano. NGS Soluções Genômicas: Pilar Drummond Sampaio Corrêa Mariani. FZEA-USP Pirassununga: Mirele Daiana Poleti; Raul Machado Neto; Ricardo Augusto Brassaloti; Ricardo Haddad; Rodrigo Tocantins Calado. FAMERP-SJRP: Cecília Artico Banho; Sandra Coccuzzo Sampaio; Svetoslav Nanev Slavov; Vagner Fonseca; Vincent Louis Viala |
| EPI_ISL_1821070 | HOSPITAL DEPARTAMENTAL MARIA INMACULADA | Instituto Nacional de Salud- Dirección de Investigación en Salud Pública | Carlos Franco-Muñoz; Carmen Osorio; Diana Malo; Diego A. Álvarez-Díaz; Diego Andrés Prada; Gerardo Santamaría; Hector Alejandro Ruiz-Moreno; Jhonnatan Reales-González; Jorge Rivera; Juan Camilo Martinez; Julian Naizaque; Katherine Laiton-Donato; Lisseth Pardo; Magdalena Wiesner; Marcela Mercado-Reyes; Maria T. Herrera-Sepúlveda; Marta Lopez Blanco; Martha Lucia Ospina Martinez; Paola Rojas; Sergio Gomez; Sheryll Corchuelo; Ángela Alarcon Cruz |
| EPI_ISL_2534428 | HOSPITAL UNIVERSITARIO SON ESPASES | HOSPITAL UNIVERSITARIO SON ESPASES | Hospital Universitario Son Espases; Servicio de Microbiología |
| EPI_ISL_1684024, EPI_ISL_2178977, EPI_ISL_2227876, EPI_ISL_2365428, EPI_ISL_2365429, EPI_ISL_2365432, EPI_ISL_2365435, EPI_ISL_2365436, EPI_ISL_2365437, EPI_ISL_2365438, EPI_ISL_2365439, EPI_ISL_2365440 | | | |
| see above | HOSPITAL UNIVERSITARIO VIRGEN DE LA ARRIXACA | Instituto de Salud Carlos III | A. Monzón; ANTONIO; F. Casas; I. Jiménez; I. MORENO DOCON; I. MORENO PARRADO; I.MORENO PARRADO; Iglesias-Caballero; LAURA; M. Camarero; M. Sandonís; P. Zaballos; S. Camarero; S. Cuesta; S. Iglesias-Caballero; S. Pozo; S. Varona; Sandonís; V. Camarero; V. Vázquez-Morón; Vázquez-Morón |
| EPI_ISL_6113844 | Hassan Institute of Medical Sciences | INSACOG-KA , NIMHANS | Ananthapadmanabha Kotambail; Anita S Desai; Chetan G K; Ellango Ramasamy; Gautham Arunachal Udupi; Mahesh Kumar.C.S; Sony Sharma; V Ravi |
| EPI_ISL_7337440, EPI_ISL_7337444, EPI_ISL_7337451, EPI_ISL_7337453 | Helen Joseph Laboratory | National Institute for Communicable Diseases of the National Health Laboratory Service | Amoako DG; Bhiman JN; Everatt J; Ismail A; Mahlangu B; Mnguni A; Mohale T; Ntuli N; Scheepers C |

| | | | |
|---|---|---|---|
| EPI_ISL_5240584, EPI_ISL_6335737, EPI_ISL_6336627, EPI_ISL_6407427, EPI_ISL_6407792, EPI_ISL_6633069, EPI_ISL_6633223, EPI_ISL_6633498, EPI_ISL_7128557, EPI_ISL_7496832, EPI_ISL_7696024, EPI_ISL_7697330, EPI_ISL_7942504, EPI_ISL_7942528, EPI_ISL_8181500 | | | |
| see above | Helix | Centers for Disease Control and Prevention Division of Viral Diseases, Pathogen Discovery | Benjamin Rambo-Martin; Christopher Gulvick; Clinton Paden; Dakota Howard; Dhwani Batra; Duncan MacCannell; Erisa Sula; Helix CA; Jason Caravas; Kristine Lacek; Matthew Schmerer; Peter Cook; Scott Sammons; Shatavia Morrison; Tymeckia Kendall; Victoria Caban Figueroa; Yvette Unoarumhi |
| EPI_ISL_1340400, EPI_ISL_1479253, EPI_ISL_1512076, EPI_ISL_1581369, EPI_ISL_1734626, EPI_ISL_1734962, EPI_ISL_2247513, EPI_ISL_2269652, EPI_ISL_2270494, EPI_ISL_2320285, EPI_ISL_2440340, EPI_ISL_2440421 | | | |
| see above | Helix/Illumina | Centers for Disease Control and Prevention Division of Viral Diseases, Pathogen Discovery | Adrian Paskey; Alexandre Bolze; Ary Ascencio; Ben L. Rambo-Martin; Benjamin Rambo-Martin; Brad Sickler; Charlotte Rivera-Garcia; Christine Tran; Christopher Gulvick; Clinton R. Paden; Dakota Howard; Darlene Wagner; David Becker; Dhwani Batra; Duncan MacCannell; Efren Sandoval; Eileen de Feo; Elizabeth Cirulli; Eric Allen; Geraint Levan; James Lu; Jan Antico; Jason Caravas; Jason Nguyen; Jimmy Ramirez; Jingtao Liu; Kara Moser; Kelly Schiabor Barrett; Kim Gietzen; Magnus Isaksson; Marc Laurent; Matthew Schmerer; Matthew Tolentino; Nicole L. Washington; Peter W. Cook; Phil Febbo; Ryan Cho; Scott Sammons; Shannon Wickline; Shatavia Morrison; Sherry Wang; Simon White; Summer Galloway; Suxiang Tong; Tyler Cassens; William Lee; Yvette Unoarumhi |
| EPI_ISL_850594 | Helix/Illumina | Genomics and Discovery, Respiratory Viruses Branch, Division of Viral Diseases, Centers for Disease Control and Prevention | Alexandre Bolze; Ary Ascencio; Ben L. Rambo-Martin Eileen de Feo; Brad Sickler; Charlotte Rivera-Garcia; Christine Tran; Clinton R. Paden; David Becker; Dhwani Batra; Duncan MacCannell; Efren Sandoval; Elizabeth Cirulli; Eric Allen; Geraint Levan; James Lu; Jan Antico; Jason Nguyen; Jimmy Ramirez; Jingtao Liu; Kelly Schiabor Barrett; Kim Gietzen; Magnus Isaksson; Marc Laurent; Matthew Tolentino; Nicole L. Washington; Peter W. Cook; Phil Febbo; Ryan Cho; Shannon Wickline; Sherry Wang; Simon White; Summer Galloway; Suxiang Tong; Tyler Cassens; William Lee |
| EPI_ISL_7621721 | Histopath | NSW Health Pathology - Institute of Clinical Pathology and Medical Research; Westmead Hospital; University of Sydney | Arnott A.; Draper J.; Gall M.; Martinez E.; Rockett R.; Sintchenko V.; on behalf of ICPMR |
| EPI_ISL_6590782 | Home Quarantine Taskforce | Hong Kong Department of Health | Alan K.L. Tsang; Edman T.K. Lam; Ken H.L. Ng; Peter C.W. Yip; Rickjason C.W. Chan |
| EPI_ISL_7795897, EPI_ISL_8206250 | Hopital | National Reference Center for Viruses of Respiratory Infections, Institut Pasteur, Paris | Angela Brisebarre; Camille Capel; Christophe Malabat; Corinne Maufrais; Etienne Simon-Lorière; Frédéric Lemoine; Isabelle WORCEL; Julien Fumey; Louise Lefrançois; Marianne BURGARD; Marion Barbet; Maud Vanpeene; Méline Bizard; Slim El Khiari; Sylvie Behillil; Sylvie Van der Werf; Vincent Enouf |
| EPI_ISL_1443196, EPI_ISL_1443197, EPI_ISL_1443198 | Hospital Aliança | Hospital São Rafael - IDOR | Aquiles Assunção Camelier; Aurea Angelica Paste; Bruno Solano de Freitas Souza; Carolina Kymie Vasques Nonaka; Elves A.P. Maciel; Isadora Cristina de Siqueira; Karoline Almeida Félix de Sousa; Margarida Celia L. C. Neves; Tiago Gräf; Victor Costa Araujo; Yasmin Santos Freitas Macêdo |
| EPI_ISL_1477056, EPI_ISL_1647948, EPI_ISL_1647949, EPI_ISL_1647950, EPI_ISL_1647956, EPI_ISL_1964218, EPI_ISL_2080925, EPI_ISL_2080928, EPI_ISL_2178834, EPI_ISL_2402462, EPI_ISL_2402466, EPI_ISL_2402498, EPI_ISL_2402505, EPI_ISL_2402508, EPI_ISL_2402528, EPI_ISL_6781211 | | | |
| see above | Hospital General Universitario Gregorio Marañón | Hospital General Universitario Gregorio Marañón | Cristina Rodriguez-Grande; Darío García de Viedma; Jorge Rodríguez-Grande; Julia Suárez; Laura Pérez-Lago; Marta Herranz Martin; Patricia Muñoz; Pedro Sola Campoy; Pilar Catalán; Sergio Buenestado Serrano; Victor Manuel de la Cueva; Victor de la Cueva |
| EPI_ISL_7174193 | Hospital Regional Antofagasta (HRA) | Laboratorio de Genómica Microbiana, Unidad de Bioquímica, Departamento Biomédico, Facultad de Ciencias de la Salud, Universidad de Antofagasta, Chile. | Alexander Pérez; Alexandra Galetović; Benito Gómez-Silva; Camila Salazar-Ardiles; Carlos Cornejo.; Gabriel Peña; Jorge Araya; Martín Vargas; Milton Urrutia |
| EPI_ISL_2026434 | Hospital Sharp | Microbial Genomics Laboratory | Alejandra Garcia-Gasca; Bruno Gomez-Gil; Daniel Fregoso-Rueda; Julissa Enciso-Ibarra |
| EPI_ISL_2313115, EPI_ISL_2397352, EPI_ISL_2397354, EPI_ISL_2466270, EPI_ISL_2466274, EPI_ISL_7597319 | Hospital Universitari Vall d'Hebron - Vall d'Hebron Institut de Recerca | Hospital Universitari Vall d'Hebron - Vall d'Hebron Institut de Recerca | Alejandra González-Sánchez; Andrés Antón; Ariadna Rando; Carla Castillo; Cristina Andrés; Damir Garcia-Cehic; Josep Quer; Juliana Esperalba; Karen García; Maria Carmen Martin; Maria Gema Codina; Maria Piñana; Rodrigo Vásquez; Tomàs Pumarola |
| EPI_ISL_2171664 | Hospital Universitario de Guadalajara | SeqCOVID-SPAIN consortium/IBV(CSIC) | Alejandro González Praetorius y Mariela Martínez Ramírez and SeqCOVID-SPAIN consortium |
| EPI_ISL_1821208 | Hospital de Base de Bauru Hospital de Base Setima Regiao | Instituto Adolfo Lutz, Interdiciplinary Procedures Center, Strategic Laboratory | Caio Vinicius Dias Lopes; Claudia Regina Gonçalves; Claudio Tavares Sacchi; Erica Valessa Ramos Gomes; Karoline Rodrigues Campos; Leonardo Jose Tadeu de Araujo |
| EPI_ISL_1961030, EPI_ISL_8209087 | Houston Health Dept. | Houston Health Dept. | Adolfo Lara; Adolpho Lara; Pamela Brown; Ryker Penn; Yanlai Lai |
| EPI_ISL_1074330, EPI_ISL_1077898, EPI_ISL_1078682, EPI_ISL_1235839, EPI_ISL_1236499, EPI_ISL_1305695, EPI_ISL_5373948, EPI_ISL_5374035, EPI_ISL_5375352, EPI_ISL_5375453, EPI_ISL_5377910, EPI_ISL_5382700, EPI_ISL_5382729, EPI_ISL_5382906, EPI_ISL_5383133, EPI_ISL_5384691, EPI_ISL_5385277, EPI_ISL_5387434, EPI_ISL_5387451, EPI_ISL_6948795, EPI_ISL_6949033, EPI_ISL_6949101, EPI_ISL_6949167, EPI_ISL_6990116, EPI_ISL_7081316 | | | |
| see above | Houston Methodist Hospital | Houston Methodist Hospital | Ilya J. Finkelstein; James J. Davis; Jessica Cambric; Jimmy Gollihar; Kristina Reppond; Layne Pruitt; Madison N. Shyer; Marcus Nguyen; Matthew Ojeda Saavedra; Paul A. Christensen; Prasanti Yerramilli; Randall J. Olsen; Robert Olson; Ryan Gadd; S. Wesley Long; Sishir Subedi; and James M. Musser |
| EPI_ISL_6316088 | Hrvatski zavod za javno zdravstvo | Hrvatski zavod za javno zdravstvo | Anita Jurić; Dragan Jurić; Irena Tabain; Ivana Ferenčak; Josipa Kuzle |
| EPI_ISL_982211 | Hôpital Henri Mondor | Department of Virology, Henri Mondor University Hospital, Assistance Publique Hôpitaux de Paris, Université Paris-Est Créteil, INSERM U955 | Alexandre Soulier; Christophe Rodriguez; Elisabeth Trawinski; Guillaume Gricourt; Jean-Michel Pawlotsky; Melissa N'Debi; Slim Fourati; Vanessa Demontant |
| EPI_ISL_2384050 | IA State Hygienic Laboratory | Centers for Disease Control and Prevention Division of Viral Diseases, Pathogen Discovery | Alison Laufer Halpin; Ben L. Rambo-Martin; Clinton R. Paden; Dakota Howard; Darlene Wagner; Dave Wentworth; Dhwani Batra; Jasmine Padilla; Justin Lee; Katie Dillon; Krista Queen; Kristen Knipe; Kristine Lacek; Mark Burroughs; Matthew Schmerer; Mili Sheth; Peter Cook; Sam Shepard; Sarah Nobles; Shoshona Le; Suxiang Tong; Vivien Dugan; Yvette Unoarumhi |
| EPI_ISL_2189596, EPI_ISL_2189638, EPI_ISL_2189641, EPI_ISL_2189726, EPI_ISL_2189770, EPI_ISL_2189823 | ICMR-National Institute of Virology | NCDC Delhi, Biotechnology Division | A Walimbe; A.Awhale; A.Titkare; B.Apoorva; G.Divekar; H.Kengale; Hema Gogia; Hemlata Lall; K P.Shinde; K.Iyengar; K.Korabu; Kalaiarasan Ponnusamy; M L Choudhary; M.Das; Mahesh S Dhar; Manoj K Singh; Meena Datta; Partha Rakshit; Preeti Madan; Priyanka Singh; R.Verma; Radhakrishnan V. S; Robin Marwal; S Jadhav; S.Bhorekar; S.Jadhav; S.Shelkande; Sandhya Kabra; Sujeet K Singh; T.Sanjeevi; U.Saha; Uma Sharma; V Malik; V.Autude; V.Vipat; Z. Sayyed |
| EPI_ISL_1703878, EPI_ISL_1704617, EPI_ISL_1704620, EPI_ISL_1841274, EPI_ISL_1841293, EPI_ISL_1841317, EPI_ISL_1841357, EPI_ISL_1841382, EPI_ISL_1928407, EPI_ISL_1928439, EPI_ISL_1928462, EPI_ISL_1970534, EPI_ISL_2131556, EPI_ISL_2546321, EPI_ISL_2546403, EPI_ISL_2965870, EPI_ISL_4950364, EPI_ISL_6460809, EPI_ISL_8049967, EPI_ISL_8049980, EPI_ISL_8050002, EPI_ISL_8050179, EPI_ISL_8164050 | | | |
| see above | ICMR-National Institute of Virology - INSACOG | NIV Influenza | Dr. Varsha Potdar and NIC Team; NIV; Potdar; Pune; Varsha on behalf of National Influenza Centre |
| EPI_ISL_7602776 | IHU Mediterranee Infection | IHU Mediterranee Infection | Philippe Colson et al. |
| EPI_ISL_1282363 | IMD - MVZ Labor Greifswald | Robert Koch Institute | |
| EPI_ISL_2341029, EPI_ISL_2341041 | INDRE | Instituto Nacional de Medicina Genomica | Cedro-Tanda A; Cisneros-Villanueva M; Garcia-Cardenas FJ; Gisela Barrera-Badillo; Gonzalez-Barrera D; Gonzalez-Woge MA; Herrera-Montalvo LA.; Hidalgo-Miranda A; Irma Lopez-Martinez; Jose Ernesto Ramirez González; Mendoza-Vargas A; Munguia-Garza P; Orjuela-Rodríguez M; Rangel-DeLeon D; Reyes-Grajeda JP |
| EPI_ISL_7226881 | INSACOG-IGIB and Kerala State SARS-COV-2 Genome Surveillance Programme (GENESCOV2) | INSACOG and GENESCOV2 at CSIR Institute of Genomics and Integrative Biology | INSACOG-IGIB and Kerala State SARS-COV-2 Genome Surveillance Programme (GENESCOV2) |
| EPI_ISL_6885484, EPI_ISL_6885665 | INSACOG-MANIPUR | National Institute of Biomedical Genomics – INSACOG | Arindam Maitra; Kh. Ranjana Devi; L. Shivadutta Singh; Nidhan Kumar Biswas; R.K.Manojkumar Singh; Saumitra Das; Sreedhar Chinnaswamy |
| EPI_ISL_6392219, EPI_ISL_6394529, EPI_ISL_6395068 | INSACOG-Mizoram | National Institute of Biomedical Genomics – INSACOG | Arindam Maitra; Gracy Laldinmawii; N Senthil Kumar; Nidhan Kumar Biswas; Saumitra Das; Sreedhar Chinnaswamy; Swagnik Roy |
| EPI_ISL_5757253, EPI_ISL_5758002 | INSACOG-Mizoram | National Institute of Biomedical Genomics – INSACOG | Arindam Maitra; Gracy Laldinmawii; N Senthil Kumar; Nidhan Kumar Biswas; Saumitra Das; Sreedhar Chinnaswamy; Swagnik Roy |
| EPI_ISL_6392921 | INSACOG-Sikkim | National Institute of Biomedical Genomics – INSACOG | Arindam Maitra; Dr. Shrijana Gurung; Kaden Zangmu Bhutia; Nidhan Kumar Biswas; Saumitra Das; Sreedhar Chinnaswamy Tshering Ongmu Bhutia |
| EPI_ISL_1589834, EPI_ISL_6886192, EPI_ISL_6886276 | INSACOG-WB | National Institute of Biomedical Genomics – INSACOG | Ajay Chakraborti; Arindam Maitra; Bhaswati Bandyopadhyay; Nidhan Kumar Biswas; Saumitra Das; Sreedhar Chinnaswamy; Tamal Ghosh |
| EPI_ISL_7883751 | INSIDE | Instituto Butantan | Antonio Jorge Martins; Claudia Renata dos Santos Barros; David Schlesinger; Debora Botequio Moretti; Dimas Tadeu Covas; Elaine Cristina Marqueze; Elaine Vieira Santos; Evandra Strazza Rodrigues; Heidge Fukumasu; Jayme Augusto de Souza-Neto; José Salvatore Leister Patané; Luiz Alcantara; Luiz Lehmann Coutinho; Maria Carolina Elias; Maurício Lacerda Nogueira; Rafael dos Santos Bezerra; Raul Machado Neto; Rejane Maria Tommasini Grotto; Ricardo Haddad; Sandra Coccuzzo Sampaio Vessoni; Simone Kashima; Svetoslav Nanev Slavov; Vincent Louis Viala |
| EPI_ISL_2458116 | INSPI-CRN DE INFLUENZA Y OTROS VIRUS RESPIRATORIOS | NIC-INSPI | Alfredo Bruno; Domenica de Mora.; Jimmy Garcés; Johanna Laines; Lizbeth Patiño; Manuel Gonzalez; Maritza Olmedo; Michelle Páez |
| EPI_ISL_2405162 | INTERLAB | Omics Sciences Laboratory | Darlyn Amaya; Derly Andrade Molina; Gabriel Morey León; Juan Carlos Fernández Cadena; Rubén Armas González |
| EPI_ISL_7017947 | Idaho Bureau of Laboratories | Center for Global Health, University of New Mexico Health Sciences Center | Aimee Ceniseros; Christian Loera; Christopher Ball; Darrell Dinwiddie; Daryl Domman; Emalee Fischer; Jesse Young; Kurt Schwalm; Matthew Burns; R. Beukelman; Robert L. Voermans; Valerie Morley |
| EPI_ISL_4435445, EPI_ISL_5850409 | Idaho Bureau of Laboratories | Idaho Bureau of Laboratories | "R. Beukelman; Aimee Ceniseros; Christian Loera; Christopher Ball; Christopher Ball"; Matthew Charles Burns; R. Beukelman; Robert L. Voermans |
| EPI_ISL_2403017 | Illinois Department of Public Health - Springfield Lab | Illinois Department of Public Health - Springfield Lab | Bryan Sim; Gordon McCall |
| EPI_ISL_4847185 | Imágenes Diagnósticas de Mazatlán | Microbial Genomics Laboratory | ; Alejandra García-Gasca; Alejandra Hernández-Terán; Alejandro Sánchez-Flores; Alfredo Herrera-Estrella; Alicia Ocaña-Mondragón; Andreu Comas-García; Angel Gustavo Salas-Lais; Antonio Loza Román; Bernardo Martínez-Miguel; Blanca Taboada; Brenda Irasema Maldonado-Meza; Bruno Gómez-Gil; Carla Ivón Herrera-Najera; Carlos F. Arias; Celia Boukadida; Clara Esperanza Santacruz-Tinoco; Concepción Grajales-Muñiz; Consorcio Mexicano de Vigilancia Genómica (CoViGen-Mex). Authors (in alphabetical order): Julio Elias Alvarado-Yaah; Cristóbal Cháidez-Quiróz; Célida Duque Molina; Célida Martínez- Rodríguez; Daniel Fregoso-Rueda; Daniel Lira Morales; Eduardo Becerril-Vargas; Fernando Fontove-Herrera; Fidencio Mejía-Nepomuceno; Francisco Pulido; Gloria Elena Espinosa-Ayala; Gloria María Molina-Salinas; Gloria Vazquez; Hector Esteban Paz-Juárez; Hector Montoya-Fuentes; Helen Haydee Fernanda Ramírez-Plascencia; Irvin González-López; Jean Pierre González; Jesús Hernández; Joel Armando Vázquez-Pérez.; Jorge Salas-Hernández; José Antonio Enciso-Moreno; José Arturo Martínez-Orozco; José Esteban Muñoz-Medina; José de Jesús Nuñez-Contreras; Juan Bautista Chale-Dzul; Julissa Enciso-Ibarra; Luis Alberto Ochoa-Carrera; Margarita Matías-Florentino; Mario Mújica-Sánchez; Marissa Perez-Garcia; María Guadalupe Santiago-Mauricio; María Guadalupe de Jesús Mireles-Rivera; Nelly Sélem-Mojica; Pavel Isa; Ricardo Ciria Merce; Ricardo Grande; Rosa María Gutiérrez Rios; Santiago Ávila-Ríos; Selene Zárate; Susana Lopez; Verónica Mata-Haro; Victor Eduardo García-Arias; Víctor Hugo Borja-Aburto |
| EPI_ISL_2230966 | InDRE | Instituto Nacional de Medicina Genomica | Cedro-Tanda A; Garcia-Cardenas FJ; Gisela Barrera-Badillo; Gonzalez-Barrera D; Gonzalez-Woge MA; Herrera-Montalvo LA.; Hidalgo-Miranda A; Irma Lopez-Martinez; Jose Ernesto Ramirez González; Mendoza- |

| | | | |
|---|---|---|---|
| | | | Vargas A; Munguia-Garza P; Rangel-DeLeon D; Reyes-Grajeda JP |
| EPI_ISL_1587585, EPI_ISL_2323648, EPI_ISL_2375714, EPI_ISL_5100182, EPI_ISL_6401623, EPI_ISL_6659079, EPI_ISL_7444339, EPI_ISL_7700992, EPI_ISL_8032788 | | | |
| see above | Infinity Biologix | Centers for Disease Control and Prevention Division of Viral Diseases, Pathogen Discovery | Adrian Paskey; Benjamin Rambo-Martin; Chirayu Goswami; Christian Bixby; Christopher Gulvick; Clinton Paden; Clinton R. Paden; Dakota Howard; Darlene Wagner; Dhwani Batra; Duncan MacCannell; Erisa Sula; Jason Caravas; Jonathan Schultz; Kara Moser; Kristine Lacek; Matthew Schmerer; Peter Cook; Peter W. Cook; Robin Grimwood; Russ Hager; Scott Sammons; Shatavia Morrison; Tymeckia Kendall; Victoria Caban Figueroa; Yihe Wang; Yvette Unoarumhi |
| EPI_ISL_1913033 | Institut National d'Hygiène | Laboratoire de Biotechnologie | Abdelomunim Essabbar; Fatima El Falaki; Hicham Oumzil; Lahcen Belyamani and Azeddine Ibrahimi; Mohamed Rhajaoui; Mouna Ouadghiri; Saaïd Amzazi; Tarik Aanniz |
| EPI_ISL_1817985, EPI_ISL_2277721 | Institute for Infectious Diseases | Institute for Infectious Diseases | Alban Ramette; Christian Baumann; Cora Sägesser; Franziska Suter-Riniker; Miguel A Terrazos Miani; Pascal Bittel; Peter Keller; Stefan Neuenschwander; Stephen L Leib |
| EPI_ISL_2342533, EPI_ISL_2342534 | Institute for Infectious Diseases, University of Bern, Switzerland | Institute for Infectious Diseases, University of Bern, Switzerland | Alban Ramette; Christian Baumann; Cora Sägesser; Franziska Suter-Riniker; Miguel A Terrazos Miani; Pascal Bittel; Peter Keller; Stefan Neuenschwander; Stephen L Leib |
| EPI_ISL_2399456 | Institute of Epidemiology, Disease Control and Research (IEDCR) | IEDCR-ideSHi-icddr,b | Firdausi Qadri; Hassan Afrad; Manjur Hossain Khan; Sadia Rahman; Tahmina Shirin |
| EPI_ISL_2341665, EPI_ISL_2341701, EPI_ISL_2341941 | Institute of Life Sciences, Bhubaneswar | Institute of Life Sciences - INSACOG | Ajay Parida; Amol M. Kanampalliwar; Arup Ghosh; Atimukta Jha; INSACOG Consortium; Omprakash Shiriwas; Punit Prasad; Rajeeb Swain; Rupesh Dash; Safal Walia; Sana Fatma; Shifu Aggarwal; Sunil K. Raghav |
| EPI_ISL_2522055, EPI_ISL_5212538, EPI_ISL_5212684, EPI_ISL_6248953 | Institute of Microbiology and Immunology, Faculty of Medicine, University of Ljubljana | Institute of Microbiology and Immunology, Faculty of Medicine, University of Ljubljana | Alen Suljič; Andraž Celar; Cene Skubic; Domen Lazar; Dominika Šturm; Doroteja Vlaj; Mario Poljak; Matic Brvar; Miša Korva; Patricija Pozvek; Samo Zakotnik; Tatjana Avšič – Županc; Tina Gabrovšek; Tina Živič; Tomaž Mark Zorec; Špela Pleh |
| EPI_ISL_1896680, EPI_ISL_2100436, EPI_ISL_2361476 | Institute of Microbiology, Universidad San Francisco de Quito | Institute of Microbiology, Universidad San Francisco de Quito | Belén Prado-Vivar; Bernardo Darquea; Bernardo Gutiérrez; Fernada Zurita; Gabriel Trueba; Hermelinda Paguay; Jorge Montaño; Juan José Guadalupe; Luis Morales; Mayra Beltrán; Michelle Grunauer; Monica Becerra-Wong; Patricio Rojas-Silva; Paúl Cárdenas; Sully Márquez; Verónica Barragán |
| EPI_ISL_8057202 | Institute of Virology, Biomedical Research Center of the Slovak Academy of Sciences, Bratislava | Faculty of Mathematics, Physics and Informatics, Comenius University, Bratislava | Boris Klempa; Brona Brejova; Jozef Nosek; Juraj Kopacek; Kristina Borsova; Lubomira Lukacikova; Martina Lickova; Martina Nebohacova; Monika Slavikova; Sabina Fumacova Havlikova; Tomas Vinar; Veronika Vanova; Viktoria Cabanova |
| EPI_ISL_6133033, EPI_ISL_6133142 | Institute of Virology, Medical Center, University of Freiburg, Freiburg, Germany | Institute of Virology, Clinial Virus Genomics, Medical Center, University of Freiburg, Freiburg, Germany | Hajo Grundmann; Jonas Fuchs; Lena Jaki; Lisa Kern; Marcus Panning; Sandra Reuter |
| EPI_ISL_1017697 | Institute of Virology, Vaccines and Sera ''Torlak'' | Institute of microbiology and Immunology, Faculty of Medicine, University of Belgrade | Jankovic, M.; Jovanovic, T.; Knezevic, A.; Milicevic, O.; Sekler, M.; Tesovic, B.; Vidanovic, D. |
| EPI_ISL_2348759, EPI_ISL_2348775, EPI_ISL_2348778 | Instituto Nacional De Investigación En Salud Pública-Crn De Influenza Y Otros Virus Respiratorios | NIC-Instituto Nacional de Investigación en Salud Pública | Alfredo Bruno; Domenica de Mora.; Jimmy Garcés; Johanna Laines; Lizbeth Patiño; Manuel Gonzalez; Maritza Olmedo; Michelle Páez |
| EPI_ISL_1824465 | Instituto Nacional de Medicina Genomica | Centro de Investigación en Enfermedades Infecciosas (CIENI), Instituto Nacional de Enfermedades Respiratorias (INER) | Arriaga-Canon C; Avila-Rios S; Boukadida C; Cedro-Tanda A; Herrera-Montalvo LA; Hidalgo-Miranda A; Matias-Florentino M; Mendoza-Vargas A; Perez-Garcia M; Reyes-Grajeda JP; Reyes-Teran G |
| EPI_ISL_2276104, EPI_ISL_2276105, EPI_ISL_2443082 | Instituto Nacional de Medicina Genomica | Instituto Nacional de Medicina Genomica | Cedro-Tanda A; Cisneros- Villanueva M; Cisneros-Villanueva M; Escobar-Arrazola; Gonzalez-Barrera D; Herrera-Montalvo LA.; Hidalgo-Miranda A; M; Mendoza-Vargas A; Munguia-Garza P; Orjuela-Rodríguez M; Ramirez-Vega O; Rangel-DeLeon D; Reyes-Grajeda JP |
| EPI_ISL_887434 | Instituto Nacional de Saude (INS), Mozambique | KRISP, KZN Research Innovation and Sequencing Platform | Giandhari J; Nadia Sitoe; Nalia Ismael; Nedio Mabunda; Paulo Arnaldo; Pillay S; Tegally H; Wilkinson E; de Oliveira T |
| EPI_ISL_7571148, EPI_ISL_8215666 | International Centre for Diarrhoeal Disease Research, Bangladesh (icddr,b) | International Centre for Diarrhoeal Disease Research, Bangladesh (icddr,b) | Afrad MH; Bhuiyan TR; Firdausi Qadri; Hamza O; Islam T; Khanam F |
| EPI_ISL_5823735 | Israel Central Virology laboratory | Israel Central Virology laboratory | Danit Sofer; Efrat Dahan Bucris; Efrat Glick-Saar; Ella Mendelson; Gideon Rechavi; Michal Mandelboim; Miranda Geva; Neta Zuckerman; Omer Asraf; Oran Erster; Orna Mor |
| EPI_ISL_944225, EPI_ISL_1674066, EPI_ISL_1762881, EPI_ISL_1763877, EPI_ISL_2183528 | Israel Central Virology laboratory | Israel National Consortium for SARS-CoV-2 sequencing | Assaf Rokney; Dana Bar-Ilan; David A. Zeevi; Efrat Dahan Bucris; Efrat Glick-Saar; Efrat Rorman; Ella Mendelson; Ephraim Fass; Eva Nachum; Gal Zizelski Valenci; Gideon Rechavi; Israel Nissan; Joseph Jaffe; Maya Davidovich Cohen; Michal Mandelboim; Miranda Geva; Mor Rubinstein; Neta Zuckerman; Netanel Abu; Omer Murik; Omri Nayshool; Oran Erster; Orna Mor; Tzvia Mann |
| EPI_ISL_8090589 | Istituto Dermopatico dell'Immacolata IDI-IRCCS (lab Genetica)-Rome | University Campus Bio-Medico of Rome (UCBM) | Angeletti S.; De Florio L.; Fogolari M.; Francesconi M.; Lintas C.; Riva E.; Veralli R. |
| EPI_ISL_2455768 | Istituto Zooprofilattico Sperimentale del Mezzogiorno | TIGEM | Antonio Grimaldi Patrizia Annunziata Francesco Panariello Biancamaria Pierri Claudia Tiberio Teresa Giuliano Valentina Bouche Chiara Colantuono Maria Concetta Cuomo Denise Di Concilio Lucio Di Filippo Anna Manfredi Marcello Salvi Antonio Limone Luigi Atripaldi Pellegrino Cerino Andrea Ballabio Davide Cacchiarelli |
| EPI_ISL_2465164 | Istituto Zooprofilattico Sperimentale del Mezzogiorno | Telethon Institute of Genetics and Medicine (TIGEM) | Antonio Grimaldi Patrizia Annunziata Francesco Panariello Biancamaria Pierri Claudia Tiberio Teresa Giuliano Valentina Bouche Chiara Colantuono Maria Concetta Cuomo Denise Di Concilio Lucio Di Filippo Anna Manfredi Marcello Salvi Antonio Limone Luigi Atripaldi Pellegrino Cerino Andrea Ballabio Davide Cacchiarelli |
| EPI_ISL_2195855 | J.W. Ruby Memorial Hospital | WVU and Marshall University Combined Genomics Core Facilities | James Denvir; Peter Perrotta; Peter Stoilov; Ryan Percifield; Wesley Kimble |
| EPI_ISL_1939939, EPI_ISL_2272824 | JIPMER Puducherry | inStem NCBS - INSACOG | Uma Ramakrishnan Dasaradhi Palakodeti Aswin SaiNarain |
| EPI_ISL_1305922 | Johns Hopkins Hospital Department of Pathology | Johns Hopkins Hospital Department of Pathology | Adannaya Amadi; C. Paul Morris; Chun Huai Luo; Heba H. Mostafa; Matthew Schwartz |
| EPI_ISL_1939969 | KAMAKSHIPALYA | inStem NCBS - INSACOG | Uma Ramakrishnan Dasaradhi Palakodeti Aswin SaiNarain |
| EPI_ISL_7802583 | KSL Diagnostics | University at Buffalo Genomics and Bioinformatics Core | Alyssa Pohlman; Amanda Boccolucci; Brandon Marzullo; Donald Yergeau; Jennifer Surtees; Jonathan Bard; Natalie Lamb; Norma Nowak |
| EPI_ISL_1382559, EPI_ISL_7592436, EPI_ISL_7603588 | KU Leuven, Rega Institute, Clinical and Epidemiological Virology | KU Leuven, Rega Institute, Clinical and Epidemiological Virology | Bert Vanmechelen; Joan Marti-Carerras; Piet Maes; Tony Wawina-Bokalanga |
| EPI_ISL_7622027 | Kaiser Permanente NW Reginal Lab | OHSU MM Lab | Amber Halse; Jeannine Lama; Xuan Qin; Yun Wu |
| EPI_ISL_8038015 | Kaiser Permanente Southern California | Helix | Helix; Kaiser Permanente Southern California |
| EPI_ISL_2227411, EPI_ISL_2248747, EPI_ISL_2271715, EPI_ISL_4600347, EPI_ISL_8318061 | Kansas Health and Environmental Lab | Kansas Health and Environmental Lab | Amanda Bradley; Ben Olsen; Carrie Welch; Gary Buruss; Jonathan Barnell; Katherine Wiggins; Meg Wise; Mike Grose; Victor Anderson; and Phil Adam |
| EPI_ISL_2227447 | Kansas Health and Environmental Lab | Wichita State University - Molecular Diagnostics Lab | Ben Olsen; Jonathan Barnell; Katherine Wiggins; Mike Grose; and Phil Adam |
| EPI_ISL_5330813 | Kantonsspital Winterthur | Institute of Medical Virology | Alexandra Trkola; Annette Audigé; Catharine Aquino; Cyril Shah; Daniel Ehrsam; Gabriela Ziltener; Guido Bloemberg; Hubert Rehrauer; Isabel Stürmer; Joel Wirz; Jon Huder; Jürg Böni; Kevin Steiner; Maria Grünberg; Maryam Zaheri; Michael Huber; Riccarda Capaul; Stefan Schmutz; Verena Kufner; Weihong Qi |
| EPI_ISL_7196112 | Karolinska University Hospital Huddinge | Karolinska University Hospital | Annelie Bjerkner; Isak Sylvin; Jan Albert; Karolina Ininbergs; Lina Guerra Blomqvist; Lynda Eneh; Martin Ekman; Martina Wahlund; Robert Dyrdak; Sandra Broddesson; Tanja Normark; Tobias Allander; Valtteri Wirta; Zhibing Yun |
| EPI_ISL_6036682, EPI_ISL_7120693 | Karolinska University Hospital Solna | Karolinska University Hospital | Annelie Bjerkner; Isak Sylvin; Jan Albert; Karolina Ininbergs; Lina Guerra Blomqvist; Lynda Eneh; Martin Ekman; Martina Wahlund; Robert Dyrdak; Sandra Broddesson; Tanja Normark; Tobias Allander; Valtteri Wirta; Zhibing Yun |
| EPI_ISL_2433577, EPI_ISL_2433587 | Kasemrad international hospital | Division of Genomic Medicine and Innovation support,Department of Medical Sciences, Ministry of Public Health, Thailand | Archawin Rojanawiwat; Jirapha Pakdee; Natthakul Bunneang; Nuanjun Wichukchinda; Penpitcha Thawong; Pilailuk Akkapaiboon Okada; Pundharika Piboonsiri; Surakameth Mahasirimongkol; Waritta Sawaengdee |
| EPI_ISL_6684756, EPI_ISL_6684904, EPI_ISL_6684971, EPI_ISL_6685029, EPI_ISL_6690922, EPI_ISL_6691256, EPI_ISL_7986388 | | | |
| see above | Kasturba Hospital Molecular Lab - INSACOG | Kasturba hospital WGS | Jayanthi Shastri; Vidushi Chitalia |
| EPI_ISL_5332789, EPI_ISL_6370286 | Kentucky State Public Health Lab | Kentucky State Public Health Lab | Anjayya Tummala; Chitralipi Kattunga; Joshua Tobias; Karim George; Matthew Johnson; Rachel Zinner; Rhonda Lucas; Vaneet Arora; William Grooms |
| EPI_ISL_745122 | Khayelitsha Hospital wc KHA | National Health Laboratory Service (NHLS), Tygerberg | Bronwyn Kleinhans; Eduan Wilkindon; Gert van Zyl; Houriiyah Tegally; Kayla Delaney; Susan Engelbrecht; Tulio de Oliveira; Wolfgang Preiser |
| EPI_ISL_1239240, EPI_ISL_1239249, EPI_ISL_1239258, EPI_ISL_1239489 | Kimberley Hospital, National Health Laboratory Services, Northern Cape, South Africa | National Institute for Communicable Diseases of the National Health Laboratory Service | Amoako DG; Bhiman JN; Ismail A; Mahlangu B; Mohale T; Ntuli N; Scheepers C |
| EPI_ISL_5250101, EPI_ISL_5923243 | Klinika za infektivne bolesti "Dr. Fran Mihaljević" | Hrvatski zavod za javno zdravstvo | Anita Jurić; Dragan Jurić; Irena Tabain; Ivana Ferenčak; Josipa Kuzle |
| EPI_ISL_2471983 | Klinikum Ernst von Bergmann gemeinnützige GmbH - stationärer Bereich | Robert Koch Institute | |
| EPI_ISL_8228500 | Klinikum Stuttgart | Robert Koch Institute | |
| EPI_ISL_6023173 | Klinički bolnički centar Zagreb | Hrvatski zavod za javno zdravstvo | Anita Jurić; Dragan Jurić; Irena Tabain; Ivana Ferenčak; Josipa Kuzle |

| | | | |
|---|---|---|---|
| EPI_ISL_696467 | Knysna CDC wc WLC & NHLS/UCT | KRISP, KZN Research Innovation and Sequencing Platform | Arash Iranzadeh; Bruna Galvao; Carolyn Williamson; Deelan Doolabh; Diana Hardie; Emanuel James San; Houriiyah Tegally; Innocent Mudau; Jennifer Giandhari; Kruger Marais; Lynn Tyers; Marvin Hsiao; Stephen Korsman; Sureshnee Pillay; Tulio de Oliveira |
| EPI_ISL_700419, EPI_ISL_700510 | Knysna Hospital wc KNY | NHLS/UCT | Arash Iranzadeh; Bruna Galvao; Carolyn Williamson; Deelan Doolabh; Diana Hardie; Houriiyah Tegally; Innocent Mudau; Kruger Marais; Lynn Tyers; Marvin Hsiao; Stephen Korsman |
| EPI_ISL_745164 | Kraaifontein CHC wc KFP | National Health Laboratory Service (NHLS), Tygerberg | Bronwyn Kleinhans; Eduan Wilkindon; Gert van Zyl; Houriiyah Tegally; Kayla Delaney; Susan Engelbrecht; Tulio de Oliveira; Wolfgang Preiser |
| EPI_ISL_2140673 | Kwanokuthula CDC wc KWA | NHLS/UCT | Arash Iranzadeh; Bruna Galvao; Carolyn Williamson; Deelan Doolabh; Diana Hardie; Innocent Mudau; Kruger Marais; Lynn Tyers; Marvin Hsiao; Stephen Korsman |
| EPI_ISL_8034822 | LA Office of Public Health Laboratories | Centers for Disease Control and Prevention Division of Viral Diseases, Pathogen Discovery | Alex Burgin; Ben Rambo-Martin; Clinton Paden; Dakota Howard; Dave Wentworth; Dhwani Batra; Jasmine Padilla; Justin Lee; Krista Queen; Kristen Knipe; Kristine Lacek; Mark Burroughs; Matthew Schmerer; Meghan Bentz; Mili Sheth; Peter Cook; Sam Shepard; Sarah Nobles; Suxiang Tong; Vivien Dugan; Yvette Unoarumhi |
| EPI_ISL_1941773 | LA SOURCE | Laboratory of genomics and metagenomics | Claire Bertelli; Damien Jacot; Gilbert Greub; Sebastien Aeby; Trestan Pillonel |
| EPI_ISL_7910270 | LABM MOULIN-HUART-BY | CERBA HealthCare | Bénédicte Roquebert; Laura Verdurme; Sabine Trombert; Stéphanie Haim-Boukobza |
| EPI_ISL_4606332 | LABORATOIRE CERBALLIANCE PLT VILLON | CNR Virus des Infections Respiratoires - France SUD | Antonin Bal; Bruno Lina; Gregory Destras; Gwendolyne Burfin; Hadrien Regue; Laurence Josset; Martine Valette; Quentin Semanas |
| EPI_ISL_6289030 | LABORATOIRE LABAZUR | CNR Virus des Infections Respiratoires - France SUD | Antonin Bal; Bruno Lina; Bruno Simon; Gregory Destras; Gwendolyne Burfin; Hadrien Regue; Laurence Josset; Martine Valette; Quentin Semanas |
| EPI_ISL_1821062, EPI_ISL_1821063, EPI_ISL_1821064, EPI_ISL_1821065 | LABORATORIO CLINICO SYNLAB | Instituto Nacional de Salud- Dirección de Investigación en Salud Pública | Carlos Franco-Muñoz; Carmen Osorio; Diana Malo; Diego A. Álvarez-Díaz; Diego Andrés Prada; Gerardo Santamaría; Hector Alejandro Ruiz-Moreno; Jhonnatan Reales-González; Jorge Rivera; Juan Camilo Martinez; Julian Naizaque; Katherine Laiton-Donato; Lisseth Pardo; Magdalena Wiesner; Marcela Mercado-Reyes; Maria T. Herrera-Sepúlveda; Marta Lopez Blanco; Martha Lucia Ospina Martinez; Paola Rojas; Sergio Gomez; Sheryll Corchuelo; Ángela Alarcon Cruz |
| EPI_ISL_1746637 | LABORATORIUM BADAŃ KLINICZNYCH WSSE w OPOLU | 1. National Institute of Public Health - National Institute of Hygiene; 2. Eurofins Genomics Europe Sequencing GmbH | ECDC COVID-19 WGS support team; Eurofins Genomics Europe Sequencing Team; Gierczyńki Rafał; Sadkowska-Todys Małgorzata; Wołkowicz Tomasz; Zacharczuk Katarzyna |
| EPI_ISL_918499, EPI_ISL_918500, EPI_ISL_918501, EPI_ISL_918506 | LACEN - Laboratório Central de Saúde Pública do Amazonas | Evandro Chagas Institute | A.M.; Barbagelata; E.C.; E.M.A.; Ferreira; J.A.; Junior; K.C.; L.C.; L.S.; M.C.; P.S.; Pinheiro; Santos; Silva; Sousa; Sousa Junior; W.D.C.; da Silva |
| EPI_ISL_940616 | LACEN-PI DR. Costa Alvarenga | Instituto Adolfo Lutz, Interdiciplinary Procedures Center, Strategic Laboratory | Claudia Regina Gonçalves; Claudio Tavares Sacchi; Erica Valessa Ramos Gomes; Karoline Rodrigues Campos |
| EPI_ISL_8221894, EPI_ISL_8226561 | LADR Zentrallabor DR. Kramer & Kollegen Geesthacht | Robert Koch Institute | |
| EPI_ISL_1322409 | LAM SELARL 2A-2B | CNR Virus des Infections Respiratoires - France SUD | Antonin Bal; Bruno Lina; Bruno Simon; Gregory Destras; Gwendolyne Burfin; Hadrien Regue; Laurence Josset; Martine Valette; Quentin Semanas |
| EPI_ISL_6647957, EPI_ISL_6698792, EPI_ISL_6704864, EPI_ISL_6704870, EPI_ISL_6704872, EPI_ISL_6704874, EPI_ISL_6704875 | | | |
| see above | LANCET LABORATORY | National Institute for Communicable Diseases of the National Health Laboratory Service | Amoako DG; Bhiman JN; Everatt J; Glass A; Ismail A; Mahlangu B; Mnguni A; Mohale T; Ntuli N; Scheepers C; Viana R; Wolter N |
| EPI_ISL_8296477 | LBM CERBALIANCE OCCITANIE | CNR Virus des Infections Respiratoires - France SUD | Antonin Bal; Bruno Lina; Bruno Simon; Gregory Destras; Gwendolyne Burfin; Hadrien Regue; Laurence Josset; Martine Valette; Quentin Semanas |
| EPI_ISL_2289230 | LBM ST BENOIT | CNR Virus des Infections Respiratoires - France SUD | Antonin Bal; Bruno Lina; Bruno Simon; Gregory Destras; Gwendolyne Burfin; Hadrien Regue; Laurence Josset; Martine Valette; Quentin Semanas |
| EPI_ISL_6606076 | LBM UNIBIO VALENTIN | CNR Virus des Infections Respiratoires - France SUD | Antonin Bal; Bruno Lina; Bruno Simon; Gregory Destras; Gwendolyne Burfin; Hadrien Regue; Laurence Josset; Martine Valette; Quentin Semanas |
| EPI_ISL_1820927, EPI_ISL_1820929 | LDSP BARRANQUILLA | Instituto Nacional de Salud- Dirección de Investigación en Salud Pública | Carlos Franco-Muñoz; Carmen Osorio; Christian Romero; Diana Malo; Diego A. Álvarez-Díaz; Diego Andrés Prada; Gerardo Santamaría; Hector Alejandro Ruiz-Moreno; Jhonnatan Reales-González; Jorge Rivera; Juan Camilo Martinez; Julian Naizaque; Katherine Laiton-Donato; Lisseth Pardo; Magdalena Wiesner; Marcela Mercado-Reyes; Maria T. Herrera-Sepúlveda; Marta Lopez Blanco; Martha Lucia Ospina Martinez; Paola Rojas; Patricia del Portillo; Sergio Gomez; Sheryll Corchuelo; Ángela Alarcon Cruz |
| EPI_ISL_2362576 | LDSP CESAR | Instituto Nacional de Salud- Dirección de Investigación en Salud Pública | Carlos Franco-Muñoz; Carmen Osorio; Diana Malo; Diego A. Álvarez-Díaz; Diego Andrés Prada; Gerardo Santamaría; Hector Alejandro Ruiz-Moreno; Jhonnatan Reales-González; Jorge Rivera; Juan Camilo Martinez; Julian Naizaque; Katherine Laiton-Donato; Lisseth Pardo; Magdalena Wiesner; Marcela Mercado-Reyes; Maria T. Herrera-Sepúlveda; Marta Lopez Blanco; Martha Lucia Ospina Martinez; Paola Rojas; Sergio Gomez; Sheryll Corchuelo; Ángela Alarcon Cruz |
| EPI_ISL_1821069 | LDSP FRONTERIZO | Instituto Nacional de Salud- Dirección de Investigación en Salud Pública | Carlos Franco-Muñoz; Carmen Osorio; Diana Malo; Diego A. Álvarez-Díaz; Diego Andrés Prada; Gerardo Santamaría; Hector Alejandro Ruiz-Moreno; Jhonnatan Reales-González; Jorge Rivera; Juan Camilo Martinez; Julian Naizaque; Katherine Laiton-Donato; Lisseth Pardo; Magdalena Wiesner; Marcela Mercado-Reyes; Maria T. Herrera-Sepúlveda; Marta Lopez Blanco; Martha Lucia Ospina Martinez; Paola Rojas; Sergio Gomez; Sheryll Corchuelo; Ángela Alarcon Cruz |
| EPI_ISL_1821076 | LDSP SANTANDER | Instituto Nacional de Salud- Dirección de Investigación en Salud Pública | Carlos Franco-Muñoz; Carmen Osorio; Diana Malo; Diego A. Álvarez-Díaz; Diego Andrés Prada; Gerardo Santamaría; Hector Alejandro Ruiz-Moreno; Jhonnatan Reales-González; Jorge Rivera; Juan Camilo Martinez; Julian Naizaque; Katherine Laiton-Donato; Lisseth Pardo; Magdalena Wiesner; Marcela Mercado-Reyes; Maria T. Herrera-Sepúlveda; Marta Lopez Blanco; Martha Lucia Ospina Martinez; Paola Rojas; Sergio Gomez; Sheryll Corchuelo; Ángela Alarcon Cruz |
| EPI_ISL_1582987 | LDSP VICHADA | Instituto Nacional de Salud- Dirección de Investigación en Salud Pública | Carlos Franco-Muñoz; Carmen Osorio; Diana Malo; Diego A. Álvarez-Díaz; Diego Andrés Prada; Gerardo Santamaría; Hector Alejandro Ruiz-Moreno; Jhonnatan Reales-González; Juan Camilo Martinez; Julian Naizaque; Katherine Laiton-Donato; Lisseth Pardo; Magdalena Wiesner; Marcela Mercado-Reyes; Maria T. Herrera-Sepúlveda; Marta Lopez Blanco; Martha Lucia Ospina Martinez; Paola Rojas; Sergio Gomez; Sheryll Corchuelo; Ángela Alarcon Cruz |
| EPI_ISL_7740798 | LEBOWAKGOMO LABORATORY | National Institute for Communicable Diseases of the National Health Laboratory Service | Amoako DG; Bhiman JN; Everatt J; Ismail A; Mahlangu B; Mnguni A; Mohale T; Ntuli N; Scheepers C; Wolter N |
| EPI_ISL_1912939 | LESP Nuevo Leon | Instituto de Diagnostico y Referencia Epidemiologicos (INDRE) | Abril Rodriguez-Maldonado; Ariadna Medina-Benitez; Claudia Wong-Arambula; Ernesto Ramirez-Gonzalez.; Gisela Barrera-Badillo; Irma Lopez-Martinez; Joaquin Quiroz-Mercado; Lucia Hernandez-Rivas; Natividad Cruz-Ortiz; Sergio Rangel-Guerrero; Tatiana Nunez-Garcia; Vanessa Rivero-Arredondo |
| EPI_ISL_1857253 | LESP Yucatan | Instituto de Diagnostico y Referencia Epidemiologicos (INDRE) | Abril Rodriguez-Maldonado; Ariadna Medina-Benitez; Claudia Wong-Arambula; Ernesto Ramirez-Gonzalez.; Gisela Barrera-Badillo; Irma Lopez-Martinez; Joaquin Quiroz-Mercado; Lucia Hernandez-Rivas; Natividad Cruz-Ortiz; Sergio Rangel-Guerrero; Tatiana Nunez-Garcia; Vanessa Rivero-Arredondo |
| EPI_ISL_2443458 | LHUB-ULB | Labo Klinische Biologie, UZA | Basil Britto Xavier; Christine Lammens; Herman Goossens; Jasmine Coppens; Marie Le Mercier; Veerle Matheeussen |
| EPI_ISL_2422366 | LKO | Jessa | Berden et al. on behalf of the Jessa_cmdLab |
| EPI_ISL_1696700 | LSUHS Emerging Viral Threat Laboratory | LSUHS Emerging Viral Threat Laboratory | Alexander J. Mijalis; Andrew D. Yurochko; Christopher G. Kevil; Gregory L. Ware; Jennifer L. Carroll; Jeremy P. Kamil; John A. Vanchiere; Maarten Van Diest; Rona S. Scott |
| EPI_ISL_1969245 | Lab RSUP DR Mohammad Hoesin Palembang | National Institute of Health Research and Development | Arie Ardiansyah Nugraha; Hana Apsari Pawestri; Hartanti Dian Ikawati; Kartika Dewi Puspa; Krisna Pangesti; Nelly Puspandari; Subangkit; Triyani Soekarso; Vivi Setiawaty |
| EPI_ISL_1532957, EPI_ISL_1827004, EPI_ISL_2192420, EPI_ISL_5432710, EPI_ISL_7339435 | Lab voor klinische biologie | Lab voor klinische biologie | Bruno Verhasselt; Hannelore Hamerlinck; Marija Janevska; May-Linh Truong |
| EPI_ISL_8132050 | Lab. Microbiologia e Virologia Cotugno A.O. dei Colli | TIGEM | Antonio Grimaldi Patrizia Annunziata Francesco Panariello Claudia Tiberio Teresa Giuliano Valentina Bouche Chiara Colantuono Lucio Di Filippo Anna Manfredi Marcello Salvi Antonio Limone Luigi Atripaldi Andrea Ballabio Davide Cacchiarelli |
| EPI_ISL_6769090 | Lab. Virología y Genética Universidad Simón Bolívar | Laboratorio de Biologia Molecular, Universidad Cooperativa de Colombia, Santa Marta | Andrew S. Muñoz-Gamba; Antonio Acosta; Camila Gonzalez; Daniel B. Ramírez-Osorio; Johana Zuñiga; José A. Usme-Ciro; Paula A. Quintero-Cortés; Yesid Bello |
| EPI_ISL_2470445 | LabKom - Labor Hannover MVZ GmbH | Robert Koch Institute | |
| EPI_ISL_2029727, EPI_ISL_2293624, EPI_ISL_2535240, EPI_ISL_5945568, EPI_ISL_6885101, EPI_ISL_6886939, EPI_ISL_7789429, EPI_ISL_7921490 | | | |
| see above | Labo Analyses Med | National Reference Center for Viruses of Respiratory Infections, Institut Pasteur, Paris | Angela Brisebarre; BENEDICTE BACCOUCH; Camille Capel; Christophe Malabat; Corinne Maufrais; Etienne Simon-Lorière; Frédéric Lemoine; Jean Boyer; Jonas Amzalag; Julien Fumey; Louise Lefrançois; Marion Barbet; Maud Vanpeene; Melanie Caron; Méline Bizard; Pierre Lechat; S. COCCO; Slim El Khiari; Sylvie Behillil; Sylvie Van der Werf; Thierry Guffond; Vincent Enouf |
| EPI_ISL_2422226 | Labo Klinische Biologie, UZA | Labo Klinische Biologie, UZA | Basil Britto Xavier; Christine Lammens; Herman Goossens; Jasmine Coppens; Marie Le Mercier; Veerle Matheeussen |
| EPI_ISL_1154883, EPI_ISL_2260405 | Labor 28 MVZ GmbH | Robert Koch Institute | |
| EPI_ISL_1982616, EPI_ISL_1989656, EPI_ISL_2382623, EPI_ISL_2466775, EPI_ISL_2466824, EPI_ISL_2504036 | Labor Berlin Charité Vivantes GmbH / Institut für Virologie | Charité Universitätsmedizin Berlin, Institut für Virologie/Labor Berlin | Barbara Mühlemann; Christian Drosten; Christine Stephan; Peter Menzel; Rolf Schwarzer; Terry Jones; Victor M Corman |
| EPI_ISL_7163629 | Labor Dr. Fenner und Kollegen | Heinrich Pette Institute, Leibniz Institute for Experimental Virology | Adam Grundhoff; Alexis Robitaille; Johannes Knobloch; Martin Aepfelbacher; Nicole Fischer; Thomas Günther |
| EPI_ISL_1439856 | Labor Dr. Heidrich & Kollegen MVZ GmbH Hamburg | Robert Koch Institute | |
| EPI_ISL_4932484 | Labor Dr. Wisplinghoff - Köln | Robert Koch Institute | |
| EPI_ISL_2469859 | Labor Lübeck bzw. Laborärztliche Gemeinschaftspraxis Lübeck | Robert Koch Institute | |
| EPI_ISL_4444427, EPI_ISL_4611776, EPI_ISL_6204147 | Labor Mönchengladbach MVZ Dr. Stein + Kollegen GbR | Robert Koch Institute | |
| EPI_ISL_2111598 | Labor ZOTZ\|KLIMAS; MVZ Dusseldorf-Centrum | Robert Koch Institute | |

| | | | |
|---|---|---|---|
| EPI_ISL_2390702, EPI_ISL_8241387 | Labor ZOTZ\|KLIMAS; MVZ Düsseldorf-Centrum | Robert Koch Institute | |
| EPI_ISL_2313259 | Laborarztpraxis Osnabrück | Robert Koch Institute | |
| EPI_ISL_7735810 | Laboratoire Ana-L | Department of Virology, Henri Mondor University Hospital, Assistance Publique Hôpitaux de Paris, Université Paris-Est Créteil, INSERM U955 | Alexandre Soulier; Christophe Rodriguez; Elisabeth Trawinski; Guillaume Gricourt; Jean-Michel Pawlotsky; Melissa N'Debi; Slim Fourati; Vanessa Demontant |
| EPI_ISL_6641348 | Laboratoire Carage | Institut Pasteur de la Guyane | Anne Lavergne; Antoine Enfissi; Dominique Rousset |
| EPI_ISL_1696878, EPI_ISL_7662490 | Laboratoire de santé publique du Québec | Laboratoire de santé publique du Québec | Guillaume Bourque; Ioannis Ragoussis; Jesse Shapiro; Mark Lathrop and Judith Fafard on behalf of the CoVSeQ research group; Mark Lathrop and Michel Roger on behalf of the CoVSeQ research group (http://covseq.ca/researchgroup); Sandrine Moreira |
| EPI_ISL_6187897, EPI_ISL_8313741 | Laboratoires d'analyses medicales - Ketterthill | Laboratoire national de sante, Microbiology, Microbial Genomics Platform | Anke Wienecke-Baldacchino; Caroline Scheiber; Catherine Ragimbeau; Elodie Solarino; Fatu Djabi; Jessica Tapp; Lise Pignon; Raoul Salmon; Serge Vedy; Sibel Berger; Tamir Abdelrahman; Virginie Jover |
| EPI_ISL_2157480 | Laboratorio Central de Saude Publica do Estado de Minas Gerais (LACEN/MG) | Laboratory of Respiratory Viruses and Measles, Oswaldo Cruz Institute, FIOCRUZ | Alice Sampaio Rocha; Ana Carolina Mendonca; Andre Felipe Leal Bernardes; Anna Carolina Paixao; Elisa Cavalcante Pereira; Fernando Motta; Luciana Appolinario; Marilda Siqueira on behalf of the Fiocruz COVID-19 Genomic Surveillance Network; Paola Resende; Renata Serrano Lopes; Taina Venas |
| EPI_ISL_7733244 | Laboratorio CQRC | CQRC_QUALITY CONTROL CHEMICAL BIOLOGICAL RISK_AOOR Villa Sofia Cervello Palermo | Brunacci G.; Contino F.; Di Gaudio F. |
| EPI_ISL_2101747, EPI_ISL_2385647 | Laboratorio Central Noel Nutels | Bioinformatics Laboratory / LNCC | Alessandra P Lamarca; Alexandra L Gerber; Amilcar Tanuri; Ana Paula de C Guimaraes; Ana Tereza R Vasconcelos; Andrea Cony Cavalcanti; Caio Luiz Pereira Ribeiro; Cassia Alves; Cintia Policarpo; Claudia Maria Braga de Mello; Cristiane Gomes da Silva; Diana Mariani; Douglas Terra Machado; Flavio Dias da Silva; Gleidson da Silva de Oliveira; Leandro Magalhaes de Souza; Liliane Cavalcante; Luiz G P de Almeida; Marcio Henrique de Oliveira Garcia; Mario Sergio Ribeiro; Ronaldo da Silva F Jr; Silvia Carvalho |
| EPI_ISL_2091380 | Laboratorio Central de Epidemiología (LCE) | Centro de Investigación en Enfermedades Infecciosas (CIENI), Instituto Nacional de Enfermedades Respiratorias (INER) | Alejandro Sanchez-Flores; Alfredo Herrera-Estrella; Alicia Ocaña-Mondragón; Angel Gustavo Salas-Lais; Bernardo Martínez-Miguel; Blanca Taboada; Brenda Irasema Maldonado-Meza; Carla Ivón Herrera-Najera; Carlos F. Arias; Celia Boukadida; Clara Esperanza Santacruz-Tinoco; Concepción Grajales-Muñiz; Consorcio Mexicano de Vigilancia Genómica (CoViGen-Mex). Authors (in alphabetical order): Julio Elias Alvarado-Yaah; Célida Duque Molina; Fernando Fontove-Herrera; Francisco Pulido; Gloria Elena Espinosa-Ayala; Gloria María Molina-Salinas; Gloria Vazquez; Hector Esteban Paz-Juárez; Hector Montoya-Fuentes; Helen Haydee Fernanda Ramírez-Plascencia; José Antonio Enciso-Moreno; José Esteban Muñoz-Medina; José de Jesús Nuñez-Contreras; Juan Bautista Chale-Dzul; Luis Alberto Ochoa-Carrera; Margarita Matías-Florentino; María Guadalupe Santiago-Mauricio; María Guadalupe de Jesús Mireles-Rivera; Nelly Sélem-Mojica; Pavel Isa; Ricardo Grande; Santiago Ávila-Ríos; Víctor Eduardo Garcia-Arias; Víctor Hugo Borja-Aburto |
| EPI_ISL_1416458, EPI_ISL_1416510, EPI_ISL_2391613, EPI_ISL_2391666 | Laboratorio Central de Epidemiología (LCE) | Instituto de Biotecnología de la UNAM | ; Alejandra Garcia-Gasca; Alejandra Hernández-Terán; Alejandro Sanchez-Flores; Alfredo Herrera-Estrella; Alicia Ocaña-Mondragón; Andreu Comas-García; Angel Gustavo Salas-Lais; Antonio Loza Román; Bernardo Martínez-Miguel; Blanca Taboada; Brenda Irasema Maldonado-Meza; Bruno Gomez-Gil; Carla Ivón Herrera-Najera; Carlos F. Arias; Celia Boukadida; Clara Esperanza Santacruz-Tinoco; Concepción Grajales-Muñiz; Consorcio Mexicano de Vigilancia Genómica (CoViGen-Mex). Authors (in alphabetical order): Julio Elias Alvarado-Yaah; Cristóbal Cháidez-Quiróz; Célida Duque Molina; Célida Martínez- Rodríguez; Daniel Fregoso-Rueda; Daniel Lira Morales; Eduardo Becerril-Vargas; Fernando Fontove-Herrera; Fidencio Mejía-Nepomuceno; Francisco Pulido; Gloria Elena Espinosa-Ayala; Gloria Elena Espinoza-Ayala; Gloria María Molina-Salinas; Gloria Vazquez; Hector Esteban Paz-Juárez; Hector Montoya-Fuentes; Helen Haydee Fernanda Ramírez-Plascencia; Irvin González-López; Jean Pierre González; Joel Armando Vázquez-Pérez.; Jorge Ivan Salinal-Nevarez; Jorge Salas-Hernández; José Antonio Enciso-Moreno; José Arturo Martínez-Orozco; José Esteban Muñoz-Medina; José de Jesús Nuñez-Contreras; Juan Bautista Chale-Dzul; Julissa Enciso-Ibarra; Luis Alberto Ochoa-Carrera; Margarita Matías-Florentino; Mario Mújica-Sánchez; María Guadalupe Santiago-Mauricio; María Guadalupe de Jesús Mireles-Rivera; Nelly Sélem-Mojica; Pavel Isa; Ricardo Grande; Rosa María Gutierrez Rios; Santiago Ávila-Ríos; Selene Zárate; Susana Lopez; Víctor Eduardo Garcia-Arias; Víctor Hugo Borja-Aburto |
| EPI_ISL_2157414 | Laboratorio Central de Saude Publica do Esatado de Alagoas (LACEN/AL) | Laboratory of Respiratory Viruses and Measles, Oswaldo Cruz Institute, FIOCRUZ | Alice Sampaio Rocha; Ana Carolina Mendonca; Anderson Brandao Leite; Anna Carolina Paixao; Elisa Cavalcante Pereira; Fernando Motta; Luciana Appolinario; Marilda Siqueira on behalf of the Fiocruz COVID-19 Genomic Surveillance Network; Paola Resende; Renata Serrano Lopes; Taina Venas |
| EPI_ISL_3048824, EPI_ISL_3048826 | Laboratorio Central de Saude Publica do Estado do Rio Grande do Sul (LACEN-RS) | Laboratório de Biologia Molecular da Universidade Federal de Ciências da Saúde de Porto Alegre | Adriana Seixas; Ana B. G. Veiga; Ana Paula Muterle Varela; Fabiana Quoos Mayer; Fernando Hayashi Sant'Anna; Janira Prichula; Letícia Garay Martins; Richard Steiner Salvato; Tatiana Schäffer Gregianini |
| EPI_ISL_2271701 | Laboratorio Central, Ministerio de Salud Cordoba | Instituto de Patologia Vegetal (CIAP-INTA) on behalf of 'Proyecto Argentino Interinstitucional de genomica de SARS-CoV-2' (PAIS Consortium) | Barbas, G.; Castro, G.; Debat, HJ.; FD; Fernandez; M; M.B.; Marquez, N.; Pisano; Re, V. |
| EPI_ISL_2007538 | Laboratorio Central, Ministerio de Salud Córdoba | Instituto de Patologia Vegetal (CIAP-INTA) on behalf of 'Proyecto Argentino Interinstitucional de genomica de SARS-CoV-2' (PAIS Consortium) | A; Amadio; Barbas, G.; Castro, G.; Debat, HJ.; FD; Fernández; Irazoqui; M; M.B.; Marquez, N.; Pisano; Re, V. |
| EPI_ISL_8217327 | Laboratorio Clinico Imbanaco | "Laboratorio de biotecnología, Universidad Icesi" | "María I. Gutiérrez López; Adrián Camilo Rodríguez Ararat; Diana M. Florez Giraldo; Marcela Mercado; Maria F. Villegas Torres; Paola A. Caicedo Burbano"; Programa Nacional de Caracterización Genómica de SARS-CoV-2 |
| EPI_ISL_4646654 | Laboratorio Clinico Toledo | RCMI-Center for Research Resources, Ponce Research Institute | Andrea Arias-García; Raphael Sánchez-Torres; Vanessa Rivera-Amill |
| EPI_ISL_7174204 | Laboratorio Clínico, Sección Biología Molecular. Hospital Dr. Carlos Cisternas, Calama. Seremía Salud Antofagasta. | Laboratorio de Genómica Microbiana, Unidad de Bioquímica, Departamento Biomédico, Facultad de Ciencias de la Salud, Universidad de Antofagasta, Chile. | Alexander Pérez; Alexandra Galetović; Benito Gómez-Silva; Camila Salazar-Ardiles; Carlos Cornejo.; Gabriel Peña; Jorge Araya; Martín Vargas; Milton Urrutia |
| EPI_ISL_6823635 | Laboratorio Clínico, Sección Biología Molecular. Hospital Dr. Carlos Cisternas, Calama. Seremía Salud Antofagasta. | Laboratorio de Genómica Microbiana, Unidad de Bioquímica, Departamento Biomédico, Facultad de Ciencias de la Salud, Universidad de Antofagasta, Chile. Avenida Angamos 601, Antofagasta, Chile. | Alexander Pérez; Alexandra Galetović; Benito Gómez-Silva; Camila Salazar-Ardiles; Carlos Cornejo.; Gabriel Peña; Jorge Araya; Martín Vargas; Milton Urrutia |
| EPI_ISL_7731368 | Laboratorio Estatal de Salud Publica de Nuevo Leon | Laboratorio Estatal de Salud Publica de Nuevo Leon | (in alphabetical order) Consuelo Treviño-Garza; Eduardo Isaac de la Rosa-Moreno; Else del Carmen Garcia-Garcia; Gloria Alejandra Jasso-de la Peña; Manuel Enrique de la O-Cavazos; Yadira Hilerio Garcia |
| EPI_ISL_7740535 | Laboratorio Nacional de Salud | Asociación de Salud Integral / Clínica Familiar "Luis Ángel García" | Ana S. Gonzalez-Reiche; Claudia Rangel; Danicela Mercado; Eduardo Arathon; Hilda Ruiz; Luis Aguirre; Luis Rivas; Narda Medina; Oscar Bonilla; Osmar Gamboa. |
| EPI_ISL_7938424 | Laboratorio de Biología Molecular Universidad Cooperativa de Colombia Sede Santa Marta | Laboratorio de Biologia Molecular, Universidad Cooperativa de Colombia, Santa Marta | Andres Rojas-Gulloso; Andrew S. Muñoz-Gamba; Beatriz Maestre; Camila Gonzalez; Daniel B. Ramírez-Osorio; Edwin Cortes; Emy S. Torres; Johanna Zúñiga; José A. Usme-Ciro; Paula A. Quintero-Cortés |
| EPI_ISL_1582988, EPI_ISL_1582989, EPI_ISL_1582990 | Laboratorio de Biología Molecular – Universidad del Magdalena | Instituto Nacional de Salud- Dirección de Investigación en Salud Pública | Carlos Franco-Muñoz; Carmen Osorio; Diana Malo; Diego A. Álvarez-Díaz; Diego Andrés Prada; Gerardo Santamaría; Hector Alejandro Ruiz-Moreno; Jhonnatan Reales-González; Juan Camilo Martinez; Julian Naizaque; Katherine Laiton-Donato; Lisseth Pardo; Magdalena Wiesner; Marcela Mercado-Reyes; Maria T. Herrera-Sepúlveda; Marta Lopez Blanco; Martha Lucia Ospina Martinez; Paola Rojas; Sergio Gomez; Sheryll Corchuelo; Ángela Alarcon Cruz |
| EPI_ISL_833140, EPI_ISL_1068149, EPI_ISL_1068150, EPI_ISL_1068151, EPI_ISL_1068154, EPI_ISL_1068160, EPI_ISL_1068169, EPI_ISL_1068198, EPI_ISL_1068226, EPI_ISL_1068243, EPI_ISL_1068248, EPI_ISL_1261122, EPI_ISL_2777374 | | | |
| see above | Laboratorio de Ecologia de Doencas Transmissiveis na Amazonia, Instituto Leonidas e Maria Deane - Fiocruz Amazonia | Laboratorio de Ecologia de Doencas Transmissiveis na Amazonia, Instituto Leonidas e Maria Deane - Fiocruz Amazonia | André Corado; Debora Duarte; Felipe Naveca; Felipe Naveca on behalf of the Fiocruz COVID-19 Genomic Surveillance Network; Fernanda Nascimento; George Silva; Karina Pessoa; Luciana Gonçalves; Maria Júlia Brandão; Matilde Mejía; Michele Jesus; Valdinete Nascimento; Victor Souza; Ágatha Costa |
| EPI_ISL_1629792, EPI_ISL_1629795, EPI_ISL_1629799, EPI_ISL_1629800, EPI_ISL_1629801, EPI_ISL_1629802, EPI_ISL_1629804, EPI_ISL_1629805, EPI_ISL_1629806, EPI_ISL_1629807 | | | |
| see above | Laboratorio de Genómica Microbiana, Universidad Peruana Cayetano Heredia | Laboratorio de Genómica Microbiana, Universidad Peruana Cayetano Heredia | Alejandra Dávila-Barclay; Diego Cuicapuza; Guillermo Salvatierra; Janet Huancachoque; Lenin Maturrano; Luis González; Pablo Tsukayama; Pedro E. Romero; Pool Marcos |
| EPI_ISL_5994968 | Laboratorio de Infectologia y Virologia Molecular | Laboratory of Molecular Virology, School of Medicine, Pontificia Universidad Catolica de Chile | Ana Maria Contreras; Andres E. Munoz-Marcos; Carlos Palma; Catalina Pardo-Roa; Constanza Maldonado; Eileen Serrano; Erick Salinas; Estefany Poblete; Eugenia L. Fuentes Luppichini; Francisco Melo; Jennifer Angulo; Jorge Levican; Leonardo I. Almonacid; M. Belen Leyton; Marcela Ferres; Maria Jose Avendano; Rafael A. Medina.; Tamara Garcia-Salum |
| EPI_ISL_1111127, EPI_ISL_1111130, EPI_ISL_1111143, EPI_ISL_1111151, EPI_ISL_1111291, EPI_ISL_1111295, EPI_ISL_1111296, EPI_ISL_1111302, EPI_ISL_1111318, EPI_ISL_1111331 | | | |
| see above | Laboratorio de Referencia Nacional de Virus Respiratorio. Instituto Nacional de Salud Perú | Laboratorio de Referencia Nacional de Enteropatógenos. Instituto Nacional de Salud del Perú | Fiorella Orellana Peralta; Iris Silva Molina; Junior Caro Castro; Ronnie Gavilan Chavez; Veronica Hurtado Vela; Willi Quino Sifuentes |
| EPI_ISL_1629787 | Laboratorio de Referencia Nacional de Virus Respiratorios, Instituto Nacional de Salud Peru | Laboratorio de Genómica Microbiana, Universidad Peruana Cayetano Heredia | Alejandra Dávila-Barclay; Diego Cuicapuza; Guillermo Salvatierra; Janet Huancachoque; Lenin Maturrano; Luis González; Maribel Huaringa; Nancy Rojas; Pablo Tsukayama; Pedro E. Romero; Pool Marcos; Priscilla Lope |
| EPI_ISL_5201374 | Laboratorio de Referencia Nacional de Virus Respiratorios. Centro Nacional de Salud Publica. Instituto Nacional de Salud Peru. | Laboratorio de Referencia Nacional de Virus Respiratorios. Centro Nacional de Salud Publica. Instituto Nacional de Salud Peru. | Carlos Padilla Rojas; Henri Bailon Calderon; Iris Silva Molina; Joseph Huayra Niquen; Lely Solari Zerpa; Luis Barcena Flores; Marco Galarza Perez; Nancy Rojas Serrano; Nieves Sevilla Castañeda; Omar Caceres Rey; Orson Mestanza Millones; Princesa Medrano Alhuay; Priscila Lope Pari; Sandra Morales Ruiz; Sara Gordillo Vilchez; Steve Acedo Lazo; Veronica Hurtado Vela; Victor Jimenez Vasquez; Wendy Lizarraga Olivares |
| EPI_ISL_2536789 | Laboratorio de Referencial Nacional de Virus Respiratorios | Laboratorio de Referencial Nacional de Virus Respiratorios | Carlos Padilla Rojas; Henri Bailon Calderon; Iris Silva Molina; Joseph Huayra Niquen; Lely Solari Zerpa; Luis Barcena Flores; Marco Galarza Perez; Nancy Rojas Serrano; Omar Caceres Rey; Orson Mestanza Millones; Priscila Lope Pari; Sandra Morales Ruiz; Steve Acedo Lazo; Veronica Hurtado Vela |
| EPI_ISL_2007496 | Laboratorio de Virología del Hospital de Niños Dr. Ricardo Gutierrez | Área de Secuenciación del Laboratorio de Virología del Hospital de Niños Dr. Ricardo Gutierrez on behalf of 'Proyecto Argentino Interinstitucional de genomica de SARS-CoV-2' (PAIS Consortium) | A; Acevedo; Acuña; Alexay; Alvarez Lopez; Barreda Frank; C; D; E; G; Goya; Grandis; Jacques; LE; Labarta; Lusso; M; ME; MI; Medina; Mistchenko; N; Nabaes Jodar; Natale; O; S; Streitenberger; Thomas; Valinotto; Viegas, M.; Villegas |
| EPI_ISL_8124586 | Laboratorio di Microbiologia | Laboratorio di Microbiologia, Istituto di Medicina di Laboratorio EOLAB, Ente Ospedaliero Cantonale | Martinetti Lucchini Gladys; Valeria Spina |
| EPI_ISL_1622927 | Laboratorium Medyczne LAB-MED | 1. National Institute of Public Health - National Institute of Hygiene; 2. Eurofins Genomics Europe Sequencing GmbH | ECDC COVID-19 WGS support team; Eurofins Genomics Europe Sequencing Team; Gierczyńki Rafał; Sadkowska-Todys Małgorzata; Wołkowicz Tomasz; Zacharczuk Katarzyna |
| EPI_ISL_1339300, EPI_ISL_1515082, EPI_ISL_1515705, EPI_ISL_1681962, EPI_ISL_1684014, EPI_ISL_1684031, EPI_ISL_1798592, EPI_ISL_1799131, EPI_ISL_1928696, EPI_ISL_1929653, EPI_ISL_2184345, EPI_ISL_2184498, EPI_ISL_2184778, EPI_ISL_2185180, EPI_ISL_2185347, EPI_ISL_2305093, EPI_ISL_2305230, EPI_ISL_2306096, EPI_ISL_2306260, EPI_ISL_2306821, EPI_ISL_2306832, EPI_ISL_2306883, EPI_ISL_2398442, EPI_ISL_2398958, EPI_ISL_2480323, EPI_ISL_2480423, EPI_ISL_2480552, EPI_ISL_2480559, EPI_ISL_2480975, EPI_ISL_2482173, EPI_ISL_2482277, EPI_ISL_4334218, EPI_ISL_4336979, EPI_ISL_4337158, EPI_ISL_4338594, EPI_ISL_4483421, EPI_ISL_4485585, EPI_ISL_4811306, EPI_ISL_4813515, EPI_ISL_4818750, EPI_ISL_5079312, EPI_ISL_5086180, EPI_ISL_5086593, EPI_ISL_5087148, EPI_ISL_5090980, EPI_ISL_5174888, EPI_ISL_5175297, EPI_ISL_5178702, EPI_ISL_5183932, EPI_ISL_5238639, EPI_ISL_5264310, EPI_ISL_5561955, EPI_ISL_5565353, EPI_ISL_5571981, EPI_ISL_5574169, EPI_ISL_5576266, EPI_ISL_5576271, EPI_ISL_5869980, EPI_ISL_5870225, EPI_ISL_5870458, EPI_ISL_5870856, EPI_ISL_5870938, EPI_ISL_5873824, EPI_ISL_5875402, EPI_ISL_6077767, EPI_ISL_6077835, EPI_ISL_6077987, EPI_ISL_6078396, EPI_ISL_6087170, EPI_ISL_6088798, EPI_ISL_6088891, EPI_ISL_6088982, EPI_ISL_6089223, EPI_ISL_6089308, EPI_ISL_6388972, EPI_ISL_6391037, EPI_ISL_6391233, EPI_ISL_6391430, EPI_ISL_6530305, EPI_ISL_6531721, EPI_ISL_6532115, EPI_ISL_6532607, EPI_ISL_6535692, EPI_ISL_6540272, EPI_ISL_6543189, EPI_ISL_6788791, EPI_ISL_6791183, EPI_ISL_6793749, EPI_ISL_6906146, EPI_ISL_7246167, EPI_ISL_7247137, EPI_ISL_7249286, EPI_ISL_7253996, EPI_ISL_7663310, EPI_ISL_7667664, EPI_ISL_7669903, EPI_ISL_7670280, EPI_ISL_7670784, EPI_ISL_7772731, EPI_ISL_7773168, EPI_ISL_7911133, EPI_ISL_8023903, EPI_ISL_8024435, EPI_ISL_8028927, EPI_ISL_8029359, EPI_ISL_8029475, EPI_ISL_8154565, EPI_ISL_8154588, EPI_ISL_8155847, EPI_ISL_8156754, EPI_ISL_8263992 | | | |

| | | | |
|---|---|---|---|
| see above | Laboratory Corporation of America | Centers for Disease Control and Prevention Division of Viral Diseases, Pathogen Discovery | Adrian Paskey; Amanda Douglas; Amanda Suchanek; Andrea Throop; Ayla Burns; Ben L. Rambo-Martin; Benjamin Rambo-Martin; Bobbi Croy; Brian Krueger; Brian Norvell; Christopher Gulvick; Christos Petropoulos; Clinton Paden; Clinton R. Paden; Craig Lukasik; Dakota Howard; Darlene Wagner; Debbie Boles; Dhwani Batra; Duncan MacCannell; Eyad Almasri; Goran Stevovic; Howard Engler; Hrushikesh Deshmukh; Jake Humphrey; Jana Schroth; Jason Caravas; Joe Voshell; John Pruitt; Jonathan Meltzer; Jonathan Williams; Kara Moser; Kimberly Wagner; Kristine Lacek; Lax Iyer; Lisa Pfefferle; Lyndon Tilson; Manoj Jain; Marcia Eisenberg; Mary Ann Cristobal; Mary Cristobal; Mary Williamson; Matthew Robinson; Matthew Schmerer; Michael Levandoski; Mike Sapeta; Mindy Nye; Minoo Agarwal; Mohan Kolli; Nuthawin Charoensri; Oren Cohen; Peter Cook; Peter W. Cook; Prashant Gupta; Qian Zeng; Rama Ghatti; Scott Parker; Scott Ryan; Scott Sammons; Shatavia Morrison; Stanley Letovsky; Steven Ragan; Suresh Babu Selvaraju; Suresh Selvaraju; Susan Countryman; Susan Hicks; Suxiang Tong; Suzanne Dale; Thomas Urban; Tim Kuphal; Tricia Zwiefelhofer; Tymeckia Kendall; Victoria Caban Figueroa; Vincent Drouillon; Yvette Unoarumhi |
| EPI_ISL_1021210, EPI_ISL_1160917, EPI_ISL_1163438 | Laboratory Corporation of America | Respiratory Viruses Branch, Division of Viral Diseases, Centers for Disease Control and Prevention | Amanda Douglas; Amanda Suchanek; Andrea Throop; Ayla Burns; Ben L. Rambo-Martin; Bobbi Croy; Brian Krueger; Brian Norvell; Christos Petropoulos; Clinton R. Paden; Craig Lukasik; Dakota Howard; Dhwani Batra; Duncan MacCannell; Eyad Almasri Debbie Boles; Goran Stevovic; Howard Engler; Hrushikesh Deshmukh; Jake Humphrey; Jana Schroth; Joe Voshell; John Pruitt; Jonathan Meltzer; Jonathan Williams; Kimberly Wagner; Lax Iyer; Lyndon Tilson; Manoj Jain; Marcia Eisenberg; Mary Ann Cristobal; Mary Williamson; Michael Levandoski; Mike Sapeta; Mindy Nye; Minoo Agarwal; Mohan Kolli; Nuthawin Charoensri; Oren Cohen; Peter W. Cook; Prashant Gupta; Qian Zeng; Rama Ghatti; Scott Parker; Scott Ryan; Stanley Letovsky; Steven Ragan; Suresh Babu Selvaraju; Susan Countryman; Susan Hicks; Suxiang Tong; Suzanne Dale; Thomas Urban; Tim Kuphal; Tricia Zwiefelhofer; Vincent Drouillon |
| EPI_ISL_7908575 | Laboratory for Molecular Diagnostics,IPHMN | Eurofins Genomics Europe Sequencing GmbH | Danijela Vujošević; Marija Govedarica |
| EPI_ISL_2249440, EPI_ISL_2249444 | Laboratório Central de Saúde Pública do Rio de Janeiro | Coordenação Geral de Laboratórios de Saúde Pública (CGLAB/DAEVS/SVS/MS) | Vagner Fonseca; et al. |
| EPI_ISL_1630158 | Laboratório de Microbiologia Molecular - Universidade FEEVALE | Laboratório de Microbiologia Molecular - Universidade FEEVALE | Alana Witt Hansen; Fernando Rosado Spilki; Flávio Silveira; Fágner Henrique Heldt; Juliana Schons Gularte; Juliane Deise Fleck; Mariana Soares da Silva; Matheus Nunes Weber; Meriane Demoliner; Micheli Filippi.; Paula Rodrigues de Almeida |
| EPI_ISL_1464650 | Laboratório de Virologia - UNIFESP | Laboratory of Respiratory Viruses and Measles, Oswaldo Cruz Institute, FIOCRUZ | Alice Sampaio Rocha; Ana Carolina Mendonca; Anna Carolina Paixao; Fernando Motta; Luciana Appolinario; Marilda Siqueira on behalf of the Fiocruz COVID-19 Genomic Surveillance Network; Nancy Belei; Paola Resende; Renata Serrano Lopes |
| EPI_ISL_7883651 | Lancet | NHLS/UCT | Arash Iranzadeh; Bruna Galvao; Carolyn Williamson; Deelan Doolabh; Diana Hardie; Gert Marais; Innocent Mudau; Luicer Olubayo; Lynn Tyers; Marvin Hsiao; Nokuzola Mbhele; Rageema Joseph; Stephen Korsman |
| EPI_ISL_6913991, EPI_ISL_6913992, EPI_ISL_6913993, EPI_ISL_6913994, EPI_ISL_6913995, EPI_ISL_6913996, EPI_ISL_6913997, EPI_ISL_6913998, EPI_ISL_6913999, EPI_ISL_6914003, EPI_ISL_6914004, EPI_ISL_6914007 | | | |
| see above | Lancet Laboratories | National Institute for Communicable Diseases of the National Health Laboratory Service | Amoako DG; Bhiman JN; Everatt J; Ismail A; Mahlangu B; Mnguni A; Mohale T; Ntuli N; Scheepers C; Wolter N |
| EPI_ISL_1350204 | Landesgesundheitsamt Baden-Württemberg | Robert Koch Institute | |
| EPI_ISL_2471861, EPI_ISL_2471887 | Landesuntersuchungsanstalt für das Gesundheits- und Veterinärwesen (LUA) Sachsen | Robert Koch Institute | |
| EPI_ISL_8254490 | Leids Universitair Medisch Centrum | Leiden University Medical Center | Stefan Boers |
| EPI_ISL_2521713 | Libramont | Plateforme de testing Namuroise | Degosserie Jonathan; Demars Aurore; Denis Olivier; Lesly Nyinkeu Kemamen; Maschietto Céline; Mullier François; Nobis Chloé; Otto Gaetan |
| EPI_ISL_8219939 | Limbach - MVZ Humangenetik Ulm | Robert Koch Institute | |
| EPI_ISL_2471514 | Limbach - MVZ Labor Ravensburg Labor Dr. Gärtner | Robert Koch Institute | |
| EPI_ISL_7808370 | Lurie Children's Hospital | RIPHL at Rush University Medical Center | Alyse Kittner; Christine Inguglia; David Williams; Edith Perez; Felix Araujo Perez; Hannah Barbian; Jason Rippe; Kevin Kunstman; Larry Kociolek; Laura Furtado; Marieta Hyde; Sofiya Bobrovska; Stefan Green |
| EPI_ISL_2249631, EPI_ISL_2443817, EPI_ISL_2443841 | M Health Fairview | Minnesota Department of Health, Public Health Laboratory | Alexandra Lorentz; Jacob Garfin; Matt Plumb; and Xiong Wang |
| EPI_ISL_1575150, EPI_ISL_2291826, EPI_ISL_2322718, EPI_ISL_2322723, EPI_ISL_2425409 | MD PHL | MD PHL | Maryland Department of Health Laboratories Administration |
| EPI_ISL_1155516, EPI_ISL_1353161 | MDI Limbach Berlin GmbH | Robert Koch Institute | |
| EPI_ISL_4437757, EPI_ISL_7074563 | MDI Limbach Berlin GmbH; MVZ Labor Berlin | Robert Koch Institute | |
| EPI_ISL_2272599 | MEPHI, Aix Marseille University | MEPHI, Aix Marseille University | Anthony LEVASSEUR |
| EPI_ISL_1663543, EPI_ISL_1663548, EPI_ISL_1663549 | MGM Medical College, Jamshedpur | Institute of Life Sciences - INSACOG | Ajay Parida; Amol M. Kanampalliwar; Arup Ghosh; Atimukta Jha; INSACOG Consortium; Punit Prasad; Rajeeb Swain; Rupesh Dash; Safal Walia; Shifu Aggarwal; Sunil K. Raghav |
| EPI_ISL_2422326 | MONTEFIORE MEDICAL CENTER LABORATORIES | Wadsworth Center, New York State Department of Health | Alexis Russell; Catharine Prussing; Daryl M. Lamson; Erasmus Schneider; Erica Lasek-Nesselquist; John Kelly; Jonathan Plitnick; Kirsten St. George; Matthew Shudt; Melissa A Leisner; Navjot Singh |
| EPI_ISL_1709905, EPI_ISL_6494029, EPI_ISL_6494066, EPI_ISL_7908077, EPI_ISL_7908123 | MSHS Clinical Microbiology Laboratories | MSHS Pathogen Surveillance Program | Adolfo García-Sastre; Adriana van de Guchte; Ajay Obla; Alberto Paniz-Mondolfi; Ana S. Gonzalez-Reiche; Angela Amoako; Aria Rooker; Ashley Salimbangon; Betsaida Salom Melo; Bremy Alburquerque; Brianne Ciferri; Charles Gleason; Christian Cognigni; Daniel Floda; Deena R. Altman; Denise Jurczyszak; Emilia Mia Sordillo; Gintaras Deikus; Giulio Kleiner; Gopi Patel; Hala Alshammary; Harm van Bakel; Irina Oussenko; Jayeeta Dutta; Jose Polanco; Juan Soto; Julia Matthews; Katherine Beach; Kathryn Twyman; Kayla Russo; Keith Farrugia; Komal Srivastava; Levy Sominsky; Mahmoud Awawda; Marta Luksza; Matthew M. Hernandez; Melissa Gitman; Michael D. Nowak; Mitchell J. Sullivan; Nancy Francoeur; Robert Sebra; Sarah Schaefer; Shelcie Fabre; Shwetha Hara Sridhar; Viviana Simon; Ying-Chih Wang; Zain Khalil; Zenab Khan |
| EPI_ISL_2471718, EPI_ISL_5118119, EPI_ISL_5118143 | MVZ Dr. Eberhard & Partner Dortmund | Robert Koch Institute | |
| EPI_ISL_8224350, EPI_ISL_8237821 | MVZ Labor Dr. Fenner und Kollegen (Standort Hamburg) | Robert Koch Institute | |
| EPI_ISL_2472488, EPI_ISL_7097556 | MVZ Labor Dr. Limbach & Kollegen GbR | Robert Koch Institute | |
| EPI_ISL_2266056 | MVZ Labor Krone GbR | Robert Koch Institute | |
| EPI_ISL_4444191 | MVZ Medizinisches Labor Hannover GmbH | Robert Koch Institute | |
| EPI_ISL_2471481, EPI_ISL_4621233, EPI_ISL_7750982 | MVZ für Laboratoriumsmedizin und Mikrobiologie Koblenz-Mittelrhein (Labor Koblenz) | Robert Koch Institute | |
| EPI_ISL_2342063, EPI_ISL_2342293 | Mahatma Gandhi Memorial Medical College & Hospital Jamshedpur | Institute of Life Sciences - INSACOG | Ajay Parida; Amol M. Kanampalliwar; Arup Ghosh; Atimukta Jha; INSACOG Consortium; Omprakash Shiriwas; Punit Prasad; Rajeeb Swain; Rupesh Dash; Safal Walia; Sana Fatma; Shifu Aggarwal; Sunil K. Raghav |
| EPI_ISL_1936208, EPI_ISL_1936226, EPI_ISL_1936291 | Main Chemical Laboratories Egypt Army | Main Chemical Laboratories Egypt Army | Abdullah Salama; AbedElrahman Zekri; Ahmed Gad; Mohamed Abdel-Monem; Mohamed El-Esawi; Mohamed Seadawy; Mohamed Shamel; Mostfa Elhoseiny |
| EPI_ISL_8214278 | Maine Health and Environmental Testing Laboratory | Tewhey Lab, The Jackson Laboratory | Barter, M.; Dewey, H.; H. and Tewhey, R.; Iosue, F.; Lynch, R.; Matluk, N.; Munger |
| EPI_ISL_5083931, EPI_ISL_6652527, EPI_ISL_7790934, EPI_ISL_8078673, EPI_ISL_8078769, EPI_ISL_8122810, EPI_ISL_8125878, EPI_ISL_8129805, EPI_ISL_8263012 | | | |
| see above | Mako Medical | Centers for Disease Control and Prevention Division of Viral Diseases, Pathogen Discovery | Benjamin Rambo-Martin; Christopher Gulvick; Clinton Paden; Dakota Howard; Dhwani Batra; Duncan MacCannell; Erisa Sula; Jason Caravas; Kristine Lacek; Lauren Moon; Matthew Schmerer; Matthew Tugwell; Peter Cook; Scott Sammons; Shatavia Morrison; Tymeckia Kendall; Victoria Caban Figueroa; Yvette Unoarumhi |
| EPI_ISL_8164246 | Malawi Liverpool Wellcome Trust Clinical Research Program | Malawi Liverpool Wellcome Trust Clinical Research Program | Belson Kutambe; Ben Morton; Catherine Anscombe; Kondwani Jambo; Mavis Menyere; Philip Ashton; Sam Lissauer |
| EPI_ISL_6931510 | Mangalam Invitro Laboratory, Rajkot | Gujarat Biotechnology Research Centre | Apurvasinh Puvar; Bhadreshsinh Gohil; Chaitanya Joshi; Dinesh Kumar; Janvi Raval; Madhvi Joshi; Nimesh Patel; Nitin Savaliya; Nitin Shukla; Priyank Chavda; Rahul Thumbar; Ramesh Pandit; Sonal Sharma; Zarna Patel |
| EPI_ISL_2157021, EPI_ISL_2157022, EPI_ISL_2157026 | Mapmygenome | CSIR-Centre for Cellular and Molecular Biology-INSACOG | Amareshwar Vodapalli; Ara Sreenivas; Archana Bharadwaj Siva; B Himasri; Blessy B John; Divya Tej Sowpati; Karthik Bharadwaj Tallapaka; Lamuk Zaveri; Payel Mukherjee; Rakesh K Mishra; Sharath Chandra Thota; Shreekant Verma; Sofia Banu; Tulasi Nagabandi; Valli Nagalakshmi Undamatla; Viswagithe S L |
| EPI_ISL_2161520, EPI_ISL_5640364, EPI_ISL_6041031 | Maryland Genomics, Institute for Genome Sciences, University of Maryland School of Medicine | Maryland Genomics, Institute for Genome Sciences, University of Maryland School of Medicine | Aditya; Claire M; Fraser; Hazen; Holly; Humphrys; Ivette; Jacques; Jonathan; Kranthi; Lim; Lisa D; Luke J; Mehta; Mike; Ott; Ravel; Roussey; Sadzewicz; Sandra; Santana-Cruz; Tallon; Tracy; Vavikolanu |
| EPI_ISL_6668353 | Mass General Brigham | Mass General Brigham | A.E.; Adams, G.; Anahtar, M.; B.L.; B.W.; Bauer, M.; Birren; Branda, J.; Carter, A.; Cerrato, F.; Chaluvadi, S.; Chapman; Cusick, C.; D.J.; DeRuff, K.; E. and Sabeti; Flowers, K.; Gallagher, G.; Gladden-Young, A.; Gnirke, A.; Harris, J.; J.E.; K.J.; LaRocque, R.; Lagerborg, K.; Lemieux; Lin; Loreth, C.; MacInnis; Neumann, A.; Normandin, E.; P.C.; Park; Pierce, V.; Reilly, S.; Rosenberg; Rudy, M.; Ryan, E.; S.B.; Shaw, B.; Siddle; Slater, D.; Smole, S.; Tomkins-Tinch, C.; Turbett, S. |
| EPI_ISL_1710630, EPI_ISL_2229483 | Massachusetts State Public Health Laboratory | Massachusetts State Public Health Laboratory | Andrew Lang; Glen Gallagher; Sandra Smole; Timelia Fink |
| EPI_ISL_1372192 | Mayo Clinic & Mayo Clinic Laboratories | Minnesota Department of Health, Public Health Laboratory | Alexandra Lorentz; Jacob Garfin; Matt Plumb; and Xiong Wang |
| EPI_ISL_1196004 | Mbabane PHU | National Institute for Communicable Diseases of the National | Amoako DG; Bhiman JN; Ismail A; Mahlangu B; Maphalala GP; Mohale T; Ntuli N; Scheepers C |

| | | | |
|---|---|---|---|
| | | Health Laboratory Service | |
| EPI_ISL_8229971, EPI_ISL_8249289 | Medizinisch-Diagnostisches Labor Kempten allgäulab | Robert Koch Institute | |
| EPI_ISL_2389102, EPI_ISL_2470840 | Medizinische Laboratorien Düsseldorf | Robert Koch Institute | |
| EPI_ISL_2390357 | Medizinisches Labor Ostsachsen MVZ GbR Nebenbetriebsstätte Dresden | Robert Koch Institute | |
| EPI_ISL_2318999 | Megalab, Molecular and Cytogenetics Diagnostics | Department for Virology, Molecular Biology and Genome Research, R. G. Lugar Center for Public Health Research, National Center for Disease Control and Public Health (NCDC) of Georgia. | Adam Kotorashvili; Amiran Gamkrelidze.; Ana Papkiauri; Ann Machablishvili; Anna Kasradze; Davit Tsaguria; Ekaterine Khmaladze; Ekaterine Zangaladze; Ekaterine Zhghenti; Giorgi Gogoladze; Giorgi Tomashvili; Gvantsa Brachveli; Gvantsa Chanturia; Irma Burjanadze; Ketevan Sidamonidze; Khatuna Zakhashvili; Lela Sabadze; Lela Urushadze; Magda Dgebuadze; Maia Alkhazashvili; Mari Gavashelidze; Mariam Zakalashvili; Marine Murtskhvaladze; Meri Pantsulaia; Nato Kotaria; Nino Berishvili; Nino Chikhovani; Paata Imnadze; Roena Sukhiashvili; Tamar Jashiashvili; Tata Imnadze; Tea Tevdoradze |
| EPI_ISL_1906153, EPI_ISL_1906559 | Michigan Department of Health and Human Services, Bureau of Laboratories | Michigan Department of Health and Human Services, Bureau of Laboratories | Blankenship HM; Riner D; Soehnlen MK |
| EPI_ISL_1892989 | Microbiologia CATLAB | Can Ruti SARS-CoV-2 Sequencing Hub (HUGTiP/IrsiCaixa/IGTP) | Alba Sánchez; Anna Not; Antoni E Bordoy; Bonaventura Clotet; Cristina Casañ; Cristina Esteban; Francesc Catala-Moll; Gemma Clara; Ignacio Blanco; Marc Noguera-Julian; Maria Casadellà; Mariona Parera; Mercedes Guerrero; Montserrat Giménez; Pere-Joan Cardona; Pilar Armengol; Roger Paredes; Verónica Saludes; and Elisa Martró on behalf of the Can Ruti SARS-CoV-2 Sequencing Hub. |
| EPI_ISL_2379289 | Microbiological Diagnostic Unit - Public Health Laboratory (MDU-PHL) | MDU-PHL | M.L.; N.L.; Sait; Seemann T.; Sherry |
| EPI_ISL_2281296, EPI_ISL_2372777, EPI_ISL_2434043, EPI_ISL_2448026 | Microbiology Department, Laboratori Clínic Metropolitana Nord. Hospital Universitari Germans Trias i Pujol | Can Ruti SARS-CoV-2 Sequencing Hub (HUGTiP/IrsiCaixa/IGTP) | Alba Snchez; Alba Sánchez; Anna Not; Antoni E Bordoy; Bonaventura Clotet; Cristina Casa; Cristina Casañ; Cristina Esteban; Francesc Catala-Moll; Gemma Clara; Ignacio Blanco; Marc Noguera-Julian; Maria Casadell; Maria Casadellà; Mariona Parera; Mercedes Guerrero; Montserrat Gimnez; Montserrat Giménez; Pere-Joan Cardona; Pilar Armengol; Roger Pared; Roger Paredes; Vernica Saludes; Verónica Saludes; and Elisa Martr on behalf of the Can Ruti SARS-CoV-2 Sequencing Hub.; and Elisa Martró on behalf of the Can Ruti SARS-CoV-2 Sequencing Hub. |
| EPI_ISL_2351304 | Microbiology Department, Laboratori Clínic Metropolitana Nord. Hospital Universitari Germans Trias i Pujol. | Can Ruti SARS-CoV-2 Sequencing Hub (HUGTiP/IrsiCaixa/IGTP) | Alba Sánchez; Anna Not; Antoni E Bordoy; Bonaventura Clotet; Cristina Casañ; Cristina Esteban; Francesc Catala-Moll; Gemma Clara; Ignacio Blanco; Marc Noguera-Julian; Maria Casadellà; Mariona Parera; Mercedes Guerrero; Montserrat Giménez; Pere-Joan Cardona; Pilar Armengol; Roger Paredes; Verónica Saludes; and Elisa Martró on behalf of the Can Ruti SARS-CoV-2 Sequencing Hub. |
| EPI_ISL_848199, EPI_ISL_2510665, EPI_ISL_2510677, EPI_ISL_2510678, EPI_ISL_2510681, EPI_ISL_6530931 | Microbiology Department. Complexo Hospitalario Universitario de Vigo | Microbiology Department. Complexo Hospitalario Universitario de Vigo | Alfaya N; Alonso I; Alvarez M; Cabrera JJ; Carballo R; Complexo Hospitalario Universitario de Vigo (CHUVI). EPICOVIGAL.; Cores O; Cortizo S; Davina C; Martinez L; Mediero G; Microbiology Department; Pena I; Perez S; Potel C; Regueiro B; Rey S; Vasallo FJ; del-Campo V |
| EPI_ISL_1303259 | Microbiology Division, South Carolina Department of Health and Environmental Control | Microbiology Division, South Carolina Department of Health and Environmental Control | Flores, H.; Freeman, J. |
| EPI_ISL_873191 | Microbiology Division, South Carolina Department of Health and Environmental Control (SC DHEC) | Microbiology Division, South Carolina Department of Health and Environmental Control (SC DHEC) | Flores, H.; Freeman, J. |
| EPI_ISL_2259158, EPI_ISL_2420756, EPI_ISL_2420759, EPI_ISL_5852818, EPI_ISL_8255058, EPI_ISL_8255079 | Microvida | Microvida | Jaco J. Verweij; Jaco Verweij; Joep J. J. M. Stohr; Joep Stohr; Suzan D. Pas |
| EPI_ISL_1621293 | Middlemore Hospital | Institute of Environmental Science and Research (ESR) | Anja Werno; Antje van der Linden; Arlo Upton; Chris Mansell; David Hammer; Dragana Drinkovic; Erasmus Smit; Gary McAuliffe; Hana Sofia Andersson; Hermes Perez; James Ussher; Jill Sherwood; Jing Wang; Joep de Ligt; Josh Freeman; Julia Howard; Juliet Elvy; Lauren Jelly; Mary DeAlmeida; Matt Blakiston; Matt Storey; Matthew Rogers; Max Bloomfield; Michael Addidle; Michelle Balm; Muhammad Faisal; Nikki Freed; Olin Silander; Olivia Stroeven; Rachel Boyle; Sally Roberts; SallyAnn Harbison; Sarah Jefferies; Sharmini Muttaiyah; Susan Morpeth; Susan Taylor; Timothy Blackmore; Vani Sathyendran; Veronica Playle; Virginia Hope; Xiaoyun Ren |
| EPI_ISL_4265220, EPI_ISL_4589931, EPI_ISL_4958473, EPI_ISL_5060199, EPI_ISL_5859190, EPI_ISL_5859281, EPI_ISL_5859365, EPI_ISL_5859427, EPI_ISL_5859460, EPI_ISL_5859464, EPI_ISL_6133642, EPI_ISL_6569760, EPI_ISL_6574677, EPI_ISL_6575041, EPI_ISL_6816164, EPI_ISL_7420112, EPI_ISL_7421273, EPI_ISL_7874448, EPI_ISL_8056091, EPI_ISL_8164444, EPI_ISL_8243568, EPI_ISL_8243580, EPI_ISL_8308604, EPI_ISL_8308612, EPI_ISL_8308632, EPI_ISL_8308633, EPI_ISL_8308634, EPI_ISL_8308647, EPI_ISL_8308648 | | | |
| see above | Ministry of Health Turkey | Ministry of Health Turkey | Fatma Bayrakdar; Gulay Korukluoglu; Gülay Korukluoğlu; Suleyman Yalcin; Süleyman Yalcin; Yasemin Cosgun; Yasemin Coşgun |
| EPI_ISL_1713881, EPI_ISL_5852623, EPI_ISL_7660022 | Ministry of Public Health / Hamad Medical Corporation | Biomedical Research Center (BRC), Qatar University / Qatar Genome Project (QGP) | Asmaa A. Al-Thani. MOPH and HMC: Abdullatif Al-Khal; Asmaa A. Al-Thani. MOPH and HMC: Abdullatif Al-Khal; BRC: Fatiha M. Benslimane; BRC: Fatiha M. Benslimane; BRC: Heba A. Al-Khatib; Chadi Saad; Dana Al-Batesh; Dana Al-Batsh; Dina Elgakhlab QGP: Fatima H. Al-Kuwari; Dina Elgakhlab. QGP: Fatima H. Al-Kuwari; Einas A. E. Al-Kuwari; Fatiha M. Benslimane; Fatma Hassan Ali; Hadeel T. Mohammed; Hadi M. Yassine; Hamad E. Al-Romaihi; Hamda Alromaihi; Heba A. Al-Khatib; Maria K. Smatti; Mashael A. Al-Bader; Mohammed Al-Thani; Muna A. S. Al-Maslamani; Oal Al-Jamal; Ola Al-Jamal; Peter V. Coyle; Reham A. El-Kahlout. QBB: Tasneem Al-Hamad; Reham A. El-Kahlout. QBB: Tasneem Al-Hamad; Roberto Bertollini; Salih Al-Marri; Swapna Thomas |
| EPI_ISL_2249526, EPI_ISL_2249612, EPI_ISL_2443810, EPI_ISL_6371398 | Minnesota Department of Health, Public Health Laboratory | Minnesota Department of Health, Public Health Laboratory | Alexandra Lorentz; Alyssa Mondelli; Elizabeth Horn; Jacob Garfin; Kelly Pung; Matt Plumb; Sarah Namugenyi; and Xiong Wang |
| EPI_ISL_2140682 | Mitchells Plain Hospital wc MPH | NHLS/UCT | Arash Iranzadeh; Bruna Galvao; Carolyn Williamson; Deelan Doolabh; Diana Hardie; Innocent Mudau; Kruger Marais; Lynn Tyers; Marvin Hsiao; Stephen Korsman |
| EPI_ISL_7899848 | Molecular Biology Laboratory, Westchester Medical Center | Genomics Core Laboratory, New York Medical College | Brahmaraju Mopidevi; Humayun K. Islam; Jian Zhuge; Rania Hatab; Salomon Amar; Utsav Pandey |
| EPI_ISL_1250514, EPI_ISL_1250517, EPI_ISL_1250531 | Molecular Diagnostics Services (MDS) | KRISP, KZN Research Innovation and Sequencing Platform | Emmanuel SJ; Giandhari J; Khan S; Lessells R; Maslo C; Mdlalose K; Pillay S; Sitharam L; Tegally H; Wilkinson E; York D; de Oliveira T |
| EPI_ISL_6901255 | Molecular diagnostic laboratory of Federal Budget Institution of Science "Central Research Institute of Epidemiology" of The Federal Service on Customers' Rights Protection and Human Well-being Surveillance | Group of Genomics and Postgenomic Technologies of Central Research Institute of Epidemiology | Akimkin V.G.; Buharina A.Y.; Chudinov I.K.; Dolotova S.M.; Goncharov S.E.; Kondrasheva L.Y.; Korneenko E.V.; Nadtoka M.I.; Roev G.V.; Samoilov A.E.; Selezov S.Y.; Shipulina O.Y.; Sinitsyn S.O.; Smirnova Y.S.; Speranskaya A.S.; Svetlichnyj D.V.; Vyhodceva A.V.; Zuev S.N. |
| EPI_ISL_6112627, EPI_ISL_6112848 | Mysuru Medical College and Research Institute | INSACOG-KA , NIMHANS | Ananthapadmanabha Kotambail; Anita S Desai; Chetan G K; Ellango Ramasamy; Gautham Arunachal Udupi; Mahesh Kumar.C.S; Sony Sharma; V Ravi |
| EPI_ISL_1588107 | NB-Hôpital Georges L. Dumont | National Microbiology Laboratory (NML) | Anna Majer; Anneliese Landgraff; CanCOGeN's metadata curation team; Darian Hole; Elsie Grudeski; Gary Van Domselaar; Grace Seo; Guillaume Desnoyers; Jennifer Tanner; Kirsten Biggar; Madison Chapel; Morag Graham; Natalie Knox; Nathalie Bastien; Philip Mabon; Public Health Agency of Canada CanCOGeN team; Rhiannon Huzarewich; Richard Garceau; Russell Mandes; Shari Tyson; Timothy Booth; Yan Li |
| EPI_ISL_1663375, EPI_ISL_2341673, EPI_ISL_2341791, EPI_ISL_2342630, EPI_ISL_2342937, EPI_ISL_2342964 | NCCS, Pune | Institute of Life Sciences - INSACOG | Ajay Parida; Amol M. Kanampalliwar; Arup Ghosh; Atimukta Jha; INSACOG Consortium; Omprakash Shiriwas; Punit Prasad; Rajeeb Swain; Rupesh Dash; Safal Walia; Sana Fatma; Shifu Aggarwal; Sunil K. Raghav |
| EPI_ISL_7545660 | NHLS Livingstone Laboratory | CERI, Centre for Epidemic Response and Innvoation, Stellenbosch University and KRISP, KZN Research Innovation and Sequencing Platform, UKZN. | Arisha Maharaj; Giandhari J; Naidoo Y; Oluwakemi Laguda-Akingba and Nokukhanya Mdlalose; Pillay S; Ramphal U; Ramphal Y; San JE; Tegally H; Tshiabuila D; Wilkinson E; de Oliveira T |
| EPI_ISL_7381211 | NHLS Port Elizabeth Laboratory | CERI, Centre for Epidemic Response and Innvoation, Stellenbosch University and KRISP, KZN Research Innovation and Sequencing Platform, UKZN. | Arisha Maharaj; Giandhari J; Moir M; Naidoo Y; Oluwakemi Laguda-Akingba and Nokukhanya Mdlalose; Pillay S; Ramphal U; Ramphal Y; San JE; Tegally H; Tshiabuila D; Wilkinson E; de Oliveira T; van Wyk S |
| EPI_ISL_912471, EPI_ISL_912476, EPI_ISL_912477, EPI_ISL_912508, EPI_ISL_7714415 | NHLS Universitas Academic | UFS Virology | D Goedhals; Emmanuel Ogunbayo; MM Nyaga; MT Mogotsi; P Nthiga; PA Bester; T de Oliveira |
| EPI_ISL_660605, EPI_ISL_660606, EPI_ISL_660609, EPI_ISL_660611, EPI_ISL_678566, EPI_ISL_736931, EPI_ISL_736933, EPI_ISL_736938, EPI_ISL_736943, EPI_ISL_736947, EPI_ISL_736959, EPI_ISL_825092, EPI_ISL_825095, EPI_ISL_825096, EPI_ISL_825103, EPI_ISL_825105, EPI_ISL_825107, EPI_ISL_825120, EPI_ISL_825124, EPI_ISL_825128, EPI_ISL_825134, EPI_ISL_825454 | | | |
| see above | NHLS-IALCH | KRISP, KZN Research Innovation and Sequencing Platform | ChimukangaraB; Giandhari J; Khan S; Lessells R; Mdlalose K; Pillay S; Tegally H; Wilkinson E; York D; de Oliveira T |
| EPI_ISL_860580, EPI_ISL_860597, EPI_ISL_860619 | NHLS-IALCH | KRISP, KZn Research Innovation and Sequencing Platform | Giandhari J; Khan S; Lessells R; Mdlalose K; Pillay S; Tegally H; Wilkinson E; York D; de Oliveira T |
| EPI_ISL_8308350 | NIH | National Institute of Hygiene | Bimouhen Abderrahman; Fatima El Falaki; Hassan Ihazmade; Hicham Oumzil; Zakia Regragui |
| EPI_ISL_1939985, EPI_ISL_1939986, EPI_ISL_1939999 | NIMHANS | inStem NCBS - INSACOG | Uma Ramakrishnan Dasaradhi Palakodeti Aswin SaiNarain |
| EPI_ISL_8038378 | NIV | NCL, Pune | Ajinkya Khilari; Anu Raghunathan; Bhagyashree Litikar; Dhanasekaran Shanmugam; Divya Niveditha; Jugal Kanekar; Shikha Takur |
| EPI_ISL_6794325 | NJ Public Health and Environmental Laboratories | Centers for Disease Control and Prevention Division of Viral Diseases, Pathogen Discovery | Alex Burgin; Ben Rambo-Martin; Clinton Paden; Dakota Howard; Dave Wentworth; Dhwani Batra; Jasmine Padilla; Justin Lee; Krista Queen; Kristen Knipe; Kristine Lacek; Mark Burroughs; Matthew Schmerer; Meghan Bentz; Mili Sheth; Peter Cook; Sam Shepard; Sarah Nobles; Suxiang Tong; Vivien Dugan; Yvette Unoarumhi |
| EPI_ISL_2443095 | NJDOH, Public Health and Environmental Laboratories | NJ_PHEL | Allison Roder; Byeong Jeong; Chelsea San Filippo; Dana Woell; Lindsey Bodnar; Ryan Pachucki; Shiv K. Verma |
| EPI_ISL_1195754, | NYU Langone Health | Departments of Pathology and Medicine, New York University | Adriana Heguy; Christian Marier; Dacia Dimartino; Emily Guzman; Gael Westby; Guiqing Wang; Paolo Cotzia; Paul Zappile; Peter Meyn; Sitharam Ramaswami; Yutong Zhang |

| | | | |
|---|---|---|---|
| EPI_ISL_1790027, EPI_ISL_2202730 | | School of Medicine | |
| EPI_ISL_5738714, EPI_ISL_6104093, EPI_ISL_7326613, EPI_ISL_7326865, EPI_ISL_7328323, EPI_ISL_7803573 | National Center of Infectious and Parasitic Diseases | National Center of Infectious and Parasitic Diseases | Alexiev et al |
| EPI_ISL_2162392, EPI_ISL_2162417, EPI_ISL_2162445, EPI_ISL_2225689, EPI_ISL_2332759 | National Centre For Cell Science | National Centre For Cell Science – INSACOG | Ajay Pillai; Dhiraj Paul; INSACOG Consortium team; Manoj Kumar Bhat; Mitali Inamdar; Mohak P Gujare; Shivang P. Bhanushali; Yogesh Shouche |
| EPI_ISL_2382960, EPI_ISL_2382970, EPI_ISL_2382996 | National Centre For Cell Science | National Centre For Cell Science-INSACOG | Ajay Pillai; Dhiraj Paul; INSACOG Consortium team; Manoj Kumar Bhat; Mitali Inamdar; Mohak P Gujare; Shivang P. Bhanushali; Yogesh Shouche |
| EPI_ISL_2107025, EPI_ISL_2107066, EPI_ISL_2107071, EPI_ISL_2107075, EPI_ISL_2107083, EPI_ISL_2107088 | National Centre For Cell Science – INSACOG | National Centre For Cell Science | Ajay Pillai; Dhiraj Paul; INSACOG Consortium team; Manoj Kumar Bhat; Mitali Inamdar; Mohak P Gujare; Shivang P. Bhanushali; Yogesh Shouche |
| EPI_ISL_2233595 | National Centre for Disease Control | CDFD -INSACOG | Ashwin Dalal; Asmita Gupta; Divya Vashisht; Murali Bashyam; Pratyusha Bala; Vinay Donipadi |
| EPI_ISL_2460533, EPI_ISL_2461035, EPI_ISL_2461158, EPI_ISL_2461509, EPI_ISL_2461699, EPI_ISL_2461701, EPI_ISL_2462007 | | | |
| see above | National Centre for Disease Control (NCDC) Biotechnology Division, Delhi | NCDC Delhi, Biotechnology Division INSACOG | Hema Gogia; Hemlata Lall; Kalaiarasan Ponnusamy; Mahesh S Dhar; Manoj K Singh; Meena Datta; Partha Rakshit; Preeti Madan; Priyanka Singh; Radhakrishnan V. S; Robin Marwal; Sandhya Kabra; Sujeet K Singh; Uma Sharma |
| EPI_ISL_5327871 | National Health Laboratory Service, Eastern Cape, South Africa | CERI, Centre for Epidemic Response and Innvoation, Stellenbosch University and KRISP, KZN Research Innovation and Sequencing Platform, UKZN. | Giandhari Jennifer; Naidoo Yeshnee; Oluwakemi Laguda-Akingba; Pillay Sureshnee; San James; Tegally Houriiyah; Tshiabuila Derek; Wilkinson Eduan; Yajna Ramphal; de Oliveira Tulio |
| EPI_ISL_6795195, EPI_ISL_6795199, EPI_ISL_7015212, EPI_ISL_7545692, EPI_ISL_7545711, EPI_ISL_7545758, EPI_ISL_7747543 | | | |
| see above | National Health Laboratory Service, Kwazulu-Natal, South Africa | CERI, Centre for Epidemic Response and Innvoation, Stellenbosch University and KRISP, KZN Research Innovation and Sequencing Platform, UKZN. | Arisha Maharaj; Giandhari J; Naidoo Y; Nokukhanya Mdlalose; Pillay S; Ramphal U; Ramphal Y; San JE; Tegally H; Tshiabuila D; Wilkinson E; de Oliveira T |
| EPI_ISL_944140, EPI_ISL_944166, EPI_ISL_1250482 | National Health Laboratory Service, South Africa | KRISP, KZN Research Innovation and Sequencing Platform | Emmanuel SJ; Giandhari J; Khan S; Laguda-Akingba O; Lessells R; Maslo C; Mdlalose K; Pillay S; Sitharam L; Tegally H; Wilkinson E; York D; de Oliveira T |
| EPI_ISL_6795188, EPI_ISL_6795189, EPI_ISL_6795190, EPI_ISL_6795193, EPI_ISL_7715355, EPI_ISL_7715357 | National Health Laboratory Services, Virology | CERI, Centre for Epidemic Response and Innvoation, Stellenbosch University and KRISP, KZN Research Innovation and Sequencing Platform, UKZN. | Arisha Maharaj; Avani Bharuthram; Florette Treurnicht; Giandhari J; Kathleen Subramoney; Naidoo Y; Nkhensani Mtileni; Pillay S; Ramphal U; Ramphal Y; San JE; Tegally H; Tshiabuila D; Wilkinson E; de Oliveira T |
| EPI_ISL_6699730, EPI_ISL_6699732, EPI_ISL_6699733, EPI_ISL_6699735, EPI_ISL_6699737, EPI_ISL_6699740, EPI_ISL_6699745, EPI_ISL_6699752, EPI_ISL_6699754, EPI_ISL_6699762, EPI_ISL_6699764, EPI_ISL_6782079, EPI_ISL_6810484 | | | |
| see above | National Health Laboratory Services, Virology, Charlotte Maxeke Johannesburg hospital, Parktown, Johannesburg, Gauteng | CERI, Centre for Epidemic Response and Innvoation, Stellenbosch University and KRISP, KZN Research Innovation and Sequencing Platform, UKZN. | Amoaka D; Arisha Maharaj; Avani Bharuthram; Bester P; Bhiman J; Engelbrecht S; Everatt J; Florette Treurnicht; Goedhals D; Hardie D; Hsiao M; Iranzadeh A; Kathleen Subramoney; Lessells R; Makatini Z; Maponga T; Mdlalose N; Mlisana K; Moir M; NGS-SA (Scheepers C; Naidoo Y; Nkhensani Mtileni; Nyaga M) Giandhari J; Oluwakemi M; Pillay S; Preiser W; Ramphal U; Ramphal Y; San JE; Tegally H; Tshiabuila D; Venter M; Wilkinson E; Williamson C; de Oliveira T; von Gottberg A |
| EPI_ISL_6939035, EPI_ISL_6939041, EPI_ISL_6939067 | National Influenza Centre | National Influenza Centre | ; Benjamiin B. Lindsey; Benjamin H. Foulkes; Bless Seyram Agbenyo; Bright Adu; Ernest Asiedu; Franklin Asiedu-Bekoe; Hilda Opoku Frempong; Ivy A. Asante; Joseph Oliver-Commey; Joyce Appiah-Kubi; Keren Okyerebea Attiku; Linda Boatemaa; Lorreta Kwasah; Mathew D. Parker; Michael Marks; Mildred Adusei-Poku; Quaneeta Mohktar; Sharon Hsu; Thushan I de Silva; William K. Ampofo |
| EPI_ISL_7456451 | National Institute For Communicable Diseases Of The National Health Laboratory Service | National Institute for Communicable Diseases of the National Health Laboratory Service | Amoako DG; Bhiman JN; Everatt J; Ismail A; Mahlangu B; Mnguni A; Mohale T; Ntuli N; Scheepers C; Wolter N |
| EPI_ISL_1239851 | National Institute for Communicable Diseases,National Health Laboratory Services, Gauteng, South Africa | National Institute for Communicable Diseases of the National Health Laboratory Service | Amoako DG; Bhiman JN; Ismail A; Mahlangu B; Mohale T; Ntuli N; Scheepers C |
| EPI_ISL_1447440 | National Institute of Infectious Diseases (NIID) | National Institute of Infectious Diseases (NIID) | Hideka Miura; Kentaro Itokawa; Kiyoko Okamoto; Kumiko Araki; Makoto Kuroda; Masanori Hashino; Rina Tanaka; Seiichiro Fujisaki; Shinichiro Hirai; Tsuyoshi Sekizuka |
| EPI_ISL_2433936 | National Institute of Laboratory Medicine and Referral Center | Genomic Research Lab, BCSIR | Abu Sayeed Mohammad Mahmud; Arifa Akram; Asish Kumar Ghosh; Barna Goswami; Eshrar Osman; Iffat Jahan; Mahmuda Yeasmin; Md. Ahasan Habib; Md. Kamrul Islam; Md. Maruf Ahmed Molla; Md. Murshed Hasan Sarkar; Md. Salim Khan; Mohammad Mohi Uddin; Mohammad Samir Uzzaman; Shahina Akter; Tanjina Akhter Banu; Tasnim Nafisa |
| EPI_ISL_7261835 | National Laboratory for Health, Environment and Food, OMM, Koper | NLZOH (National Laboratory for Health, Environment and Food) / CISLD (Clinical Institute of Special Laboratory Diagnostics), University Children's Hospital, University Medical Center Ljubljana | Aleksander Kocuvan; Aleksander Mahnic; Alenka Štorman; Ana Grom; Barbara Jenko Bizjan; Gašper Strugar; Kaja Tominc; Katarina Kozmos; Maja Rupnik; Marko Pokorn; Maruša Debeljak; Maša Jarčič; Mitja Rak / Jernej Kovač; Nika Gobec; Robert Šket; Sandra Janezic; Tadej Battelino; Tina Cvetković; Tine Tesovnik; Tjaša Žohar Čretnik |
| EPI_ISL_1191867, EPI_ISL_1191876, EPI_ISL_1191918, EPI_ISL_1191929, EPI_ISL_1191961 | National Microbiology Reference Laboratory | Quadram Institute Bioscience | Agnes Juru; Ana-Victoria Gutierrez; Andrew J. Page; Andrew Tarupiwa; Charles Nyagupe; David Baker; Faustinos T Takawira; Gaetan Thilliez; Gemma Kay; Hlanai Gumbo; Justin O'Grady; Kenneth K Maeka; Leonardo de Oliveira Martins; Muchaneta Mugabe; Raiva Simbi; Robert Kingsley; Sekesai Zinyowera; Tapfumanei Mashe; Thanh Le Viet |
| EPI_ISL_2483841 | National Platform bis UMONS/Jolimont | National Platform bis UMONS/Jolimont | Florian Juszczak; François Dufrasne; Gautier Detry; Guillaume Bayon-Vicente; Ruddy Wattiez |
| EPI_ISL_7405740 | National Public Health Laboartory | National Public Health Laboratory | National Public Health Laboratory Team |
| EPI_ISL_6963506 | National Public Health Laboratory | Central Department Of Biotechnology, Tribhuvan University | Krishna Das Manandhar |
| EPI_ISL_7391092 | National Public Health Laboratory | NPHL | National Public Health Laboratory Team |
| EPI_ISL_1816949, EPI_ISL_1897619, EPI_ISL_2349863, EPI_ISL_2508963, EPI_ISL_7604616, EPI_ISL_8092733 | National Public Health Laboratory, National Centre for Infectious Diseases | National Public Health Laboratory, National Centre for Infectious Diseases | Grace Jie Yin Ngan; Katherine Ching; Lin Cui; Raymond Tzer Pin Lin; Royce Ang; Samuel Loo; Tze Minn Mak; Zhenyang Zhou |
| EPI_ISL_1273394 | National Reference Laboratory - Ministry of Health Maseru Lesotho | National Institute for Communicable Diseases of the National Health Laboratory Service | Amoako DG; Banda R; Bhiman JN; Gorova V; Ismail A; Mahlangu B; Mathabo M; Mohale T; Mooko M; Ntuli N; Scheepers C |
| EPI_ISL_789092, EPI_ISL_1092834, EPI_ISL_1117765, EPI_ISL_1252275, EPI_ISL_1398626, EPI_ISL_2029174, EPI_ISL_2088054, EPI_ISL_2362991, EPI_ISL_2399927, EPI_ISL_2533657, EPI_ISL_7406068, EPI_ISL_7406089, EPI_ISL_7776634 | | | |
| see above | National Virus Reference Laboratory | National Virus Reference Laboratory | Charlene Bennet; Charlene Bennett; Cillian F De Gascun; Dana Alalwan; Daniel Hare; Gabriel Gonzalez; Guerrino Macori; Jonathan Dean; Michael Carr; Seamus Fanning; Zoe Yandle |
| EPI_ISL_7167335 | Nebraska Medicine | UNMC COVID-19 Response Team | UNMC COVID-19 Response Team |
| EPI_ISL_678562, EPI_ISL_678563, EPI_ISL_678564, EPI_ISL_678573, EPI_ISL_678595, EPI_ISL_678596 | Netcare | KRISP, KZN Research Innovation and Sequencing Platform | ChimukangaraB; Giandhari J; Khan S; Lessells R; Mdlalose K; Pillay S; Tegally H; Wilkinson E; York D; de Oliveira T |
| EPI_ISL_8161852 | New Mexico Department of Health Scientific Laboratory | Center for Global Health, University of New Mexico Health Sciences Center | Anastacia Griego; Darrell Dinwiddie; Daryl Domman; Jesse Young; Joseph Hicks; Kurt Schwalm; Michael Edwards; Twila Kunde; Valerie Morley |
| EPI_ISL_1114694 | New Mexico Department of Health Scientific Laboratory | New Mexico Department of Health Scientific Laboratory | Anastacia Griego-Fisher; D'eldra Malone; Ellie Johnson; Jennifer Benoit |
| EPI_ISL_1715411 | Nigerian Centre for Disease Control (NCDC) | African Centre of Excellence for Genomics of Infectious Diseases, Redeemer's University | I.B. et al; Olawoye |
| EPI_ISL_2361904 | Noguchi Memorial Institute for Medical Research, University of Ghana, Legon, Ghana | Institute of Tropical Medicine, Universitätsklinikum Tübingen, Germany | Abraham Kwabena Anang; Bright Adu; Dorothy Yeboah-Manu; Hilda Opoku Frempong; John Kofi Odoom; Joseph Humphrey Kofi Bonney; Joyce Appiah-Kubi; Keren Okyerebea Attiku; Le Thi Kieu Linh; Quaneeta Mohktar; Sivaramakrishna Rachakonda; Srinivas-reddy Pallerla; Thirumalaisamy P Velavan |
| EPI_ISL_8065958, | Noguchi Memorial Institute for Medical Research, University of Ghana, Legon, Ghana | Noguchi Memorial Institute for Medical Research, University | Bright Adu; Hilda Opoku Frempong; John K. Odoom; Joseph K. H. Bonney; Joyce Appiah-Kubi; Keren O. Attiku; Mohktar; Peace Uche; Quaneeta; Seyram B. Avonyo |

| | | | |
|---|---|---|---|
| EPI_ISL_8065959 | | of Ghana, Legon, Ghana | |
| EPI_ISL_2156591 | Northern Mindanao TB Regional Center | Philippine Genome Center | Alethea R. de Guzman; Anna Ong-Lim; Arianne A. Zamora; Asia Louisa U. Chong; Benedict A. Maralit; Candice Francheska B. Tambaoan; Carlo M. Lapid; Celia Carlos; Devon Ray Pacial; Edsel Maurice Salvaña; El King D. Morado; Elcid Aaron R. Pangilinan; Eva Maria Cutiongco-de la Paz; Francis A. Tablizo; Irish Coleen A. Asin; Jaime C. Montoya; Jan Michael C. Yap; Jo-Hannah S. Llames; John Q. Wong; Joshua Gregor A. Dizon; Juan Antonio R. Magalang; Karol Sophia Agape R. Padilla; Kenneth M. Kim; Kris P. Punayan; Marc Edsel C. Ayes; Marc Jerrone R. Castro; Maria Rosario Singh-Vergeire and Cynthia P. Saloma; Maria Sofia L. Yangzon; Marissa Alejandria; Razel Nikka M. Hao; Rianna Patricia S. Cruz; Sheila Mae M. Araiza |
| EPI_ISL_8119538 | Northstar Pathology Lab, Ahmedabad | Gujarat Biotechnology Research Centre | Akhilesh Modi; Apurvasinh Puvar; Bhadreshsinh Gohil; Chaitanya Joshi; Disha Vora; Janvi Raval; Jaykumar Rangani; Jigar Suthar; Madhvi Joshi; Nimesh Patel; Nitin Savaliya; Nitin Shukla; Priyank Chavda; Ramesh Pandit; Roshani Mishra; Sonal Sharma; Tasnim Trivedi; Zarna Patel |
| EPI_ISL_7976184 | Northwestern Memorial Hospital | Northwestern University - Center for Pathogen Genomics and Microbial Evolution | Chad J. Achenbach; Chao Qi; Egon A. Ozer; Judd F. Hultquist; Lacy M. Simons; Lawrence J. Jennings; Michael G. Ison; Ramon Lorenzo-Redondo; Taylor J. Dean |
| EPI_ISL_2009345 | Northwestern Memorial Hospital | Northwestern University - Ozer Lab | Chad J. Achenbach; Chao Qi; Egon A. Ozer; Judd F. Hultquist; Lacy M. Simons; Lawrence J. Jennings; Michael G. Ison; Ramon Lorenzo-Redondo; Taylor J. Dean |
| EPI_ISL_2323331 | Norwegian Institute of Public Health, Department of Virology | Norwegian Institute of Public Health, Department of Virology | Atiya R Ali; Debech Nadia; Engebretsen Serina Beate; Garcia Llorente Ignacio; Hilde Elshaug; Hilde Vollan; Jon Bråte; Kamilla Heddeland Instefjord; Karoline Bragstad; Kathrine Stene-Johansen; Line Victoria Moen; Marie Paulsen Madsen; Olav Hungnes; Pedersen Benedikte Nevjen; Rasmus Riis Kopperud |
| EPI_ISL_7624125 | OHSU Lab Services Molecular Microbiology Lab | Oregon SARS-CoV-2 Genome Sequencing Center | Alec J. Hirsch; Andrew C. Adey; Benjamin N. Bimber; Brendan L. O'Connell; Brian J. O'Roak; Cierra LaBlanc; Daniel N. Streblow; Destine Krenik; Donna Hansel; Guang Fan; Rabeka Ali; Ruth V. Nichols; Sonia Acharya; William B. Messer; Xuan Qin |
| EPI_ISL_875872 | Ohio Department of Health Laboratory | Ohio Department of Health Laboratory | Caitlin McDonnell; Eric Brandt; Erica Leasure; Glen McGillivary; Heather Blankenship; Holmes; Jennifer; Kelsey Florek; Keoni Omura; Kirtana Ramadugu; Quanta Brown; and Tammy Bannerman |
| EPI_ISL_1968639 | Omics Sciences Laboratory | Omics Sciences Laboratory | Darlyn Amaya; Derly Andrade Molina; Gabriel Morey León; Juan Carlos Fernández Cadena; Katheryn Sacheri Viteri; Rubén Armas González |
| EPI_ISL_6938727 | Orange County Public Health Laboratory | Fulgent Genetics | Becky Tsai; Benafsh Sapra; Doreen Ng; Harry Gao; James Xie; John Gao; Joseph Fierro; Mickey Li; Yan Meng |
| EPI_ISL_6937429 | Orange County Public Health Laboratory | Orange County Public Health Laboratory | Angelica Torres; Julia Wolfe |
| EPI_ISL_4472078, EPI_ISL_7363006 | Oslo University Hospital, Department of Microbiology | Norwegian Institute of Public Health, Department of Virology | Arvind Yegambaram Meenakshi Sundaram; Cathrine Fladeby; Garcia Llorente Ignacio; Gregor D. Gilfillan; Hilde Elshaug; Hilde Vollan; Jon Bråte; Kamilla Heddeland Instefjord; Karoline Bragstad; Kathrine Stene-Johansen; Line Victoria Moen; Lise Andresen; Mariann Nilsen; Mona Holberg-Petersen; Olav Hungnes; Pedersen Benedikte Nevjen; Pål Marius Bjørnstad; Rasmus Riis Kopperud; Teodora Plamenova Ribarska |
| EPI_ISL_6696153, EPI_ISL_7466967, EPI_ISL_7777006 | Ostfold Hospital Trust - Kalnes, Centre for Laboratory Medicine, Section for gene technology and infection serology | Norwegian Institute of Public Health, Department of Virology | Atiya R Ali; Debech Nadia; Engebretsen Serina Beate; Garcia Llorente Ignacio; Hilde Elshaug; Hilde Vollan; Jon Bråte; Kamilla Heddeland Instefjord; Karoline Bragstad; Kathrine Stene-Johansen; Line Victoria Moen; Marie Paulsen Madsen; Olav Hungnes; Pedersen Benedikte Nevjen; Rasmus Riis Kopperud |
| EPI_ISL_7605742 | PORT ELIZABETH LABORATORY | National Institute for Communicable Diseases of the National Health Laboratory Service | Amoako DG; Bhiman JN; Everatt J; Ismail A; Mahlangu B; Mnguni A; Mohale T; Ntuli N; Scheepers C; Wolter N |
| EPI_ISL_8048085 | PRVKP FKUI | National Institute of Health Research and Development | Arie Ardiansyah Nugraha; Fajar Nur Sulistiyohadi; Hana Apsari Pawestri; Hartanti Dian Ikawati; Kartika Dewi Puspa; Krisna Pangesti; Nelly Puspandari; Subangkit; Vivi Setiawaty |
| EPI_ISL_700451, EPI_ISL_700478, EPI_ISL_700551, EPI_ISL_1040679 | Pacaltsdorp Clinic wc PAC | NHLS/UCT | Arash Iranzadeh; Bruna Galvao; Carolyn Williamson; Deelan Doolabh; Diana Hardie; Houriiyah Tegally; Innocent Mudau; Kruger Marais; Lynn Tyers; Marvin Hsiao; Stephen Korsman |
| EPI_ISL_696470, EPI_ISL_696474 | Pacaltsdorp Clinic wc PAC & NHLS/UCT | KRISP, KZN Research Innovation and Sequencing Platform | Arash Iranzadeh; Bruna Galvao; Carolyn Williamson; Deelan Doolabh; Diana Hardie; Emanuel James San; Houriiyah Tegally; Innocent Mudau; Jennifer Giandhari; Kruger Marais; Lynn Tyers; Marvin Hsiao; Stephen Korsman; Sureshnee Pillay; Tulio de Oliveira |
| EPI_ISL_7834578 | Palapye Primary Hospital Laboratory | Botswana Harvard HIV Reference Laboratory | Boitumelo Zuze; Botshelo Radibe; Dorcas Maruapula; Joseph Makhema; Keoratile Ntshambiwa; Kgomotso Moruisi; Legodile Kooepile; Mosepele Mosepele; Mphaphi B. Mbulawa; Ontlametse T. Bareng; Pamela Smith-Lawrence; Roger Shapiro; Sefetogi Ramaologa; Shahin Lockman; Sikhulile Moyo; Simani Gaseitsiwe; Thongbotho Mphoyakgosi; Wonderful T. Choga |
| EPI_ISL_1258797, EPI_ISL_1306295, EPI_ISL_1543466, EPI_ISL_1634155, EPI_ISL_1828077, EPI_ISL_2235075, EPI_ISL_2446069, EPI_ISL_2446070, EPI_ISL_4546800, EPI_ISL_5017831, EPI_ISL_5287521, EPI_ISL_6000122, EPI_ISL_7128782, EPI_ISL_7851641, EPI_ISL_8093537, EPI_ISL_8093984, EPI_ISL_8094012, EPI_ISL_8094081, EPI_ISL_8094236, EPI_ISL_8094933, EPI_ISL_8306248 | | | |
| see above | Pandemic Response Lab - NYC | Pandemic Response Lab, R&D | Alex Carpio; Cybill del Castillo; Dylan Law; Haiping Hao; Henry Lee; Isabel Fernandez Escapa; Jon Laurent; Katharine Nelson; Melissa Hopkins; Michael Hammerling; Pradeep Bugga; Shinyoung Clair Kang; Sol Rey; William Ward |
| EPI_ISL_8109026 | Pangenomics International Pvt. Ltd, Ahmedabad | Gujarat Biotechnology Research Centre | Akhilesh Modi; Apurvasinh Puvar; Bhadreshsinh Gohil; Chaitanya Joshi; Disha Vora; Janvi Raval; Jaykumar Rangani; Madhvi Joshi; Nimesh Patel; Nitin Savaliya; Nitin Shukla; Priyank Chavda; Ramesh Pandit; Roshani Mishra; Shashi Singh; Sonal Sharma; Tasnim Trivedi; Zarna Patel |
| EPI_ISL_8190648 | Pathocare Pathology laboratory,Vadodara | Gujarat Biotechnology Research Centre | Akhilesh Modi; Apurvasinh Puvar; Bhadreshsinh Gohil; Chaitanya Joshi; Disha Vora; Janvi Raval; Jaykumar Rangani; Madhvi Joshi; Mitul Vakani; Nimesh Patel; Nitin Savaliya; Nitin Shukla; Priyank Chavda; Ramesh Pandit; Roshani Mishra; Sonal Sharma; Tasnim Trivedi; Zarna Patel |
| EPI_ISL_1430426, EPI_ISL_1928171, EPI_ISL_5716246, EPI_ISL_7190236 | Pathogen Genomics Center, National Institute of Infectious Diseases | Pathogen Genomics Center, National Institute of Infectious Diseases | Kentaro Itokawa; Makoto Kuroda; Masanori Hashino; Rina Tanaka; Tsuyoshi Sekizuka |
| EPI_ISL_2397204 | Plateforme de testing Namuroise | Plateforme de testing Namuroise | Degosserie Jonathan; Denis Olivier; Lesly Nyinkeu Kemamen; Mullier François; Nicolas Gilliard; Otto Gaetan |
| EPI_ISL_2246065, EPI_ISL_2422205, EPI_ISL_4059529 | Platform BIS UZA/UAntwerpen | Labo Klinische Biologie, UZA | Basil Britto Xavier; Christine Lammens; Herman Goossens; Ines Verbesselt; Jasmine Coppens; Kathleen Holemans; Marie Le Mercier; Veerle Matheeussen |
| EPI_ISL_2447802, EPI_ISL_2447805, EPI_ISL_2447809, EPI_ISL_2447816 | Port Elizabeth Laboratory | National Institute for Communicable Diseases of the National Health Laboratory Service | Amoako DG; Bhiman JN; Ismail A; Mahlangu B; Mohale T; Ntuli N; Scheepers C |
| EPI_ISL_712070, EPI_ISL_712072, EPI_ISL_712074 | Port Elizabeth Provincial Hospital, National Health Laboratory Services, Eastern Cape, South Africa | National Institute for Communicable Diseases of the National Health Laboratory Service | Allam M; Bhiman JN; Ismail A; Mahlangu B; Mohale T; Ntuli N |
| EPI_ISL_7849096 | PrimBio Research Institute LLC | PrimBio Research Institute LLC | Diana Z. Ye; Yang Liu |
| EPI_ISL_2471711 | Procomcure Biotech Germany GmbH | Robert Koch Institute | |
| EPI_ISL_2543561 | Provincial Public Health Reference Laboratory (PPHRL) P&SHD | Provincial Public Health Reference Laboratory (PPHRL) P&SHD | ANDLEEB HANIF |
| EPI_ISL_2341693, EPI_ISL_2341724, EPI_ISL_2341790, EPI_ISL_2341869, EPI_ISL_2341871 | Pt. Jawahar Lal Nehru Memorial Medical College, Raipur | Institute of Life Sciences - INSACOG | Ajay Parida; Amol M. Kanampalliwar; Arup Ghosh; Atimukta Jha; INSACOG Consortium; Omprakash Shiriwas; Punit Prasad; Rajeeb Swain; Rupesh Dash; Safal Walia; Sana Fatma; Shifu Aggarwal; Sunil K. Raghav |
| EPI_ISL_6826086 | Public Health Authority of the Slovak Republic | Laboratory of Genomics and Bioinformatics, Comenius University Science Park | Anna Gičová; Diana Rusňáková; Jakub Styk; Jaroslav Budiš; Miroslav Böhmer; Tatiana Sedláčková; Tomáš Szemes |
| EPI_ISL_2293205 | Public Health Laboratory, Minnesota Department of Health | University of Minnesota Genomics Center | Benjamin Auch; Corbin Dirkx; Daryl M. Gohl; Jaquelyn Kuriger-Laber; John Garbe; and Chris Carlson |
| EPI_ISL_7876369, EPI_ISL_8027342, EPI_ISL_8027362 | Public Health Ontario Laboratory | Public Health Ontario Laboratory | Aimin Li; Alireza Eshaghi; Andre Villegas; Ashleigh Sullivan; Christine Frantz; Dean Maxwell; Esha Joshi; Jared Simpson; Jennifer L Guthrie; Jonathan B Gubbay; Karthikeyan Sivaraman; Lawrence Heisler; Matthew Watson; Michael CY Li; Michael Laszloffy; Nahuel Fittipaldi; Philip Banh; Richard de Borja; Samir N Patel; Sandeep Nagra; Sandra Zittermann; Sarah Teatero; Vanessa G Allen; Yao Chen; Yogi Sundaravadanam |
| EPI_ISL_1193871, EPI_ISL_1220509, EPI_ISL_1252963, EPI_ISL_1254462, EPI_ISL_1267144, EPI_ISL_1339935, EPI_ISL_1511736, EPI_ISL_1702672, EPI_ISL_1737939, EPI_ISL_1989890, EPI_ISL_2372191, EPI_ISL_2397343, EPI_ISL_2439400, EPI_ISL_2439474, EPI_ISL_2439619, EPI_ISL_2439785, EPI_ISL_2440645, EPI_ISL_2440735, EPI_ISL_2440752, EPI_ISL_2451299, EPI_ISL_2488852, EPI_ISL_2526898, EPI_ISL_2529276, EPI_ISL_4566277, EPI_ISL_5068094, EPI_ISL_5068206, EPI_ISL_5081880, EPI_ISL_5320653, EPI_ISL_5389379, EPI_ISL_5418220, EPI_ISL_5418974, EPI_ISL_5586019, EPI_ISL_5586369, EPI_ISL_5720043, EPI_ISL_5929735, EPI_ISL_5962796, EPI_ISL_6001849, EPI_ISL_6399338, EPI_ISL_6399491, EPI_ISL_6403339, EPI_ISL_6656764, EPI_ISL_6657153, EPI_ISL_6658106, EPI_ISL_6783373, EPI_ISL_7124309, EPI_ISL_7125584, EPI_ISL_7236969, EPI_ISL_7451409, EPI_ISL_7947415, EPI_ISL_7947945 | | | |
| see above | Quest Diagnostics Incorporated | Centers for Disease Control and Prevention Division of Viral Diseases, Pathogen Discovery | A. Gerasimova; A. Perez; Adrian Paskey; B. Anderson; Ben L. Rambo-Martin; Benjamin Rambo-Martin; Christopher Gulvick; Clinton Paden; Clinton R. Paden; Dakota Howard; Darlene Wagner; Dhwani Batra; Duncan MacCannell; Erisa Sula; F. Lacbawan; I. A. Shlyakhter; I. Shlyakhter; Jason Caravas; K. Livingston; K.E. Livingston; Kara Moser; Kristine Lacek; L. Bernstein; L.E. Bernstein; M. Hua; Matthew Schmerer; P. Tanpaiboon; Peter Cook; Peter W. Cook; R. Kagan; R. M. Kagan; R. Owen; R. Rolando; R. V. Rolando; S. H. Rosenthal; S. Rosenthal; Scott Sammons; Shatavia Morrison; Suxiang Tong; Tymeckia Kendall; Victoria Caban Figueroa; Y. Liu; Yvette Unoarumhi |
| EPI_ISL_1139676 | Quest Diagnostics Incorporated | Respiratory Viruses Branch, Division of Viral Diseases, Centers for Disease Control and Prevention | A. Gerasimova; A. Perez; B. Anderson; Ben L. Rambo-Martin; Clinton R. Paden; Dakota Howard; Dhwani Batra; Duncan MacCannell; F. Lacbawan; I. A. Shlyakhter; K.E. Livingston; L.E. Bernstein; M. Hua; P. Tanpaiboon; Peter W. Cook; R. M. Kagan; R. Owen; R. V. Rolando; S. H. Rosenthal; Suxiang Tong; Y. Liu |
| EPI_ISL_2450331 | RH HOSAHALLI | INSACOG-KA, NIMHANS | Ananthapadmanabha Kotambail; Anita S Desai; Anson Kunjumon George; Chetan G K; Chitra Pattabiraman; Darshan Sreenivas; Gautham Arunachal Udupi; V Ravi |
| EPI_ISL_7740812 | ROB FERREIRA LABORATORY | National Institute for Communicable Diseases of the National Health Laboratory Service | Amoako DG; Bhiman JN; Everatt J; Ismail A; Mahlangu B; Mnguni A; Mohale T; Ntuli N; Scheepers C; Wolter N |
| EPI_ISL_5365937 | Rahila Research and Reference Laboratory | LabGenetix | Madeeha Shahzad Lodhi; Malik Ihsan Ullah Khan; Muhammad Tahir Khan; Muhammad Zeeshan Anwar; Sumaira Sharif |
| EPI_ISL_8267122 | Ramathibodi Hospital | Center for Medical Genomics Ramathibodi Hospital | Anek mungaomklang; Insee Sensorn; Inthuorn Ratchatasetthakul; Kanyarat Janthima; Parsakorn Tapaopong; Poramate Jiaranai; Vichapon Tiacharoen; Wasun Chantratita |
| EPI_ISL_2249873, EPI_ISL_2319973 | Reditus Laboratories | Reditus Laboratories | Alexa Eichelberger; Cassy Philips; Joshua J. Geltz; M.S.; Ph.D.; Rex Dyer; Robert M. Sgambelluri |
| EPI_ISL_6136150 | Regional Medical Sciences Center 1 Chiang Mai | National Institute of Health, Department of Medical Sciences, Ministry of Public Health, Thailand | Archawin Rojanawiwat; Natchaya Khiadsang; Nuttida Thongpramul; Pakorn Piromtong; Pilailuk Okada; Sirikanda Wimol; Siripaporn Phuygun; Sunthareeya Waicharoen; Suratchana Mitrat; Thanutsapa Thanadachakul |
| EPI_ISL_2362433 | Rhode Island Department of Health | Infectious Disease Program, Broad Institute of Harvard and MIT | Adams, G.; Azevedo, K.; B.L.; B.W.; Bauer, M.; Birren; Carter, A.; Chaluvadi, S.; D.J.; DeRuff, K.; Gladden-Young, A.; Huard, R.; J.E.; K.J.; King, E.; Lagerborg, K.; Lemieux; Loreth, C.; Miller, A.; Normandin, E.; P.C.; Park; Pearlman, L.; Reilly, S.; Rudy, M.; Sabeti; Siddle; Tomkins-Tinch, C.; and MacInnis |
| EPI_ISL_2285243 | Rob Ferreira | National Institute for Communicable Diseases of the National | Amoako DG; Bhiman JN; Ismail A; Mahlangu B; Mohale T; Ntuli N; Scheepers C |

| | | | |
|---|---|---|---|
| | | Health Laboratory Service | |
| EPI_ISL_7042061 | Rothen Medizinische Laboratorien AG | Clinical Bacteriology, University Hospital Basel | Adrian Egli; Alfredo Mari; Fanny Wegner; Hans Hirsch; Helena MB Seth-Smith; Ingrid Steffen; Julia Bielicki; Karoline Leuzinger; Manuel Battegay; Tim Roloff |
| EPI_ISL_1969246 | Rumah Sakit Umum Daerah Palangkaraya | National Institute of Health Research and Development | Arie Ardiansyah Nugraha; Hana Apsari Pawestri; Hartanti Dian Ikawati; Kartika Dewi Puspa; Krisna Pangesti; Nelly Puspandari; Subangkit; Triyani Soekarso; Vivi Setiawaty |
| EPI_ISL_1597723 | Rush University Medical Center | Abbott | Alan Landay; Ana Olivo; Ana Vallari; Barbara Harris; Cheryl Jennings; Gavin Cloherty; Mary Hayden; Mary Rodgers; Stefan Green; Todd Meyer |
| EPI_ISL_2341728, EPI_ISL_2341856 | S.C.B. Medical College and Hospital, Cuttack | Institute of Life Sciences - INSACOG | Ajay Parida; Amol M. Kanampalliwar; Arup Ghosh; Atimukta Jha; INSACOG Consortium; Omprakash Shiriwas; Punit Prasad; Rajeeb Swain; Rupesh Dash; Safal Walia; Sana Fatma; Shifu Aggarwal; Sunil K. Raghav |
| EPI_ISL_1927404 | SARS-CoV-2 testing team, National Institute of Infectious Diseases | Pathogen Genomics Center, National Institute of Infectious Diseases | Hazuka Y Furihata; Kentaro Itokawa; Makoto Kuroda; Masanori Hashino; Masumichi Saito; Naomi Nojiri; Nozomu Hanaoka; Rina Tanaka; Sana Uchikoba; Tsuguto Fujimoto; Tsuyoshi Sekizuka |
| EPI_ISL_745157 | SAS Saldanha VPA | National Health Laboratory Service (NHLS), Tygerberg | Bronwyn Kleinhans; Eduan Wilkindon; Gert van Zyl; Houriiyah Tegally; Kayla Delaney; Susan Engelbrecht; Tulio de Oliveira; Wolfgang Preiser |
| EPI_ISL_1445272 | SERV DE VIG SANITARIA EPIDEMIO E CTRL DE ZOONOZES GUARUJA | Instituto Butantan / Mendelics | Antonio Jorge Martins; Bibiana Santos; Claudia Renata dos Santos Barros; David Schlesinger; Debora Botequio Moretti; Dimas Tadeu Covas; Elaine Cristina Marqueze; Elaine Vieira dos Santos; Erika Freitas; Evandra Strazza Rodrigues; Flavia Aburjaile; José Salvatore Leister Patané; João Paulo Kitajima; Luiz Carlos Junior de Alcantara; Maria Carolina Elias; Marta Giovanetti; Rafael dos Santos Bezerra; Raul Machado Neto; Ricardo Haddad; Rodrigo Tocantins Calado.; Sandra Coccuzzo Sampaio; Simone Kashima; Svetoslav Nanev Slavov; Vagner Fonseca; Vincent Louis Viala |
| EPI_ISL_961748 | SIESP CHIETI DRIVE IN LANCIANO | Istituto Zooprofilattico Sperimentale dell'Abruzzo e Molise "G. Caporale" | Ancora M; Calistri P; Cammà C; Curini V; Di Domenico M; Di Pasquale A; Lorusso A; Mangone I; Marcacci M; Puglia I; Rinaldi A; Savini G; Scialabba S |
| EPI_ISL_1580587 | SIESP DIPARTIMENTO DI PREVENZIONE TERAMO TERAMO(TERAMO) | Istituto Zooprofilattico Sperimentale dell'Abruzzo e Molise "G. Caporale" | Ancora M; Calistri P; Cammà C; Caporale M; Curini V; Delli Compagni E; Di Domenico M; Di Lollo Valeria; Di Pasquale A; Lorusso A; Mangone I; Marcacci M; Puglia I; Rinaldi A; Savini G; Scialabba S |
| EPI_ISL_2341727 | SLN Medical College and Hospital, Koraput | Institute of Life Sciences - INSACOG | Ajay Parida; Amol M. Kanampalliwar; Arup Ghosh; Atimukta Jha; INSACOG Consortium; Omprakash Shiriwas; Punit Prasad; Rajeeb Swain; Rupesh Dash; Safal Walia; Sana Fatma; Shifu Aggarwal; Sunil K. Raghav |
| EPI_ISL_6262602 | SURA | Gencore - Universidad de los Andes | Cristian Barrera; David González; Gabriela Ariza; Luisa Sacristan; Marcela Guevara; Marcela Mercado; Silvia Restrepo |
| EPI_ISL_1960042 | SURA | Universidad Nacional de Colombia - Laboratorio Genómico One Health | Andres F. Cardona-Rios; Carlos Franco-Muñoz; Carolina Muñoz-Arango; Celeny Ortiz; Daniel O. Maldonado-Perez; Diego A. Álvarez-Díaz; Hector Alejandro Ruiz-Moreno; Idabely Betancur Ortiz; Jorge E. Osorio; Juan P. Hernandez-Ortiz; Karl A Ciuoderis; Katherine Laiton-Donato; Laura Silvana Perez; Lina M. Hurtado; Marcela Mercado-Reyes; Maria Angélica Maya; Maria Stella López; Rita Almanza Payares; Sandra Ines Cano; Simón Villegas Velásquez |
| EPI_ISL_8020418, EPI_ISL_8266488 | SYNLAB | GIGA Medical Genomics | Bouchra Boujemla; Claire Gourzonès; Cécile Meex; Keith Durkin; Laurent Gillet; Maria Artesi; Marie-Pierre Hayette; Nadine Cambisano; Nathalie Renotte; Olivier Ek; Sébastien Bontems; Vincent Bours |
| EPI_ISL_2387829 | SYNLAB MVZ Heidelberg | Robert Koch Institute | |
| EPI_ISL_2124208, EPI_ISL_2264430, EPI_ISL_2469623, EPI_ISL_4436571, EPI_ISL_4938078, EPI_ISL_6441339, EPI_ISL_6953026, EPI_ISL_7068667, EPI_ISL_8226687 | | | |
| see above | SYNLAB MVZ Leinfelden-Echterdingen | Robert Koch Institute | |
| EPI_ISL_2116505, EPI_ISL_2126725, EPI_ISL_2267643, EPI_ISL_2315215, EPI_ISL_2386328, EPI_ISL_2386369, EPI_ISL_2386476, EPI_ISL_2387527, EPI_ISL_2387702, EPI_ISL_2389648, EPI_ISL_2389795, EPI_ISL_2469524, EPI_ISL_2469678, EPI_ISL_4223673, EPI_ISL_8247569 | | | |
| see above | SYNLAB MVZ Leverkusen | Robert Koch Institute | |
| EPI_ISL_1849712 | SYNLAB MVZ Neustadt a. d. WeinstraÃŸe | Robert Koch Institute | |
| EPI_ISL_2267692 | SYNLAB MVZ Trier | Robert Koch Institute | |
| EPI_ISL_5122798 | SYNLAB MVZ Weiden | Robert Koch Institute | |
| EPI_ISL_2178755, EPI_ISL_2341036, EPI_ISL_2341039, EPI_ISL_2443074, EPI_ISL_2443075, EPI_ISL_2443077, EPI_ISL_2444505, EPI_ISL_7172864 | | | |
| see above | Salud Digna | Instituto Nacional de Medicina Genomica | Abraham Campos-Romero; Cedro-Tanda A; Cisneros-Villanueva M; Cruz-Islas Jazmin; Escobar-Arrazola; Escobar-Arrazola MA; Garnica-Lopez Dora; Gonzalez-Barrera D; Herrera-Montalvo LA.; Hidalgo-Miranda A; Luna-Ruiz Marco; M; Mendoza-Vargas A; Moreno-Camacho José Luis; Munguia-Garza P; Orjuela-Rodríguez M; Ramirez-Vega O; Rangel-DeLeon D; Reyes-Grajeda JP; Rodriguez-Gallegos Jorge; Yair Alfaro-Mora |
| EPI_ISL_6257960 | Salud Digna, A.C | Andersen lab at Scripps Research | Abraham Garcia Gil; Jose Luis Moreno Camacho; Marco Antonio Luna Ruiz-Esparza; Miguel A. Fernandez Rojas; SEARCH Alliance with Abraham Campos Romero |
| EPI_ISL_2155673 | San Antonio City of Ilagan Hospital | Philippine Genome Center | Alethea R. de Guzman; Anna Ong-Lim; Arianne A. Zamora; Asia Louisa U. Chong; Benedict A. Maralit; Candice Francheska B. Tambaoan; Carlo M. Lapid; Celia Carlos; Devon Ray Pacial; Edsel Maurice Salvaña; El King D. Morado; Eva Maria Cutiongco-de la Paz; Francis A. Tablizo; Irish Coleen A. Asin; Jaime C. Montoya; Jan Michael C. Yap; Jo-Hannah S. Llames; John Q. Wong; Joshua Gregor A. Dizon; Juan Antonio R. Magalang; Karol Sophia Agape R. Padilla; Kenneth M. Kim; Kris P. Punayan; Marc Edsel C. Ayes; Marc Jerrone R. Castro; Maria Rosario Singh-Vergeire and Cynthia P. Saloma; Maria Sofia L. Yangzon; Marissa Alejandria; Razel Nikka M. Hao; Rianna Patricia S. Cruz; Sheila Mae M. Araiza |
| EPI_ISL_5428613, EPI_ISL_6490496, EPI_ISL_7189309, EPI_ISL_8162934 | San Diego County Public Health Laboratory | San Diego County Public Health Laboratory | Ashleigh Murphy Schafer; Brett Austin; Maria Victorio |
| EPI_ISL_2233906 | Scientific Platform Pasteur USP, University of Sao Paulo | Scientific Platform Pasteur USP, University of Sao Paulo | Cunha; M.P.; Muxel; S.M. and Minoprio, P. |
| EPI_ISL_696475 | Sedgefield Clinic wc SGE & NHLS/UCT | KRISP, KZN Research Innovation and Sequencing Platform | Arash Iranzadeh; Bruna Galvao; Carolyn Williamson; Deelan Doolabh; Diana Hardie; Emanuel James San; Houriiyah Tegally; Innocent Mudau; Jennifer Giandhari; Kruger Marais; Lynn Tyers; Marvin Hsiao; Stephen Korsman; Sureshnee Pillay; Tulio de Oliveira |
| EPI_ISL_2179709, EPI_ISL_2179750 | Servicio Microbiología Hospital La Paz | Servicio Microbiología Hospital La Paz | Elie Dahdouh; Fernando Lázaro; Jesús Mingorance Cruz; Rubén Cáceres |
| EPI_ISL_7752333 | Servicio Virosis Respiratorias-Departamento Virologia-INEI | Instituto Nacional Enfermedades Infecciosas C.G.Malbran | Avaro M.; Baumeister E.; Benedetti E.; Campos J.; Cisterna D.; Dattero ME; De Belder D.; Haim MS.; Lorenzo F.; Molina V.; Perandones C.; Poklepovich T.; Pontoriero A.; Russo M.; Sanchez Loria J.; Tuduri E. |
| EPI_ISL_2135148, EPI_ISL_2135258, EPI_ISL_2135260, EPI_ISL_2135695, EPI_ISL_2135729, EPI_ISL_2136106, EPI_ISL_2136147, EPI_ISL_2136172, EPI_ISL_2140059, EPI_ISL_2140131, EPI_ISL_2140134, EPI_ISL_2158728, EPI_ISL_2158789 | | | |
| see above | Servicio Virosis Respiratorias-Departamento Virología-INEI | Instituto Nacional Enfermedades Infecciosas C.G.Malbran | Avaro M.; Baumeister E.; Benedetti E.; Campos J.; Cisterna D.; Dattero ME; Lorenzo F.; Molina V.; Perandones C.; Poklepovich T.; Pontoriero A.; Russo M.; Tuduri E. |
| EPI_ISL_2104445, EPI_ISL_2283637 | Servicio de Microbiologia Hospital Ramon y Cajal | Servicio de Microbiologia Hospital Ramon y Cajal | Galan JC; Martinez L. Abreu M; Ponce M; y Gonzalez-Alba JM |
| EPI_ISL_1922390 | Servicio de Microbiología Clínica (Complejo Hospitalario de Navarra, Pamplona) | Centro de Secuenciación NASERTIC | Ana Miqueleiz; Ana Navascués; Carmen Ezpeleta Baquedano |
| EPI_ISL_6769199 | Servicios Medicos Olimpus | Laboratorio de Biologia Molecular, Universidad Cooperativa de Colombia, Santa Marta | Andrew S. Muñoz-Gamba; Camila Gonzalez; Daniel B. Ramírez-Osorio; Johana Zuñiga; José A. Usme-Ciro; Mayerlin Vasquez Cardenas; Paula A. Quintero-Cortés; Pricelis Paulin Polanco Fontalvo |
| EPI_ISL_6795212, EPI_ISL_6901899 | Shamir Medical Center (Asaf Harofe) | Shamir Medical Center (Asaf Harofe) | Abu Hamad Ramzia; Adina Bar Chaim; Alona Frenkel; Anna Vishnevsky; Chen Weiner; Nir Rainy; Patricia Benveniste-Lekovitz; Reut Sorek Abramovich; Yevgeni Yegorov |
| EPI_ISL_5867481, EPI_ISL_5867514, EPI_ISL_5867532 | Sharp HealthCare Laboratory | Andersen lab at Scripps Research | Art Mendoza; Cathy Woerle; Jacquelyn Berumen; Liam McGinnis; Omid Bakhtar; SEARCH Alliance San Diego with Aaron Harding |
| EPI_ISL_2438027 | Somer | Universidad Nacional de Colombia - Laboratorio Genómico One Health | Andres F. Cardona-Rios; Carlos Franco-Muñoz; Carolina Muñoz-Arango; Celeny Ortiz; Daniel O. Maldonado-Perez; Diego A. Álvarez-Díaz; Hector Alejandro Ruiz-Moreno; Idabely Betancur Ortiz; Jorge E. Osorio; Juan P. Hernandez-Ortiz; Karl A Ciuoderis; Katherine Laiton-Donato; Laura Silvana Perez; Lina M. Hurtado; Marcela Mercado-Reyes; Maria Angélica Maya; Maria Stella López; Rita Almanza Payares; Sandra Ines Cano; Simón Villegas Velásquez |
| EPI_ISL_2125310 | Sonic - Labor Staber Bayreuth | Robert Koch Institute | |
| EPI_ISL_7768808 | Sonic - MVZ Medizinisches Labor Bremen GmbH | Robert Koch Institute | |
| EPI_ISL_7870635 | South Goa District Hospital | RF-IISER Pune | Aurnab Ghose; Joy Merwin Monteiro; Krishanpal Karmodiya |
| EPI_ISL_7380295 | Southern Nevada Public Health Laboratory | Southern Nevada Public Health Laboratory | Michael Picker |
| EPI_ISL_6652189 | Stadtspital Triemli | Institute of Medical Virology | Alexandra Trkola; Annette Audigé; Catharine Aquino; Cyril Shah; Daniel Ehrsam; Gabriela Ziltener; Guido Bloemberg; Hubert Rehrauer; Isabel Stürmer; Joel Wirz; Jon Huder; Jürg Böni; Kevin Steiner; Maria Grünberg; Maryam Zaheri; Michael Huber; Riccarda Capaul; Stefan Schmutz; Verena Kufner; Weihong Qi |
| EPI_ISL_7854370 | StarMed Healthcare | UNC Charlotte Environmental Monitoring Laboratory | Cynthia Gibas; Jannatul Ferdous; Jessica Schlueter; Juan Bolanos; Kevin Lambirth; Samuel Kunkleman; Torri Weathers |
| EPI_ISL_1914584, EPI_ISL_1914588, EPI_ISL_2307108 | State Virus Research and Diagnostic Laboratory (VRDL), AIIMS Raipur | State Virus Research and Diagnostic Laboratory (VRDL), AIIMS Raipur | Anudita Bhargava; Jiyan Chandrawanshi; Kuldeep Sharma; MD Rafiiqullah Khan; Priyanka Singh; Pushpendra Singh; Sanjay Singh Negi; Somya Sharma |
| EPI_ISL_8312200 | Sweden, Vastragotaland, Unilabs | Unilabs, Eskilstuna, Sweden | Emma Arvidsson |
| EPI_ISL_1200068, EPI_ISL_2032726, EPI_ISL_2205764, EPI_ISL_2206506, EPI_ISL_2207557, EPI_ISL_2212960, EPI_ISL_2221199, EPI_ISL_2411733, EPI_ISL_2414409, EPI_ISL_2414793, EPI_ISL_2415846, EPI_ISL_2417416, EPI_ISL_5362628, EPI_ISL_6177667, EPI_ISL_6177877, EPI_ISL_6179778, EPI_ISL_6709426 | | | |
| see above | Swedish national genomic surveillance program of SARS-CoV-2 | The Public Health Agency of Sweden | Alma Brolund; Maria Lind Karlberg; Maximilian Riess; Swedish national genomic surveillance program of SARS-CoV-2 |
| EPI_ISL_6324509 | Szpital Bródnowski | 1. Tricity SARS-CoV-2 sequencing consortium: University of Gdansk, Medical University of Gdansk, Vaxican Ltd., Invicta Ltd. 2. National Institute of Public Health - National Institute of Hygiene, Warsaw, Poland | Celina Cybulska; Karolina Gackowska; Katarzyna Groth; Katarzyna Zacharczuk; Krystyna Bienkowska Szewczyk; Lukasz Rabalski; Maciej Grzybek; Maciej Kosinski; Magdalena Nowakowska; Marcin Lubocki; Małgorzata Sadkowska-Todys; Tomasz Wołkowicz |
| EPI_ISL_7949953, EPI_ISL_7950070 | Szpital Solec | 1. Tricity SARS-CoV-2 sequencing consortium: University of Gdansk, Medical University of Gdansk, Vaxican Ltd., Invicta Ltd. 2. National Institute of Public Health - National Institute of Hygiene, Warsaw, Poland | Celina Cybulska; Karolina Gackowska; Katarzyna Groth; Katarzyna Zacharczuk; Krystyna Bienkowska Szewczyk; Lukasz Rabalski; Maciej Grzybek; Maciej Kosinski; Magdalena Nowakowska; Marcin Lubocki; Małgorzata Sadkowska-Todys; Tomasz Wołkowicz |
| EPI_ISL_1935627 | TEMPUS LABS INC | Wadsworth Center, New York State Department of Health | Alexis Russell; Catharine Prussing; Daryl M. Lamson; Erasmus Schneider; Erica Lasek-Nesselquist; John Kelly; Jonathan Plitnick; Kirsten St. George; Matthew Shudt; Melissa A Leisner; Navjot Singh |
| EPI_ISL_1113569, | TGen North | TGen North | "Jolene Bowers; Ashlyn Pfeiffer; Brett Van Tassel; Chris French; Darrin Lemmer; Dave Engelthaler; Dave Engelthaler"; Hayley Yaglom; Heather Centner; Megan Folkerts; The Arizona COVID Genomics Union |

| | | | |
|---|---|---|---|
| EPI_ISL_2281712, EPI_ISL_7660611 | | | (ACGU)" |
| EPI_ISL_5900584, EPI_ISL_6252700, EPI_ISL_7709982 | TPMG Regional Laboratory | California Department of Public Health | Emily Smith on behalf of CDPH-COVIDNet and UCLA Technology Center for Genomics & Bioinformatics; Emily Smith on behalf of CDPH-COVIDNet and UCR Institute for Integrative Genome Biology; Emily Smith on behalf of CDPH-COVIDNet and UCSF Center for Advanced Technology |
| EPI_ISL_7605623, EPI_ISL_8182751 | TSHEPONG LABORATORY | National Institute for Communicable Diseases of the National Health Laboratory Service | Amoako DG; Bhiman JN; Everatt J; Ismail A; Kekana D; Mahlangu B; Mnguni A; Mohale T; Ntuli N; Scheepers C; Wolter N |
| EPI_ISL_1961124 | TXDSHS | TXDSHS | Anita Pokharel; Bonnie Oh; Chun Wang; Grace Kubin; Jenny Zhang; Lorraine Rodriguez; Maliha Rahman; Mayela Pedrueza; Myong Koag; Rachel Lee; Rashmi Tuladhar |
| EPI_ISL_7456440 | Tambo Memorial Laboratory | National Institute for Communicable Diseases of the National Health Laboratory Service | Amoako DG; Bhiman JN; Everatt J; Ismail A; Mahlangu B; Mnguni A; Mohale T; Ntuli N; Scheepers C; Wolter N |
| EPI_ISL_2342337, EPI_ISL_2342353 | Tata Main Hospital, Jharkhand | Institute of Life Sciences - INSACOG | Ajay Parida; Amol M. Kanampalliwar; Arup Ghosh; Atimukta Jha; INSACOG Consortium; Omprakash Shiriwas; Punit Prasad; Rajeeb Swain; Rupesh Dash; Safal Walia; Sana Fatma; Shifu Aggarwal; Sunil K. Raghav |
| EPI_ISL_2135033, EPI_ISL_2501624, EPI_ISL_5597436 | Texas Children's Hospital | Texas Children's Microbiome Center | Ila Singh; James Dunn; James Versalovic; Jennifer K. Spinler; Ruth Ann Luna |
| EPI_ISL_6941462, EPI_ISL_6941470 | The Caribbean Public Health Agency | Carrington Lab, Department of Preclinical Sciences, Faculty of Medical Sciences, The University of the West Indies, St Augustine Campus | Anushka Ramjag; Arianne Brown-Jordan; Avery Hinds; Christine V. F. Carrington; Christopher Oura; Gabriel Escobar; Irad Potter; Nikita S. D. Sahadeo; Nuno Faria; Oliver Pybus; Risha Singh; Sarah Hill; SueMin Nathaniel; Vernie Ramkissoon |
| EPI_ISL_8317746 | The Jackson Laboratory | The Jackson Laboratory | Adams M; Kelly K; Li L; Long E; Omerza G; Renzette N |
| EPI_ISL_1838086 | The National Centre for Cell Science | CSIR-Centre for Cellular and Molecular Biology-INSACOG | Ajay Pillai; Amareshwar Vodapalli; Ara Sreenivas; Archana Bharadwaj Siva; B Himasri; Blessy B John; Dhiraj Paul; Divya Tej Sowpati; INSACOG Consortium team; Karthik Bharadwaj Tallapaka; Lamuk Zaveri; Manoj Kumar Bhat; Mitali Inamdar; Mohak P Gujare; Onkar Kulkarni; Payel Mukherjee; Rakesh K Mishra; Sharath Chandra Thota; Shivang P. Bhanushali; Shreekant Verma; Sofia Banu; Sonal Manik Chavan; Tulasi Nagabandi; Valli Nagalakshmi Undamatla; Viswagithe S L; Yogesh Shouche |
| EPI_ISL_960139 | Thembalethu CDC wc THC | National Health Laboratory Service/UCT | Arash Iranzadeh; Bruna Galvao; Carolyn Williamson; Deelan Doolabh; Diana Hardie; Innocent Mudau; Kruger Marais; Lynn Tyers; Marvin Hsiao; Stephen Korsman |
| EPI_ISL_696500 | Thembalethu CDC wc THC & NHLS/UCT | KRISP, KZN Research Innovation and Sequencing Platform | Arash Iranzadeh; Bruna Galvao; Carolyn Williamson; Deelan Doolabh; Diana Hardie; Emanuel James San; Houriiyah Tegally; Innocent Mudau; Jennifer Giandhari; Kruger Marais; Lynn Tyers; Marvin Hsiao; Stephen Korsman; Sureshnee Pillay; Tulio de Oliveira |
| EPI_ISL_7850799 | Tulane University School of Medicine | Tulane University School of Medicine | Di Tian |
| EPI_ISL_6015848, EPI_ISL_8306804, EPI_ISL_8306836 | Tulare County Public Health Laboratory | Tulare County Public Health Laboratory | Denise L Lopez; Joshua Kootstra |
| EPI_ISL_745149, EPI_ISL_745180, EPI_ISL_745183 | Tygerberg Hospital wc TBH | National Health Laboratory Service (NHLS), Tygerberg | Bronwyn Kleinhans; Eduan Wilkindon; Gert van Zyl; Houriiyah Tegally; Kayla Delaney; Susan Engelbrecht; Tulio de Oliveira; Wolfgang Preiser |
| EPI_ISL_7316761 | UAB Diagnostikos laboratorija | Institute of Biotechnology, Life Sciences Center, Vilnius University | Albertas Timinskas; Alma Gedvilaite; Danguole Ziogiene; Emilija Vasiliunaite; Milda Norkiene |
| EPI_ISL_7709503 | UCSC Genomics Institute | UCSC Genomics Institute | A. Marm Kilpatrick; Angie Hinrichs; Beth Shapiro; Brian Young; Bryan Thornlow; Ciara Wanket; David Haussler; Eric Beraut; Hugh Olsen; Ikenna Anigbogu; Isabel Bjork; Jakob McBroome; Jeremy Sanford; Joshua Kapp; Mark Akeson; Maximilian Haeussler; Michael Stone; Miten Jain; Molly Cassatt-Johnstone; Namrita Dhillon; Russell Corbett-Detig; Terren Chang; Yatish Turakhia |
| EPI_ISL_8266573 | UMC Groningen, Clinical Virology, Department of Medical Microbiology and Infection Prevention | UMC Groningen, Clinical Virology, Department of Medical Microbiology and Infection Prevention | Alexander Friedrich; Coretta Van Leer-Buter; Erley Lizarazo-Forero; Hubert Niesters; Lilli Gard; Marjolein Knoester; Monika Fliss; Sigrid Rosema; Xuewei Zhou |
| EPI_ISL_8150842, EPI_ISL_8150897 | UMass Memorial Medical Center | Center for Microbiome Research | Doyle V. Ward; Karl J. Simon; Richard T. Ellison III; Timothy F. Kowalik |
| EPI_ISL_2136285, EPI_ISL_2136287, EPI_ISL_2136315, EPI_ISL_2136322, EPI_ISL_2321621 | UNIGEM | Universidad Nacional de Colombia - Laboratorio Genómico One Health | Andres F. Cardona-Rios; Carlos Franco-Muñoz; Carolina Muñoz-Arango; Celeny Ortiz; Daniel O. Maldonado-Perez; Diego A. Álvarez-Díaz; Hector Alejandro Ruiz-Moreno; Idabely Betancur Ortiz; Jorge E. Osorio; Juan P. Hernandez-Ortiz; Karl A Ciuoderis; Katherine Laiton-Donato; Laura Silvana Perez; Lina M. Hurtado; Marcela Mercado-Reyes; Maria Angélica Maya; Maria Stella López; Rita Almanza Payares; Sandra Ines Cano; Simón Villegas Velásquez |
| EPI_ISL_1820925, EPI_ISL_1820935 | UNIVERSIDAD DE CARTAGENA | Instituto Nacional de Salud- Dirección de Investigación en Salud Pública | Carlos Franco-Muñoz; Carmen Osorio; Christian Romero; Diana Malo; Diego A. Álvarez-Díaz; Diego Andrés Prada; Gerardo Santamaría; Hector Alejandro Ruiz-Moreno; Jhonnatan Reales-González; Jorge Rivera; Juan Camilo Martinez; Julian Naizaque; Katherine Laiton-Donato; Lisseth Pardo; Magdalena Wiesner; Marcela Mercado-Reyes; Maria T. Herrera-Sepúlveda; Marta Lopez Blanco; Martha Lucia Ospina Martinez; Paola Rojas; Patricia del Portillo; Sergio Gomez; Sheryll Corchuelo; Ángela Alarcon Cruz |
| EPI_ISL_1820926 | UNIVERSIDAD DE Magdalena | Instituto Nacional de Salud- Dirección de Investigación en Salud Pública | Carlos Franco-Muñoz; Carmen Osorio; Christian Romero; Diana Malo; Diego A. Álvarez-Díaz; Diego Andrés Prada; Gerardo Santamaría; Hector Alejandro Ruiz-Moreno; Jhonnatan Reales-González; Jorge Rivera; Juan Camilo Martinez; Julian Naizaque; Katherine Laiton-Donato; Lisseth Pardo; Magdalena Wiesner; Marcela Mercado-Reyes; Maria T. Herrera-Sepúlveda; Marta Lopez Blanco; Martha Lucia Ospina Martinez; Paola Rojas; Patricia del Portillo; Sergio Gomez; Sheryll Corchuelo; Ángela Alarcon Cruz |
| EPI_ISL_1820936 | UNIVERSIDAD DEL SINU | Instituto Nacional de Salud- Dirección de Investigación en Salud Pública | Carlos Franco-Muñoz; Carmen Osorio; Christian Romero; Diana Malo; Diego A. Álvarez-Díaz; Diego Andrés Prada; Gerardo Santamaría; Hector Alejandro Ruiz-Moreno; Jhonnatan Reales-González; Jorge Rivera; Juan Camilo Martinez; Julian Naizaque; Katherine Laiton-Donato; Lisseth Pardo; Magdalena Wiesner; Marcela Mercado-Reyes; Maria T. Herrera-Sepúlveda; Marta Lopez Blanco; Martha Lucia Ospina Martinez; Paola Rojas; Patricia del Portillo; Sergio Gomez; Sheryll Corchuelo; Ángela Alarcon Cruz |
| EPI_ISL_5403140 | UOC Patologia Clinica, Ospedale Fabrizio Spaziani, Frosinone, Italy | University Campus Bio-Medico of Rome (UCBM) | Angeletti S.; De Florio L.; Fogolari M.; Francesconi M.; Lintas C.; Riva E.; Veralli R. |
| EPI_ISL_5428014, EPI_ISL_7412973, EPI_ISL_7831238 | US Air Force School of Aerospace Medicine | US Air Force School of Aerospace Medicine | Anthony Fries; Carol Garrett; Clarise Starr; Cole Anderson; Deanna Muehleman; Elizabeth Macias; Fritz Castillo; Jennifer Black; Jennifer Meyer; Kelsey Lanter; Sarah Purves; William Gruner |
| EPI_ISL_1366175, EPI_ISL_1405247, EPI_ISL_1620982, EPI_ISL_1826132, EPI_ISL_2002340, EPI_ISL_2002342, EPI_ISL_2002836, EPI_ISL_2031294, EPI_ISL_2031505, EPI_ISL_2432613, EPI_ISL_2432699, EPI_ISL_2544979, EPI_ISL_2545184 | | | |
| see above | UW Virology Lab | UW Virology Lab | Alexander Greninger; Hong Xie; Keith R Jerome; Lasata Shrestha; Meei-Li Huang; Michelle Lin; Nathan Breit; Noah R. Baker; Patrick Mathias; Pavitra Roychoudhury; Ricardo Perez; Robert J. Livingston; Saraswathi Sathees; Sean Ellis; Shah Mohamed Bakhash; Tien V. Nguyen |
| EPI_ISL_1220045 | Universidad de Magdalena | Instituto Nacional de Salud- Dirección de Investigación en Salud Pública | Carlos Franco-Muñoz; Diego A. Álvarez-Díaz; Diego Andrés Prada; Gerardo Santamaría; Hector Alejandro Ruiz-Moreno; Jhonnatan Reales-González; Julian Naizaque; Katherine Laiton-Donato; Magdalena Wiesner; Marcela Mercado-Reyes.; Maria T. Herrera-Sepúlveda; Martha Lucia Ospina Martínez; Sheryll Corchuelo |
| EPI_ISL_1582994 | Universidad del Atlántico Laboratorio de Investigación en Biología Molecular | Instituto Nacional de Salud- Dirección de Investigación en Salud Pública | Carlos Franco-Muñoz; Carmen Osorio; Diana Malo; Diego A. Álvarez-Díaz; Diego Andrés Prada; Gerardo Santamaría; Hector Alejandro Ruiz-Moreno; Jhonnatan Reales-González; Juan Camilo Martinez; Julian Naizaque; Katherine Laiton-Donato; Lisseth Pardo; Magdalena Wiesner; Marcela Mercado-Reyes; Maria T. Herrera-Sepúlveda; Marta Lopez Blanco; Martha Lucia Ospina Martinez; Paola Rojas; Sergio Gomez; Sheryll Corchuelo; Ángela Alarcon Cruz |
| EPI_ISL_7380139 | University Clinical Hospital of Mostar, Department of Microbiology and Molecular Diagnostics | University of Sarajevo, Veterinary Faculty, Laboratory for Molecular Diagnostic and Research | Goletic S; Goletic T; Herceg I; Hodzic A; Jazic A; Nicevic M; Ostojic M; Sabic E; Skocibusic S; Softic A; Terzic I |
| EPI_ISL_6961017 | University Hospital of Northern Norway, Department for Microbiology and Infectious Disease Control | Norwegian Institute of Public Health, Department of Virology | Atiya R Ali; Debech Nadia; Engebretsen Serina Beate; Garcia Llorente Ignacio; Hilde Elshaug; Hilde Vollan; Jon Bråte; Kamilla Heddeland Instefjord; Karoline Bragstad; Kathrine Stene-Johansen; Line Victoria Moen; Marie Paulsen Madsen; Olav Hungnes; Pedersen Benedikte Nevjen; Rasmus Riis Kopperud |
| EPI_ISL_2091086 | University Hospitals Translational Laboratory (UHTL), University Hospitals | University Hospitals Translational Laboratory (UHTL), University Hospitals | Alouani, D.; Sadri, N.; Song, X. |
| EPI_ISL_2490628, EPI_ISL_6512613, EPI_ISL_7096228 | University Hospitals of Geneva, Laboratory of Virology | HUG, Laboratory of Virology and the Health2030 Genome Center | Aline Mamin; Ana Rita Goncalves; Cedric Howald; Deborah Penet; Emmanouil Dermitzakis; Francisco Perez; Henri Pegeot; Ioannis Xenarios; Keith Harshman; Laurent Kaiser; Lorenzo Cerutti; Melyssa Elies; Samuel Cordey |
| EPI_ISL_6604483 | University of Chicago Medicine | RIPHL at Rush University Medical Center | Alyse Kittner; Edith Perez; Felix Araujo Perez; Kevin Kunstman; Laura Furtado; Lauren Megger; Marieta Hyde; Nedra Love; Sofiya Bobrovska; Stefan Green |
| EPI_ISL_4548766 | University of Liège COVID-19 testing center | GIGA Medical Genomics | Bouchra Boujemla; Claire Gourzonès; Cécile Meex; Keith Durkin; Laurent Gillet; Maria Artesi; Marie-Pierre Hayette; Nadine Cambisano; Nathalie Renotte; Olivier Ek; Sébastien Bontems; Vincent Bours |
| EPI_ISL_873065, EPI_ISL_1753673, EPI_ISL_5987105, EPI_ISL_6425352, EPI_ISL_7145203 | University of Michigan Clinical Microbiology Laboratory | Lauring Lab, University of Michigan, Department of Microbiology and Immunology | Gilbert; Valesano |
| EPI_ISL_1827549, EPI_ISL_1827554, EPI_ISL_2245470 | University of Wisconsin-Madison AIDS Vaccine Research Laboratories | University of Wisconsin-Madison AIDS Vaccine Research Laboratories | Gage Moreno; Katarina Braun; et al. AIDS Vaccine Research Laboratories |
| EPI_ISL_977352 | University of Zambia, School of Veterinary Medicine | UNZAVET and PATH | Daniel Bridges; Mulenga Mwenda-Chimfwembe; Ngonda Saasa |
| EPI_ISL_1570323 | UniversitÃ¤tsklinikum Heidelberg | Robert Koch Institute | |
| EPI_ISL_7765829 | Universitätsklinikum Heidelberg | Robert Koch Institute | |
| EPI_ISL_1364891 | Universitätsklinikum Knappschaftskrankenhaus Bochum | Bundeswehr Institute of Microbiology | Alexandra Rehn; Enrico Georgi; Malena Bestehorn-Willmann; Markus Antwerpen; Mathias Walter; Mike Pillukat; Roman Wölfel; Sabine Zange |
| EPI_ISL_5907107, EPI_ISL_5907165 | Universitätsmedizin Mannheim | Robert Koch Institute | |

| | | | |
|---|---|---|---|
| EPI_ISL_7924595 | Uniwersyteckie Centrum Kliniczne | 1. Tricity SARS-CoV-2 sequencing consortium: University of Gdansk, Medical University of Gdansk, Vaxican Ltd., Invicta Ltd. 2. National Institute of Public Health - National Institute of Hygiene, Warsaw, Poland | Celina Cybulska; Karolina Gackowska; Katarzyna Groth; Katarzyna Zacharczuk; Krystyna Bienkowska Szewczyk; Lukasz Rabalski; Maciej Grzybek; Maciej Kosinski; Magdalena Nowakowska; Marcin Lubocki; Małgorzata Sadkowska-Todys; Tomasz Wołkowicz |
| EPI_ISL_2291791, EPI_ISL_4176377, EPI_ISL_7742672 | Utah Public Health Laboratory | Utah Public Health Laboratory | Erin L. Young; John Arnn; Kelly F. Oakeson; Olinto Linares-Perdomo; Pooja Gupta; Tara Gallagher; Tom Iverson |
| EPI_ISL_7971547 | VERMAAK LABORATORY | National Institute for Communicable Diseases of the National Health Laboratory Service | Amoako DG; Bhiman JN; Everatt J; Ismail A; Kekana D; Mahlangu B; Mnguni A; Mohale T; Ntuli N; Scheepers C; Wolter N |
| EPI_ISL_1445235 | VIGILANCIA EPIDEMIOLOGICA | Instituto Butantan / Mendelics | Antonio Jorge Martins; Bibiana Santos; Claudia Renata dos Santos Barros; David Schlesinger; Debora Botequio Moretti; Dimas Tadeu Covas; Elaine Cristina Marqueze; Elaine Vieira dos Santos; Erika Freitas; Evandra Strazza Rodrigues; Flavia Aburjaile; José Salvatore Leister Patané; João Paulo Kitajima; Luiz Carlos Junior de Alcantara; Maria Carolina Elias; Marta Giovanetti; Rafael dos Santos Bezerra; Raul Machado Neto; Ricardo Haddad; Rodrigo Tocantins Calado.; Sandra Coccuzzo Sampaio; Simone Kashima; Svetoslav Nanev Slavov; Vagner Fonseca; Vincent Louis Viala |
| EPI_ISL_2493171 | VIGILANCIA EPIDEMIOLOGICA DE IBATE | Instituto Butantan | Antonio Jorge Martins; Claudia Renata dos Santos Barros; David Schlesinger; Debora Botequio Moretti; Dimas Tadeu Covas; Elaine Cristina Marqueze; Elaine Vieira Santos; Evandra Strazza Rodrigues; Heidge Fukumasu; Jayme Augusto de Souza-Neto; José Salvatore Leister Patané; Luiz Alcantara; Luiz Lehmann Coutinho; Maria Carolina Elias; Maurício Lacerda Nogueira; Rafael dos Santos Bezerra; Raul Machado Neto; Rejane Maria Tommasini Grotto; Ricardo Haddad; Sandra Coccuzzo Sampaio Vessoni; Simone Kashima; Svetoslav Nanev Slavov; Vincent Louis Viala |
| EPI_ISL_2210288 | VIGILANCIA EPIDEMIOLOGICA DE IBATE | Instituto Butantan / Mendelics | Antonio Jorge Martins; Claudia Renata dos Santos Barros; David Schlesinger; Debora Botequio Moretti; Dimas Tadeu Covas; Elaine Cristina Marqueze; Elaine Vieira Santos; Evandra Strazza Rodrigues; Heidge Fukumasu; Jayme Augusto de Souza-Neto; José Salvatore Leister Patané; Luiz Alcantara; Luiz Lehmann Coutinho; Maria Carolina Elias; Maurício Lacerda Nogueira; Rafael dos Santos Bezerra; Raul Machado Neto; Rejane Maria Tommasini Grotto; Ricardo Haddad; Sandra Coccuzzo Sampaio Vessoni; Simone Kashima; Svetoslav Nanev Slavov; Vincent Louis Viala |
| EPI_ISL_7133387, EPI_ISL_7133418 | VRDL Government Medical College,Patiala Punjab | NIV Influenza | Dr.Rupinder Bakshi |
| EPI_ISL_2001193 | VRDL, Goverment Medical College (GMC), Surat | Gujarat Biotechnology Research Centre | Chaitanya Joshi; Dinesh Kumar; Janvi Raval; Madhvi Joshi; Neeta Khandelwal; Nitesh Shah; Nitin Savaliya; Ramesh Pandit; Sonal Sharma; Twinkle Soni; Umang Mishra; Zarna Patel; Zuber Saiyed |
| EPI_ISL_2360348, EPI_ISL_2360350, EPI_ISL_2360367, EPI_ISL_2360390, EPI_ISL_2360416, EPI_ISL_2360597, EPI_ISL_2360635, EPI_ISL_2360759, EPI_ISL_2360817, EPI_ISL_2360854, EPI_ISL_2360877, EPI_ISL_2360916, EPI_ISL_2360926, EPI_ISL_2360932, EPI_ISL_2360948, EPI_ISL_2360952, EPI_ISL_2360980, EPI_ISL_2360987 | | | |
| see above | Vaccines and Infectious Diseases Analytics Research Unit (VIDA) | KRISP, KZn Research Innovation and Sequencing Platform | Baillie Vicky; Giandhari Jennifer; Madhi Shabir; Naidoo Yeshnee; Pillay Sureshnee; San James; Tegally Houriiyah; Wilkinson Eduan; de Oliveira Tulio; du Plessis Jeanine |
| EPI_ISL_940855, EPI_ISL_940879, EPI_ISL_940880 | Vaccines and Infectious Diseases Analytics Research Unit (VIDA) | KRISP, KZN Research Innovation and Sequencing Platform | Baillie Vicky; Giandhari Jennifer; Madhi Shabir; Naidoo Yeshnee; Pillay Sureshnee; Tegally Houriiyah; de Oliveira Tulio; du Plessis Jeanine |
| EPI_ISL_1132711, EPI_ISL_1132715 | Vaccines and Infectious Diseases Analytics Research Unit (VIDA) | KRISP, KZn Research Innovation and Sequencing Platform | Baillie Vicky; Giandhari Jennifer; Madhi Shabir; Naidoo Yeshnee; Pillay Sureshnee; Tegally Houriiyah; de Oliveira Tulio; du Plessis Jeanine |
| EPI_ISL_2521368, EPI_ISL_7693665 | Valais Hospital, Central Institute | Valais Hospital, Central Institute | Alexis Dumoulin; Cedric Howald; Deborah Penet; Emmanouil Dermitzakis; Henri Pegeot; Ioannis Xenarios; Keith Harshman; Lorenzo Cerutti; Melyssa Elies |
| EPI_ISL_1662284 | Veer Surendra Sai Institute of Medical Sciences and Research, Burla, Sambalpur | Institute of Life Sciences - INSACOG | Ajay Parida; Amol M. Kanampalliwar; Arup Ghosh; Atimukta Jha; INSACOG Consortium; Punit Prasad; Rajeeb Swain; Rupesh Dash; Safal Walia; Shifu Aggarwal; Sunil K. Raghav |
| EPI_ISL_2428123, EPI_ISL_2428174 | Viesoji istaiga Klaipedos universitetine ligonine | National Public Health Surveillance Laboratory | Ana Steponkiene; Danas Baksa; Jelena Razmuk; Lukas Vasionis; Lukas Zemaitis; Migle Gabrielaite; Svajune Muralyte |
| EPI_ISL_737488, EPI_ISL_899751, EPI_ISL_1129363, EPI_ISL_2152117, EPI_ISL_2308041, EPI_ISL_2544315, EPI_ISL_2544321, EPI_ISL_2544460, EPI_ISL_2544462, EPI_ISL_6720837, EPI_ISL_7372061 | | | |
| see above | Viollier AG | Department of Biosystems Science and Engineering, ETH Zürich | Andrea Patrignani; Andreia Cabral de Gouvea; Catharine Aquino; Chaoran Chen; Christian Beisel; Christiane Beckmann; Christoph Noppen; Daniel Ehrsam; David Dreifuss; Deborah Penet; Doris Popovic; Elodie Burcklen; Emmanouil Dermitzakis; Griffin White; Henri Pegeot; Ina Nissen; Ioannis Xenarios; Isabel Stürmer; Ivan Topolsky; Jay Tracy; Katharina Jahn; Keith Harshman; Kim Philipp Jablonski; Lara Fuhrmann; Laura Neff; Lennart Opitz; Lorenzo Cerutti; Louis du Plessis; Maria Domenica Moccia; Maurice Redondo; Mirjam Feldkamp; Natascha Santacroce; Niko Beerenwinkel; Noemie Santamaria de Souza; Olivier Kobel; Philipp Jablonski; Ralph Schlapbach; Rebecca Denes; Sarah Nadeau; Simon Grüter; Sophie Seidel; Tanja Stadler; Timothy Sykes |
| EPI_ISL_2344462 | Virginia Division of Consolidated Laboratory Services | Virginia Division of Consolidated Laboratory Services | Virginia DCLS |
| EPI_ISL_4503229, EPI_ISL_4748028, EPI_ISL_4748048 | Virology Unit, Institut Pasteur du Cambodge | Virology Unit, Institut Pasteur du Cambodge | Cecile Troupin; Chau Darapheak; Chin Savuth; Erik A Karlsson; Jurre Y Siegers; Kraing Sidonn; Leakhena Pum; Ly Sovann; Sophoannadedh Rath; Veasna Duong; Yi Sengdoeurn |
| EPI_ISL_745146 | Vredendal Hospital wc VRE | National Health Laboratory Service (NHLS), Tygerberg | Bronwyn Kleinhans; Eduan Wilkindon; Gert van Zyl; Houriiyah Tegally; Kayla Delaney; Susan Engelbrecht; Tulio de Oliveira; Wolfgang Preiser |
| EPI_ISL_2247027 | WSSE w Krakowie | 1. Academic Center for Pathomorphological and Genetic-Molecular Diagnostics ltd, Bialystok, Poland 2. National Institute of Public Health - National Institute of Hygiene, Warsaw, Poland | Anetta Sulewska; Jacek Nikliński; Janusz Dzięcioł; Joanna Kiśluk; Katarzyna Zacharczuk; Konrad Raczkowski; Magdalena Nowakowska; Małgorzata Sadkowska-Todys; Piotr Karabowicz; Piotr Majewski; Przemysław Biecek. Joanna Reszeć; Radosław Charkiewicz; Tomasz Wołkowicz |
| EPI_ISL_5509534 | WSSE w Olsztynie | Wojewodzka Stacja Sanitarno-Epidemiologiczna w Olsztynie, Laboratorium Badan Epidemiologiczno-Klinicznych | Aleksandra Kobiatko; Barbara Dolinska; Emilia Tarabasz; Ewa Liszewska; Marta Lukian; Monika Czerminska; Patryk Bielecki; Paulina Rozycka; Sylwia Krzetowska; Tomasz Jakubczak |
| EPI_ISL_2204896 | WV Labs | WVU and Marshall University Combined Genomics Core Facilities | James Denvir; Peter Perrotta; Peter Stoilov; Ryan Percifield; Wesley Kimble |
| EPI_ISL_5771409 | Wakayama Prefectural Research Center of Environment and Public Health | Wakayama Prefectural Research Center of Environment and Public Health | Akari Kajino; Fumio Terasoma; Hideyuki Yanai; Kentaro Itokawa; Makoto Kuroda; Masanori Hashino; Rina Tanaka; Tsuyoshi Sekizuka; Yasuyuki Hujimoto; Yosuke Hamajima |
| EPI_ISL_7899139, EPI_ISL_7899225 | Wake Forest School of Medicine, Internal Medicine | Wake Forest School of Medicine, Internal Medicine | Gregory Hawkins; Kimberly Reeves |
| EPI_ISL_7041044 | Weill Cornell Medicine | New York Genome Center | Amy Baldwin; Andre Corvelo; Arryn Craney; Chris Mason; Dayna M. Oschwald; Hanna Rennert; Lars F Westblade; Margaret Elizabeth Ross; Melissa Cushing; Michael Zody; Olivier Elemento; Priya Velu; Samantha Fennessey; Tom Maniatis |
| EPI_ISL_2343271, EPI_ISL_5392918, EPI_ISL_5393305 | Willis-Knighton Medical Center Hospital Laboratory | LSUHS Emerging Viral Threat Laboratory | Adrian Almodovar; Alexander Mijalis; Andrew D. Yurochko; April N. Johnson; Christopher G. Kevil; Gregory L. Ware; Jennifer L. Carroll; Jeremy P. Kamil; John A. Vanchiere; Joseph A. Bocchini; Krista Queen; Lorie M. Atkins; Maarten Van Diest; Rona S. Scott |
| EPI_ISL_2018547 | Wisconsin State Laboratory of Hygiene Communicable Disease Division | Wisconsin State Laboratory of Hygiene Communicable Disease Division | Abigail C. Shockey; Alicia J. Mooney; Erika M. Hanson; Kelsey R. Florek; Richard Griesser; Sara Wagner; Tonya Danz |
| EPI_ISL_2350856, EPI_ISL_2500298 | Wojewodzka Stacja Sanitarno-Epidemiologiczna w Olsztynie, Laboratorium Badan Epidemiologiczno-Klinicznych | Wojewodzka Stacja Sanitarno-Epidemiologiczna w Olsztynie, Laboratorium Badan Epidemiologiczno-Klinicznych | Aleksandra Kobiatko; Barbara Dolinska; Emilia Tarabasz; Ewa Liszewska; Marta Lukian; Monika Czerminska; Patryk Bielecki; Paulina Rozycka; Sylwia Krzetowska; Tomasz Jakubczak |
| EPI_ISL_888938, EPI_ISL_2444763 | Wyoming Public Health Laboratory | Wyoming Public Health Laboratory | Ashley Norberg; Bailey Bowcutt; Brian Dominguez; Brittany Oher; Cari Sloma; Channing Weber; Chayse Rowley; Elliot Thomasson; Jim Mildenberger; Lynette Gumbleton; Marley Goetz; Noah Hull; Sam Britz; Taylor Fearing; and Rob Christensen; and Wanda Manley |
| EPI_ISL_1447603, EPI_ISL_5644840, EPI_ISL_5645262, EPI_ISL_6173741, EPI_ISL_7674042 | Yale Clinical Virology Lab | Grubaugh Lab - Yale School of Public Health | Anderson Brito; Annie Watkins; Chaney Kalinich; Chantal Vogels; Isabel Ott; Jessica Rothman; Joseph Fauver; Kendall Billig; Mallery Breban; Marie L. Landry; Mary Petrone; Nathan Grubaugh; Tara Alpert; Tobias Koch |
| EPI_ISL_2020812 | Yale Clinical Virology Lab | Yale Center for Genomic Analysis | Brooke Sullivan; Curt Scharfe; Irina Tikhonova; Kaya Bilguvar; Shrikant Mane |
| EPI_ISL_5330267, EPI_ISL_5849308 | Yale Pathology Labs | Yale Pathology Labs | Angelique Levi; Brian Daley; Chen Liu; Guangxiao Yang; Jianhui Wang; Jinglan Wang; John Sinard; Katherine Fajardo; Kevin Schofield; Minghao Zhong; Monica Talmor; Pei Hui; Peter Gershkovich; Richard Bouffard; Stephanie Weirsman; Susan Bell; Sylvia White |
| EPI_ISL_6795838, EPI_ISL_6795842, EPI_ISL_6795850, EPI_ISL_6825390, EPI_ISL_6825392, EPI_ISL_6825397, EPI_ISL_7015178, EPI_ISL_7015180, EPI_ISL_7015181, EPI_ISL_7015204, EPI_ISL_7015207, EPI_ISL_7015208, EPI_ISL_8128438 | | | |
| see above | ZARV/NHLS, Department Medical Virology, University of Pretoria | CERI, Centre for Epidemic Response and Innvoation, Stellenbosch University and KRISP, KZN Research Innovation and Sequencing Platform, UKZN. | Adriano Mendes; Amoaka D; Amy Strydom; Arisha Maharaj; Bester P; Bhiman J; Engelbrecht S; Everatt J; Giandhari J; Goedhals D; Hardie D; Hsiao M; Iranzadeh A; Lessells R; Makatini Z; Maponga T; Mdlalose N; Micheala Davids; Mlisana K; Moir M; NGS-SA (Scheepers C; Naidoo Y; Nyaga M) Giandhari J; Oluwakemi M; Pillay S; Preiser W; Ramphal U; Ramphal Y; San JE; Sim Mayaphi and Marietjie Venter; Tegally H; Tshiabuila D; Venter M; Wilkinson E; Williamson C; de Oliveira T; von Gottberg A |
| EPI_ISL_6329340 | ZLC Zentrallabor Chur | Microbiology | Dominique Fabien Hilti; Faina Wehrli; Lorenz Risch; Martin Risch; Nadia Wohlwend; Sinem Kas; Thomas Bodmer |
| EPI_ISL_8038985 | ZOTZ KLIMAS MVZ Düsseldorf-Centrum GbR ÜBAG für Labormedizin, Genetik, Zytologie, Pathologie | Center of Medical Microbiology, Virology, and Hospital Hygiene, University of Duesseldorf | Alexander Dilthey; Andreas Walker; Daniel Strelow; Jessica Nicolai; Jörg Timm; Katrin Hoffmann; Klaus Pfeffer; Lisanna Hülse; Malte Kohns Vasconcelos; Maximilian Damagnez; Nadine Lübke; Patrick Finzer; Rainer Zotz; Tobias Wienemann; Torsten Houwaart |
| EPI_ISL_7880377 | Zavod za javno zdravstvo Dubrovačko-Neretvanske županije | Hrvatski zavod za javno zdravstvo | Anita Jurić; Dragan Jurić; Irena Tabain; Ivana Ferenčak; Josipa Kuzle |
| EPI_ISL_6031057 | Zavod za javno zdravstvo Međimurske županije | Hrvatski zavod za javno zdravstvo | Anita Jurić; Dragan Jurić; Irena Tabain; Ivana Ferenčak; Josipa Kuzle |
| EPI_ISL_6023455 | Zavod za javno zdravstvo Osječko-Baranjske županije | Hrvatski zavod za javno zdravstvo | Anita Jurić; Dragan Jurić; Irena Tabain; Ivana Ferenčak; Josipa Kuzle |
| EPI_ISL_5393693 | Zavod za javno zdravstvo Zagrebačke županije | Hrvatski zavod za javno zdravstvo | Irena Tabain; Ivana Ferenčak |
| EPI_ISL_8108836 | Zdravotní ústav Ústí nad Labem | Institute of Medical Microbiology and Virology, University Hospital Carl Gustav Carus, TU Dresden | Alexa Laubner; Alexander Dalpke; Anett Zabzinski; Eva Patrasová; Fabian Rost; Grit Mehnert; Ivana Stiborová; Jitka Pohořská; Lenka Šimůnková; Marlena Stadtmüller; Montserrat Palau de Miquel; Romana Mikešová; Susanne Reinhardt; Sylke Winkler; Sylvia Klemroth |
| EPI_ISL_2470883 | amedes Göttingen | Robert Koch Institute | |
| EPI_ISL_6952622 | amedes MVZ Halle/Leipzig | Robert Koch Institute | |
| EPI_ISL_7423744 | amedes MVZ Hannover | Robert Koch Institute | |

| | | | |
|---|---|---|---|
| EPI_ISL_1565242 | amedes MVZ fÃ¼r Laboratoriumsdiagnostik Raubling GmbH | Robert Koch Institute | |
| EPI_ISL_2470945 | amedes MVZ für Labordiagnostik Rhein-Main | Robert Koch Institute | |
| EPI_ISL_2491648 | center de prélévement COVID ISSOIRE | CHU Clermont-Ferrand, service de virologie | Bisseux Maxime; Combes Patricia; Henquell Cécile; Mirand Audrey |
| EPI_ISL_2263159, EPI_ISL_2469808, EPI_ISL_5665900 | labopart - Medizinische Laboratorien Dresden | Robert Koch Institute | |
| EPI_ISL_2151686 | laboratoire Belle Epine | Department of Virology, Henri Mondor University Hospital, Assistance Publique Hôpitaux de Paris, Université Paris-Est Créteil, INSERM U955 | Alexandre Soulier; Christophe Rodriguez; Elisabeth Trawinski; Guillaume Gricourt; Jean-Michel Pawlotsky; Melissa N'Debi; Slim Fourati; Vanessa Demontant |
| EPI_ISL_7760559, EPI_ISL_7760663 | nordlab - Partnerschaftspraxis für Laboratoriumsmedizin | Robert Koch Institute | |
